# Ancestral and derived feeding morphology during rapid ecological divergence within a species

**DOI:** 10.64898/2026.11.19.764305

**Authors:** Guðbjörg Ósk Jónsdóttir, Kenedy Annejulee Williams, Nína Guðrún Baldursdóttir, Sarah Elizabeth Steele, Arnar Pálsson

## Abstract

Derived characters that may have emerged via positive selection can be identified by comparing species, but also possibly subgroups within species. Allopatric and sympatric polymorphism observed in many salmonids is thought to emerge mainly in landlocked populations, which have lost the anadromous lifestyle of their ancestors. Populations of Arctic charr (*Salvelinus alpinus*) are an example of such polymorphism, commonly diverging along a pelagic-benthic axis. Populations of Icelandic charr can be anadromous or landlocked, like in Lake Þingvallavatn where four morphs coexist differing in habitat use, preferred prey and external feeding morphology. Three of these morphs are also caught in the neighboring Lake Úlfljótsvatn, despite prey availability in the two lakes differing notably. We studied variation of craniofacial morphology in Icelandic charr populations and compared the ecologically specialized sympatric morphs to five anadromous charr populations and to two non-specialized lake populations, as well as across two connected lakes. The results indicate minor variation among anadromous populations and that the benthic lake forms are derived. There was clear phenotypic difference between benthic and pelagic morphs within Þingvallavatn but no differences between the morphs in Úlfljótsvatn. Patterns of variation between corresponding morphs across the two lakes suggest genetics separation and phenotypic plasticity.

## Introduction

New adaptive traits and species can arise jointly when natural selection on populations in distinct environments drives the evolution of particular traits, and eventually reproductive barriers (Schluter, 2000). This can occur in allopatric, sympatric and parapatric geographical contexts (Rundle & Schluter, 2004) but a key driver is ecological speciation matching organisms to specific environment(s). This form of speciation is best viewed as a continuum, ranging from little or no divergence to complete divergence between groups (Hendry, 2017), that can shift along this continuum due to altered conditions while reproductive isolation is incomplete. This is driven by adaptive divergence, that is adaptations of different populations to their distinct environments (Hendry, 2009, 2017; Räsänen & Hendry, 2008; Rundle & Nosil, 2005). Generally, it is assumed that during adaptive divergence individuals have higher fitness in their home environment (which they have adapted to) in comparison with individuals migrating in from other locations (Hendry & Gonzalez, 2008; Kawecki & Ebert, 2004; Schluter, 2000). How common local adaptations are in nature is debated (Hendry & Gonzalez, 2008), though meta-analyses suggest this to be rather common while maladapted populations are also reported (Fraser et al., 2011; Hereford, 2009; Leimu & Fischer, 2008). One meta-analysis on salmonids estimated that the frequency of local adaptation was around 50% to 70% (Fraser et al., 2011). On average, local populations had 1.20 times higher fitness compared to non-local, though with considerable variability (Fraser et al., 2011). Definition of groups or populations as ancestral vs. derived can help identify if natural selection favored specific traits during local adaptation.

Intraspecific phenotypic variation stems from variations in genetic and environmental factors, their interplay and chance (Hendry, 2017; Kawecki & Ebert, 2004; Schwander & Leimar, 2011). Phenotypic variation in complex traits is often influenced by a genetic architecture including alleles of small to moderate effects, across many genes (Hendry, 2017). Environment can affect phenotype,_phenotypic plasticity is defined as the ability of a genotype to produce different phenotypes in response to varying environmental cues (Bradshaw, 1965; West-Eberhard, 2003). Phenotypic plasticity can either be adaptive or non-adaptive, and may evolve just as trait means (Ghalambor et al., 2007). Plasticity in novel environments may aid adaptive evolution (West-Eberhard, 1989, 2003) supported by meta-analyses on transplants experiments (Stamp & Hadfield, 2020).

For complex anatomies of animals, it is often hard to identify which traits represent adaptations and which are mere “spandrels” or secondary features not directly influenced by positive selection. We use feeding morphology of freshwater fish in the Northern Hemisphere to study seemingly adaptive divergence within species and to explore phenotypic plasticity in the wild (Parsons et al., 2011; Snorrason & Skúlason, 2004). After the last glacial retreat, many freshwater habitats opened and were colonized by anadromous fish species. Due to the minor interspecific competition and diversity of unutilized niches (Snorrason & Skúlason, 2004), resource polymorphisms arose in many systems (Smith & Skúlason, 1996; Snorrason & Skúlason, 2004). We leverage the variation within and between Icelandic populations of Arctic charr (*Salvelinus alpinus*) and focus on their craniofacial variation as it aligns with feeding specializations and may have adaptive value. Teleost are thought to respond readily to changes in environmental factors, in part due to their developmental flexibility in body size and maturation as well as the great number of “independent” skeletal elements (Gidmark et al., 2019). Arctic charr is known for extensive polymorphism through its geographic distribution (Gíslason et al., 1999; Jónsdóttir et al., 2024; Klemetsen, 2010; Kurta et al., 2025; Noakes, 2008; Woods et al., 2013) and rearing experiments have demonstrated that charr has developmental plasticity in form and behavior in response to feeding on pelagic vs benthic food (Dellinger et al., 2023; Küttner et al., 2013; Parsons et al., 2011).

Anadromous Arctic charr colonized Iceland following the last glacial retreat approximately <11,000 years ago (Saemundsson, 1992; Snorrason & Skúlason, 2004) and subsequently dispersed around the island and became isolated in some areas (Brachmann et al., 2021; Gíslason et al., 1999; Kapralova et al., 2011). Now Arctic charr is found in lakes and rivers all around Iceland, both as anadromous (putative ancestral form) or non-anadromous (landlocked) (Brachmann et al., 2021; Gíslason et al., 1999; Sandlund et al., 1992). Some waterbodies have monomorphic populations (i.e. have only one morph) while others are polymorphic (i.e. have two or more morphs). Generally, it is assumed that phenotypic and ecological variation in anadromous Arctic charr (and in salmonids in general) is low relative to derived morphs, but this is rarely tested (Esin et al., 2018; Kristjánsson et al., 2012; Sandlund et al., 1992). Population genetics have documented repeated loss of anadromy in Arctic charr in Canada and repeated colonization in Europe and Asia (Jacobs et al., 2020; Salisbury et al., 2022). Unpublished research indicates Icelandic anadromous Arctic charr split into at least four genetic subgroups (Xiao et al., Unpublished).

Landlocked charr have repeatedly diverged along the benthic-pelagic axis in multiple lakes and springs resulting in similar morph pairs or groups in Iceland (Brachmann et al., 2021; Gíslason et al., 1999; Kapralova et al., 2011; Sandlund et al., 1992). Sympatric morphs show substantial phenotypic variation and, in some cases differ genetically, but others appear influenced by plasticity (Brachmann et al., 2021; Gíslason et al., 1999; Kurta et al., 2025). The most dramatic example of polymorphism in Iceland are the four morphs in lake Þingvallavatn (Sandlund et al., 1992; Snorrason et al., 1989). They differ in adult size, habitat use, spawning time, life history, diet, morphology (Jonsson et al., 1988; Jónsdóttir et al., 2024; Malmquist, 1992; Malmquist et al., 1985; Sandlund et al., 1992; Sandlund et al., 1987; Skúlason, Snorrason, et al., 1989; Snorrason et al., 1989) but curiously only split into three genetic clusters (Guðbrandsson et al., 2019; Xiao et al., 2026). These morphs most likely diverged after a single colonization (Xiao et al., 2026). It was postulated that high habitat heterogeneity in the lake caused rapid evolution of these morphs - driven by disruptive selection on traits associated with their distinct feeding resources (Brachmann et al., 2021; Smith & Skúlason, 1996). Two of the morphs primarily feed on benthic invertebrates (large benthivorous (LB-) and small benthivorous (SB-) charr) while the other two morphs primarily capture prey in the water column (planktivorous (PL-) and piscivorous (PI-) charr) (Malmquist, 1992; Malmquist et al., 1985). Importantly for our study, these morphs are also found in the smaller and shallower neighboring lake Úlfljótsvatn (Jóhannsson & Jónsson, 2000; Woods et al., 2012). From Þingvallavatn a river, Efra Sog, drains into lake Úlfljótsvatn, which has (Aðalsteinsson et al., 1989) far less habitat diversity (Jóhannsson & Jónsson, 2000; Jónsson & Jóhannsson, 2024). Currently only downstream migration occurs for fish, but upstream migration was possible before the construction of a hydroelectric power station in 1959 at the outlet of Þingvallavatn (Jónasson, 1992). All four Þingvallavatn morphs can be found in Úlfljótsvatn, though SB-charr is very rare (Jóhannsson & Jónsson, 2000; Jónsson & Jóhannsson, 2024). Xiao et al. (2026) found that the LB- and PI-morphs in Úlfljótsvatn differ genetically from the LB- and PI-morphs in Þingvallavatn. In contrast, the PL-charr in the two lakes were genetically inseparable (Xiao et al., 2026). We will refer to the same morphs in the two lakes as “sister” morphs, for simplification. In head shape, the PL-<u>“</u>sister<u>”</u> morphs showed little separation, while the PI- and LB-<u>“</u>sister<u>”</u> morphs differed more notably between the two lakes (Xiao et al., 2026). This system allows us to gauge phenotypic plasticity in nature, specifically in the PL-<u>“</u>sister<u>”</u> morphs that are genetically similar but use different prey resources.

Here we aimed to characterize ancestral and derived feeding related traits (jaw bone shape) in Arctic charr. We examined several populations of anadromous Arctic charr, representing a putative ancestral form, and among charr in lakes that are either monomorphic or polymorphic. Anadromous charr and the charr in monomorphic landlocked lakes are considered to have a piscivore/pelagic lifestyle, but in lakes with two or more morphs (the Þingvallavatn/Úlfljótsvatn system), pelagic and benthic morphs are found and the latter hypothesized to be derived forms. We compared these sympatric morphs within and between lakes to test if and how they differ from the anadromous and monomorphic populations. The research questions are:

1. Are there differences between anadromous charr populations in bone size and/or shape? (how variable are the putative ancestral populations?)
2. Which bone size and shape traits are derived in the specialized lake morphs? (which traits have evolved most, relative to the ancestral phenotypes?)
3. Are there differences in bone size or shape between “sister” morphs in the Þingvallavatn and Úlfljótsvatn system? (Is there evidence of phenotypic plasticity in specific traits, that associate with resource use?)

## Material and methods

### Data collection and measurements

Sampling and killing of wild fishes in Iceland, requires a permit from the Directorate of Fisheries in Iceland (https://island.is/s/fiskistofa). Two authors of this study (XXX and XXX) which oversaw all sampling efforts have such permits (#0460/2021-2.0 and #2025-05-30-0558). Sexually mature Arctic charr were collected from nine different waterbodies, five rivers and four lakes (Figure 1A). Anadromous Arctic charr (AN-charr) were collected from five different rivers (Vífilsdalsá, Lambeyrarkvísl, Víðidalsá, Gufudaslá and Brúará) around Iceland using electrofishing, seine nets and angling (fish euthanized with single blow to head) during the falls of 2023 and 2024. When sampling from Vífilsdalsá, Lambeyrarkvísl and Víðidalsá mature males and females were squeezed for eggs and sperm which were fertilized and deposited into the riverbeds as a countermeasure to protect the local stocks. Charr in lakes were collected using composite nets laid overnight (fish died in net), seine nets and angling (fish euthanized with single blow to head) from four different lakes in Iceland. Two of those lakes are monomorphic (Hestvatn and Hítarvatn) and two are polymorphic (Þingvallavatn, four morphs and Úlfljótsvatn, three morphs). Data from lake Þingvallavatn was collected earlier (Jónsdóttir et al., 2024). All other lake populations (Úlfljótsvatn, Hestvatn and Hítarvatn) were sampled in the summer/fall of 2024. Older samples from Hestvatn, collected in the summer of 2011 were added to increase the sample size of that population.

**Figure 1:**
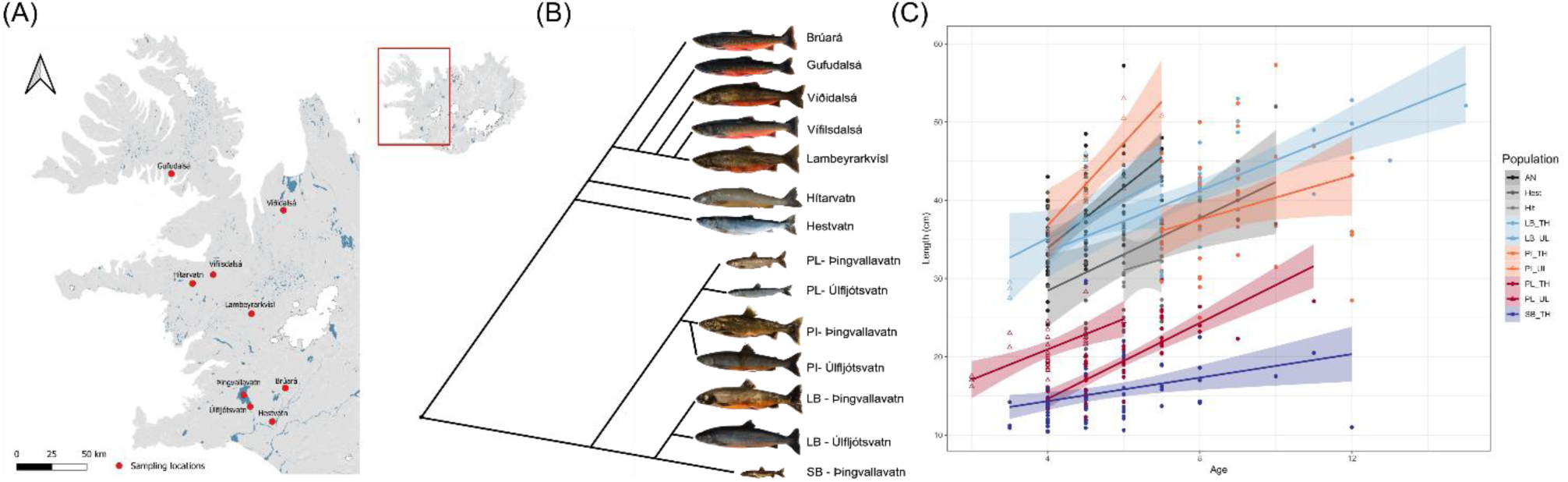
The putative ancestral forms (anadromous charr) tend to be large, but some of the sympatric morphs from the two connected landlocked lakes in South Iceland (Þingvallavatn and Úlfljótsvatn) have reduced body size. (A) Map of the nine sampling locations in West Iceland (see inset), made using the program QGIS (https://qgis.org/). (B) Phylogenetic tree shows the subdivision of the five Anadromous populations (top), two monomorphic lake populations (Hítarvatn and Hestvatn) and the sympatric morphs from the two connected lakes. Tree is a schematic, based on unpublished SNP data from ddRAD-seq data (Han Xiao unpublished data). (C) Scatter plot showing the relationship between age (x-axis) and fork length (y-axis) colored by population for the 417 specimens. Hítarvatn individuals are included in this figure for visualization purposes. We sampled 66 (16%) from five anadromous populations and 349 (84%) from lakes (classification in Supplementary material 3). Sex ratio was slightly skewed, 173 (42%) were female and 219 (53%) males, for 23 (5%) fish we could not identify the sex.

All individuals were measured (weight and length measurement), generally fork length (FL) was used for body length measurements, however only total length (TL) was measured for the Hítarvatn individuals (countermeasures are described below). When possible, sex and sexual maturity were evaluated by dissection. This was difficult for older samples (Hestvatn 2011) as well as individuals provided by landowners (Hestvatn 2024) or recreational fishermen (Hítarvatn, Gufudalsá and Brúará) - still for the majority of samples this data was available. Otoliths were used for age determination under 4X magnification in a stereomicroscope. Information on morph classifications for the charr from Þingvallvatn can be see here (Jónsdóttir et al., 2024), same method was used to classify fish from Úlfljótsvatn.

In this study we focused on the morphology of six paired jaw bones (dentary, articular-angular, quadrate, premaxilla, maxilla and supramaxilla). They were extracted from the head following our maceration protocol (Jónsdóttir, 2022; Jónsdóttir et al., 2024). Bones from both sides (left and right) were arranged stereotypically with a size scale and photographed with a Canon EOS 77D, 50 mm Microlens (Jónsdóttir et al., 2024). To capture the shape of the bones we registered 2D landmarks on the bones from the left side of the head using *tpsdig2* (http://www.sbmorphometrics.org/soft-dataacq.html), using predefined landmarks and principles (Jónsdóttir, 2022; Jónsdóttir et al., 2024). We used the same landmarking scheme as in (Jónsdóttir et al., 2024) but decided to re-landmark the fish for lake Þingvallavatn for this study. For all charr populations we landmarked all bones at least twice with a two-week interval between runs. Repeatability was tested on a subsample and proved high for all bones (Supplementary materials 1 and 2). The same researcher conducted all landmarking for this analysis. Figures of all bones and external figures of most specimens are hosted on Figshare+ project XXXX:XXXX, and for Þingvallavatn in project: Data on craniofacial diversity in Arctic charr from Lake Thingvallavatn Iceland (doi:10.25452/figshare.plus.25118825). Landmark data, flatfiles and R-scripts are available on Github ().

### Statistical analysis

Most analyses follow Jónsdóttir et al. (2024), briefly summarized here. All statistical analysis were done using R (4.4.2)(R Core Team, 2024) in RStudio (2024.12.1+563)(Posit team, 2025). The landmark data was analyzed using the package *geomorph (4.0.10*)(Adams et al., 2025; Baken et al., 2021; Collyer & Adams, 2018, 2024). For all analysis we used aligned shape variables, aligned using Generalized Procrustes Analysis (GPA), function: *gpagen*. Using the function *plotOuliers* we identified outliers and removed them if bones were damaged. Repeatability of the landmark sets were tested in three ways: 1) calculating the general intraclass correlation, 2) using Procrustes ANOVA (*procD.lm*) to characterize the measurement error, and 3) using two-block partial least squares (PLS) analysis (*two.b.pls*) to assess the degree of association between the two trials.

For each of the six bones, centroid size (CS) was calculated (CS_bone_; indicates use of CS of a bone and applied to each bone separately). We then used Procrustes ANCOVA (*procD.lm*) to examine the effects of body size (log_e_ FL), population and their interaction on bone size (CS_bone_). For all tests we used a Bonferroni corrected threshold, α = 0.008. Hítarvatn individuals were not included in this analysis. Due to discrepancies in the method of length measurement among populations. We used bone size (CS_bone_) as the size covariate for further shape analysis, which was shown to be highly correlated to TL and FL in Arctic charr (Jónsdóttir et al., 2024) and this was confirmed in this dataset before statistical analysis. Therefore, Hítarvatn individuals were included in the following analysis. We used Procrustes ANCOVA (*procD.lm*) to examine the effects of size (CS_bone_), population and their interaction on bone shape, Bonferroni corrected threshold, α = 0.008. We compared three models to determine the best fit: null (shape = CS_bone_), reduced (shape = CS_bone_ + population) and full model (shape = CS_bone_ x population). When the full model had the best fit, we tested for significant differences in both vector angles (vector correlation) and vector lengths (rate of shape change per unit covariate). For visualization, Principal Component Analysis (PCA, using *plotTangentSpace*) was used. If a bone did not show significant differences in allometry among populations, size corrected shape variables were used for visualization. Shape variation along PC axes was summarized with deformation grids and vector plots from *shape*.*predictor*.

## Results

In total 415 sexually mature Arctic charr were analyzed and we found significant population, sex and population by sex effects on body size (fork length and weight, Supplementary material 4). Body size was rather similar between the anadromous populations, the monomorphic lakes and the PI- and LB-charrs, but predictably the SB- and PL-charr from Þingvallavatn/Úlfljótsvatns were smaller (Supplementary materials 3, 5 and 6, SB-charr the smallest). There was only a significant population and sex effect on age, no interaction detected (Supplementary materials 4 and 5). However, maturity had significant population effect and a population by sex interaction but showed no sex dependence (Supplementary material 4 and 5).

Bone landmarking was highly repeatable (Supplementary materials 1 and 2), both the correlation coefficients (r = 0.83 – 0.93) and association between replicates (r-PLS = 0.81 – 0.99) were high for all bones and proportion of variation due to measurement error was low relative to variation among individual (R^2^ = 0.001 – 0.004 vs. R^2^ = 0.91 – 0.97, Supplementary material 2). We explored sexual dimorphisms in bone size and shape (Supplementary materials 7 and 8) and generally found a significant but weak sex effect on all bones for both sets of traits. Population by sex interactions where generally non-significant for both bone size and shape. Thus, for the following analyses we mainly explored population and size effects and dropped sex as a term.

### Bone size variation by populations

Bone size correlated strongly with body size and varied significantly by population (Table 1). However, the population by body size interaction accounted only for a small fraction of the variation (R^2^ < 0.6%). Further analysis revealed that the allometric differences were mainly driven by differences in vector lengths (vector angle differences were never significant) between the larger bodied charr populations (AN-, PI- and LB-charr) and smaller bodied populations (SB- and PL-charr) (Supplementary materials 9 and 10). Therefore, we concluded that the bone size allometry was not truly different by populations and assumed common allometry for the subsequent analyses.

**Table 1:** Bone size relates to body size and differs slightly by populations. Influence of body size (FL, fork length) and population on bone size (CS) variation in six jaw bones. Results from Procrustes ANCOVA, with AN-charr treated as five separate populations. Hítarvatn individuals excluded from this analysis.

| **Bone Size** | **Log<sub>e</sub> FL** |  |  |  | **Population** |  |  |  | **Population x Log<sub>e</sub> FL Interaction** |  |  |  |
|---|---|---|---|---|---|---|---|---|---|---|---|---|
|  | **R<sup>2</sup>** | **F** | **Z** | **P** | **R<sup>2</sup>** | **F** | **Z** | **P** | **R<sup>2</sup>** | **F** | **Z** | **P** |
| Dentary | 94.32% | 12326.80 | 17.71 | **\*0.001** | 2.30% | 23.10 | 11.89 | **\*0.001** | 0.43% | 4.37 | 4.59 | **\*0.001** |
| Articular-angular | 96.01% | 14581.77 | 16.88 | **\*0.001** | 1.10% | 12.85 | 9.50 | **\*0.001** | 0.36% | 4.15 | 4.63 | **\*0.001** |
| Quadrate | 95.62% | 14903.36 | 16.90 | **\*0.001** | 1.64% | 19.71 | 13.52 | **\*0.001** | 0.29% | 3.46 | 3.81 | **\*0.001** |
| Premaxilla | 92.97% | 10047.96 | 16.07 | **\*0.001** | 2.88% | 23.94 | 12.49 | **\*0.001** | 0.57% | 4.76 | 4.73 | **\*0.001** |
| Maxilla | 94.50% | 11973.67 | 16.58 | **\*0.001** | 2.05% | 19.93 | 11.82 | **\*0.001** | 0.42% | 4.10 | 4.58 | **\*0.001** |
| Supramaxilla | 94.35% | 10713.62 | 15.32 | **\*0.001** | 2.07% | 18.11 | 9.69 | **\*0.001** | 0.25% | 2.20 | 2.27 | 0.014 |

R2 = coefficient of determination; F = F-statistic; Z = Z-statistic; P = P-value, P values below the Bonferroni corrected threshold (α = 0.008) are shown in bold and with asterisk.

The bone size differed between the populations (Table 1, Figure 2 and Supplementary material 11), but as the allometric trajectories did not vary by population (above) we analyzed the residuals for each bone (Supplementary materials 12 and 13). First off, bone size varied by AN-charr populations, and had a similar pattern for all bones (individuals from river Víðidalsá with the largest bones, Supplementary materials 11 and 14). When AN-populations were pooled, they generally had large bones (Figure 2, Supplementary materials 11, 13 and 15). The large-bodied fish from Hestvatn had similar bone sizes as the other larger lake morphs (LB- and PI-charr), though this varied a bit (Figure 2, Supplementary materials 15). The sympatric morphs in Þingvallavatn differed markedly, however the same morphs within Úlfljótsvatn appeared to have mostly similar bone sizes (Figure 2, Supplementary materials 14 and 15). Across the two connected lakes, it appeared that the Úlfljótsvatn morphs had smaller bones than their corresponding “sister” morph in Þingvallavatn. Though this was only formally significant in three pairwise tests of eighteen possible tests (Supplementary materials 14 and 15), a sign test showed that this tendency was significant (p = 0.008).

**Figure 2:**
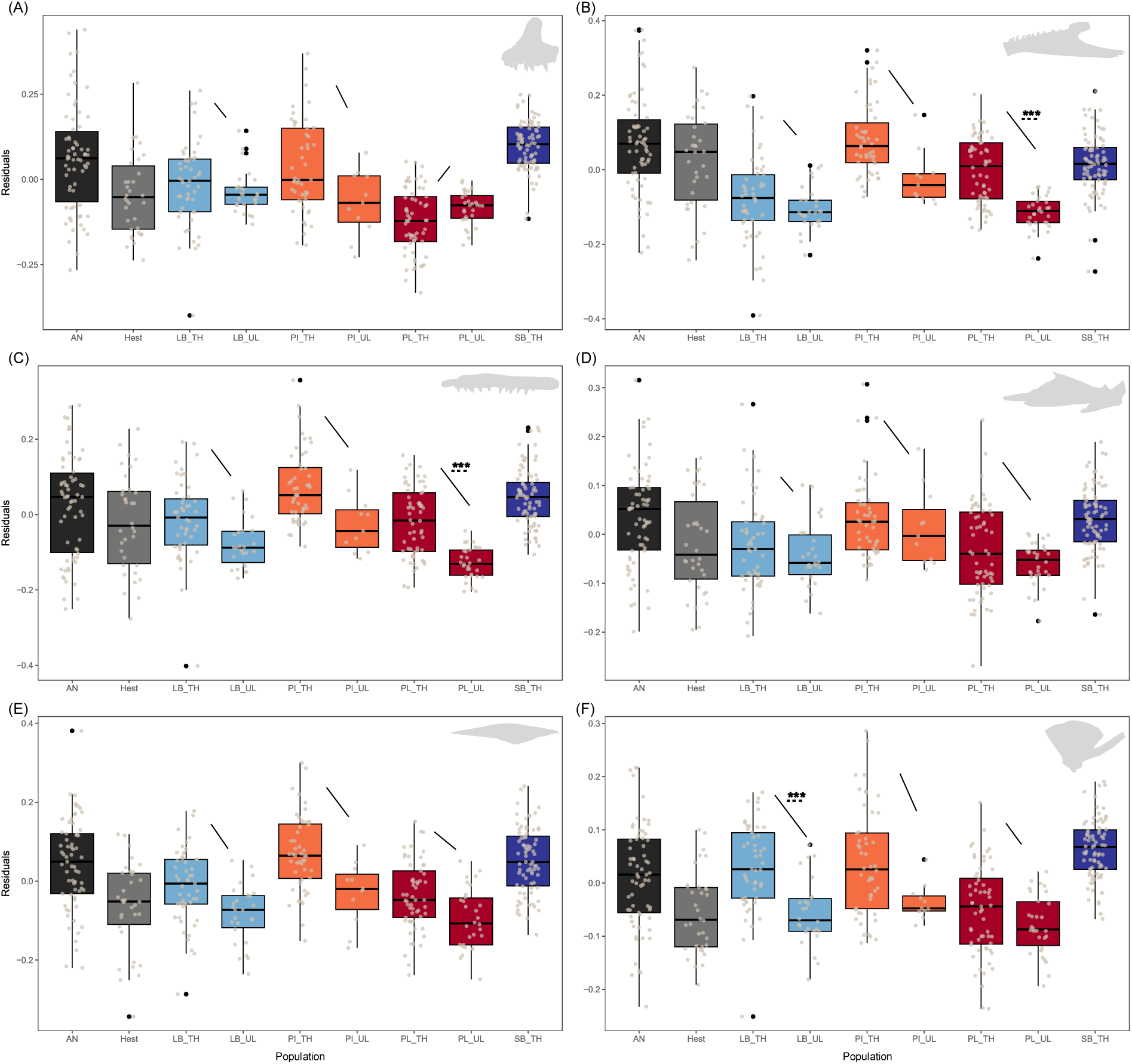
Bone size variation by populations of Arctic charr, analyzed as residuals from common linear allometry for each bone. Boxplots show the residuals by population, in this picture the five AN-charr populations were pooled (black) and Hítarvatn individuals were excluded. Color code follows figure 1. Upper jaw bones on left side, from top A) premaxilla, C) maxilla and E) supramaxilla. Lower jaw bones on right side, from top: B) dentary, D) articular-angular and F) quadrate. In the upper right corner of each graph is an illustration of each bone. Arrow on each graph visualizes the reoccurring pattern of fish from Úlfljótsvatn having smaller bone then their corresponding “sister” morph in Þingvallavatn. Three asterisks are by arrows with significant post-hoc Tukey tests (Supplementary materials 14 and 15).

### Jaw bone shape allometry and population differences

Procrustes ANOVAs showed that the shape of six jaw bones varied by population and with bone size. Bone shape and size associated significantly in all bones (Table 2), in general more strongly in the lower jaw bones (allometry explained around 20 – 30% of variation for the lower jaw bones (dentary, articular-angular and quadrate) and 10 – 20% for the upper jaw bones (premaxilla, maxilla and supramaxilla)). Shape of all bones varied by populations, strongly in four (dentary, articular-angular, maxilla and premaxilla) where around 18 – 24% of the shape variation was explained by population, but weaker for the quadrate and supramaxilla (11-13% of the variation). The allometric trajectories by population were significant for all bones (Table 2), however, subsequent comparisons of vector correlation angles and lengths (Supplementary materials 16 and 17) indicated only substantial changes in the allometry by population for two bones (dentary and premaxilla, Figure 3B and D). In both bones, were these allometric differences mainly driven by differences between the derived Þingvallavatn morphs, however some differences between anadromous and lake charr were also detected (Supplementary materials 16 and 17). Below we will present mean shape differences from the full model (shape = CS_bone_ X population) only for the dentary and premaxilla and results from the reduced model (shape = CS_bone_ + population) for other bones. All visualizations, unless otherwise noted, have been corrected for size to remove the effects of allometry on shape.

**Figure 3:**
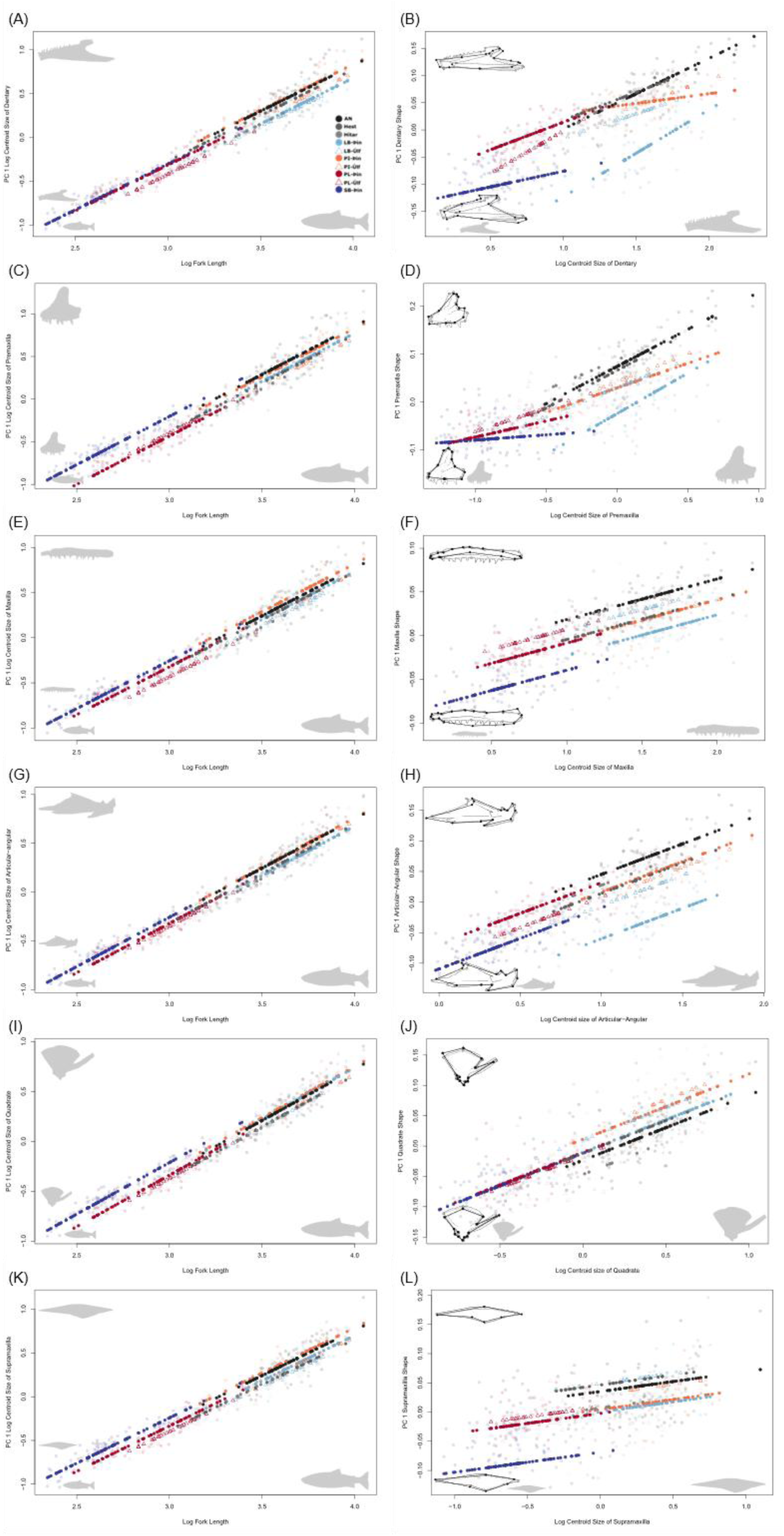
Allometry of size (left) and shape (right) for six jaw bones. From top to bottom: A) and B) dentary, C) and D) premaxilla, E) and F) maxilla, G) and H) articular-angular, I) and J) quadrate and K) and L) supramaxilla. Inset are pictures of each bone (and fish). Left panels (A, C, E, G, I, K): relationship between bone size (y) and fork length (x) by group; Right panels (B, D, F, H, J, L): relationship between bone shape (y) and bone size (x). On the right are inset associated shape changes related to each component, grey outlines the mean shape, and the black the two extremes for each PC. On all figures are shown both predicted values (filled symbols) from a linear regression and individual values (faded).

**Table 2:** Bone shape is affected by size and population. Influence of bone size and population on bone shape variation in six jaw bones in Arctic charr. Results from Procrustes ANCOVA, with AN-charr treated as five separate populations.

| **Bone Shape** | **Log<sub>e</sub> CS** |  |  |  | **Population** |  |  |  | **Population x Log<sub>e</sub> CS Interaction** |  |  |  |
|---|---|---|---|---|---|---|---|---|---|---|---|---|
|  | **R<sup>2</sup>** | **F** | **Z** | **P** | **R<sup>2</sup>** | **F** | **Z** | **P** | **R<sup>2</sup>** | **F** | **Z** | **P** |
| Dentary | 32.14% | 304.96 | 5.61 | **\*0.001** | 24.23% | 17.68 | 14.82 | **\*0.001** | 3.06% | 2.23 | 5.38 | **\*0.001** |
| Articular-angular | 24.56% | 185.24 | 7.19 | **\*0.001** | 21.49% | 12.47 | 12.18 | **\*0.001** | 2.90% | 1.68 | 3.52 | **\*0.001** |
| Quadrate | 25.73% | 164.58 | 7.82 | **\*0.001** | 11.44% | 5.63 | 9.46 | **\*0.001** | 3.28% | 1.61 | 3.02 | **\*0.002** |
| Premaxilla | 19.40% | 127.92 | 7.50 | **\*0.001** | 18.17% | 9.22 | 14.27 | **\*0.001** | 3.73% | 1.89 | 5.14 | **\*0.001** |
| Maxilla | 18.30% | 121.37 | 8.32 | **\*0.001** | 20.08% | 10.25 | 14.36 | **\*0.001** | 3.73% | 1.90 | 3.99 | **\*0.001** |
| Supramaxilla | 9.99% | 51.65 | 8.59 | **\*0.001** | 12.60% | 5.01 | 9.96 | **\*0.001** | 4.34% | 1.73 | 2.84 | **\*0.001** |

### Minor differences among Anadromous populations in bone shape

The five anadromous populations were used to explore the variation among potential “ancestral” forms of charr. There were minor differences among the five AN-charr populations (Supplementary material 18 and 19), with few significant pairwise differences between populations seen (only in three bones, dentary, articular-angular and maxilla, Figure 4A and Supplementary material 19). In all cases, were the differences between fish from river Víðidalsá in the north and other locations (dentary, vs. rivers Gufudalsá and Víðidalsá p = 0.001, articular-angular, vs. rivers Brúará and Víðidalsá p = 0.007 and maxilla, vs. rivers Lambeyrarkvísl and Víðidalsá p = 0.001, Supplementary material 19). In the dentary, the AN-populations mainly diverged along PC1 (54% of the total variation) and PC3 (7.6%). This PC also separated the lake charr and the sympatric morphs (see below). For the articular-angular AN-populations varied on PC2 (16%) and PC4 (8.5%), but neither axis separated benthic and pelagic morphs. For the maxilla, AN-populations diverged along PC1 (30%) which did not separate the benthic and pelagic morphs (Supplementary material 18).

**Figure 4:**
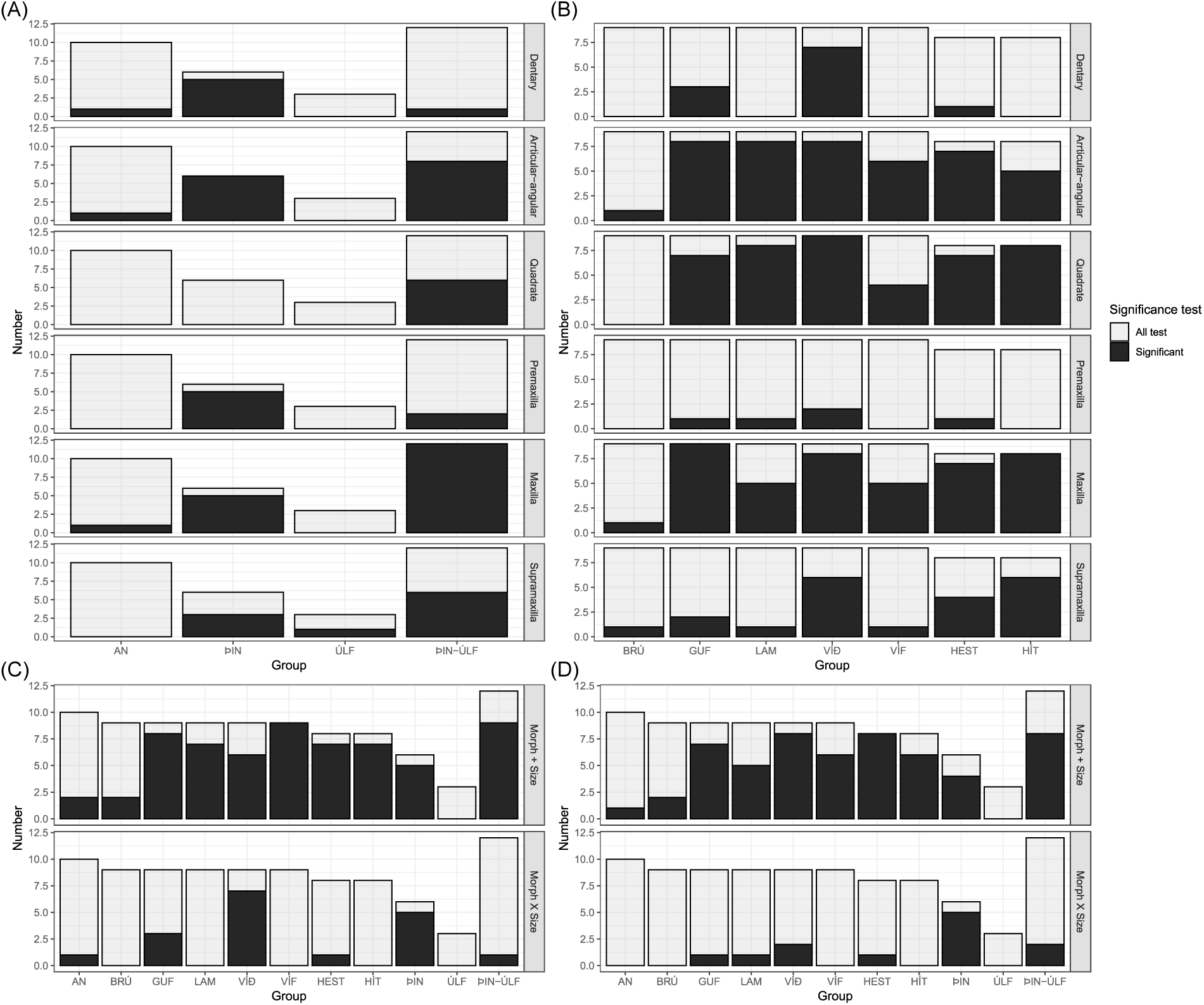
Few significant pairwise differences in bone shape between Anadromous populations, some among sympatric morphs. Bar plots show the number of significant pairwise tests for each bone shape per population/morph. Gray colored bars are all tests performed, and black the number of significant ones. A) Number of tests within categories or location, the five AN-charr populations (AN), between the four morphs in Þingvallavatn (Þin), between the three morphs in Úlfljótsvatn (ÚLF) and between Þingvallavatn and Úlfljótsvatn (ÞIN-ÚLF). B) Number of test when comparing each AN-charr populations (BRÚ, GUF, LAM, VÍÐ, VÍF) to lake charr and comparing nonspecialized lake charr to derived sympatric morphs (HEST, HÍT). C) Number of tests for dentary shape, when using either the reduced (shape = CS_bone_ + population) and full model (shape = CS_bone_ x population). D) Number of tests on premaxilla shape, when using either the reduced (shape = CS_bone_ + population) and full model (shape = CS_bone_ x population). Figures were made based on Supplementary materials 19 to 21.

### Pelagic bone shape appears to be ancestral

Since variation among the AN-charr populations were minor, we grouped them and tested for differences relative to the lake populations. AN-charr diverged from at least one lake morph for each bone (Supplementary material 20). In general, the bone shape of the monomorphic charr populations aligned with the AN-charr, and appeared more pelagic (on the PC that separated pelagic and benthic morphs). Specifically, for the articular-angular and maxilla shape the AN-charr differs from almost all the lake morphs, except PI-charr from Úlfljótsvatn (Supplementary material 20). For both bones the AN-charr generally overlapped in shape space with the pelagic morphs along the main benthic vs. pelagic separating axis (PC4 (6%) for maxilla and PC1 (32%) for articular-angular, Figure 5B and D). Weaker differences were seen in the dentary and premaxilla, with the AN-charr only significantly differing from some charr in lakes (Supplementary material 20). For the dentary the PCA (Figure 5D) revealed a similar pattern to the articular-angular and the maxilla, with AN-charr generally overlapping with the pelagic morphs along the main benthic vs. pelagic separating axis (PC1, 54%). However, the premaxilla pattern was different (Figure 5A), the AN-charr landed in the middle on the main benthic vs. pelagic separating axis (PC2, 19%). While for the quadrate and supramaxilla pairwise analysis indicated differences between AN-charr and lake morphs (Supplementary material 20), the PCAs did not reveal clear separation along any of the axes (Supplementary material 22). We ran complementary CVA analysis for these two bones which mainly showed AN-charr separating from the SB-charr, with consistent results (Supplementary material 23).

**Figure 5:**
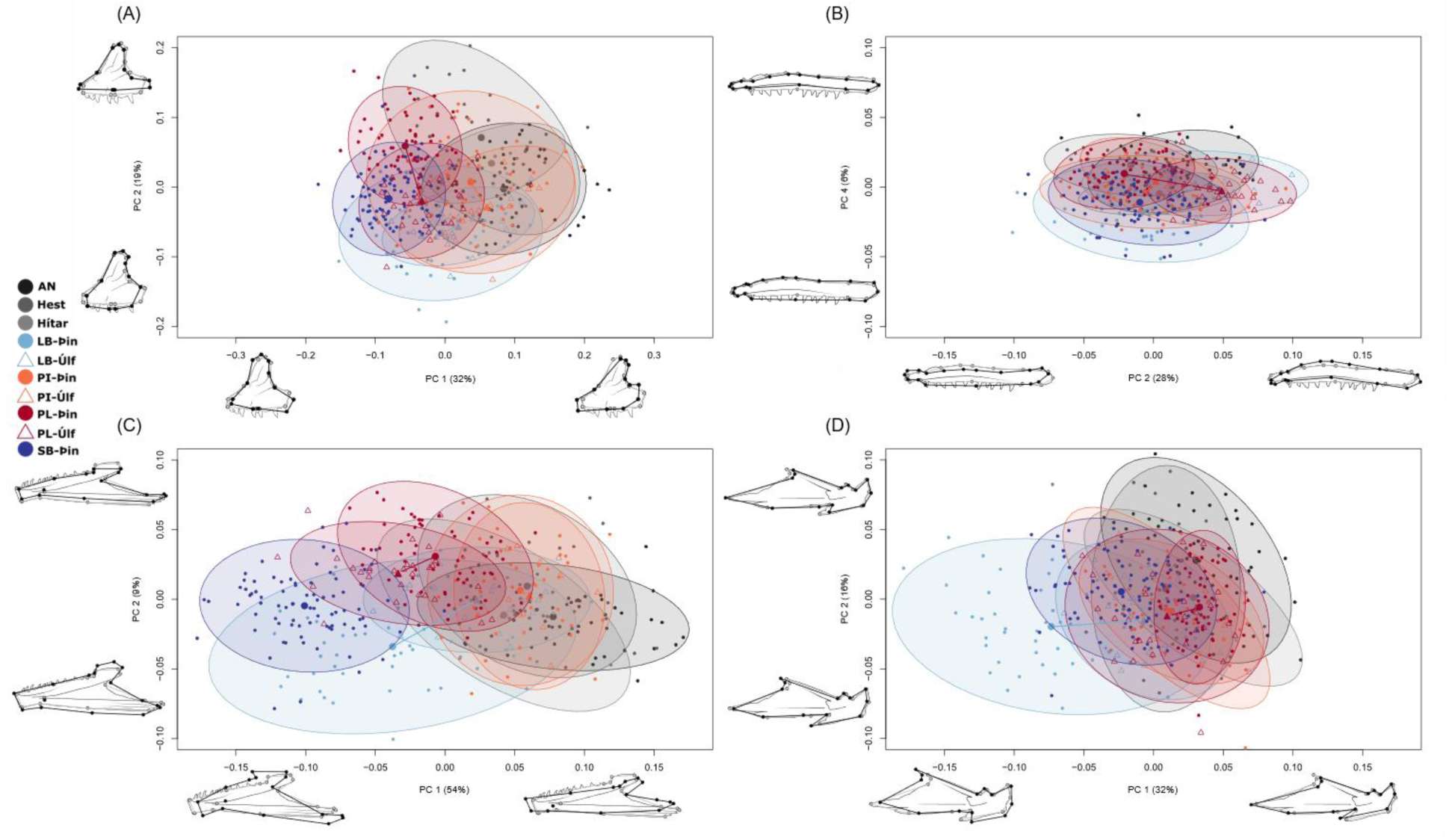
Substantial shape variation by charr morphs and populations in three jaw bones. PC-plot with deformation grids depict the variation in shape for two upper jaw bones (A) premaxilla and B) maxilla) and two lower jaw bones (C) dentary and (D) articular-angular). Note that the effects of size were not removed for the premaxilla (A) and dentary (C), due to differences in allometric trajectories between groups. Size effects were removed for the maxilla (B) and articular-angular (D) plots. None of the deformation grids are magnified, grey outline represents mean shape and black lines the two extremes for each PC. Each dot or triangle represents an individual and the ellipses 95% CI for the distribution by group (large dot or triangle represents the mean shape of each group in these dimensions).

Next, we compared the Hestvatn and Hítarvatn charr (non-specialized pelagic fish from lakes with only one morph), to the sympatric morphs from Þingvallavatn and Úlfljótsvatn. Charr from Hestvatn diverged from all the sympatric morphs (expect the PI-charr from Úlfljótsvatn) for the articular-angular and maxilla bone (Supplementary material 19 and 20). For both bones, the Hestvatn fish overlapped in shape space with pelagic morphs on the main benthic vs. pelagic separating axis (PC4 (6%) for maxilla and PC1 (32%) for articular-angular, Figure 5B and D). For the dentary and premaxilla shape the charr from Hestvatn only separated significantly from the Þingvallavatn PI- and SB-charr (Supplementary material 19 and 20). The fish from Hítarvatn, generally differed from all derived lake morphs for the maxilla shape. For the articular-angular, Hítarvatn charr differed from LB-charr from both lakes and the smaller Þingvallavatn morphs (PL- and SB-charr). However, for both the dentary and premaxilla, Hítarvatn charr did not diverge from any of the sympatric morphs (Supplementary material 19 and 20). Similar to the Hestvatn charr, Hítarvatn charr generally clustered/overlapped with the other pelagic morphs in shape space for all bones (Figure 5).

### Clear separation of sympatric morphs in large lake, but none in connected smaller lake

Previously we documented large differences in bone shape between the sympatric Þingvallavatn morphs (Jónsdóttir et al., 2024) - that data is included here. However, while the same morphs (except SB-charr) are common in the smaller downstream lake Úlfljótsvatn, none of them differed significantly from one another in the shape of any bone (figure 3, Supplementary material 19 and 20).

In detail, the strongest differences between the Þingvallavatn morphs were in the two most frontal bones (dentary and premaxilla). The morphs clearly diverged in lower jaw shape (Supplementary material 19 and 20), strongest in the dentary along PC1 (54%) which mainly separates benthic and pelagic morphs (Figure 5C). A similar but weaker pattern was seen in the articular-angular (Supplementary material 19 and 20), where morphs separated along PC1 (32%) – along a benthic and pelagic axis (Figure 5D). The quadrate shape on the other hand showed no separation of the Þingvallavatn morphs (Supplementary material 19 and 20). In the upper jaw the clearest morphs differences were in the premaxilla (Supplementary material 19 and 20). The morphs mainly separated along PC2 (19%), delineating benthic and pelagic fish (differences in the posterior and anterior limbs of the premaxilla). For the maxilla, the Þingvallavatn morphs mainly differed in the width of the bone (along PC4, 6%). Generally, the supramaxilla shape did not vary by morph (Supplementary materials 19 to 23).

### Plasticity and genetic divergence affect jaw shape in distinct morphs

To explore plasticity in the Þingvallavatn and Úlfljótsvatn system we tested for morphological differences in “sister” morphs across the two lakes. Of those PL-charr are genetically very similar across the two lakes, but the LB- and PI-charr differ subtly between them (Xiao et al., 2026). Pairwise tests revealed significant differences between “sister” morphs in four bones (maxilla, articular-angular, quadrate and supramaxilla, Figures 5 and 6 and Supplementary material 19 and 20). The general tendencies were shifts towards the middle (in shape space) in the environmentally more homogeneous Úlfljótsvatn. PL-charr in that lake had more benthic phenotype and LB-charr more pelagic.

**Figure 6:**
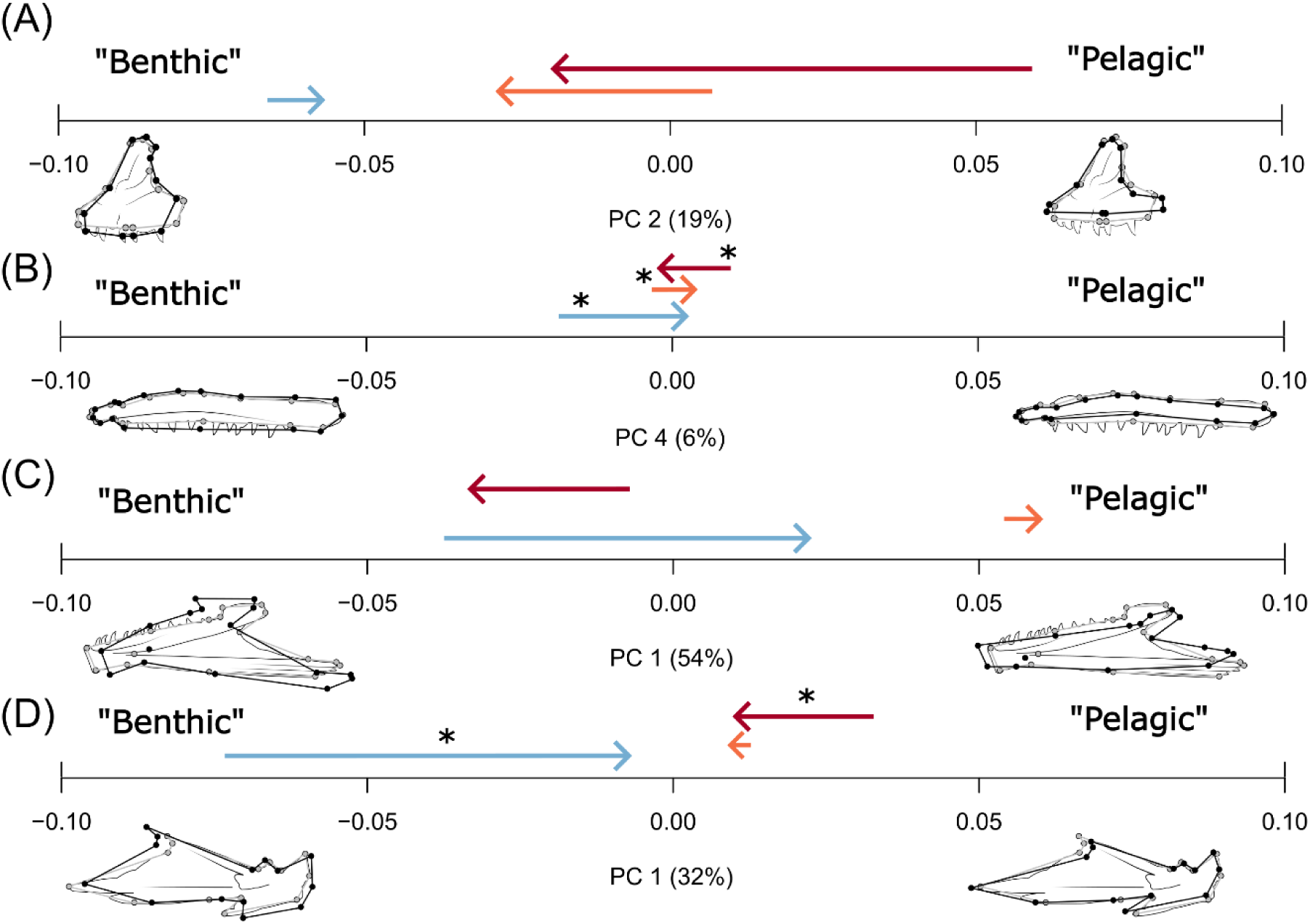
Indication of plastic shift in PL-charr by lake and genetic influence on shape in LB-charr. Bone shape differences between “sister” morphs across Þingvallavatn and Úlfljótsvatn. Visualized are shape differences between the morphs in the two lakes (mean per group) along the main benthic vs. pelagic shape axis for each bone (from top to bottom: premaxilla (A), maxilla (B), dentary (C) and articular-angular (D)). Based on PCA (see Figure 5). Arrows are colored by morph and point from Þingvallavatn (average shape) towards Úlfljótsvatn (average shape). An asterisk (*) by an arrow indicates significant pairwise tests (Supplementary material 19). Note, the dentary and premaxilla shape estimates were not size corrected (to account for differences in allometry).

In more detail, for the maxilla differences were seen between all “sister” morphs (LB-LB, PL-PL and PI-PI). In the articular-angular, there were differences between the LB- and PL-morphs by lakes, but not the PI-charr (the supramaxilla showed the same pattern). As mentioned above, no PCs showed clear difference between any morph pairs for the quadrate or the supramaxilla (Supplementary material 22). For the articular-angular and maxilla we saw similar patterns in the differences between the Þingvallavatn and Úlfljótsvatn “sister” morphs, along the shape axis capturing benthic vs. pelagic separation (Figure 5B and 5D and Figure 6B and 6D). For the maxilla, along PC4 (benthic vs. pelagic) the PL-charr from Úlfljótsvatn were slightly closer to the maximum (“more benthic”) then the PL-charr from Þingvallavatn. However, the LB-charr from Úlfljótsvatn was slightly closer to the minimum (“more pelagic”) relative to the LB-charr of Þingvallavatn (Figure 6B). A similar pattern was observed for the articular-angular along PC1 (main benthic vs. pelagic axis), the PL-charr from Úlfljótsvatn were slightly closer to the minimum (“more benthic”) than the PL-charr from Þingvallavatn and the LB-charr from Úlfljótsvatn closer to the maximum (“more pelagic”) compared to the LB-charr from Þingvallavatn (Figure 6D). Despite the dentary and premaxilla not showing significant differences between any of the “sister” morphs (Supplementary material 19 and 20), the same patterns were seen in the PCA (Figure 5A and 5C and Figure 6A and 6C). For both bones the PL-charr from Úlfljótsvatn appear to have a “more benthic” shape compared to PL-charr from Þingvallavatn and LB-charr from Úlfljótsvatn have a “more pelagic” shape compared to the LB-charr from Þingvallavatn (Figure 6A and 6C). Other pairwise comparisons (not between “sister” morphs) across the lakes were also significant with the maxilla and articular-angular having the most significant pairwise comparisons (Supplementary material 19 and 20), not described further here.

## Discussions

Salmonids and other northern freshwater fishes show impressive phenotypic diversity and patterns consistent with parallel and repeated adaptation of traits to freshwater niches after losing anadromy (Brachmann et al., 2021; Jacobs et al., 2020; Kapralova et al., 2011; Salisbury et al., 2022). Most studies have explored putative derived populations, with the allopatric and sympatric polymorphism in the *Salvelinus* genus being prominent examples (Brachmann et al., 2021; Jacobs et al., 2020; Kristjánsson et al., 2012; Salisbury et al., 2022; Snorrason et al., 1994). Here we studied putative ancestral and derived populations and found the former to be rather uniform in jaw bone shape, the benthic forms to have derived shape, and an indication of plastic response in bone shape in one morph most likely reflecting niche differences.

### Minor variation among the anadromous ancestral groups

Research on *Salvelinus* diversity has generally focused on characterizing derived morphs, assuming that anadromous charr has relatively low phenotypic diversity, at least compared to derived morphs (Esin et al., 2018; Kristjánsson et al., 2012; Sandlund et al., 1992). If used, most papers only use anadromous charr from a single river system to represent an outgroup or putative ancestor (Brachmann et al., 2021; Dellinger et al., 2023), notable exception being population genetic study on Canadian populations (Salisbury et al., 2022). We tested whether anadromous charr from different regions in Iceland had different phenotypes. Unpublished data indicates genetic separation of anadromous charr by these regions (Xiao et al., Unpublished). Generally, the anadromous charr differed in size by rivers (both fork length and weight). With river Víðidalsá in north Iceland having the longest and heaviest fish while river Gufudalsá (west) had the shortest and lightest fish. Similar pattern was seen in bone size with fish from river Víðidalsá having the largest bones, fish from rivers Brúará (south) and Gufudalsá having the smallest bones and fish from rivers Lambeyrarkvísl and Vífilsdalsá (both west) falling in the middle. Despite these size variations, there were only minor variations in bone shape among the anadromous populations. Notably, the bone shape differences among the anadromous populations appeared on distinct axis of shape (not the ones separating the sympatric morphs). Obviously, the anadromous populations may vary in other phenotypic traits not examined in this study or differ more strongly on a larger geographic scale.

The minimal variation in jaw bone shape argues that adult anadromous jaw phenotypes are constrained (though relative size of bones may vary). This may be because piscivory (Collar et al., 2009) or other aspects of the anadromous charr life cycle places high functional demands on their bone shape suppressing divergence. There was some variation within each anadromous population, in most traits. We only examined variation in adult anadromous morphology, but the juvenile anadromous charr utilize benthic habitats in rivers (while adults operate in pelagic waters of estuaries and the sea) (Klemetsen et al., 2003). Generally, such ontogenetic niche shifts are often accompanied by remodeling of feeding elements (Gidmark et al., 2019; Karagic et al., 2020). Therefore, it is possible anadromous charr vary more in bone shape across each individual’s lifetime rather than across different river systems. It is also possible, though not very likely, that there are more notable bone shape differences in juveniles of anadromous charr.

We used anadromous charr and charr from the monomorphic lakes Hestvatn and Hítarvatn as putative outgroups for the derived Arctic charr morphs from lakes Þingvallavatn and Úlfljótsvatn. Generally, this assumption held. Despite some minor between group differences the putative outgroups tended to cluster together and aligned with the pelagic morphs for the most variable bones (dentary, articular-angular and maxilla). This was clearest for the dentary and articular-angular where these outgroups fell firmly on the “pelagic side” - even being “more pelagic” than the derived pelagic lake morphs. The shape changes in the benthic forms lead to shorter and boxier lower jaw, which is consistent with subterminal jaw morphology, hypothetically an adaptation to bottom feeding (Jónsdóttir et al., 2024; Skúlason, Noakes, et al., 1989). However, the premaxilla showed a different pattern, the anadromous charr fell in the middle, between the derived benthic and pelagic lake morphs. For this bone, the charr from Hestavatn and Hítarvatn sat on the “pelagic side”, having similar shape to the derived pelagic lake morphs. Independent evolution of modular anatomy is well known, and the definition of the derived state (“benthic” or “pelagic”) varies by bone. As most groups overlapped for maxilla shape, it was not possible to define ancestral and derived forms for this bone. Jointly the data suggests that the pelagic morphs in sympatry have a phenotype closer to the ancestral phenotype, but the benthic morphs may be more derived (for most bones). This agrees with previous hypotheses which postulated that the pelagic morphs in lake Þingvallavatn (PI- and PL-charr) resemble the anadromous ancestor (Parsons et al., 2010; Sandlund et al., 1992; Skúlason, Noakes, et al., 1989). But the premaxilla shape seemingly evolved both in the sympatric pelagic and benthic morphs in the lake. These results allow us to firmly demarcate derived forms of seemingly important functional feeding elements in Arctic charr. It would be interesting to see if the same holds in other species within the genus, within salmonids or even other teleost orders, that show similar ecological divergence (Avery et al., 2026; Bernatchez et al., 2010; Esin et al., 2018; Osborne et al., 2026). Similar feeding elements (dentary and premaxilla) have been found to differ in shape between benthic and pelagic populations of Trinidadian guppies (*Poecilia reticulata*), though the shape divergence is not the same as the benthic – pelagic divergence in Arctic charr (Avery et al., 2026).

### Differences in shape of feeding related bones in two connected lakes

The connected lakes Þingvallavatn and Úlfljótsvatn are landlocked and most likely were colonized once by charr, that radiated into four distinct morphs (Xiao et al., 2026). Þingvallavatn is larger and deeper, has multiple distinct habitats and hosts the four sympatric morphs, but Úlfljótsvatn is smaller and shallower, with more uniform habitat. We analyzed variation in body size and bone morphology in the three morphs common in both lakes. We saw neither differences in body length nor weight between “sister” morphs in two lakes, however the Úlfljótsvatn fish were significantly younger. This pattern (Úlfljótsvatn charr being of a similar size as older Þingvallavatn charr) indicates the Úlfljótsvatn morphs have higher growth rate than their Þingvallavatn “sister” morphs, due to growth plasticity (Karjalainen et al., 2016; Skulason & Smith, 1995). Growth rates of fish are affected by many biotic and abiotic factors, for example temperature (Jobling, 1983), food availability, and population density (Amundsen et al., 2007). Available data suggests the two lakes have similar water temperature, Supplementary material 24 (Malmquist et al., 2020). Systematic surveys have been done in lake Þingvallavatn (Snorrason, Unpublished), but not in Úlfljótsvatn, and no standardized data is available on food availability in the two lakes. Stomach content analyses indicate less prey diversity in Úlfljótsvatn, with snails dominating the menu (Sandlund et al., 1992; Xiao et al., 2026). However, we cannot confirm which factors differ between the lakes. The observed differences in maturity, Úlfljótsvatn morphs generally having less average maturity then their Þingvallavatn “sister” morph (Supplementary material 5), may reflect that the sampling took place at different times (Úlfljótsvatn in early September / Þingvallavatn late September to early October). In lake Þingvallavatn the spawning time of LB-charr is August, the PL- and PI- charr in later September (Sandlund et al., 1992), but it is unknown when (and even if) spawning occurs in lake Úlfljótsvatn.

External head shape was previously examined for these morphs (Xiao et al., 2026) and revealed that “sister” morphs in the two lakes generally overlapped in head shape, especially the two PL-morphs (Xiao et al., 2026). Our previous study on the Þingvallavatn charr morphs revealed more pronounced differences in bone shape than external head/body shape (Jónsdóttir et al., 2024). Bone shape may thus better capture ecological variation than metrics of external shape and we maintain that our results of significant bone shape variation do not contradict previous findings (Xiao et al., 2026). The bone size and shape across the lakes showed an interesting pattern, though not always significant. A consistent tendency for both bone shape and size were some significant pairwise differences between the Þingvallavatn morphs while no significant differences were observed between any of the Úlfljótsvatn morphs. Therefore, in bone shape and size the Úlfljótsvatn morphs appear more similar to each other, but the Þingvallavatn morphs more distinct.

The morphs in Þingvallavatn appear to be diverging, possibly due to positive selection due to diet differences (Snorrason et al., 1994). The opposite pattern was seen in Úlfljótsvatn, less difference was found between morphs. As ecological divergence can be sensitive to environmental factors (Hendry, 2017) this is not surprising. Xiao et al. (2026) found the PL-charr in the two lakes to be genetically very similar, but more differences were between the PI- and LB-<u>“</u>sister<u>”</u> morphs. Interestingly, the population genetics also provide evidence for more hybridization (gene flow into PI- from LB-charr) in lake Úlfljótsvatn compared to lake Þingvallavatn (Xiao et al., 2026), possibly due to reduced segregation in spawning sites or increased overlap of spawning periods. Lake Úlfljótsvatn has a restricted pelagic zone and the main prey items (crustacean zooplankton) of adult PL-charr in Þingvallavatn (Malmquist, 1992; Malmquist et al., 1985) are scarce. Thus in Úlfljótsvatn, all morphs feed on the same prey resource (snails) (Jónsson & Jóhannsson, 2024) which have the same seasonal fluctuations (same peaks), while distinct niches provide unique food sources in Þingvallavatn (Malmquist et al., 1985). This could have two consequences, i) more similar spawning times for the morphs and increased likelihood of hybridization and ii) the development of pelagic morphs might respond to altered food sources (plasticity).

### Plasticity and genetic divergence influence bone shape variation

Genetic differentiation and hybridization may explain the shifts in bone shape in the LB-charr in Úlfljótsvatn. However, since the PL-charr are genetically very similar in the two lakes (Xiao et al., 2026), genetics can hardly explain the shape variation observed. As was mentioned above, stomach contents (Jónsson & Jóhannsson, 2024; Xiao et al., 2026) show that adult PL-charr in lake Úlfljótsvatn primarily feed on benthic freshwater snails, which is rarely observed in adult PL-charr in lake Þingvallavatn (Malmquist, 1992; Malmquist et al., 1985). We postulate that the bone shape differences in the PL-charr between the two lakes reflect plastic responses, known in the species (Brachmann et al., 2021; Küttner et al., 2013; Parsons et al., 2011).

Similar external morphology and genetics of the two PL-charr “sister” morphs (Xiao et al., 2026) as well as the large population size and considerable mobility of PL-charr in Þingvallavatn (Sandlund et al., 1992; Snorrason et al., 1992) indicate substantial rates of migration of PL-charr from lake Þingvallavatn to lake Úlfljótsvatn (Xiao et al., 2026). This immigration may maintain genetic similarity and prevent local extinction of the PL-charr in lake Úlfljótsvatn. It is unknown what fraction of PL-charr in lake Úlfljótsvatn are recent immigrants, and if they arrived as juveniles or adults. Newly migrated PL-charr are expected to resemble more the Þingvallavatn PL-charr. The PL-charr in the two lakes did overlap somewhat in shape of several bones, but we observed a pattern where PL-charr from Úlfljótsvatn had more “benthic” bone shape (compared to PL-charr in Þingvallavatn). The possible explanations are: 1) Plasticity, PL-charr feed on snails develop benthic jaw bone shape. 2) Differential survival, majority of the PL-charr come from migrating juveniles from lake Þingvallavatn and only those with more benthic morphology survive to adulthood. 3) and 4) Local adaptation, PL-charr born in lake Úlfljótsvatn (“local” population) have diverged in bones shape through 3) genetics or 4) plasticity. Also, some combination of all two - four scenarios could explain the findings. The similar genetics suggest substantial migration from lake Þingvallavatn, and it is more likely that small fish are pulled by the current in river Efra Sog. If spawning occurs in Úlfljótsvatn, it is possible that the Úlfljótsvatn PL-charr represent a mix of individuals hatched in the lake (“local”) and PL-charr from lake Þingvallavatn (“migrant”). To disentangle this, tracking data on individuals, genetic analyses of many fish in both lakes (including juveniles) or stable isotope analyses might be useful.

Previous experiments rearing Þingvallavatn morphs under different dietary conditions have shown that external morphology (including head measures) responds to benthic vs pelagic dietary treatment (Küttner et al., 2013; Parsons et al., 2010), and also their behavior (Dellinger et al., 2023). Specifically, PL-charr from lake Þingvallavatn reared on a benthic-like diet showed more “benthic-like” head morphology than individuals reared on a pelagic-like diet (Küttner et al., 2013). This agrees with the results on the PL-charr from Úlfljótsvatn (“reared on benthic diet”) appeared to have more “benthic” jaw bone morphology than their genetically identical (Xiao et al., 2026) Þingvallavatn “sister” morph (“reared on pelagic diet”). The Úlfljótsvatn system can be viewed as a natural “rearing experiment” where fish of genetically “pelagic” stock are reared in a novel benthic environment. It was proposed that the benthic morphs in Þingvallavatn are paedomorphic, i.e. retain juvenile characteristic as adults (Skúlason, Noakes, et al., 1989). Therefore, the PL-charr from Úlfljótsvatn may retain more juvenile traits than their Þingvallavatn “sister” morph, because they mainly feed on snails in the benthos. To test this hypothesis an ontogenetic comparison of juvenile PL-charr reared in the two lakes is needed. Currently we only studied adults, and juvenile bone shape is hard to measure because of size (Jónsdóttir, unpublished data). Therefore, we cannot confirm that the PL-charr in Úlfljótsvatn are retaining more juvenile traits than the PL-charr in Þingvallavatn.

Previously it was hypothesized that the PL-charr in Úlfljótsvatn have a lower fitness than their Þingvallavatn “sister” morph (Xiao et al., 2026). Though we did not measure fitness, the data does not support this idea. The Úlfljótsvatn charr are in similar body condition as the Þingvallavatn charr and even grow faster (see above). Reproductive success may be a different story, as phenotypic plasticity is not necessarily adaptive (Ghalambor et al., 2007). Thus, we cannot assume that the apparent phenotypic plasticity means that the PL-charr in the smaller lake are adapting. A better explanation may be that the ecological settings that promote sympatric divergence may be particularly strong in the larger lake Þingvallavatn, but weaker in the smaller more uniform downstream lake Úlfljótsvatn. It is clear that the PL-charr can survive in lake Úlfljótsvatn, but we cannot conclude that the observed change in PL-charr bone morphology is an example of adaptive phenotypic plasticity. It may more likely be due to a response to being forced to feed almost exclusively on snails. It may be that rearing experiments with these morphs, ideally sampled from both lakes and appropriate outgroups, could allow us to disentangle these scenarios.

## Conclusions

Arctic charr is known for showing great phenotypic variation and extensive polymorphism, sometimes referred to as the “charr problem” (Klemetsen, 2010), however why charr show such extensive variation is not truly known. We examined craniofacial variation in putative ancestral and derived populations and confirmed the benthic morphology to be generally derived. The results also uphold a common, though rarely stated assumption, that anadromous charr do not vary much in feeding phenotypes in their natural habitats. It would be interesting to see if that holds in other populations or salmonid taxa. Arctic charr has the ability to respond to environmental changes with body size and external morphology through phenotypic plasticity (Brachmann et al., 2021; Küttner et al., 2013; Parsons et al., 2011). However, we lack estimates of the ancestral plasticity, which may be important for emergence of polymorphism and divergence (West-Eberhard, 2003). Therefore, the capacity for phenotypic plasticity in anadromous charr and populations which have newly colonized freshwaters, could be explored in future studies using rearing experiments, other natural situations like this or transplanted stocks of known origin.

We explored potential phenotypic plasticity in the unique Þingvallavatn-Úlfljótsvatn system focusing on feeding related traits. This system could be further used to study in more detail variation of individual plasticity, the genotype by environment interaction (GxE) and individual by environment interaction (IxE) (Brommer, 2013; Nussey et al., 2007). Such analysis could also explore patterns over multiple bones or traits (like teeth counts) tackling questions about adaptability and plasticity? Could it be that less variable bones serve more basal structural needs, while others may have more evolvability or developmental plasticity, sensitive to prey types? It was proposed that the evolution of a modified pharyngeal jaws allowed the oral jaw in cichlids and labrids to diversify into various different shapes (Liem, 1973; Mabuchi et al., 2007). Studies of the patterns of variation in jaw shape and teeth traits across several groups of salmonids and fishes could provide powerful systems to explore the relationships of modular development, genetic and GxE evolution for the emergence of adaptations.

## Supporting information

All supplementary materials in one file

## Acknowledgments

This study was financially supported by the Icelandic Research Fund, Ph.D. grant no. 2410464-051 to GOJ and grant no. 2511413-051 to AP as well as the Science and Research Fund of South Iceland, Icelandic Genetic Resource Council in Agriculture and the Directorate of Fisheries. We were granted access and got advice from Hörður Hjartarson for sampling in Vífilsdalsá and Gunnar Þorgeirsson for Víðidalsá. We also thank Sigrún Reynisdóttir (Hestvatn), Kristján F. Sveinbjörnsson (Hítarvatn), Ragnar Danner (Brúará) and Guðmundur Steinbach (Gufudalsá) for providing fish. Rafn Sigurðsson we thank for assisting in Víðidalsá. For assistance with sampling and processing of fish from Úlfljótsvatn, Vífilsdalsá and Lambeyrarkvísl we thank Benóny Jónsson, Jóhannes Guðbrandsson, Magnús Jóhannsson and Sigurður Már Einarsson of the Icelandic Marine & Freshwater Research Institute. We also thank Benóný Jónsson for providing the Úlfljótsvatn water temperature data. We thank Ian Dworkin for reviewing and editing the manuscript. Finally, we thank members of the Arctic charr group at the University of Iceland for their help during the project, especially Ruhila Goswami, Laura-Marie von Elm, Samuel Tersigni and Nahal Eskafi for assisting with processing and aging of fish.

