## Supplementary material for "Ancestral and derived feeding morphology during rapid ecological divergence within a species": All supplementary materials in one file

Supplementary material for manuscript entitled: Ancestral and derived feeding morphology during rapid ecological divergence within a species

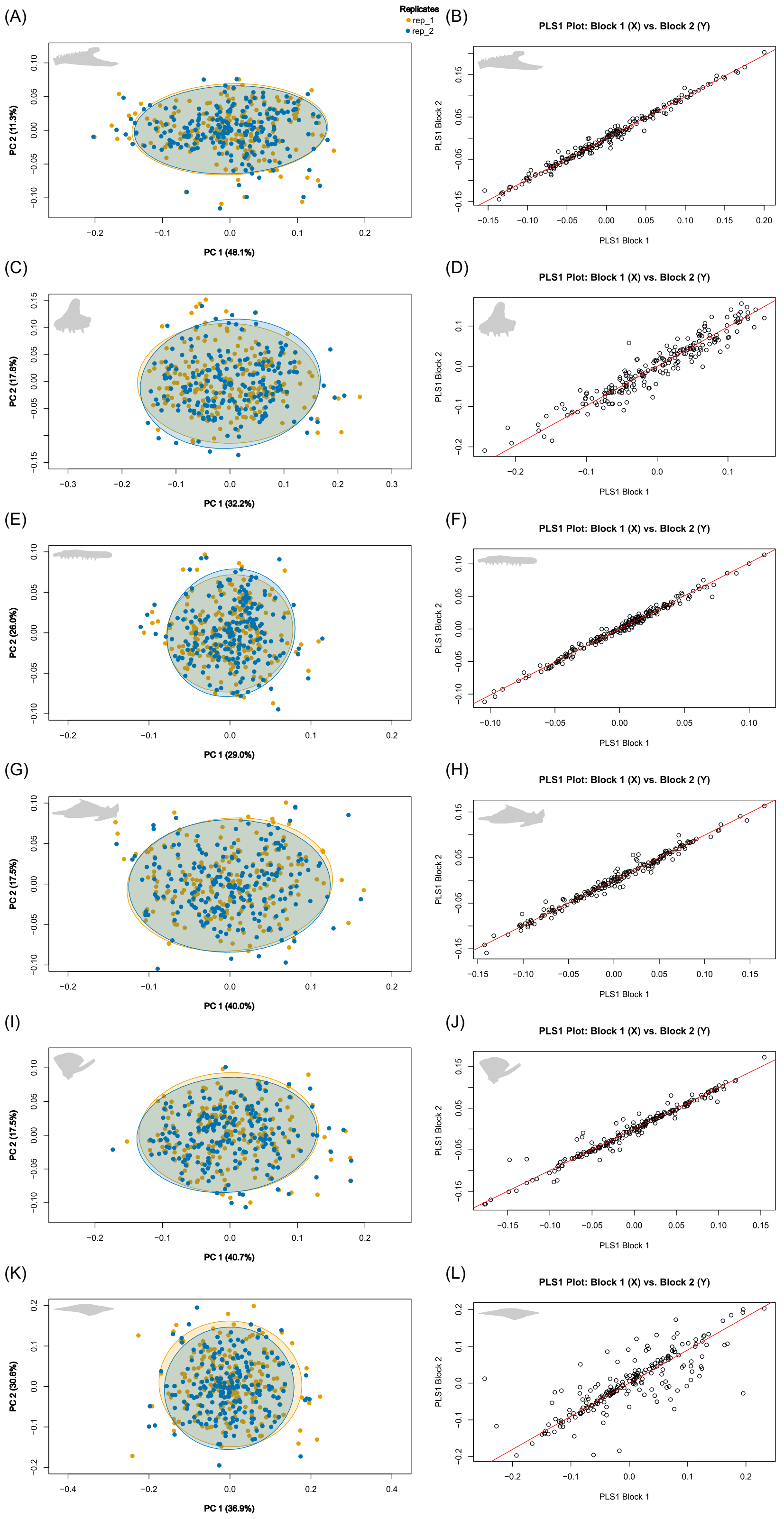


***Supplementary material 1****: On left are PC-plots of specimens showing shape differences between the two replicates (rep1 and rep2), for A) dentary, C) premaxilla, E) maxilla, G)*

*articular-angular, I) quadrate and K) supramaxilla (from top to bottom). Each dot represents an individual and the ellipses represent 95% CI for the distribution by replicates (large dot represents the mean for each replicate in these dimensions). On right are ordination plot (from partial least squares analysis (PLS)) showing the degree of association between replicates for B) dentary, D) premaxilla, F) maxilla, H) articular-angular, J) quadrat and L) supramaxilla (from top to bottom). Block 1 (x-axis) is replicate 1 and Block 2 (y-axis) is replicate 2. For all bones association between the replicates was always significant (p < 0.001) and the Pearson correlation coefficient (r-PLS) was high, dentary: 0.993, premaxilla: 0.952, maxilla: 0.993, articular-angular: 0.99, quadrate: 0.979 and supramaxilla: 0.81.*

***Supplementary material 2****: ANOVA results from tests of the influence of individual variation and measurement error (replicates) effects on shape variation, analyzed with Procrustes ANCOVA. The general intraclass correlation was also calculated by hand and gave these results: dentary: 0.934, premaxilla: 0.829, maxilla: 0.876, articular-angular: 0.871, quadrate: 0.904 and supramaxilla: 0.831.*

| Dataset/Bone | Individuals (id) | | | | Replicates (rep) | | | | Residuals |
|---|---|---|---|---|---|---|---|---|---|
| | $R^2$ | F | Z | P | $R^2$ | F | Z | P | $R^2$ |
| Dentary | 0.967 | 29.34 | 29.63 | ***0.001** | 0.000 | 1.57 | 1.21 | 0.122 | 0.033 |
| Articular-angular | 0.935 | 14.68 | 31.21 | ***0.001** | 0.001 | 3.51 | 1.96 | 0.018 | 0.064 |
| Quadrate | 0.952 | 20.85 | 25.80 | ***0.001** | 0.002 | 10.93 | 5.34 | ***0.001** | 0.046 |
| Maxilla | 0.938 | 16.09 | 24.74 | ***0.001** | 0.004 | 13.54 | 5.60 | ***0.001** | 0.058 |
| Premaxilla | 0.914 | 11.15 | 29.37 | ***0.001** | 0.004 | 9.31 | 5.73 | ***0.001** | 0.082 |
| Supramaxilla | 0.915 | 11.05 | 10.24 | ***0.001** | 0.002 | 4.64 | 2.59 | ***0.005** | 0.083 |

R2 = coefficient of determination; F = F-statistic; Z = Z-statistic; P = P-value, < 0.008 in bold with asterisks.

***Supplementary material 3****: Summary statistics for all Arctic charr groups used in this study. Shows number of individuals (N), average fork length (FL), weight, age and maturity and standard deviation (SD) for length, weight, age and maturity.*

| Population | Sex | N | $FL_{Mean}$(cm)[1] | $Weight_{Mean}$(g) | $Age_{Mean}$ | $Maturity_{mean}$[2] |
|---|---|---|---|---|---|---|
| AN Brúará | F | 3 | 42.53 ± 3.55 | 831.00 ± 186.11 | 5.33 ± 0.58 | NA |
| | M | 3 | 41.50 ± 6.38 | 748.00 ± 120.40 | 5.67 ± 1.15 | NA |
| | All | 6 | 42.02 ± 4.65 | 789.50 ± 147.38 | 5.50 ± 0.84 | NA |
| AN Gufudalsá | F | 11 | 32.98 ± 4.14 | 440.91 ± 215.71 | 4.63 ± 0.67 | 1.00 ± 0.00 |
| | M | 8 | 30.04 ± 4.03 | 322.50 ± 94.68 | 4.43 ± 0.67 | 1.00 ± 0.00 |
| | All | 20 | 32.02 ± 4.31 | 395.50 ± 177.78 | 4.58 ± 0.61 | 1.00 ± 0.00 |
| AN Lambeyrarkvísl | F | 0 | NA | NA | NA | NA |
| | M | 12 | 35.68 ± 3.51 | 543.58 ± 168.47 | 4.17 ± 0.39 | NA |
| | All | 12 | 35.68 ± 3.51 | 543.58 ± 168.47 | 4.17 ± 0.39 | NA |
| AN Vífilsdalsá | F | 4 | 36.38 ± 4.27 | 484.28 ± 210.87 | 4.00 ± 0.00 | 7.00 ± 0.00 |
| | M | 6 | 36.30 ± 4.36 | 527.88 ± 204.03 | 4.33 ± 0.52 | NA |
| | All | 10 | 36.33 ± 4.08 | 510.44 ± 196.10 | 4.20 ± 0.42 | 7.00 ± 0.00 |
| AN Víðidalsá | F | 11 | 40.30 ± 3.83 | 653.91 ± 165.87 | 5.91 ± 0.30 | 6.33 ± 0.58 |
| | M | 7 | 46.86 ± 5.11 | 1145.63 ± 475.60 | 5.86 ± 0.90 | NA |
| | All | 18 | 42.85 ± 5.35 | 845.13 ± 396.06 | 5.89 ± 0.58 | 6.33 ± 0.58 |
| AN | F | 29 | 37.21 ± 5.30 | 568.04 ± 226.58 | 5.10 ± 0.86 | 3.00 ± 2.81 |
| | M | 36 | 37.19 ± 7.04 | 625.93 ± 369.56 | 4.71 ± 0.93 | 1.00 ± 0.00 |
| | All | 66 | 37.20 ± 6.22 | 598.28 ± 310.97 | 4.89 ± 0.90 | 2.27 ± 2.47 |
| Hestvatn | F | 8 | 28.64 ± 7.41 | 206.16 ± 193.90 | 6.13 ± 1.56 | 2.80 ± 0.84 |
| | M | 11 | 32.79 ± 8.81 | 446.24 ± 319.96 | 5.80 ± 1.40 | 2.00 ± 0.82 |

| | | | | | | |
|---|---|---|---|---|---|---|
| | All | 34 | 33.89 ± 8.07 | 513.34 ± 383.23 | 6.30 ± 1.61 | 2.33 ± 0.89 |
| Hítarvatn | F | 7 | 31.29 ± 3.30 | 265.14 ± 90.00 | 6.57 ± 0.53 | NA |
| | M | 1 | 35.10 | 358.00 | 6.00 | NA |
| | All | 8 | 31.76 ± 3.34 | 276.75 ± 89.56 | 6.50 ± 0.53 | NA |
| LB_TH | F | 17 | 42.28 ± 7.01 | 944.29 ± 381.73 | 8.20 ± 1.61 | 5.47 ± 2.32 |
| | M | 32 | 40.73 ± 6.33 | 844.66 ± 388.13 | 8.06 ± 2.20 | 6.06 ± 1.81 |
| | All | 49 | 41.27 ± 6.54 | 879.23 ± 384.91 | 8.11 ± 2.01 | 5.86 ± 2.00 |
| LB_UL | F | 11 | 39.97 ± 3.54 | 928.91 ± 308.15 | 5.20 ± 0.42 | 2.09 ± 0.54 |
| | M | 11 | 36.15 ± 6.03 | 712.76 ± 405.14 | 4.55 ± 1.44 | 1.60 ± 1.58 |
| | All | 26 | 37.24 ± 6.65 | 787.39 ± 399.41 | 4.88 ± 1.01 | 1.86 ± 1.13 |
| PI_TH | F | 14 | 38.25 ± 6.05 | 670.43 ± 337.38 | 8.93 ± 1.73 | 6.79 ± 0.58 |
| | M | 34 | 38.19 ± 8.56 | 780.21 ± 597.28 | 8.61 ± 1.73 | 5.85 ± 1.18 |
| | All | 48 | 38.20 ± 7.84 | 748.19 ± 533.39 | 8.70 ± 1.57 | 6.12 ± 1.12 |
| PI_UL | F | 7 | 42.87 ± 5.58 | 1124.86 ± 477.03 | 5.43 ± 0.98 | 3.29 ± 1.60 |
| | M | 4 | 45.88 ± 4.97 | 1466.50 ± 643.81 | 5.25 ± 0.50 | 1.25 ± 0.50 |
| | All | 11 | 43.96 ± 5.33 | 1249.09 ± 539.07 | 5.36 ± 0.81 | 2.55 ± 1.63 |
| PL_TH | F | 23 | 22.47 ± 3.30 | 106.49 ± 59.01 | 6.87 ± 1.22 | 6.83 ± 0.58 |
| | M | 38 | 16.75 ± 3.31 | 52.90 ± 33.01 | 5.14 ± 1.00 | 5.55 ± 1.16 |
| | All | 61 | 18.91 ± 4.31 | 73.10 ± 51.33 | 5.80 ± 1.38 | 6.03 ± 1.15 |
| PL_UL | F | 10 | 20.96 ± 2.37 | 105.28 ± 37.28 | 4.30 ± 1.06 | 1.00 ± 0.00 |
| | M | 19 | 21.18 ± 3.98 | 112.37 ± 75.62 | 3.90 ± 1.05 | 1.00 ± 0.00 |
| | All | 30 | 21.01 ± 3.44 | 108.39 ± 63.74 | 4.03 ± 1.03 | 1.00 ± 0.00 |
| SB_TH | F | 47 | 16.46 ± 4.36 | 59.77 ± 56.00 | 5.60 ± 1.36 | 5.26 ± 2.26 |
| | M | 33 | 13.79 ± 2.88 | 30.91 ± 23.05 | 5.34 ± 2.04 | 5.67 ± 1.22 |
| | All | 82 | 15.34 ± 3.97 | 47.54 ± 46.87 | 5.46 ± 1.66 | 5.42 ± 1.90 |
| All | F | 173 | 29.02 ± 11.27 | 404.74 ± 421.51 | 6.15 ± 1.77 | 4.72 ± 2.53 |
| | M | 219 | 29.27 ± 12.42 | 457.66 ± 497.42 | 5.97 ± 2.22 | 4.63 ± 2.32 |
| | All | 415 | 29.42 ± 11.85 | 443.13 ± 465.27 | 6.05 ± 2.01 | 4.66 ± 2.42 |

1 FL: Fork length, length from tip of snout to the posterior tip of middle caudal fin ray

2 Maturity: Estimation of maturity, from a scale of 1 to 7 (running).

Log: natural logarithmic transformation.

***Supplementary material 4****: Anova results when testing influence of population and sex effects on length (cm FL), log$_e$ weight (g), age (years) and maturity (scale 1-7). Hítarvatn fish included when testing for effects on weight and age but excluded when testing effects on length and maturity. AN-charr treated as five separate populations.*

| Variable | | Population | | Sex | | Population x Sex Interaction | |
|---|---|---|---|---|---|---|---|
| | N | F | P | F | P | F | P |
| Fork length (cm) | 383 | 158.22 | ***<< 0.001** | 20.14 | ***<< 0.001** | 3.87 | ***<< 0.001** |
| Log$_e$ weight (g) | 390 | 149.32 | ***<< 0.001** | 11.97 | ***<< 0.001** | 3.47 | ***<< 0.001** |
| Age | 381 | 32.82 | ***<< 0.001** | 21.00 | ***0.001** | 1.24 | 0.256 |
| Maturity | 334 | 63.76 | ***<< 0.001** | 3.49 | 0.063 | 2.88 | ***0.004** |

F = F-statistic, P = P – value, <0.05 in bold, with asterisk

***Supplementary material 5****: Post hoc results when testing influence of morph and sex effects on length (cm FL), $log_e$ weight (g) age (years) and maturity (scale 1-7). Hítarvatn fish included when testing for effects on weight and age but excluded when testing effects on length and maturity. AN-charr treated as five separate populations.*

| Variable | System | Pairs | Diff | LCL | UCL | P-value |
|---|---|---|---|---|---|---|
| Fork length (cm) | AN populations | AN_Gufudalsá:AN_Brúará | -0.284 | -0.572 | 0.004 | 0.058 |
| | | AN_Lambeyrarkvísl:AN_Brúará | -0.163 | -0.470 | 0.145 | 0.865 |
| | | AN_Víðidalsá:AN_Brúará | 0.018 | -0.272 | 0.308 | 1.000 |
| | | AN_Vífilsdalsá:AN_Brúará | -0.146 | -0.463 | 0.172 | 0.948 |
| | | AN_Lambeyrarkvísl:AN_Gufudalsá | 0.121 | -0.106 | 0.348 | 0.858 |
| | | AN_Víðidalsá:AN_Gufudalsá | 0.302 | 0.099 | 0.504 | *<<**0.001** |
| | | AN_Vífilsdalsá:AN_Gufudalsá | 0.138 | -0.102 | 0.378 | 0.786 |
| | | AN_Víðidalsá:AN_Lambeyrarkvísl | 0.180 | -0.049 | 0.410 | 0.298 |
| | | AN_Vífilsdalsá:AN_Lambeyrarkvísl | 0.017 | -0.246 | 0.280 | 1.000 |
| | | AN_Vífilsdalsá:AN_Víðidalsá | -0.164 | -0.406 | 0.079 | 0.555 |
| | Brúará to lake morphs | Hest:AN_Brúará | -0.330 | -0.618 | -0.042 | ***0.010** |
| | | LB_TH:AN_Brúará | -0.026 | -0.292 | 0.240 | 1.000 |
| | | LB_UL:AN_Brúará | -0.103 | -0.386 | 0.180 | 0.992 |
| | | PI_TH:AN_Brúará | -0.112 | -0.378 | 0.155 | 0.974 |
| | | PI_UL:AN_Brúará | 0.044 | -0.268 | 0.356 | 1.000 |
| | | PL_TH:AN_Brúará | -0.819 | -1.082 | -0.556 | *<<**0.001** |
| | | PL_UL:AN_Brúará | -0.695 | -0.971 | -0.420 | *<<**0.001** |
| | | SB_TH:AN_Brúará | -1.031 | -1.291 | -0.771 | *<<**0.001** |
| | Gufudalsá to lake morphs | Hest:AN_Gufudalsá | -0.047 | -0.246 | 0.153 | 1.000 |
| | | LB_TH:AN_Gufudalsá | 0.258 | 0.092 | 0.424 | *<<**0.001** |
| | | LB_UL:AN_Gufudalsá | 0.181 | -0.012 | 0.373 | 0.091 |
| | | PI_TH:AN_Gufudalsá | 0.172 | 0.006 | 0.339 | ***0.035** |
| | | PI_UL:AN_Gufudalsá | 0.328 | 0.095 | 0.561 | *<<**0.001** |
| | | PL_TH:AN_Gufudalsá | -0.535 | -0.697 | -0.374 | *<<**0.001** |
| | | PL_UL:AN_Gufudalsá | -0.412 | -0.593 | -0.230 | *<<**0.001** |
| | | SB_TH:AN_Gufudalsá | -0.747 | -0.904 | -0.590 | *<<**0.001** |
| | Lambeyrarkvísl to lake morphs | Hest:AN_Lambeyrarkvísl | -0.168 | -0.394 | 0.059 | 0.400 |
| | | LB_TH:AN_Lambeyrarkvísl | 0.137 | -0.061 | 0.335 | 0.515 |
| | | LB_UL:AN_Lambeyrarkvísl | 0.060 | -0.161 | 0.280 | 1.000 |
| | | PI_TH:AN_Lambeyrarkvísl | 0.051 | -0.147 | 0.250 | 1.000 |
| | | PI_UL:AN_Lambeyrarkvísl | 0.207 | -0.050 | 0.463 | 0.265 |
| | | PL_TH:AN_Lambeyrarkvísl | -0.656 | -0.850 | -0.462 | *<<**0.001** |
| | | PL_UL:AN_Lambeyrarkvísl | -0.533 | -0.744 | -0.322 | *<<**0.001** |
| | | SB_TH:AN_Lambeyrarkvísl | -0.868 | -1.058 | -0.678 | *<<**0.001** |
| | Víðidalsá to lake morphs | Hest:AN_Víðidalsá | -0.348 | -0.550 | -0.146 | *<<**0.001** |
| | | LB_TH:AN_Víðidalsá | -0.044 | -0.213 | 0.126 | 1.000 |
| | | LB_UL:AN_Víðidalsá | -0.121 | -0.316 | 0.074 | 0.689 |
| | | PI_TH:AN_Víðidalsá | -0.129 | -0.299 | 0.041 | 0.354 |
| | | PI_UL:AN_Víðidalsá | 0.026 | -0.209 | 0.261 | 1.000 |
| | | PL_TH:AN_Víðidalsá | -0.837 | -1.002 | -0.672 | *<<**0.001** |
| | | PL_UL:AN_Víðidalsá | -0.713 | -0.898 | -0.529 | *<<**0.001** |
| | | SB_TH:AN_Víðidalsá | -1.049 | -1.209 | -0.888 | *<<**0.001** |
| | Vífisdalsá to lake morphs | Hest:AN_Vífilsdalsá | -0.184 | -0.425 | 0.056 | 0.338 |
| | | LB_TH:AN_Vífilsdalsá | 0.120 | -0.093 | 0.333 | 0.808 |
| | | LB_UL:AN_Vífilsdalsá | 0.043 | -0.192 | 0.277 | 1.000 |
| | | PI_TH:AN_Vífilsdalsá | 0.034 | -0.179 | 0.248 | 1.000 |
| | | PI_UL:AN_Vífilsdalsá | 0.190 | -0.079 | 0.458 | 0.477 |
| | | PL_TH:AN_Vífilsdalsá | -0.673 | -0.883 | -0.463 | *<<**0.001** |
| | | PL_UL:AN_Vífilsdalsá | -0.550 | -0.775 | -0.324 | *<<**0.001** |
| | | SB_TH:AN_Vífilsdalsá | -0.885 | -1.091 | -0.679 | *<<**0.001** |
| | Hestvatn | LB_TH:Hest | 0.305 | 0.138 | 0.471 | *<<**0.001** |
| | | LB_UL:Hest | 0.227 | 0.035 | 0.420 | ***0.007** |
| | | PI_TH:Hest | 0.219 | 0.052 | 0.386 | ***0.001** |
| | | PI_UL:Hest | 0.374 | 0.141 | 0.607 | *<<**0.001** |

| | | | | | | |
|---|---|---|---|---|---|---|
| | | PL_TH:Hest | -0.489 | -0.650 | -0.327 | *<<0.001 |
| | | PL_UL:Hest | -0.365 | -0.547 | -0.184 | *<<0.001 |
| | | SB_TH:Hest | -0.700 | -0.857 | -0.544 | *<<0.001 |
| | Þingvallavatn | PI_TH:LB_TH | -0.086 | -0.211 | 0.039 | 0.528 |
| | | PL_TH:LB_TH | -0.793 | -0.911 | -0.675 | *<<0.001 |
| | | SB_TH:LB_TH | -1.005 | -1.116 | -0.893 | *<<0.001 |
| | | PL_TH:PI_TH | -0.707 | -0.826 | -0.589 | *<<0.001 |
| | | SB_TH:PI_TH | -0.919 | -1.032 | -0.807 | *<<0.001 |
| | | SB_TH:PL_TH | -0.212 | -0.316 | -0.107 | *<<0.001 |
| | Úlfljótsvatn | PI_UL:LB_UL | 0.147 | -0.080 | 0.374 | 0.620 |
| | | PL_UL:LB_UL | -0.592 | -0.766 | -0.418 | *<<0.001 |
| | | PL_UL:PI_UL | -0.739 | -0.957 | -0.522 | *<<0.001 |
| | Across Þingvallavatn and Úlfljótsvatn | LB_UL:LB_TH | -0.077 | -0.235 | 0.080 | 0.918 |
| | | PI_UL:LB_TH | 0.070 | -0.135 | 0.275 | 0.996 |
| | | PL_UL:LB_TH | -0.670 | -0.814 | -0.526 | *<<0.001 |
| | | PI_TH:LB_UL | -0.008 | -0.167 | 0.150 | 1.000 |
| | | PL_TH:LB_UL | -0.716 | -0.869 | -0.563 | *<<0.001 |
| | | SB_TH:LB_UL | -0.928 | -1.076 | -0.780 | *<<0.001 |
| | | PI_UL:PI_TH | 0.155 | -0.050 | 0.361 | 0.363 |
| | | PL_UL:PI_TH | -0.584 | -0.729 | -0.439 | *<<0.001 |
| | | PL_TH:PI_UL | -0.863 | -1.064 | -0.661 | *<<0.001 |
| | | SB_TH:PI_UL | -1.075 | -1.272 | -0.877 | *<<0.001 |
| | | PL_UL:PL_TH | 0.123 | -0.015 | 0.262 | 0.138 |
| | | SB_TH:PL_UL | -0.335 | -0.469 | -0.202 | *<<0.001 |
| $Log_e$ weight (g) | AN populations | AN_Gufudalsá:AN_Brúará | -0.769 | -1.672 | 0.134 | 0.193 |
| | | AN_Lambeyrarkvísl:AN_Brúará | -0.398 | -1.363 | 0.566 | 0.982 |
| | | AN_Víðidalsá:AN_Brúará | 0.004 | -0.905 | 0.913 | 1.000 |
| | | AN_Vífilsdalsá:AN_Brúará | -0.484 | -1.480 | 0.513 | 0.935 |
| | | AN_Lambeyrarkvísl:AN_Gufudalsá | 0.371 | -0.340 | 1.082 | 0.892 |
| | | AN_Víðidalsá:AN_Gufudalsá | 0.773 | 0.139 | 1.408 | *0.004 |
| | | AN_Vífilsdalsá:AN_Gufudalsá | 0.286 | -0.468 | 1.039 | 0.992 |
| | | AN_Víðidalsá:AN_Lambeyrarkvísl | 0.402 | -0.316 | 1.121 | 0.831 |
| | | AN_Vífilsdalsá:AN_Lambeyrarkvísl | -0.085 | -0.911 | 0.741 | 1.000 |
| | | AN_Vífilsdalsá:AN_Víðidalsá | -0.488 | -1.248 | 0.273 | 0.656 |
| | Brúará to lake morphs | Hest:AN_Brúará | -1.140 | -2.049 | -0.231 | *0.002 |
| | | Hit:AN_Brúará | -1.074 | -2.116 | -0.032 | *0.036 |
| | | LB_TH:AN_Brúará | 0.011 | -0.823 | 0.846 | 1.000 |
| | | LB_UL:AN_Brúará | -0.064 | -0.952 | 0.824 | 1.000 |
| | | PI_TH:AN_Brúará | -0.274 | -1.110 | 0.561 | 0.998 |
| | | PI_UL:AN_Brúará | 0.399 | -0.580 | 1.378 | 0.984 |
| | | PL_TH:AN_Brúará | -2.567 | -3.392 | -1.742 | *<<0.001 |
| | | PL_UL:AN_Brúará | -2.074 | -2.939 | -1.209 | *<<0.001 |
| | | SB_TH:AN_Brúará | -3.087 | -3.904 | -2.271 | *<<0.001 |
| | Gufudalsá to lake morphs | Hest:AN_Gufudalsá | -0.371 | -1.005 | 0.264 | 0.783 |
| | | Hit:AN_Gufudalsá | -0.305 | -1.118 | 0.508 | 0.993 |
| | | LB_TH:AN_Gufudalsá | 0.781 | 0.259 | 1.302 | *<<0.001 |
| | | LB_UL:AN_Gufudalsá | 0.705 | 0.101 | 1.309 | *0.007 |
| | | PI_TH:AN_Gufudalsá | 0.495 | -0.028 | 1.018 | 0.085 |
| | | PI_UL:AN_Gufudalsá | 1.168 | 0.437 | 1.899 | *<<0.001 |
| | | PL_TH:AN_Gufudalsá | -1.798 | -2.305 | -1.291 | *<<0.001 |
| | | PL_UL:AN_Gufudalsá | -1.305 | -1.874 | -0.736 | *<<0.001 |
| | | SB_TH:AN_Gufudalsá | -2.318 | -2.810 | -1.826 | *<<0.001 |
| | Lambeyrarkvísl to lake morphs | Hest:AN_Lambeyrarkvísl | -0.742 | -1.461 | -0.023 | *0.036 |
| | | Hit:AN_Lambeyrarkvísl | -0.676 | -1.556 | 0.205 | 0.350 |
| | | LB_TH:AN_Lambeyrarkvísl | 0.410 | -0.212 | 1.031 | 0.610 |
| | | LB_UL:AN_Lambeyrarkvísl | 0.334 | -0.358 | 1.027 | 0.937 |
| | | PI_TH:AN_Lambeyrarkvísl | 0.124 | -0.499 | 0.746 | 1.000 |
| | | PI_UL:AN_Lambeyrarkvísl | 0.797 | -0.008 | 1.603 | 0.055 |
| | | PL_TH:AN_Lambeyrarkvísl | -2.169 | -2.778 | -1.560 | *<<0.001 |
| | | PL_UL:AN_Lambeyrarkvísl | -1.676 | -2.338 | -1.014 | *<<0.001 |
| | | SB_TH:AN_Lambeyrarkvísl | -2.689 | -3.286 | -2.092 | *<<0.001 |

| | | | | | | |
|---|---|---|---|---|---|---|
| | Víðidalsá to lake morphs | Hest:AN_Víðidalsá | -1.144 | -1.787 | -0.501 | *<<**0.001** |
| | | Hit:AN_Víðidalsá | -1.078 | -1.898 | -0.258 | ***0.001** |
| | | LB_TH:AN_Víðidalsá | 0.007 | -0.524 | 0.539 | 1.000 |
| | | LB_UL:AN_Víðidalsá | -0.068 | -0.681 | 0.545 | 1.000 |
| | | PI_TH:AN_Víðidalsá | -0.279 | -0.812 | 0.255 | 0.890 |
| | | PI_UL:AN_Víðidalsá | 0.395 | -0.343 | 1.133 | 0.872 |
| | | PL_TH:AN_Víðidalsá | -2.571 | -3.089 | -2.054 | *<<**0.001** |
| | | PL_UL:AN_Víðidalsá | -2.078 | -2.657 | -1.500 | *<<**0.001** |
| | | SB_TH:AN_Víðidalsá | -3.091 | -3.595 | -2.588 | *<<**0.001** |
| | Vífisdalsá to lake morphs | Hest:AN_Vífilsdalsá | -0.656 | -1.417 | 0.104 | 0.177 |
| | | Hit:AN_Vífilsdalsá | -0.590 | -1.505 | 0.325 | 0.645 |
| | | LB_TH:AN_Vífilsdalsá | 0.495 | -0.174 | 1.164 | 0.413 |
| | | LB_UL:AN_Vífilsdalsá | 0.420 | -0.316 | 1.155 | 0.811 |
| | | PI_TH:AN_Vífilsdalsá | 0.209 | -0.461 | 0.880 | 0.999 |
| | | PI_UL:AN_Vífilsdalsá | 0.883 | 0.040 | 1.725 | ***0.030** |
| | | PL_TH:AN_Vífilsdalsá | -2.083 | -2.742 | -1.425 | *<<**0.001** |
| | | PL_UL:AN_Vífilsdalsá | -1.591 | -2.298 | -0.883 | *<<**0.001** |
| | | SB_TH:AN_Vífilsdalsá | -2.604 | -3.251 | -1.957 | *<<**0.001** |
| | Hestvatn and Hítarvatn | Hit:Hest | 0.066 | -0.754 | 0.886 | 1.000 |
| | | LB_TH:Hest | 1.151 | 0.620 | 1.683 | *<<**0.001** |
| | | LB_UL:Hest | 1.076 | 0.463 | 1.689 | *<<**0.001** |
| | | PI_TH:Hest | 0.865 | 0.332 | 1.399 | *<<**0.001** |
| | | PI_UL:Hest | 1.539 | 0.801 | 2.277 | *<<**0.001** |
| | | PL_TH:Hest | -1.427 | -1.944 | -0.910 | *<<**0.001** |
| | | PL_UL:Hest | -0.934 | -1.513 | -0.356 | *<<**0.001** |
| | | SB_TH:Hest | -1.947 | -2.450 | -1.444 | *<<**0.001** |
| | | LB_TH:Hit | 1.085 | 0.350 | 1.821 | *<<**0.001** |
| | | LB_UL:Hit | 1.010 | 0.213 | 1.806 | ***0.002** |
| | | PI_TH:Hit | 0.799 | 0.063 | 1.536 | ***0.020** |
| | | PI_UL:Hit | 1.473 | 0.577 | 2.369 | *<<**0.001** |
| | | PL_TH:Hit | -1.493 | -2.218 | -0.768 | *<<**0.001** |
| | | PL_UL:Hit | -1.000 | -1.771 | -0.230 | ***0.001** |
| | | SB_TH:Hit | -2.013 | -2.729 | -1.298 | *<<**0.001** |
| | Þingvallavatn | PI_TH:LB_TH | -0.286 | -0.678 | 0.106 | 0.436 |
| | | PL_TH:LB_TH | -2.578 | -2.949 | -2.208 | *<<**0.001** |
| | | SB_TH:LB_TH | -3.099 | -3.449 | -2.749 | *<<**0.001** |
| | | PL_TH:PI_TH | -2.293 | -2.665 | -1.920 | *<<**0.001** |
| | | SB_TH:PI_TH | -2.813 | -3.165 | -2.461 | *<<**0.001** |
| | | SB_TH:PL_TH | -0.520 | -0.848 | -0.192 | *<<**0.001** |
| | Úlfljótsvatn | PI_UL:LB_UL | 0.463 | -0.249 | 1.175 | 0.633 |
| | | PL_UL:LB_UL | -2.010 | -2.556 | -1.465 | *<<**0.001** |
| | | PL_UL:PI_UL | -2.473 | -3.156 | -1.790 | *<<**0.001** |
| | Across Þingvallavatn and Úlfljótsvatn | LB_UL:LB_TH | -0.075 | -0.571 | 0.420 | 1.000 |
| | | PI_UL:LB_TH | 0.388 | -0.256 | 1.031 | 0.745 |
| | | PL_UL:LB_TH | -2.086 | -2.538 | -1.634 | *<<**0.001** |
| | | PI_TH:LB_UL | -0.210 | -0.707 | 0.286 | 0.978 |
| | | PL_TH:LB_UL | -2.503 | -2.983 | -2.023 | *<<**0.001** |
| | | SB_TH:LB_UL | -3.023 | -3.488 | -2.559 | *<<**0.001** |
| | | PI_UL:PI_TH | 0.674 | 0.029 | 1.318 | ***0.031** |
| | | PL_UL:PI_TH | -1.800 | -2.254 | -1.346 | *<<**0.001** |
| | | PL_TH:PI_UL | -2.966 | -3.598 | -2.334 | *<<**0.001** |
| | | SB_TH:PI_UL | -3.486 | -4.107 | -2.866 | *<<**0.001** |
| | | PL_UL:PL_TH | 0.493 | 0.058 | 0.928 | ***0.011** |
| | | SB_TH:PL_UL | -1.013 | -1.431 | -0.595 | *<<**0.001** |
| Age | AN populations | AN_Gufudalsá:AN_Brúará | -0.944 | -3.166 | 1.277 | 0.977 |
| | | AN_Lambeyrarkvísl:AN_Brúará | -1.333 | -3.690 | 1.023 | 0.819 |
| | | AN_Víðidalsá:AN_Brúará | 0.389 | -1.833 | 2.610 | 1.000 |
| | | AN_Vífilsdalsá:AN_Brúará | -1.300 | -3.734 | 1.134 | 0.873 |
| | | AN_Lambeyrarkvísl:AN_Gufudalsá | -0.389 | -2.145 | 1.367 | 1.000 |
| | | AN_Víðidalsá:AN_Gufudalsá | 1.333 | -0.238 | 2.904 | 0.197 |
| | | AN_Vífilsdalsá:AN_Gufudalsá | -0.356 | -2.214 | 1.503 | 1.000 |

| | | | | | | |
|---|---|---|---|---|---|---|
| | | AN_Víðidalsá:AN_Lambeyrarkvísl | 1.722 | -0.034 | 3.479 | 0.061 |
| | | AN_Vífilsdalsá:AN_Lambeyrarkvísl | 0.033 | -1.984 | 2.051 | 1.000 |
| | | AN_Vífilsdalsá:AN_Víðidalsá | -1.689 | -3.548 | 0.170 | 0.120 |
| | Brúará to lake morphs | Hest:AN_Brúará | 0.444 | -1.777 | 2.666 | 1.000 |
| | | Hit:AN_Brúará | 1.000 | -1.545 | 3.545 | 0.988 |
| | | LB_TH:AN_Brúará | 2.606 | 0.563 | 4.649 | *0.002 |
| | | LB_UL:AN_Brúará | -0.643 | -2.824 | 1.539 | 0.999 |
| | | PI_TH:AN_Brúará | 3.202 | 1.159 | 5.245 | *<<0.001 |
| | | PI_UL:AN_Brúará | -0.136 | -2.528 | 2.255 | 1.000 |
| | | PL_TH:AN_Brúará | 0.300 | -1.718 | 2.318 | 1.000 |
| | | PL_UL:AN_Brúará | -1.466 | -3.579 | 0.648 | 0.525 |
| | | SB_TH:AN_Brúará | -0.006 | -2.004 | 1.991 | 1.000 |
| | Gufudalsá to lake morphs | Hest:AN_Gufudalsá | 1.389 | -0.182 | 2.960 | 0.148 |
| | | Hit:AN_Gufudalsá | 1.944 | -0.058 | 3.947 | 0.067 |
| | | LB_TH:AN_Gufudalsá | 3.551 | 2.245 | 4.857 | *<<0.001 |
| | | LB_UL:AN_Gufudalsá | 0.302 | -1.212 | 1.815 | 1.000 |
| | | PI_TH:AN_Gufudalsá | 4.147 | 2.840 | 5.453 | *<<0.001 |
| | | PI_UL:AN_Gufudalsá | 0.808 | -0.995 | 2.612 | 0.965 |
| | | PL_TH:AN_Gufudalsá | 1.244 | -0.022 | 2.511 | 0.060 |
| | | PL_UL:AN_Gufudalsá | -0.521 | -1.935 | 0.893 | 0.994 |
| | | SB_TH:AN_Gufudalsá | 0.938 | -0.296 | 2.172 | 0.365 |
| | Lambeyrarkvísl to lake morphs | Hest:AN_Lambeyrarkvísl | 1.778 | 0.021 | 3.534 | *0.044 |
| | | Hit:AN_Lambeyrarkvísl | 2.333 | 0.182 | 4.484 | *0.020 |
| | | LB_TH:AN_Lambeyrarkvísl | 3.940 | 2.415 | 5.464 | *<<0.001 |
| | | LB_UL:AN_Lambeyrarkvísl | 0.690 | -1.015 | 2.396 | 0.985 |
| | | PI_TH:AN_Lambeyrarkvísl | 4.535 | 3.011 | 6.060 | *<<0.001 |
| | | PI_UL:AN_Lambeyrarkvísl | 1.197 | -0.770 | 3.164 | 0.731 |
| | | PL_TH:AN_Lambeyrarkvísl | 1.633 | 0.143 | 3.124 | *0.017 |
| | | PL_UL:AN_Lambeyrarkvísl | -0.132 | -1.750 | 1.485 | 1.000 |
| | | SB_TH:AN_Lambeyrarkvísl | 1.327 | -0.136 | 2.789 | 0.122 |
| | Víðidalsá to lake morphs | Hest:AN_Víðidalsá | 0.056 | -1.515 | 1.626 | 1.000 |
| | | Hit:AN_Víðidalsá | 0.611 | -1.391 | 2.614 | 0.999 |
| | | LB_TH:AN_Víðidalsá | 2.217 | 0.911 | 3.524 | *<<0.001 |
| | | LB_UL:AN_Víðidalsá | -1.032 | -2.545 | 0.482 | 0.555 |
| | | PI_TH:AN_Víðidalsá | 2.813 | 1.507 | 4.120 | *<<0.001 |
| | | PI_UL:AN_Víðidalsá | -0.525 | -2.329 | 1.278 | 0.999 |
| | | PL_TH:AN_Víðidalsá | -0.089 | -1.355 | 1.178 | 1.000 |
| | | PL_UL:AN_Víðidalsá | -1.854 | -3.268 | -0.440 | *0.001 |
| | | SB_TH:AN_Víðidalsá | -0.395 | -1.629 | 0.838 | 0.998 |
| | Vífisdalsá to lake morphs | Hest:AN_Vífilsdalsá | 1.744 | -0.114 | 3.603 | 0.091 |
| | | Hit:AN_Vífilsdalsá | 2.300 | 0.065 | 4.535 | *0.037 |
| | | LB_TH:AN_Vífilsdalsá | 3.906 | 2.265 | 5.548 | *<<0.001 |
| | | LB_UL:AN_Vífilsdalsá | 0.657 | -1.154 | 2.468 | 0.994 |
| | | PI_TH:AN_Vífilsdalsá | 4.502 | 2.861 | 6.143 | *<<0.001 |
| | | PI_UL:AN_Vífilsdalsá | 1.164 | -0.895 | 3.223 | 0.821 |
| | | PL_TH:AN_Vífilsdalsá | 1.600 | -0.010 | 3.210 | 0.053 |
| | | PL_UL:AN_Vífilsdalsá | -0.166 | -1.894 | 1.563 | 1.000 |
| | | SB_TH:AN_Vífilsdalsá | 1.294 | -0.291 | 2.878 | 0.251 |
| | Hestvatn and Hítarvatn | Hit:Hest | 0.556 | -1.447 | 2.558 | 1.000 |
| | | LB_TH:Hest | 2.162 | 0.856 | 3.468 | *<<0.001 |
| | | LB_UL:Hest | -1.087 | -2.601 | 0.426 | 0.464 |
| | | PI_TH:Hest | 2.758 | 1.451 | 4.064 | *<<0.001 |
| | | PI_UL:Hest | -0.581 | -2.384 | 1.223 | 0.998 |
| | | PL_TH:Hest | -0.144 | -1.411 | 1.122 | 1.000 |
| | | PL_UL:Hest | -1.910 | -3.324 | -0.496 | *0.001 |
| | | SB_TH:Hest | -0.451 | -1.685 | 0.783 | 0.994 |
| | | LB_TH:Hit | 1.606 | -0.196 | 3.409 | 0.140 |
| | | LB_UL:Hit | -1.643 | -3.601 | 0.315 | 0.213 |
| | | PI_TH:Hit | 2.202 | 0.400 | 4.005 | *0.004 |
| | | PI_UL:Hit | -1.136 | -3.326 | 1.053 | 0.895 |
| | | PL_TH:Hit | -0.700 | -2.474 | 1.074 | 0.988 |

| | | | | | | |
|---|---|---|---|---|---|---|
| | | PL_UL:Hit | -2.466 | -4.348 | -0.584 | *0.001 |
| | | SB_TH:Hit | -1.006 | -2.757 | 0.744 | 0.802 |
| | Þingvallavatn | PI_TH:LB_TH | 0.596 | -0.376 | 1.568 | 0.722 |
| | | PL_TH:LB_TH | -2.306 | -3.224 | -1.388 | *<<0.001 |
| | | SB_TH:LB_TH | -2.613 | -3.485 | -1.741 | *<<0.001 |
| | | PL_TH:PI_TH | -2.902 | -3.820 | -1.984 | *<<0.001 |
| | | SB_TH:PI_TH | -3.209 | -4.081 | -2.336 | *<<0.001 |
| | | SB_TH:PL_TH | -0.306 | -1.118 | 0.505 | 0.992 |
| | Úlfljótsvatn | PI_UL:LB_UL | 0.506 | -1.248 | 2.261 | 0.999 |
| | | PL_UL:LB_UL | -0.823 | -2.173 | 0.528 | 0.730 |
| | | PL_UL:PI_UL | -1.329 | -2.998 | 0.340 | 0.289 |
| | Across Þingvallavatn and Úlfljótsvatn | LB_UL:LB_TH | -3.249 | -4.486 | -2.012 | *<<0.001 |
| | | PI_UL:LB_TH | -2.743 | -4.321 | -1.164 | *<<0.001 |
| | | PL_UL:LB_TH | -4.072 | -5.185 | -2.959 | *<<0.001 |
| | | PI_TH:LB_UL | 3.845 | 2.608 | 5.082 | *<<0.001 |
| | | PL_TH:LB_UL | 0.943 | -0.252 | 2.138 | 0.304 |
| | | SB_TH:LB_UL | 0.636 | -0.524 | 1.797 | 0.850 |
| | | PI_UL:PI_TH | -3.338 | -4.917 | -1.760 | *<<0.001 |
| | | PL_UL:PI_TH | -4.668 | -5.780 | -3.555 | *<<0.001 |
| | | PL_TH:PI_UL | 0.436 | -1.109 | 1.982 | 1.000 |
| | | SB_TH:PI_UL | 0.130 | -1.389 | 1.649 | 1.000 |
| | | PL_UL:PL_TH | -1.766 | -2.831 | -0.700 | *<<0.001 |
| | | SB_TH:PL_UL | 1.459 | 0.432 | 2.486 | *<<0.001 |
| Maturity | AN populations | AN_Víðidalsá:AN_Gufudalsá | 5.333 | 2.492 | 8.175 | *<<0.001 |
| | | AN_Vífilsdalsá:AN_Gufudalsá | 6.000 | 3.159 | 8.841 | *<<0.001 |
| | | AN_Vífilsdalsá:AN_Víðidalsá | 0.667 | -3.068 | 4.401 | 1.000 |
| | Gufudalsá to lake morphs | Hest:AN_Gufudalsá | 1.333 | -0.353 | 3.020 | 0.273 |
| | | LB_TH:AN_Gufudalsá | 4.857 | 3.621 | 6.093 | *<<0.001 |
| | | LB_UL:AN_Gufudalsá | 0.857 | -0.591 | 2.305 | 0.704 |
| | | PI_TH:AN_Gufudalsá | 5.125 | 3.885 | 6.365 | *<<0.001 |
| | | PI_UL:AN_Gufudalsá | 1.545 | -0.187 | 3.278 | 0.130 |
| | | PL_TH:AN_Gufudalsá | 5.033 | 3.831 | 6.234 | *<<0.001 |
| | | PL_UL:AN_Gufudalsá | 0.000 | -1.359 | 1.359 | 1.000 |
| | | SB_TH:AN_Gufudalsá | 4.425 | 3.258 | 5.592 | *<<0.001 |
| | Víðidalsá to lake morphs | Hest:AN_Víðidalsá | -4.000 | -6.952 | -1.048 | *0.001 |
| | | LB_TH:AN_Víðidalsá | -0.476 | -3.196 | 2.244 | 1.000 |
| | | LB_UL:AN_Víðidalsá | -4.476 | -7.299 | -1.653 | *<<0.001 |
| | | PI_TH:AN_Víðidalsá | -0.208 | -2.930 | 2.513 | 1.000 |
| | | PI_UL:AN_Víðidalsá | -3.788 | -6.767 | -0.809 | *0.002 |
| | | PL_TH:AN_Víðidalsá | -0.301 | -3.005 | 2.404 | 1.000 |
| | | PL_UL:AN_Víðidalsá | -5.333 | -8.112 | -2.555 | *<<0.001 |
| | | SB_TH:AN_Víðidalsá | -0.908 | -3.598 | 1.781 | 0.991 |
| | Vífisdalsá to lake morphs | Hest:AN_Vífilsdalsá | -4.667 | -7.619 | -1.714 | *<<0.001 |
| | | LB_TH:AN_Vífilsdalsá | -1.143 | -3.863 | 1.577 | 0.956 |
| | | LB_UL:AN_Vífilsdalsá | -5.143 | -7.966 | -2.320 | *<<0.001 |
| | | PI_TH:AN_Vífilsdalsá | -0.875 | -3.597 | 1.847 | 0.994 |
| | | PI_UL:AN_Vífilsdalsá | -4.455 | -7.433 | -1.476 | *<<0.001 |
| | | PL_TH:AN_Vífilsdalsá | -0.967 | -3.672 | 1.737 | 0.986 |
| | | PL_UL:AN_Vífilsdalsá | -6.000 | -8.778 | -3.222 | *<<0.001 |
| | | SB_TH:AN_Vífilsdalsá | -1.575 | -4.265 | 1.115 | 0.718 |
| | Hestvatn | LB_TH:Hest | 3.524 | 2.051 | 4.997 | *<<0.001 |
| | | LB_UL:Hest | -0.476 | -2.131 | 1.179 | 0.998 |
| | | PI_TH:Hest | 3.792 | 2.316 | 5.268 | *<<0.001 |
| | | PI_UL:Hest | 0.212 | -1.697 | 2.121 | 1.000 |
| | | PL_TH:Hest | 3.699 | 2.255 | 5.144 | *<<0.001 |
| | | PL_UL:Hest | -1.333 | -2.911 | 0.245 | 0.187 |
| | | SB_TH:Hest | 3.092 | 1.676 | 4.507 | *<<0.001 |
| | Þingvallavatn | PI_TH:LB_TH | 0.268 | -0.661 | 1.197 | 0.998 |
| | | PL_TH:LB_TH | 0.176 | -0.702 | 1.053 | 1.000 |
| | | SB_TH:LB_TH | -0.432 | -1.262 | 0.398 | 0.840 |
| | | PL_TH:PI_TH | -0.092 | -0.975 | 0.790 | 1.000 |

| | | | | | | |
|---|---|---|---|---|---|---|
| | | SB_TH:PI_TH | -0.700 | -1.535 | 0.135 | 0.196 |
| | | SB_TH:PL_TH | -0.608 | -1.385 | 0.170 | 0.288 |
| | Úlfljótsvatn | PI_UL:LB_UL | 0.688 | -1.014 | 2.391 | 0.966 |
| | | PL_UL:LB_UL | -0.857 | -2.177 | 0.463 | 0.575 |
| | | PL_UL:PI_UL | -1.545 | -3.173 | 0.082 | 0.080 |
| | Across Þingvallavatn and Úlfljótsvatn | LB_UL:LB_TH | -4.000 | -5.193 | -2.807 | *<<0.001 |
| | | PI_UL:LB_TH | -3.312 | -4.838 | -1.786 | *<<0.001 |
| | | PL_UL:LB_TH | -4.857 | -5.941 | -3.774 | *<<0.001 |
| | | PI_TH:LB_UL | 4.268 | 3.071 | 5.464 | *<<0.001 |
| | | PL_TH:LB_UL | 4.176 | 3.019 | 5.333 | *<<0.001 |
| | | SB_TH:LB_UL | 3.568 | 2.446 | 4.689 | *<<0.001 |
| | | PI_UL:PI_TH | -3.580 | -5.108 | -2.051 | *<<0.001 |
| | | PL_UL:PI_TH | -5.125 | -6.213 | -4.037 | *<<0.001 |
| | | PL_TH:PI_UL | 3.487 | 1.989 | 4.985 | *<<0.001 |
| | | SB_TH:PI_UL | 2.880 | 1.409 | 4.350 | *<<0.001 |
| | | PL_UL:PL_TH | -5.033 | -6.077 | -3.989 | *<<0.001 |
| | | SB_TH:PL_UL | 4.425 | 3.421 | 5.429 | *<<0.001 |

Diff = Estimated differnces between groups; LCL = lower confidence level; UCL = upper confidence level; P = P – value, <0.05 in bold, with asterisk

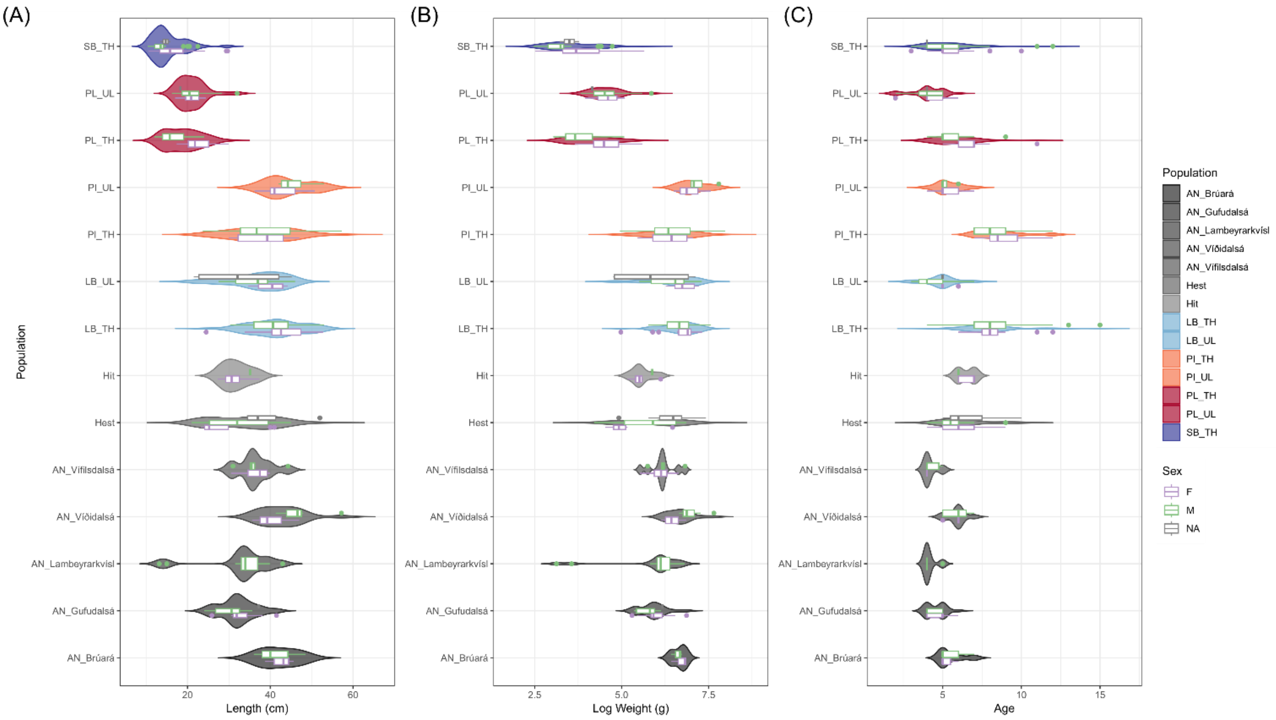


***Supplementary material 6****: Violin plots showing variation in A) length (cm), B) log weight (g) and C) age (years) of all fish used in this study. Shows variation by morph by sex, NA indicates fish that could not be sexed.  Hítarvatn individuals are included in this figure for visualization purposes*

***Supplementary material*** *7: Influence of body size (fork length, FL), population and sex on bone size variation in six jaw bones. Analyzed with Procrustes ANCOVA. Hítarvatn fish excluded from this analysis. AN-charr pooled as one population.*

| Bone Size | | Dentary | Articular-angular | Quadrate | Premaxilla | Maxilla | Supramaxilla |
|---|---|---|---|---|---|---|---|
| **$Log_e$ FL Effect** | **$R^2$** | 94.53% | 96.19% | 95.78% | 93.16% | 94.81% | 94.69% |
| | **F** | 15681.53 | 15880.27 | 15254.41 | 13183.96 | 14037.98 | 11188.12 |
| | **Z** | 18.65 | 18.08 | 18.60 | 17.83 | 16.85 | 16.38 |
| | **P** | ***0.001** | ***0.001** | ***0.001** | ***0.001** | ***0.001** | ***0.001** |
| **Morph Effect** | **$R^2$** | 1.67% | 0.66% | 1.20% | 2.34% | 1.21% | 1.36% |
| | **F** | 34.63 | 13.57 | 23.97 | 41.42 | 22.43 | 20.13 |
| | **Z** | 12.60 | 7.13 | 11.25 | 13.96 | 8.79 | 8.73 |
| | **P** | ***0.001** | ***0.001** | ***0.001** | ***0.001** | ***0.001** | ***0.001** |
| **Morph x $Log_e$ FL Interaction Effect** | **$R^2$** | 0.51% | 0.44% | 0.33% | 0.41% | 0.49% | 0.34% |
| | **F** | 10.60 | 9.13 | 6.52 | 7.21 | 9.01 | 5.03 |
| | **Z** | 7.05 | 6.37 | 4.93 | 5.41 | 6.07 | 4.31 |
| | **P** | ***0.001** | ***0.001** | ***0.001** | ***0.001** | ***0.001** | ***0.001** |
| **Sex Effects** | **$R^2$** | 0.95% | 0.51% | 0.43% | 1.19% | 1.01% | 0.50% |
| | **F** | 156.98 | 84.03 | 68.32 | 168.31 | 148.84 | 58.72 |
| | **Z** | 6.25 | 5.42 | 5.10 | 6.65 | 6.31 | 4.69 |
| | **P** | ***0.001** | ***0.001** | ***0.001** | ***0.001** | ***0.001** | ***0.001** |
| **Sex x $Log_e$ FL Interaction Effect** | **$R^2$** | 0.09% | 0.03% | 0.02% | 0.12% | 0.04% | 0.03% |
| | **F** | 14.23 | 4.60 | 3.60 | 16.35 | 5.64 | 3.34 |
| | **Z** | 3.05 | 1.71 | 1.58 | 3.02 | 1.90 | 1.45 |
| | **P** | ***0.001** | 0.040 | 0.055 | ***0.001** | 0.023 | 0.066 |
| **Morph x Sex Interaction Effects** | **$R^2$** | 0.09% | 0.02% | 0.04% | 0.18% | 0.08% | 0.09% |
| | **F** | 1.86 | 0.49 | 0.80 | 3.23 | 1.41 | 1.35 |
| | **Z** | 1.49 | -1.11 | -0.19 | 2.84 | 0.86 | 0.85 |
| | **P** | 0.063 | 0.860 | 0.558 | ***0.003** | 0.187 | 0.196 |
| **Morph x Sex x $Log_e$ FL Interaction Effect** | **$R^2$** | 0.08% | 0.05% | 0.05% | 0.14% | 0.05% | 0.12% |
| | **F** | 1.64 | 1.01 | 0.98 | 2.49 | 0.85 | 1.81 |
| | **Z** | 1.21 | 0.15 | 0.14 | 2.18 | -0.12 | 1.42 |
| | **P** | 0.109 | 0.431 | 0.448 | 0.014 | 0.546 | 0.075 |

R2 = coefficient of determination; F = F-statistic; Z = Z-statistic; P = P-value, ≤ 0.008 in bold with asterisk.

***Supplementary material 8****: Influence of bone size and population and sex on bone shape variation in six jaw bones. Analyzed with Procrustes ANCOVA. Hítarvatn fish excluded from this analysis. AN-charr pooled as one population.*

| Bone Shape | | Dentary | Articular-angular | Quadrate | Premaxilla | Maxilla | Supramaxilla |
|---|---|---|---|---|---|---|---|
| **$Log_e$ CS Effect** | **$R^2$** | 31.99% | 24.96% | 26.44% | 19.75% | 19.16% | 10.00% |
| | **F** | 276.80 | 179.73 | 154.61 | 120.50 | 117.64 | 47.98 |
| | **Z** | 5.35 | 7.38 | 7.21 | 7.93 | 8.29 | 8.44 |
| | **P** | ***0.001** | ***0.001** | ***0.001** | ***0.001** | ***0.001** | ***0.001** |
| **Morph Effect** | **$R^2$** | 21.91% | 20.62% | 8.84% | 15.78% | 17.61% | 10.60% |
| | **F** | 23.70 | 18.56 | 6.46 | 12.03 | 13.51 | 6.36 |
| | **Z** | 15.49 | 12.48 | 9.71 | 9.68 | 12.25 | 7.29 |
| | **P** | ***0.001** | ***0.001** | ***0.001** | ***0.001** | ***0.001** | ***0.001** |
| **Morph x $Log_e$ CS Interaction Effect** | **$R^2$** | 2.29% | 2.23% | 2.21% | 3.31% | 2.39% | 3.23% |
| | **F** | 2.48 | 2.00 | 1.61 | 2.53 | 1.84 | 1.94 |
| | **Z** | 4.85 | 3.97 | 2.65 | 5.87 | 3.20 | 2.86 |

| | | | | | | | |
|---|---|---|---|---|---|---|---|
| | **P** | ***0.001** | ***0.001** | ***0.002** | ***0.001** | ***0.001** | ***0.005** |
| **Sex Effects** | **$R^2$** | 1.03% | 0.81% | 0.72% | 1.38% | 1.49% | 0.71% |
| | **F** | 8.92 | 5.80 | 4.23 | 8.45 | 9.17 | 3.40 |
| | **Z** | 4.70 | 4.05 | 3.38 | 7.32 | 4.06 | 2.25 |
| | **P** | ***0.001** | ***0.001** | ***0.002** | ***0.001** | ***0.001** | 0.008 |
| **Sex x $Log_e$ CS Interaction Effect** | **$R^2$** | 0.56% | 0.55% | 0.15% | 0.22% | 0.22% | 0.13% |
| | **F** | 4.87 | 3.94 | 0.85 | 1.33 | 1.33 | 0.64 |
| | **Z** | 4.41 | 3.27 | -0.12 | 0.88 | 0.77 | -0.37 |
| | **P** | ***0.001** | ***0.001** | 0.541 | 0.196 | 0.239 | 0.647 |
| **Morph x Sex Interaction Effects** | **$R^2$** | 1.13% | 1.54% | 1.68% | 1.39% | 1.41% | 1.92% |
| | **F** | 1.22 | 1.39 | 1.23 | 1.06 | 1.08 | 1.15 |
| | **Z** | 1.27 | 1.90 | 1.15 | 0.42 | 0.52 | 0.61 |
| | **P** | 0.109 | 0.033 | 0.113 | 0.362 | 0.304 | 0.280 |
| **Morph x Sex x $Log_e$ CS Interaction Effect** | **$R^2$** | 1.11% | 1.25% | 1.49% | 1.14% | 1.54% | 2.74% |
| | **F** | 1.20 | 1.12 | 1.09 | 0.87 | 1.18 | 1.64 |
| | **Z** | 1.11 | 0.75 | 0.49 | -0.82 | 0.84 | 2.03 |
| | **P** | 0.133 | 0.221 | 0.313 | 0.783 | 0.201 | 0.019 |

R2 = coefficient of determination; F = F-statistic; Z = Z-statistic; P = P-value, ≤ 0.008 in bold with asterisk.

***Supplementary material 9****: Results from pairwise tests of bone size by populations, for differences between absolute vector lengths (D) used to examine possible bone size differences by population in five bones. These bones had significant size differences according to ANOVA model with main effects and interaction of populations by body size. Used to verify results from ANOVA and test for true allometry (i.e., different group slopes). Hítarvatn fish excluded from this analysis. AN-charr treated as five separate populations.*

| Bone | System | Pairs | D-value | UCL | Z | P-value |
|---|---|---|---|---|---|---|
| Dentary | AN populations | AN_Brúará:AN_Gufudalsá | 0.569 | 0.738 | 1.11 | 0.143 |
| | | AN_Brúará:AN_Lambeyrarkvísl | 0.767 | 0.931 | 1.30 | 0.091 |
| | | AN_Brúará:AN_Víðidalsá | 1.065 | 0.774 | 2.18 | 0.012 |
| | | AN_Brúará:AN_Vífilsdalsá | 0.612 | 0.935 | 0.95 | 0.175 |
| | | AN_Gufudalsá:AN_Lambeyrarkvísl | 0.198 | 0.630 | -0.08 | 0.548 |
| | | AN_Gufudalsá:AN_Víðidalsá | 0.497 | 0.469 | 1.72 | 0.039 |
| | | AN_Gufudalsá:AN_Vífilsdalsá | 0.043 | 0.611 | -1.32 | 0.886 |
| | | AN_Lambeyrarkvísl:AN_Víðidalsá | 0.298 | 0.651 | 0.42 | 0.352 |
| | | AN_Lambeyrarkvísl:AN_Vífilsdalsá | 0.155 | 0.780 | -0.51 | 0.689 |
| | | AN_Víðidalsá:AN_Vífilsdalsá | 0.453 | 0.639 | 1.03 | 0.152 |
| | Brúará to lake morphs | AN_Brúará:Hest | 0.340 | 0.714 | 0.45 | 0.343 |
| | | AN_Brúará:LB_TH | 0.580 | 0.725 | 1.25 | 0.112 |
| | | AN_Brúará:LB_UL | 0.598 | 0.720 | 1.28 | 0.104 |
| | | AN_Brúará:PI_TH | 0.575 | 0.703 | 1.24 | 0.113 |
| | | AN_Brúará:PI_UL | 0.903 | 0.808 | 1.81 | 0.035 |
| | | AN_Brúará:PL_TH | 0.271 | 0.729 | 0.19 | 0.447 |
| | | AN_Brúará:PL_UL | 0.456 | 0.755 | 0.81 | 0.225 |
| | | AN_Brúará:SB_TH | 0.314 | 0.723 | 0.38 | 0.370 |
| | Gufudalsá to lake morphs | AN_Gufudalsá:Hest | 0.229 | 0.328 | 0.99 | 0.175 |
| | | AN_Gufudalsá:LB_TH | 0.012 | 0.346 | -1.77 | 0.965 |
| | | AN_Gufudalsá:LB_UL | 0.029 | 0.344 | -1.29 | 0.889 |
| | | AN_Gufudalsá:PI_TH | 0.006 | 0.339 | -1.88 | 0.968 |
| | | AN_Gufudalsá:PI_UL | 0.334 | 0.559 | 0.77 | 0.243 |
| | | AN_Gufudalsá:PL_TH | 0.297 | 0.328 | 1.43 | 0.072 |
| | | AN_Gufudalsá:PL_UL | 0.113 | 0.370 | -0.10 | 0.548 |
| | | AN_Gufudalsá:SB_TH | 0.255 | 0.313 | 1.23 | 0.111 |
| | Lambeyrarkvísl to lake morphs | AN_Lambeyrarkvísl:Hest | 0.427 | 0.559 | 1.10 | 0.139 |
| | | AN_Lambeyrarkvísl:LB_TH | 0.187 | 0.593 | -0.05 | 0.538 |

| | | | | | | |
|---|---|---|---|---|---|---|
| | | AN_Lambeyrarkvísl:LB_UL | 0.169 | 0.589 | -0.14 | 0.571 |
| | | AN_Lambeyrarkvísl:PI_TH | 0.192 | 0.574 | 0.01 | 0.514 |
| | | AN_Lambeyrarkvísl:PI_UL | 0.136 | 0.709 | -0.59 | 0.719 |
| | | AN_Lambeyrarkvísl:PL_TH | 0.496 | 0.555 | 1.37 | 0.086 |
| | | AN_Lambeyrarkvísl:PL_UL | 0.311 | 0.627 | 0.57 | 0.298 |
| | | AN_Lambeyrarkvísl:SB_TH | 0.453 | 0.549 | 1.22 | 0.109 |
| | Víðidalsá to lake morphs | AN_Víðidalsá:Hest | 0.725 | 0.374 | 3.01 | ***0.001** |
| | | AN_Víðidalsá:LB_TH | 0.485 | 0.403 | 1.98 | 0.017 |
| | | AN_Víðidalsá:LB_UL | 0.468 | 0.391 | 1.88 | 0.018 |
| | | AN_Víðidalsá:PI_TH | 0.490 | 0.382 | 2.11 | 0.012 |
| | | AN_Víðidalsá:PI_UL | 0.163 | 0.594 | -0.22 | 0.585 |
| | | AN_Víðidalsá:PL_TH | 0.794 | 0.366 | 3.14 | ***0.001** |
| | | AN_Víðidalsá:PL_UL | 0.609 | 0.404 | 2.37 | ***0.002** |
| | | AN_Víðidalsá:SB_TH | 0.751 | 0.376 | 3.06 | ***0.001** |
| | Vífisdalsá to lake morphs | AN_Vífilsdalsá:Hest | 0.272 | 0.534 | 0.51 | 0.326 |
| | | AN_Vífilsdalsá:LB_TH | 0.032 | 0.534 | -1.48 | 0.925 |
| | | AN_Vífilsdalsá:LB_UL | 0.015 | 0.553 | -1.83 | 0.957 |
| | | AN_Vífilsdalsá:PI_TH | 0.037 | 0.550 | -1.39 | 0.901 |
| | | AN_Vífilsdalsá:PI_UL | 0.291 | 0.702 | 0.24 | 0.420 |
| | | AN_Vífilsdalsá:PL_TH | 0.341 | 0.531 | 0.85 | 0.217 |
| | | AN_Vífilsdalsá:PL_UL | 0.156 | 0.557 | -0.20 | 0.586 |
| | | AN_Vífilsdalsá:SB_TH | 0.298 | 0.536 | 0.67 | 0.266 |
| | Hestvatn | Hest:LB_TH | 0.240 | 0.205 | 1.90 | 0.021 |
| | | Hest:LB_UL | 0.258 | 0.215 | 1.93 | 0.022 |
| | | Hest:PI_TH | 0.235 | 0.166 | 2.27 | 0.009 |
| | | Hest:PI_UL | 0.563 | 0.485 | 1.89 | 0.018 |
| | | Hest:PL_TH | 0.069 | 0.158 | 0.35 | 0.374 |
| | | Hest:PL_UL | 0.116 | 0.249 | 0.45 | 0.364 |
| | | Hest:SB_TH | 0.026 | 0.142 | -0.65 | 0.722 |
| | Þingvallavatn | LB_TH:PI_TH | 0.005 | 0.200 | -1.82 | 0.964 |
| | | LB_TH:PL_TH | 0.309 | 0.182 | 2.56 | ***0.005** |
| | | LB_TH:SB_TH | 0.266 | 0.181 | 2.34 | ***0.003** |
| | | PI_TH:PL_TH | 0.304 | 0.154 | 3.03 | ***0.001** |
| | | PI_TH:SB_TH | 0.261 | 0.144 | 2.78 | ***0.002** |
| | | PL_TH:SB_TH | 0.043 | 0.127 | 0.02 | 0.509 |
| | Úlfljótsvatn | LB_UL:PI_UL | 0.305 | 0.488 | 0.83 | 0.217 |
| | | LB_UL:PL_UL | 0.141 | 0.263 | 0.57 | 0.310 |
| | | PI_UL:PL_UL | 0.447 | 0.522 | 1.31 | 0.089 |
| | Across Þingvallavatn and Úlfljótsvatn | LB_TH:LB_UL | 0.017 | 0.240 | -1.29 | 0.886 |
| | | LB_TH:PI_UL | 0.322 | 0.474 | 0.92 | 0.187 |
| | | LB_TH:PL_UL | 0.124 | 0.271 | 0.43 | 0.367 |
| | | LB_UL:PI_TH | 0.023 | 0.223 | -1.04 | 0.838 |
| | | LB_UL:PL_TH | 0.326 | 0.203 | 2.56 | ***0.001** |
| | | LB_UL:SB_TH | 0.283 | 0.200 | 2.30 | ***0.004** |
| | | PI_TH:PI_UL | 0.328 | 0.478 | 0.99 | 0.174 |
| | | PI_TH:PL_UL | 0.119 | 0.245 | 0.43 | 0.357 |
| | | PI_UL:PL_TH | 0.631 | 0.477 | 2.13 | 0.012 |
| | | PI_UL:SB_TH | 0.588 | 0.478 | 1.97 | 0.016 |
| | | PL_TH:PL_UL | 0.185 | 0.243 | 1.18 | 0.127 |
| | | PL_UL:SB_TH | 0.142 | 0.244 | 0.82 | 0.219 |
| Articular-angular | AN populations | AN_Brúará:AN_Gufudalsá | 0.428 | 0.651 | 0.93 | 0.172 |
| | | AN_Brúará:AN_Lambeyrarkvísl | 0.377 | 0.754 | 0.50 | 0.329 |
| | | AN_Brúará:AN_Víðidalsá | 0.753 | 0.665 | 1.81 | 0.032 |
| | | AN_Brúará:AN_Vífilsdalsá | 0.410 | 0.739 | 0.66 | 0.282 |
| | | AN_Gufudalsá:AN_Lambeyrarkvísl | 0.051 | 0.523 | -1.10 | 0.832 |
| | | AN_Gufudalsá:AN_Víðidalsá | 0.325 | 0.395 | 1.22 | 0.119 |
| | | AN_Gufudalsá:AN_Vífilsdalsá | 0.018 | 0.501 | -1.69 | 0.946 |
| | | AN_Lambeyrarkvísl:AN_Víðidalsá | 0.376 | 0.534 | 0.94 | 0.188 |
| | | AN_Lambeyrarkvísl:AN_Vífilsdalsá | 0.033 | 0.642 | -1.51 | 0.927 |
| | | AN_Víðidalsá:AN_Vífilsdalsá | 0.343 | 0.550 | 0.85 | 0.207 |
| | Brúará to lake morphs | AN_Brúará:Hest | 0.250 | 0.590 | 0.29 | 0.397 |

| | | | | | | |
|---|---|---|---|---|---|---|
| | | AN_Brúará:LB_TH | 0.462 | 0.611 | 1.15 | 0.130 |
| | | AN_Brúará:LB_UL | 0.388 | 0.608 | 0.87 | 0.202 |
| | | AN_Brúará:PI_TH | 0.388 | 0.604 | 0.90 | 0.205 |
| | | AN_Brúará:PI_UL | 0.684 | 0.687 | 1.56 | 0.052 |
| | | AN_Brúará:PL_TH | 0.106 | 0.611 | -0.60 | 0.728 |
| | | AN_Brúará:PL_UL | 0.264 | 0.633 | 0.30 | 0.394 |
| | | AN_Brúará:SB_TH | 0.201 | 0.607 | 0.04 | 0.500 |
| | Gufudalsá to lake morphs | AN_Gufudalsá:Hest | 0.178 | 0.285 | 0.85 | 0.206 |
| | | AN_Gufudalsá:LB_TH | 0.034 | 0.298 | -0.80 | 0.772 |
| | | AN_Gufudalsá:LB_UL | 0.040 | 0.289 | -0.86 | 0.790 |
| | | AN_Gufudalsá:PI_TH | 0.040 | 0.278 | -0.75 | 0.775 |
| | | AN_Gufudalsá:PI_UL | 0.256 | 0.483 | 0.59 | 0.297 |
| | | AN_Gufudalsá:PL_TH | 0.323 | 0.274 | 1.91 | 0.025 |
| | | AN_Gufudalsá:PL_UL | 0.164 | 0.314 | 0.55 | 0.301 |
| | | AN_Gufudalsá:SB_TH | 0.227 | 0.260 | 1.32 | 0.090 |
| | Lambeyrarkvísl to lake morphs | AN_Lambeyrarkvísl:Hest | 0.127 | 0.472 | -0.30 | 0.622 |
| | | AN_Lambeyrarkvísl:LB_TH | 0.085 | 0.484 | -0.64 | 0.716 |
| | | AN_Lambeyrarkvísl:LB_UL | 0.010 | 0.498 | -1.89 | 0.972 |
| | | AN_Lambeyrarkvísl:PI_TH | 0.011 | 0.474 | -1.80 | 0.962 |
| | | AN_Lambeyrarkvísl:PI_UL | 0.307 | 0.606 | 0.53 | 0.328 |
| | | AN_Lambeyrarkvísl:PL_TH | 0.272 | 0.473 | 0.67 | 0.268 |
| | | AN_Lambeyrarkvísl:PL_UL | 0.113 | 0.506 | -0.40 | 0.649 |
| | | AN_Lambeyrarkvísl:SB_TH | 0.177 | 0.462 | 0.10 | 0.467 |
| | Víðidalsá to lake morphs | AN_Víðidalsá:Hest | 0.503 | 0.311 | 2.49 | *0.002 |
| | | AN_Víðidalsá:LB_TH | 0.291 | 0.333 | 1.35 | 0.101 |
| | | AN_Víðidalsá:LB_UL | 0.366 | 0.340 | 1.71 | 0.037 |
| | | AN_Víðidalsá:PI_TH | 0.365 | 0.317 | 1.82 | 0.029 |
| | | AN_Víðidalsá:PI_UL | 0.069 | 0.496 | -0.88 | 0.797 |
| | | AN_Víðidalsá:PL_TH | 0.648 | 0.309 | 3.03 | *0.001 |
| | | AN_Víðidalsá:PL_UL | 0.489 | 0.351 | 2.23 | *0.006 |
| | | AN_Víðidalsá:SB_TH | 0.552 | 0.317 | 2.64 | *0.001 |
| | Vífisdalsá to lake morphs | AN_Vífilsdalsá:Hest | 0.160 | 0.451 | 0.04 | 0.508 |
| | | AN_Vífilsdalsá:LB_TH | 0.052 | 0.457 | -1.04 | 0.837 |
| | | AN_Vífilsdalsá:LB_UL | 0.023 | 0.479 | -1.45 | 0.911 |
| | | AN_Vífilsdalsá:PI_TH | 0.022 | 0.458 | -1.56 | 0.933 |
| | | AN_Vífilsdalsá:PI_UL | 0.274 | 0.595 | 0.39 | 0.382 |
| | | AN_Vífilsdalsá:PL_TH | 0.305 | 0.469 | 0.91 | 0.188 |
| | | AN_Vífilsdalsá:PL_UL | 0.146 | 0.479 | -0.11 | 0.565 |
| | | AN_Vífilsdalsá:SB_TH | 0.210 | 0.451 | 0.39 | 0.370 |
| | Hestvatn | Hest:LB_TH | 0.212 | 0.177 | 1.95 | 0.019 |
| | | Hest:LB_UL | 0.138 | 0.176 | 1.10 | 0.146 |
| | | Hest:PI_TH | 0.138 | 0.140 | 1.52 | 0.060 |
| | | Hest:PI_UL | 0.434 | 0.400 | 1.70 | 0.040 |
| | | Hest:PL_TH | 0.145 | 0.139 | 1.67 | 0.043 |
| | | Hest:PL_UL | 0.014 | 0.208 | -1.27 | 0.885 |
| | | Hest:SB_TH | 0.049 | 0.120 | 0.23 | 0.416 |
| | Þingvallavatn | LB_TH:PI_TH | 0.074 | 0.171 | 0.35 | 0.384 |
| | | LB_TH:PL_TH | 0.357 | 0.168 | 3.08 | *0.001 |
| | | LB_TH:SB_TH | 0.261 | 0.158 | 2.56 | *0.001 |
| | | PI_TH:PL_TH | 0.283 | 0.136 | 3.16 | *0.001 |
| | | PI_TH:SB_TH | 0.187 | 0.118 | 2.48 | *0.004 |
| | | PL_TH:SB_TH | 0.095 | 0.111 | 1.36 | 0.084 |
| | Úlfljótsvatn | LB_UL:PI_UL | 0.297 | 0.430 | 0.98 | 0.185 |
| | | LB_UL:PL_UL | 0.124 | 0.232 | 0.54 | 0.317 |
| | | PI_UL:PL_UL | 0.420 | 0.452 | 1.45 | 0.074 |
| | Across Þingvallavatn and Úlfljótsvatn | LB_TH:LB_UL | 0.075 | 0.207 | 0.14 | 0.460 |
| | | LB_TH:PI_UL | 0.222 | 0.420 | 0.60 | 0.300 |
| | | LB_TH:PL_UL | 0.198 | 0.227 | 1.34 | 0.093 |
| | | LB_UL:PI_TH | 0.001 | 0.183 | -2.45 | 0.998 |
| | | LB_UL:PL_TH | 0.282 | 0.179 | 2.53 | *0.002 |
| | | LB_UL:SB_TH | 0.187 | 0.170 | 1.79 | 0.034 |

| | | | | | | |
|---|---|---|---|---|---|---|
| | | PI_TH:PI_UL | 0.296 | 0.400 | 1.03 | 0.169 |
| | | PI_TH:PL_UL | 0.124 | 0.211 | 0.74 | 0.243 |
| | | PI_UL:PL_TH | 0.579 | 0.406 | 2.23 | ***0.007** |
| | | PI_UL:SB_TH | 0.484 | 0.414 | 1.90 | 0.020 |
| | | PL_TH:PL_UL | 0.158 | 0.197 | 1.19 | 0.113 |
| | | PL_UL:SB_TH | 0.063 | 0.195 | -0.03 | 0.530 |
| Quadrate | AN populations | AN_Brúará:AN_Gufudalsá | 0.767 | 0.812 | 1.48 | 0.066 |
| | | AN_Brúará:AN_Lambeyrarkvísl | 0.962 | 0.863 | 1.72 | 0.040 |
| | | AN_Brúará:AN_Víðidalsá | 0.878 | 0.845 | 1.70 | 0.042 |
| | | AN_Brúará:AN_Vífilsdalsá | 1.172 | 0.882 | 2.08 | 0.011 |
| | | AN_Gufudalsá:AN_Lambeyrarkvísl | 0.195 | 0.537 | 0.18 | 0.449 |
| | | AN_Gufudalsá:AN_Víðidalsá | 0.111 | 0.383 | -0.16 | 0.577 |
| | | AN_Gufudalsá:AN_Vífilsdalsá | 0.405 | 0.494 | 1.25 | 0.102 |
| | | AN_Lambeyrarkvísl:AN_Víðidalsá | 0.084 | 0.534 | -0.70 | 0.758 |
| | | AN_Lambeyrarkvísl:AN_Vífilsdalsá | 0.210 | 0.621 | -0.01 | 0.532 |
| | | AN_Víðidalsá:AN_Vífilsdalsá | 0.294 | 0.513 | 0.72 | 0.256 |
| | Brúará to lake morphs | AN_Brúará:Hest | 0.654 | 0.795 | 1.27 | 0.113 |
| | | AN_Brúará:LB_TH | 0.726 | 0.797 | 1.42 | 0.081 |
| | | AN_Brúará:LB_UL | 0.829 | 0.837 | 1.62 | 0.054 |
| | | AN_Brúará:PI_TH | 0.859 | 0.804 | 1.74 | 0.035 |
| | | AN_Brúará:PI_UL | 0.755 | 0.847 | 1.36 | 0.086 |
| | | AN_Brúará:PL_TH | 0.530 | 0.789 | 0.95 | 0.175 |
| | | AN_Brúará:PL_UL | 0.681 | 0.815 | 1.30 | 0.099 |
| | | AN_Brúará:SB_TH | 0.638 | 0.803 | 1.25 | 0.114 |
| | Gufudalsá to lake morphs | AN_Gufudalsá:Hest | 0.112 | 0.262 | 0.25 | 0.421 |
| | | AN_Gufudalsá:LB_TH | 0.041 | 0.273 | -0.76 | 0.756 |
| | | AN_Gufudalsá:LB_UL | 0.062 | 0.280 | -0.48 | 0.679 |
| | | AN_Gufudalsá:PI_TH | 0.092 | 0.266 | -0.01 | 0.522 |
| | | AN_Gufudalsá:PI_UL | 0.011 | 0.443 | -1.84 | 0.957 |
| | | AN_Gufudalsá:PL_TH | 0.236 | 0.263 | 1.40 | 0.078 |
| | | AN_Gufudalsá:PL_UL | 0.086 | 0.298 | -0.16 | 0.574 |
| | | AN_Gufudalsá:SB_TH | 0.128 | 0.250 | 0.50 | 0.327 |
| | Lambeyrarkvísl to lake morphs | AN_Lambeyrarkvísl:Hest | 0.308 | 0.450 | 0.94 | 0.176 |
| | | AN_Lambeyrarkvísl:LB_TH | 0.236 | 0.480 | 0.53 | 0.329 |
| | | AN_Lambeyrarkvísl:LB_UL | 0.133 | 0.463 | -0.15 | 0.566 |
| | | AN_Lambeyrarkvísl:PI_TH | 0.103 | 0.469 | -0.36 | 0.648 |
| | | AN_Lambeyrarkvísl:PI_UL | 0.206 | 0.608 | 0.13 | 0.458 |
| | | AN_Lambeyrarkvísl:PL_TH | 0.432 | 0.451 | 1.48 | 0.064 |
| | | AN_Lambeyrarkvísl:PL_UL | 0.281 | 0.476 | 0.75 | 0.247 |
| | | AN_Lambeyrarkvísl:SB_TH | 0.323 | 0.451 | 1.02 | 0.146 |
| | Víðidalsá to lake morphs | AN_Víðidalsá:Hest | 0.224 | 0.319 | 1.02 | 0.172 |
| | | AN_Víðidalsá:LB_TH | 0.152 | 0.328 | 0.42 | 0.362 |
| | | AN_Víðidalsá:LB_UL | 0.049 | 0.332 | -0.78 | 0.778 |
| | | AN_Víðidalsá:PI_TH | 0.019 | 0.311 | -1.44 | 0.918 |
| | | AN_Víðidalsá:PI_UL | 0.122 | 0.486 | -0.26 | 0.614 |
| | | AN_Víðidalsá:PL_TH | 0.348 | 0.321 | 1.81 | 0.029 |
| | | AN_Víðidalsá:PL_UL | 0.197 | 0.346 | 0.72 | 0.243 |
| | | AN_Víðidalsá:SB_TH | 0.239 | 0.305 | 1.21 | 0.126 |
| | Vífisdalsá to lake morphs | AN_Vífilsdalsá:Hest | 0.518 | 0.442 | 1.83 | 0.030 |
| | | AN_Vífilsdalsá:LB_TH | 0.446 | 0.460 | 1.55 | 0.056 |
| | | AN_Vífilsdalsá:LB_UL | 0.344 | 0.479 | 1.08 | 0.155 |
| | | AN_Vífilsdalsá:PI_TH | 0.313 | 0.449 | 0.98 | 0.173 |
| | | AN_Vífilsdalsá:PI_UL | 0.417 | 0.601 | 1.03 | 0.163 |
| | | AN_Vífilsdalsá:PL_TH | 0.642 | 0.460 | 2.25 | ***0.007** |
| | | AN_Vífilsdalsá:PL_UL | 0.491 | 0.487 | 1.64 | 0.047 |
| | | AN_Vífilsdalsá:SB_TH | 0.534 | 0.454 | 1.91 | 0.024 |
| | Hestvatn | Hest:LB_TH | 0.072 | 0.170 | 0.35 | 0.375 |
| | | Hest:LB_UL | 0.174 | 0.180 | 1.54 | 0.060 |
| | | Hest:PI_TH | 0.204 | 0.149 | 2.24 | 0.008 |
| | | Hest:PI_UL | 0.101 | 0.396 | -0.26 | 0.605 |
| | | Hest:PL_TH | 0.124 | 0.130 | 1.49 | 0.067 |

| | | | | | | |
|---|---|---|---|---|---|---|
| | | Hest:PL_UL | 0.026 | 0.201 | -0.85 | 0.791 |
| | | Hest:SB_TH | 0.016 | 0.123 | -0.95 | 0.817 |
| | Þingvallavatn | LB_TH:PI_TH | 0.133 | 0.165 | 1.22 | 0.110 |
| | | LB_TH:PL_TH | 0.196 | 0.149 | 2.15 | ***0.007** |
| | | LB_TH:SB_TH | 0.088 | 0.144 | 0.79 | 0.234 |
| | | PI_TH:PL_TH | 0.328 | 0.125 | 3.75 | ***0.001** |
| | | PI_TH:SB_TH | 0.220 | 0.121 | 2.86 | ***0.001** |
| | | PL_TH:SB_TH | 0.108 | 0.108 | 1.63 | 0.050 |
| | Úlfljótsvatn | LB_UL:PI_UL | 0.073 | 0.409 | -0.61 | 0.739 |
| | | LB_UL:PL_UL | 0.148 | 0.229 | 0.89 | 0.206 |
| | | PI_UL:PL_UL | 0.075 | 0.424 | -0.61 | 0.726 |
| | Across Þingvallavatn and Úlfljótsvatn | LB_TH:LB_UL | 0.103 | 0.198 | 0.54 | 0.321 |
| | | LB_TH:PI_UL | 0.029 | 0.408 | -1.19 | 0.861 |
| | | LB_TH:PL_UL | 0.045 | 0.209 | -0.48 | 0.685 |
| | | LB_UL:PI_TH | 0.030 | 0.184 | -0.71 | 0.749 |
| | | LB_UL:PL_TH | 0.298 | 0.178 | 2.67 | ***0.001** |
| | | LB_UL:SB_TH | 0.190 | 0.160 | 1.86 | 0.020 |
| | | PI_TH:PI_UL | 0.103 | 0.398 | -0.22 | 0.595 |
| | | PI_TH:PL_UL | 0.178 | 0.201 | 1.38 | 0.081 |
| | | PI_UL:PL_TH | 0.225 | 0.395 | 0.70 | 0.271 |
| | | PI_UL:SB_TH | 0.117 | 0.386 | -0.07 | 0.543 |
| | | PL_TH:PL_UL | 0.150 | 0.190 | 1.17 | 0.122 |
| | | PL_UL:SB_TH | 0.042 | 0.187 | -0.36 | 0.653 |
| Premaxilla | AN populations | AN_Brúará:AN_Gufudalsá | 0.880 | 0.887 | 1.52 | 0.054 |
| | | AN_Brúará:AN_Lambeyrarkvísl | 1.019 | 1.018 | 1.55 | 0.050 |
| | | AN_Brúará:AN_Víðidalsá | 1.543 | 0.923 | 2.61 | ***0.004** |
| | | AN_Brúará:AN_Vífilsdalsá | 0.980 | 1.034 | 1.50 | 0.064 |
| | | AN_Gufudalsá:AN_Lambeyrarkvísl | 0.139 | 0.745 | -0.57 | 0.698 |
| | | AN_Gufudalsá:AN_Víðidalsá | 0.663 | 0.551 | 1.96 | 0.018 |
| | | AN_Gufudalsá:AN_Vífilsdalsá | 0.100 | 0.684 | -0.72 | 0.755 |
| | | AN_Lambeyrarkvísl:AN_Víðidalsá | 0.524 | 0.757 | 0.96 | 0.174 |
| | | AN_Lambeyrarkvísl:AN_Vífilsdalsá | 0.039 | 0.896 | -1.64 | 0.946 |
| | | AN_Víðidalsá:AN_Vífilsdalsá | 0.563 | 0.705 | 1.16 | 0.127 |
| | Brúará to lake morphs | AN_Brúará:Hest | 0.656 | 0.852 | 1.16 | 0.131 |
| | | AN_Brúará:LB_TH | 0.755 | 0.854 | 1.38 | 0.082 |
| | | AN_Brúará:LB_UL | 0.725 | 0.848 | 1.30 | 0.096 |
| | | AN_Brúará:PI_TH | 0.894 | 0.829 | 1.69 | 0.034 |
| | | AN_Brúará:PI_UL | 1.239 | 1.010 | 1.97 | 0.014 |
| | | AN_Brúará:PL_TH | 0.481 | 0.829 | 0.71 | 0.260 |
| | | AN_Brúará:PL_UL | 0.687 | 0.883 | 1.17 | 0.114 |
| | | AN_Brúará:SB_TH | 0.545 | 0.832 | 0.87 | 0.205 |
| | Gufudalsá to lake morphs | AN_Gufudalsá:Hest | 0.224 | 0.355 | 0.80 | 0.239 |
| | | AN_Gufudalsá:LB_TH | 0.126 | 0.390 | 0.02 | 0.509 |
| | | AN_Gufudalsá:LB_UL | 0.155 | 0.390 | 0.23 | 0.425 |
| | | AN_Gufudalsá:PI_TH | 0.014 | 0.352 | -1.59 | 0.934 |
| | | AN_Gufudalsá:PI_UL | 0.359 | 0.649 | 0.69 | 0.252 |
| | | AN_Gufudalsá:PL_TH | 0.400 | 0.352 | 1.79 | 0.026 |
| | | AN_Gufudalsá:PL_UL | 0.194 | 0.415 | 0.40 | 0.370 |
| | | AN_Gufudalsá:SB_TH | 0.335 | 0.347 | 1.51 | 0.060 |
| | Lambeyrarkvísl to lake morphs | AN_Lambeyrarkvísl:Hest | 0.363 | 0.689 | 0.64 | 0.284 |
| | | AN_Lambeyrarkvísl:LB_TH | 0.265 | 0.700 | 0.24 | 0.429 |
| | | AN_Lambeyrarkvísl:LB_UL | 0.294 | 0.689 | 0.31 | 0.392 |
| | | AN_Lambeyrarkvísl:PI_TH | 0.125 | 0.677 | -0.57 | 0.719 |
| | | AN_Lambeyrarkvísl:PI_UL | 0.220 | 0.836 | -0.29 | 0.616 |
| | | AN_Lambeyrarkvísl:PL_TH | 0.539 | 0.683 | 1.24 | 0.109 |
| | | AN_Lambeyrarkvísl:PL_UL | 0.333 | 0.702 | 0.44 | 0.343 |
| | | AN_Lambeyrarkvísl:SB_TH | 0.474 | 0.683 | 1.05 | 0.157 |
| | Víðidalsá to lake morphs | AN_Víðidalsá:Hest | 0.887 | 0.433 | 3.12 | ***0.001** |
| | | AN_Víðidalsá:LB_TH | 0.789 | 0.444 | 2.74 | ***0.001** |
| | | AN_Víðidalsá:LB_UL | 0.818 | 0.491 | 2.70 | ***0.001** |
| | | AN_Víðidalsá:PI_TH | 0.649 | 0.434 | 2.38 | ***0.006** |

| | | | | | | |
|---|---|---|---|---|---|---|
| | | AN_Víðidalsá:PI_UL | 0.304 | 0.662 | 0.34 | 0.384 |
| | | AN_Víðidalsá:PL_TH | 1.062 | 0.431 | 3.65 | ***0.001** |
| | | AN_Víðidalsá:PL_UL | 0.856 | 0.473 | 2.70 | ***0.001** |
| | | AN_Víðidalsá:SB_TH | 0.998 | 0.427 | 3.46 | ***0.001** |
| | Vífisdalsá to lake morphs | AN_Vífilsdalsá:Hest | 0.324 | 0.610 | 0.61 | 0.291 |
| | | AN_Vífilsdalsá:LB_TH | 0.225 | 0.627 | 0.14 | 0.468 |
| | | AN_Vífilsdalsá:LB_UL | 0.255 | 0.628 | 0.26 | 0.421 |
| | | AN_Vífilsdalsá:PI_TH | 0.086 | 0.601 | -0.81 | 0.775 |
| | | AN_Vífilsdalsá:PI_UL | 0.259 | 0.809 | -0.04 | 0.515 |
| | | AN_Vífilsdalsá:PL_TH | 0.499 | 0.609 | 1.28 | 0.107 |
| | | AN_Vífilsdalsá:PL_UL | 0.293 | 0.624 | 0.39 | 0.386 |
| | | AN_Vífilsdalsá:SB_TH | 0.435 | 0.581 | 1.08 | 0.148 |
| | Hestvatn | Hest:LB_TH | 0.099 | 0.221 | 0.30 | 0.404 |
| | | Hest:LB_UL | 0.069 | 0.247 | -0.19 | 0.576 |
| | | Hest:PI_TH | 0.238 | 0.188 | 1.99 | 0.018 |
| | | Hest:PI_UL | 0.583 | 0.576 | 1.62 | 0.046 |
| | | Hest:PL_TH | 0.175 | 0.184 | 1.52 | 0.060 |
| | | Hest:PL_UL | 0.031 | 0.286 | -1.08 | 0.857 |
| | | Hest:SB_TH | 0.111 | 0.168 | 0.84 | 0.217 |
| | Þingvallavatn | LB_TH:PI_TH | 0.140 | 0.225 | 0.84 | 0.212 |
| | | LB_TH:PL_TH | 0.274 | 0.209 | 2.07 | 0.009 |
| | | LB_TH:SB_TH | 0.209 | 0.197 | 1.70 | 0.039 |
| | | PI_TH:PL_TH | 0.414 | 0.180 | 3.34 | ***0.001** |
| | | PI_TH:SB_TH | 0.349 | 0.170 | 3.11 | ***0.001** |
| | | PL_TH:SB_TH | 0.065 | 0.144 | 0.33 | 0.394 |
| | Úlfljótsvatn | LB_UL:PI_UL | 0.514 | 0.583 | 1.38 | 0.082 |
| | | LB_UL:PL_UL | 0.038 | 0.321 | -0.93 | 0.803 |
| | | PI_UL:PL_UL | 0.552 | 0.602 | 1.44 | 0.071 |
| | Across Þingvallavatn and Úlfljótsvatn | LB_TH:LB_UL | 0.029 | 0.268 | -0.99 | 0.820 |
| | | LB_TH:PI_UL | 0.484 | 0.573 | 1.29 | 0.100 |
| | | LB_TH:PL_UL | 0.068 | 0.302 | -0.45 | 0.673 |
| | | LB_UL:PI_TH | 0.169 | 0.251 | 0.97 | 0.171 |
| | | LB_UL:PL_TH | 0.244 | 0.233 | 1.67 | 0.042 |
| | | LB_UL:SB_TH | 0.180 | 0.217 | 1.20 | 0.120 |
| | | PI_TH:PI_UL | 0.345 | 0.552 | 0.80 | 0.242 |
| | | PI_TH:PL_UL | 0.208 | 0.275 | 1.08 | 0.139 |
| | | PI_UL:PL_TH | 0.758 | 0.579 | 2.12 | 0.012 |
| | | PI_UL:SB_TH | 0.694 | 0.562 | 1.94 | 0.018 |
| | | PL_TH:PL_UL | 0.206 | 0.265 | 1.16 | 0.118 |
| | | PL_UL:SB_TH | 0.141 | 0.259 | 0.60 | 0.299 |
| Maxilla | AN populations | AN_Brúará:AN_Gufudalsá | 0.801 | 0.801 | 1.65 | 0.051 |
| | | AN_Brúará:AN_Lambeyrarkvísl | 0.838 | 0.901 | 1.50 | 0.068 |
| | | AN_Brúará:AN_Víðidalsá | 1.028 | 0.804 | 2.07 | 0.010 |
| | | AN_Brúará:AN_Vífilsdalsá | 0.889 | 0.888 | 1.60 | 0.050 |
| | | AN_Gufudalsá:AN_Lambeyrarkvísl | 0.037 | 0.608 | -1.40 | 0.907 |
| | | AN_Gufudalsá:AN_Víðidalsá | 0.227 | 0.453 | 0.46 | 0.347 |
| | | AN_Gufudalsá:AN_Vífilsdalsá | 0.089 | 0.628 | -0.81 | 0.789 |
| | | AN_Lambeyrarkvísl:AN_Víðidalsá | 0.190 | 0.679 | -0.14 | 0.569 |
| | | AN_Lambeyrarkvísl:AN_Vífilsdalsá | 0.051 | 0.760 | -1.35 | 0.897 |
| | | AN_Víðidalsá:AN_Vífilsdalsá | 0.139 | 0.640 | -0.43 | 0.673 |
| | Brúará to lake morphs | AN_Brúará:Hest | 0.559 | 0.715 | 1.17 | 0.112 |
| | | AN_Brúará:LB_TH | 0.730 | 0.714 | 1.62 | 0.047 |
| | | AN_Brúará:LB_UL | 0.767 | 0.728 | 1.67 | 0.038 |
| | | AN_Brúará:PI_TH | 0.723 | 0.711 | 1.59 | 0.047 |
| | | AN_Brúará:PI_UL | 1.138 | 0.855 | 2.14 | 0.010 |
| | | AN_Brúará:PL_TH | 0.399 | 0.711 | 0.67 | 0.278 |
| | | AN_Brúará:PL_UL | 0.629 | 0.736 | 1.32 | 0.089 |
| | | AN_Brúará:SB_TH | 0.515 | 0.720 | 1.06 | 0.141 |
| | Gufudalsá to lake morphs | AN_Gufudalsá:Hest | 0.241 | 0.314 | 1.12 | 0.136 |
| | | AN_Gufudalsá:LB_TH | 0.070 | 0.338 | -0.49 | 0.684 |
| | | AN_Gufudalsá:LB_UL | 0.033 | 0.352 | -1.09 | 0.847 |

| | | | | | | |
|---|---|---|---|---|---|---|
| | | AN_Gufudalsá:PI_TH | 0.078 | 0.332 | -0.32 | 0.631 |
| | | AN_Gufudalsá:PI_UL | 0.337 | 0.564 | 0.78 | 0.227 |
| | | AN_Gufudalsá:PL_TH | 0.402 | 0.316 | 2.04 | 0.012 |
| | | AN_Gufudalsá:PL_UL | 0.172 | 0.366 | 0.42 | 0.351 |
| | | AN_Gufudalsá:SB_TH | 0.285 | 0.302 | 1.47 | 0.063 |
| | Lambeyrarkvísl to lake morphs | AN_Lambeyrarkvísl:Hest | 0.279 | 0.579 | 0.51 | 0.317 |
| | | AN_Lambeyrarkvísl:LB_TH | 0.108 | 0.568 | -0.55 | 0.711 |
| | | AN_Lambeyrarkvísl:LB_UL | 0.071 | 0.585 | -0.90 | 0.803 |
| | | AN_Lambeyrarkvísl:PI_TH | 0.115 | 0.569 | -0.47 | 0.694 |
| | | AN_Lambeyrarkvísl:PI_UL | 0.300 | 0.725 | 0.33 | 0.392 |
| | | AN_Lambeyrarkvísl:PL_TH | 0.439 | 0.565 | 1.22 | 0.107 |
| | | AN_Lambeyrarkvísl:PL_UL | 0.209 | 0.605 | 0.10 | 0.472 |
| | | AN_Lambeyrarkvísl:SB_TH | 0.323 | 0.558 | 0.74 | 0.237 |
| | Víðidalsá to lake morphs | AN_Víðidalsá:Hest | 0.468 | 0.374 | 2.04 | 0.014 |
| | | AN_Víðidalsá:LB_TH | 0.298 | 0.372 | 1.17 | 0.129 |
| | | AN_Víðidalsá:LB_UL | 0.261 | 0.382 | 0.92 | 0.197 |
| | | AN_Víðidalsá:PI_TH | 0.305 | 0.358 | 1.28 | 0.109 |
| | | AN_Víðidalsá:PI_UL | 0.110 | 0.591 | -0.56 | 0.703 |
| | | AN_Víðidalsá:PL_TH | 0.629 | 0.360 | 2.67 | ***0.002** |
| | | AN_Víðidalsá:PL_UL | 0.399 | 0.405 | 1.59 | 0.058 |
| | | AN_Víðidalsá:SB_TH | 0.512 | 0.364 | 2.29 | ***0.007** |
| | Vífisdalsá to lake morphs | AN_Vífilsdalsá:Hest | 0.330 | 0.550 | 0.74 | 0.247 |
| | | AN_Vífilsdalsá:LB_TH | 0.159 | 0.546 | -0.18 | 0.594 |
| | | AN_Vífilsdalsá:LB_UL | 0.122 | 0.562 | -0.48 | 0.685 |
| | | AN_Vífilsdalsá:PI_TH | 0.167 | 0.552 | -0.13 | 0.560 |
| | | AN_Vífilsdalsá:PI_UL | 0.249 | 0.730 | 0.02 | 0.495 |
| | | AN_Vífilsdalsá:PL_TH | 0.490 | 0.553 | 1.41 | 0.085 |
| | | AN_Vífilsdalsá:PL_UL | 0.261 | 0.584 | 0.36 | 0.373 |
| | | AN_Vífilsdalsá:SB_TH | 0.374 | 0.546 | 0.96 | 0.175 |
| | Hestvatn | Hest:LB_TH | 0.171 | 0.200 | 1.34 | 0.097 |
| | | Hest:LB_UL | 0.208 | 0.221 | 1.52 | 0.069 |
| | | Hest:PI_TH | 0.163 | 0.172 | 1.50 | 0.072 |
| | | Hest:PI_UL | 0.578 | 0.495 | 1.84 | 0.025 |
| | | Hest:PL_TH | 0.160 | 0.157 | 1.56 | 0.045 |
| | | Hest:PL_UL | 0.069 | 0.251 | -0.20 | 0.583 |
| | | Hest:SB_TH | 0.044 | 0.147 | -0.15 | 0.563 |
| | Þingvallavatn | LB_TH:PI_TH | 0.007 | 0.196 | -1.63 | 0.937 |
| | | LB_TH:PL_TH | 0.331 | 0.186 | 2.75 | ***0.001** |
| | | LB_TH:SB_TH | 0.215 | 0.173 | 2.02 | 0.013 |
| | | PI_TH:PL_TH | 0.324 | 0.157 | 3.18 | ***0.001** |
| | | PI_TH:SB_TH | 0.207 | 0.147 | 2.24 | ***0.002** |
| | | PL_TH:SB_TH | 0.116 | 0.135 | 1.36 | 0.087 |
| | Úlfljótsvatn | LB_UL:PI_UL | 0.371 | 0.521 | 1.06 | 0.151 |
| | | LB_UL:PL_UL | 0.139 | 0.274 | 0.55 | 0.305 |
| | | PI_UL:PL_UL | 0.509 | 0.513 | 1.55 | 0.053 |
| | Across Þingvallavatn and Úlfljótsvatn | LB_TH:LB_UL | 0.037 | 0.234 | -0.74 | 0.747 |
| | | LB_TH:PI_UL | 0.408 | 0.483 | 1.27 | 0.104 |
| | | LB_TH:PL_UL | 0.101 | 0.257 | 0.22 | 0.424 |
| | | LB_UL:PI_TH | 0.044 | 0.219 | -0.49 | 0.674 |
| | | LB_UL:PL_TH | 0.368 | 0.207 | 2.69 | ***0.002** |
| | | LB_UL:SB_TH | 0.252 | 0.201 | 2.01 | 0.013 |
| | | PI_TH:PI_UL | 0.415 | 0.494 | 1.30 | 0.096 |
| | | PI_TH:PL_UL | 0.094 | 0.237 | 0.18 | 0.458 |
| | | PI_UL:PL_TH | 0.739 | 0.510 | 2.31 | ***0.004** |
| | | PI_UL:SB_TH | 0.622 | 0.490 | 2.05 | 0.012 |
| | | PL_TH:PL_UL | 0.230 | 0.242 | 1.52 | 0.065 |
| | | PL_UL:SB_TH | 0.113 | 0.232 | 0.54 | 0.323 |
| Supramaxilla | AN populations | AN_Brúará:AN_Gufudalsá | 0.261 | 0.718 | 0.05 | 0.494 |
| | | AN_Brúará:AN_Lambeyrarkvísl | 0.462 | 0.853 | 0.58 | 0.300 |
| | | AN_Brúará:AN_Víðidalsá | 0.680 | 0.735 | 1.43 | 0.074 |
| | | AN_Brúará:AN_Vífilsdalsá | 0.358 | 0.850 | 0.28 | 0.415 |

| | | | | | | |
|---|---|---|---|---|---|---|
| | | AN_Gufudalsá:AN_Lambeyrarkvísl | 0.201 | 0.668 | -0.06 | 0.535 |
| | | AN_Gufudalsá:AN_Víðidalsá | 0.419 | 0.470 | 1.36 | 0.094 |
| | | AN_Gufudalsá:AN_Vífilsdalsá | 0.097 | 0.595 | -0.68 | 0.741 |
| | | AN_Lambeyrarkvísl:AN_Víðidalsá | 0.218 | 0.675 | -0.05 | 0.535 |
| | | AN_Lambeyrarkvísl:AN_Vífilsdalsá | 0.104 | 0.769 | -0.79 | 0.781 |
| | | AN_Víðidalsá:AN_Vífilsdalsá | 0.322 | 0.616 | 0.56 | 0.304 |
| | Brúará to lake morphs | AN_Brúará:Hest | 0.078 | 0.675 | -1.01 | 0.817 |
| | | AN_Brúará:LB_TH | 0.260 | 0.660 | 0.11 | 0.484 |
| | | AN_Brúará:LB_UL | 0.233 | 0.691 | -0.02 | 0.520 |
| | | AN_Brúará:PI_TH | 0.151 | 0.666 | -0.49 | 0.692 |
| | | AN_Brúará:PI_UL | 0.474 | 0.816 | 0.72 | 0.255 |
| | | AN_Brúará:PL_TH | 0.018 | 0.666 | -1.73 | 0.955 |
| | | AN_Brúará:PL_UL | 0.137 | 0.683 | -0.59 | 0.726 |
| | | AN_Brúará:SB_TH | 0.052 | 0.656 | -1.31 | 0.882 |
| | Gufudalsá to lake morphs | AN_Gufudalsá:Hest | 0.183 | 0.321 | 0.68 | 0.267 |
| | | AN_Gufudalsá:LB_TH | 0.001 | 0.340 | -2.42 | 0.997 |
| | | AN_Gufudalsá:LB_UL | 0.028 | 0.332 | -1.19 | 0.868 |
| | | AN_Gufudalsá:PI_TH | 0.110 | 0.320 | 0.07 | 0.487 |
| | | AN_Gufudalsá:PI_UL | 0.213 | 0.575 | 0.19 | 0.445 |
| | | AN_Gufudalsá:PL_TH | 0.243 | 0.318 | 1.13 | 0.140 |
| | | AN_Gufudalsá:PL_UL | 0.124 | 0.366 | 0.04 | 0.494 |
| | | AN_Gufudalsá:SB_TH | 0.209 | 0.311 | 0.95 | 0.182 |
| | Lambeyrarkvísl to lake morphs | AN_Lambeyrarkvísl:Hest | 0.384 | 0.598 | 0.89 | 0.194 |
| | | AN_Lambeyrarkvísl:LB_TH | 0.202 | 0.620 | 0.06 | 0.487 |
| | | AN_Lambeyrarkvísl:LB_UL | 0.229 | 0.614 | 0.19 | 0.434 |
| | | AN_Lambeyrarkvísl:PI_TH | 0.311 | 0.585 | 0.59 | 0.297 |
| | | AN_Lambeyrarkvísl:PI_UL | 0.012 | 0.759 | -1.92 | 0.972 |
| | | AN_Lambeyrarkvísl:PL_TH | 0.444 | 0.588 | 1.13 | 0.143 |
| | | AN_Lambeyrarkvísl:PL_UL | 0.325 | 0.603 | 0.62 | 0.286 |
| | | AN_Lambeyrarkvísl:SB_TH | 0.410 | 0.592 | 1.01 | 0.172 |
| | Víðidalsá to lake morphs | AN_Víðidalsá:Hest | 0.602 | 0.369 | 2.55 | ***0.002** |
| | | AN_Víðidalsá:LB_TH | 0.420 | 0.372 | 1.80 | 0.027 |
| | | AN_Víðidalsá:LB_UL | 0.447 | 0.391 | 1.84 | 0.020 |
| | | AN_Víðidalsá:PI_TH | 0.528 | 0.371 | 2.35 | ***0.005** |
| | | AN_Víðidalsá:PI_UL | 0.206 | 0.555 | 0.07 | 0.489 |
| | | AN_Víðidalsá:PL_TH | 0.661 | 0.364 | 2.77 | ***0.002** |
| | | AN_Víðidalsá:PL_UL | 0.543 | 0.410 | 2.12 | 0.010 |
| | | AN_Víðidalsá:SB_TH | 0.628 | 0.366 | 2.70 | ***0.002** |
| | Vífisdalsá to lake morphs | AN_Vífilsdalsá:Hest | 0.280 | 0.509 | 0.62 | 0.297 |
| | | AN_Vífilsdalsá:LB_TH | 0.098 | 0.519 | -0.56 | 0.706 |
| | | AN_Vífilsdalsá:LB_UL | 0.125 | 0.507 | -0.41 | 0.654 |
| | | AN_Vífilsdalsá:PI_TH | 0.206 | 0.499 | 0.19 | 0.448 |
| | | AN_Vífilsdalsá:PI_UL | 0.116 | 0.701 | -0.68 | 0.741 |
| | | AN_Vífilsdalsá:PL_TH | 0.339 | 0.498 | 0.91 | 0.196 |
| | | AN_Vífilsdalsá:PL_UL | 0.221 | 0.551 | 0.23 | 0.423 |
| | | AN_Vífilsdalsá:SB_TH | 0.306 | 0.497 | 0.76 | 0.242 |
| | Hestvatn | Hest:LB_TH | 0.182 | 0.189 | 1.48 | 0.059 |
| | | Hest:LB_UL | 0.155 | 0.200 | 1.11 | 0.148 |
| | | Hest:PI_TH | 0.074 | 0.167 | 0.33 | 0.388 |
| | | Hest:PI_UL | 0.396 | 0.466 | 1.26 | 0.097 |
| | | Hest:PL_TH | 0.059 | 0.162 | 0.13 | 0.465 |
| | | Hest:PL_UL | 0.059 | 0.241 | -0.33 | 0.641 |
| | | Hest:SB_TH | 0.026 | 0.149 | -0.64 | 0.726 |
| | Þingvallavatn | LB_TH:PI_TH | 0.109 | 0.182 | 0.75 | 0.248 |
| | | LB_TH:PL_TH | 0.242 | 0.178 | 2.15 | 0.013 |
| | | LB_TH:SB_TH | 0.208 | 0.172 | 1.99 | 0.015 |
| | | PI_TH:PL_TH | 0.133 | 0.156 | 1.29 | 0.100 |
| | | PI_TH:SB_TH | 0.100 | 0.141 | 1.02 | 0.168 |
| | | PL_TH:SB_TH | 0.033 | 0.127 | -0.29 | 0.615 |
| | Úlfljótsvatn | LB_UL:PI_UL | 0.241 | 0.504 | 0.46 | 0.334 |
| | | LB_UL:PL_UL | 0.096 | 0.271 | 0.12 | 0.463 |

| | | | | | | |
|---|---|---|---|---|---|---|
| | | PI_UL:PL_UL | 0.337 | 0.494 | 0.92 | 0.196 |
| | Across Þingvallavatn and Úlfljótsvatn | LB_TH:LB_UL | 0.027 | 0.208 | -0.98 | 0.823 |
| | | LB_TH:PI_UL | 0.214 | 0.475 | 0.35 | 0.385 |
| | | LB_TH:PL_UL | 0.123 | 0.245 | 0.51 | 0.327 |
| | | LB_UL:PI_TH | 0.081 | 0.199 | 0.18 | 0.452 |
| | | LB_UL:PL_TH | 0.214 | 0.188 | 1.81 | 0.027 |
| | | LB_UL:SB_TH | 0.181 | 0.183 | 1.56 | 0.054 |
| | | PI_TH:PI_UL | 0.322 | 0.470 | 0.95 | 0.183 |
| | | PI_TH:PL_UL | 0.015 | 0.243 | -1.35 | 0.899 |
| | | PI_UL:PL_TH | 0.455 | 0.458 | 1.53 | 0.052 |
| | | PI_UL:SB_TH | 0.422 | 0.462 | 1.40 | 0.086 |
| | | PL_TH:PL_UL | 0.118 | 0.238 | 0.54 | 0.306 |
| | | PL_UL:SB_TH | 0.085 | 0.219 | 0.16 | 0.454 |

D = pairwise distances between mean; UCL = upper confidence level, Z = Z-statistic; P = P-value, ≤ 0.008 in bold with asterisk.

***Supplementary material 10**: Results from "Significant Vector Angle test" examining possibly allometric bone size differences by population for five bones, that showed significant allometric differences according to ANOVA model with main and interaction effects (population x body size). Used to verify results from ANOVA and test for true allometry (i.e., different group slopes). Hítarvatn fish excluded from this analysis. AN-charr treated as five separate populations.*

| Bone | System | Pairs | R | Angle | UCL | Z | P-value |
|---|---|---|---|---|---|---|---|
| Dentary | AN populations | AN_Brúará:AN_Gufudalsá | 1.000 | 0.000 | 0.000 | -0.05 | 1.000 |
| | | AN_Brúará:AN_Lambeyrarkvísl | 1.000 | 0.000 | 0.000 | -0.06 | 1.000 |
| | | AN_Brúará:AN_Víðidalsá | 1.000 | 0.000 | 0.000 | -0.05 | 1.000 |
| | | AN_Brúará:AN_Vífilsdalsá | 1.000 | 0.000 | 0.000 | -0.05 | 1.000 |
| | | AN_Gufudalsá:AN_Lambeyrarkvísl | 1.000 | 0.000 | 0.000 | -0.04 | 1.000 |
| | | AN_Gufudalsá:AN_Víðidalsá | 1.000 | 0.000 | 0.000 | -0.50 | 1.000 |
| | | AN_Gufudalsá:AN_Vífilsdalsá | 1.000 | 0.000 | 0.000 | -0.54 | 1.000 |
| | | AN_Lambeyrarkvísl:AN_Víðidalsá | 1.000 | 0.000 | 0.000 | -0.04 | 1.000 |
| | | AN_Lambeyrarkvísl:AN_Vífilsdalsá | 1.000 | 0.000 | 0.000 | -0.04 | 1.000 |
| | | AN_Víðidalsá:AN_Vífilsdalsá | 1.000 | 0.000 | 0.000 | -0.55 | 1.000 |
| | Brúará to lake morphs | AN_Brúará:Hest | 1.000 | 0.000 | 0.000 | -0.05 | 1.000 |
| | | AN_Brúará:LB_TH | 1.000 | 0.000 | 0.000 | -0.05 | 1.000 |
| | | AN_Brúará:LB_UL | 1.000 | 0.000 | 0.000 | -0.05 | 1.000 |
| | | AN_Brúará:PI_TH | 1.000 | 0.000 | 0.000 | -0.05 | 1.000 |
| | | AN_Brúará:PI_UL | 1.000 | 0.000 | 0.000 | -0.05 | 1.000 |
| | | AN_Brúará:PL_TH | 1.000 | 0.000 | 0.000 | -0.05 | 1.000 |
| | | AN_Brúará:PL_UL | 1.000 | 0.000 | 0.000 | -0.05 | 1.000 |
| | | AN_Brúará:SB_TH | 1.000 | 0.000 | 0.000 | -0.05 | 1.000 |
| | Gufudalsá to lake morphs | AN_Gufudalsá:Hest | 1.000 | 0.000 | 0.000 | -0.52 | 1.000 |
| | | AN_Gufudalsá:LB_TH | 1.000 | 0.000 | 0.000 | -0.55 | 1.000 |
| | | AN_Gufudalsá:LB_UL | 1.000 | 0.000 | 0.000 | -0.50 | 1.000 |
| | | AN_Gufudalsá:PI_TH | 1.000 | 0.000 | 0.000 | -0.50 | 1.000 |
| | | AN_Gufudalsá:PI_UL | 1.000 | 0.000 | 0.000 | -0.51 | 1.000 |
| | | AN_Gufudalsá:PL_TH | 1.000 | 0.000 | 0.000 | -0.51 | 1.000 |
| | | AN_Gufudalsá:PL_UL | 1.000 | 0.000 | 0.000 | -0.52 | 1.000 |
| | | AN_Gufudalsá:SB_TH | 1.000 | 0.000 | 0.000 | -0.49 | 1.000 |
| | Lambeyrarkvísl to lake morphs | AN_Lambeyrarkvísl:Hest | 1.000 | 0.000 | 0.000 | -0.04 | 1.000 |
| | | AN_Lambeyrarkvísl:LB_TH | 1.000 | 0.000 | 0.000 | -0.04 | 1.000 |
| | | AN_Lambeyrarkvísl:LB_UL | 1.000 | 0.000 | 0.000 | -0.04 | 1.000 |
| | | AN_Lambeyrarkvísl:PI_TH | 1.000 | 0.000 | 0.000 | -0.04 | 1.000 |
| | | AN_Lambeyrarkvísl:PI_UL | 1.000 | 0.000 | 0.000 | -0.04 | 1.000 |
| | | AN_Lambeyrarkvísl:PL_TH | 1.000 | 0.000 | 0.000 | -0.04 | 1.000 |
| | | AN_Lambeyrarkvísl:PL_UL | 1.000 | 0.000 | 0.000 | -0.04 | 1.000 |

| | | | | | | | |
|---|---|---|---|---|---|---|---|
| | | AN_Lambeyrarkvísl:SB_TH | 1.000 | 0.000 | 0.000 | -0.04 | 1.000 |
| | Víðidalsá to lake morphs | AN_Víðidalsá:Hest | 1.000 | 0.000 | 0.000 | -0.53 | 1.000 |
| | | AN_Víðidalsá:LB_TH | 1.000 | 0.000 | 0.000 | -0.56 | 1.000 |
| | | AN_Víðidalsá:LB_UL | 1.000 | 0.000 | 0.000 | -0.52 | 1.000 |
| | | AN_Víðidalsá:PI_TH | 1.000 | 0.000 | 0.000 | -0.50 | 1.000 |
| | | AN_Víðidalsá:PI_UL | 1.000 | 0.000 | 0.000 | -0.52 | 1.000 |
| | | AN_Víðidalsá:PL_TH | 1.000 | 0.000 | 0.000 | -0.52 | 1.000 |
| | | AN_Víðidalsá:PL_UL | 1.000 | 0.000 | 0.000 | -0.53 | 1.000 |
| | | AN_Víðidalsá:SB_TH | 1.000 | 0.000 | 0.000 | -0.51 | 1.000 |
| | Vífisdalsá to lake morphs | AN_Vífilsdalsá:Hest | 1.000 | 0.000 | 0.000 | -0.53 | 1.000 |
| | | AN_Vífilsdalsá:LB_TH | 1.000 | 0.000 | 0.000 | -0.58 | 1.000 |
| | | AN_Vífilsdalsá:LB_UL | 1.000 | 0.000 | 0.000 | -0.53 | 1.000 |
| | | AN_Vífilsdalsá:PI_TH | 1.000 | 0.000 | 0.000 | -0.52 | 1.000 |
| | | AN_Vífilsdalsá:PI_UL | 1.000 | 0.000 | 0.000 | -0.53 | 1.000 |
| | | AN_Vífilsdalsá:PL_TH | 1.000 | 0.000 | 0.000 | -0.53 | 1.000 |
| | | AN_Vífilsdalsá:PL_UL | 1.000 | 0.000 | 0.000 | -0.54 | 1.000 |
| | | AN_Vífilsdalsá:SB_TH | 1.000 | 0.000 | 0.000 | -0.51 | 1.000 |
| | Hestvatn | Hest:LB_TH | 1.000 | 0.000 | 0.000 | -0.56 | 1.000 |
| | | Hest:LB_UL | 1.000 | 0.000 | 0.000 | -0.51 | 1.000 |
| | | Hest:PI_TH | 1.000 | 0.000 | 0.000 | -0.50 | 1.000 |
| | | Hest:PI_UL | 1.000 | 0.000 | 0.000 | -0.53 | 1.000 |
| | | Hest:PL_TH | 1.000 | 0.000 | 0.000 | -0.53 | 1.000 |
| | | Hest:PL_UL | 1.000 | 0.000 | 0.000 | -0.52 | 1.000 |
| | | Hest:SB_TH | 1.000 | 0.000 | 0.000 | -0.50 | 1.000 |
| | Þingvallavatn | LB_TH:PI_TH | 1.000 | 0.000 | 0.000 | -0.54 | 1.000 |
| | | LB_TH:PL_TH | 1.000 | 0.000 | 0.000 | -0.56 | 1.000 |
| | | LB_TH:SB_TH | 1.000 | 0.000 | 0.000 | -0.53 | 1.000 |
| | | PI_TH:PL_TH | 1.000 | 0.000 | 0.000 | -0.49 | 1.000 |
| | | PI_TH:SB_TH | 1.000 | 0.000 | 0.000 | -0.48 | 1.000 |
| | | PL_TH:SB_TH | 1.000 | 0.000 | 0.000 | -0.49 | 1.000 |
| | Úlfljótsvatn | LB_UL:PI_UL | 1.000 | 0.000 | 0.000 | -0.51 | 1.000 |
| | | LB_UL:PL_UL | 1.000 | 0.000 | 0.000 | -0.51 | 1.000 |
| | | PI_UL:PL_UL | 1.000 | 0.000 | 0.000 | -0.52 | 1.000 |
| | Across Þingvallavatn and Úlfljótsvatn | LB_TH:LB_UL | 1.000 | 0.000 | 0.000 | -0.55 | 1.000 |
| | | LB_TH:PI_UL | 1.000 | 0.000 | 0.000 | -0.55 | 1.000 |
| | | LB_TH:PL_UL | 1.000 | 0.000 | 0.000 | -0.55 | 1.000 |
| | | LB_UL:PI_TH | 1.000 | 0.000 | 0.000 | -0.50 | 1.000 |
| | | LB_UL:PL_TH | 1.000 | 0.000 | 0.000 | -0.52 | 1.000 |
| | | LB_UL:SB_TH | 1.000 | 0.000 | 0.000 | -0.49 | 1.000 |
| | | PI_TH:PI_UL | 1.000 | 0.000 | 0.000 | -0.50 | 1.000 |
| | | PI_TH:PL_UL | 1.000 | 0.000 | 0.000 | -0.51 | 1.000 |
| | | PI_UL:PL_TH | 1.000 | 0.000 | 0.000 | -0.50 | 1.000 |
| | | PI_UL:SB_TH | 1.000 | 0.000 | 0.000 | -0.48 | 1.000 |
| | | PL_TH:PL_UL | 1.000 | 0.000 | 0.000 | -0.52 | 1.000 |
| | | PL_UL:SB_TH | 1.000 | 0.000 | 0.000 | -0.49 | 1.000 |
| Articular-angular | AN populations | AN_Brúará:AN_Gufudalsá | 1.000 | 0.000 | 0.000 | -0.54 | 1.000 |
| | | AN_Brúará:AN_Lambeyrarkvísl | 1.000 | 0.000 | 0.000 | -0.55 | 1.000 |
| | | AN_Brúará:AN_Víðidalsá | 1.000 | 0.000 | 0.000 | -0.56 | 1.000 |
| | | AN_Brúará:AN_Vífilsdalsá | 1.000 | 0.000 | 0.000 | -0.57 | 1.000 |
| | | AN_Gufudalsá:AN_Lambeyrarkvísl | 1.000 | 0.000 | 0.000 | -0.57 | 1.000 |
| | | AN_Gufudalsá:AN_Víðidalsá | 1.000 | 0.000 | 0.000 | -0.58 | 1.000 |
| | | AN_Gufudalsá:AN_Vífilsdalsá | 1.000 | 0.000 | 0.000 | -0.58 | 1.000 |
| | | AN_Lambeyrarkvísl:AN_Víðidalsá | 1.000 | 0.000 | 0.000 | -0.59 | 1.000 |
| | | AN_Lambeyrarkvísl:AN_Vífilsdalsá | 1.000 | 0.000 | 0.000 | -0.59 | 1.000 |
| | | AN_Víðidalsá:AN_Vífilsdalsá | 1.000 | 0.000 | 0.000 | -0.59 | 1.000 |
| | Brúará to lake morphs | AN_Brúará:Hest | 1.000 | 0.000 | 0.000 | -0.57 | 1.000 |
| | | AN_Brúará:LB_TH | 1.000 | 0.000 | 0.000 | -0.60 | 1.000 |
| | | AN_Brúará:LB_UL | 1.000 | 0.000 | 0.000 | -0.56 | 1.000 |
| | | AN_Brúará:PI_TH | 1.000 | 0.000 | 0.000 | -0.55 | 1.000 |
| | | AN_Brúará:PI_UL | 1.000 | 0.000 | 0.000 | -0.55 | 1.000 |
| | | AN_Brúará:PL_TH | 1.000 | 0.000 | 0.000 | -0.58 | 1.000 |

| | | | | | | | |
|---|---|---|---|---|---|---|---|
| | | AN_Brúará:PL_UL | 1.000 | 0.000 | 0.000 | -0.60 | 1.000 |
| | | AN_Brúará:SB_TH | 1.000 | 0.000 | 0.000 | -0.60 | 1.000 |
| | Gufudalsá to lake morphs | AN_Gufudalsá:Hest | 1.000 | 0.000 | 0.000 | -0.57 | 1.000 |
| | | AN_Gufudalsá:LB_TH | 1.000 | 0.000 | 0.000 | -0.60 | 1.000 |
| | | AN_Gufudalsá:LB_UL | 1.000 | 0.000 | 0.000 | -0.57 | 1.000 |
| | | AN_Gufudalsá:PI_TH | 1.000 | 0.000 | 0.000 | -0.56 | 1.000 |
| | | AN_Gufudalsá:PI_UL | 1.000 | 0.000 | 0.000 | -0.58 | 1.000 |
| | | AN_Gufudalsá:PL_TH | 1.000 | 0.000 | 0.000 | -0.60 | 1.000 |
| | | AN_Gufudalsá:PL_UL | 1.000 | 0.000 | 0.000 | -0.60 | 1.000 |
| | | AN_Gufudalsá:SB_TH | 1.000 | 0.000 | 0.000 | -0.61 | 1.000 |
| | Lambeyrarkvísl to lake morphs | AN_Lambeyrarkvísl:Hest | 1.000 | 0.000 | 0.000 | -0.59 | 1.000 |
| | | AN_Lambeyrarkvísl:LB_TH | 1.000 | 0.000 | 0.000 | -0.60 | 1.000 |
| | | AN_Lambeyrarkvísl:LB_UL | 1.000 | 0.000 | 0.000 | -0.58 | 1.000 |
| | | AN_Lambeyrarkvísl:PI_TH | 1.000 | 0.000 | 0.000 | -0.57 | 1.000 |
| | | AN_Lambeyrarkvísl:PI_UL | 1.000 | 0.000 | 0.000 | -0.58 | 1.000 |
| | | AN_Lambeyrarkvísl:PL_TH | 1.000 | 0.000 | 0.000 | -0.61 | 1.000 |
| | | AN_Lambeyrarkvísl:PL_UL | 1.000 | 0.000 | 0.000 | -0.62 | 1.000 |
| | | AN_Lambeyrarkvísl:SB_TH | 1.000 | 0.000 | 0.000 | -0.62 | 1.000 |
| | Víðidalsá to lake morphs | AN_Víðidalsá:Hest | 1.000 | 0.000 | 0.000 | -0.59 | 1.000 |
| | | AN_Víðidalsá:LB_TH | 1.000 | 0.000 | 0.000 | -0.61 | 1.000 |
| | | AN_Víðidalsá:LB_UL | 1.000 | 0.000 | 0.000 | -0.59 | 1.000 |
| | | AN_Víðidalsá:PI_TH | 1.000 | 0.000 | 0.000 | -0.58 | 1.000 |
| | | AN_Víðidalsá:PI_UL | 1.000 | 0.000 | 0.000 | -0.57 | 1.000 |
| | | AN_Víðidalsá:PL_TH | 1.000 | 0.000 | 0.000 | -0.60 | 1.000 |
| | | AN_Víðidalsá:PL_UL | 1.000 | 0.000 | 0.000 | -0.62 | 1.000 |
| | | AN_Víðidalsá:SB_TH | 1.000 | 0.000 | 0.000 | -0.63 | 1.000 |
| | Vífisdalsá to lake morphs | AN_Vífilsdalsá:Hest | 1.000 | 0.000 | 0.000 | -0.59 | 1.000 |
| | | AN_Vífilsdalsá:LB_TH | 1.000 | 0.000 | 0.000 | -0.63 | 1.000 |
| | | AN_Vífilsdalsá:LB_UL | 1.000 | 0.000 | 0.000 | -0.59 | 1.000 |
| | | AN_Vífilsdalsá:PI_TH | 1.000 | 0.000 | 0.000 | -0.57 | 1.000 |
| | | AN_Vífilsdalsá:PI_UL | 1.000 | 0.000 | 0.000 | -0.59 | 1.000 |
| | | AN_Vífilsdalsá:PL_TH | 1.000 | 0.000 | 0.000 | -0.60 | 1.000 |
| | | AN_Vífilsdalsá:PL_UL | 1.000 | 0.000 | 0.000 | -0.62 | 1.000 |
| | | AN_Vífilsdalsá:SB_TH | 1.000 | 0.000 | 0.000 | -0.63 | 1.000 |
| | Hestvatn | Hest:LB_TH | 1.000 | 0.000 | 0.000 | -0.62 | 1.000 |
| | | Hest:LB_UL | 1.000 | 0.000 | 0.000 | -0.59 | 1.000 |
| | | Hest:PI_TH | 1.000 | 0.000 | 0.000 | -0.57 | 1.000 |
| | | Hest:PI_UL | 1.000 | 0.000 | 0.000 | -0.58 | 1.000 |
| | | Hest:PL_TH | 1.000 | 0.000 | 0.000 | -0.60 | 1.000 |
| | | Hest:PL_UL | 1.000 | 0.000 | 0.000 | -0.62 | 1.000 |
| | | Hest:SB_TH | 1.000 | 0.000 | 0.000 | -0.63 | 1.000 |
| | Þingvallavatn | LB_TH:PI_TH | 1.000 | 0.000 | 0.000 | -0.60 | 1.000 |
| | | LB_TH:PL_TH | 1.000 | 0.000 | 0.000 | -0.64 | 1.000 |
| | | LB_TH:SB_TH | 1.000 | 0.000 | 0.000 | -0.65 | 1.000 |
| | | PI_TH:PL_TH | 1.000 | 0.000 | 0.000 | -0.60 | 1.000 |
| | | PI_TH:SB_TH | 1.000 | 0.000 | 0.000 | -0.61 | 1.000 |
| | | PL_TH:SB_TH | 1.000 | 0.000 | 0.000 | -0.65 | 1.000 |
| | Úlfljótsvatn | LB_UL:PI_UL | 1.000 | 0.000 | 0.000 | -0.58 | 1.000 |
| | | LB_UL:PL_UL | 1.000 | 0.000 | 0.000 | -0.63 | 1.000 |
| | | PI_UL:PL_UL | 1.000 | 0.000 | 0.000 | -0.61 | 1.000 |
| | Across Þingvallavatn and Úlfljótsvatn | LB_TH:LB_UL | 1.000 | 0.000 | 0.000 | -0.62 | 1.000 |
| | | LB_TH:PI_UL | 1.000 | 0.000 | 0.000 | -0.62 | 1.000 |
| | | LB_TH:PL_UL | 1.000 | 0.000 | 0.000 | -0.64 | 1.000 |
| | | LB_UL:PI_TH | 1.000 | 0.000 | 0.000 | -0.58 | 1.000 |
| | | LB_UL:PL_TH | 1.000 | 0.000 | 0.000 | -0.61 | 1.000 |
| | | LB_UL:SB_TH | 1.000 | 0.000 | 0.000 | -0.61 | 1.000 |
| | | PI_TH:PI_UL | 1.000 | 0.000 | 0.000 | -0.58 | 1.000 |
| | | PI_TH:PL_UL | 1.000 | 0.000 | 0.000 | -0.61 | 1.000 |
| | | PI_UL:PL_TH | 1.000 | 0.000 | 0.000 | -0.60 | 1.000 |
| | | PI_UL:SB_TH | 1.000 | 0.000 | 0.000 | -0.61 | 1.000 |
| | | PL_TH:PL_UL | 1.000 | 0.000 | 0.000 | -0.64 | 1.000 |

| | | | | | | | |
|---|---|---|---|---|---|---|---|
| | | PL_UL:SB_TH | 1.000 | 0.000 | 0.000 | -0.66 | 1.000 |
| Quadrate | AN populations | AN_Brúará:AN_Gufudalsá | 1.000 | 0.000 | 0.000 | -0.08 | 1.000 |
| | | AN_Brúará:AN_Lambeyrarkvísl | 1.000 | 0.000 | 0.000 | -0.08 | 1.000 |
| | | AN_Brúará:AN_Víðidalsá | 1.000 | 0.000 | 0.000 | -0.08 | 1.000 |
| | | AN_Brúará:AN_Vífilsdalsá | 1.000 | 0.000 | 0.000 | -0.07 | 0.247 |
| | | AN_Gufudalsá:AN_Lambeyrarkvísl | 1.000 | 0.000 | 0.000 | -0.58 | 1.000 |
| | | AN_Gufudalsá:AN_Víðidalsá | 1.000 | 0.000 | 0.000 | -0.57 | 1.000 |
| | | AN_Gufudalsá:AN_Vífilsdalsá | 1.000 | 0.000 | 0.000 | 1.59 | 0.267 |
| | | AN_Lambeyrarkvísl:AN_Víðidalsá | 1.000 | 0.000 | 0.000 | -0.56 | 1.000 |
| | | AN_Lambeyrarkvísl:AN_Vífilsdalsá | 1.000 | 0.000 | 0.000 | 1.67 | 0.240 |
| | | AN_Víðidalsá:AN_Vífilsdalsá | 1.000 | 0.000 | 0.000 | 1.68 | 0.248 |
| | Brúará to lake morphs | AN_Brúará:Hest | 1.000 | 0.000 | 0.000 | -0.07 | 0.238 |
| | | AN_Brúará:LB_TH | 1.000 | 0.000 | 0.000 | -0.08 | 1.000 |
| | | AN_Brúará:LB_UL | 1.000 | 0.000 | 0.000 | -0.08 | 1.000 |
| | | AN_Brúará:PI_TH | 1.000 | 0.000 | 0.000 | -0.08 | 1.000 |
| | | AN_Brúará:PI_UL | 1.000 | 0.000 | 0.000 | -0.08 | 1.000 |
| | | AN_Brúará:PL_TH | 1.000 | 0.000 | 0.000 | -0.08 | 1.000 |
| | | AN_Brúará:PL_UL | 1.000 | 0.000 | 0.000 | -0.08 | 1.000 |
| | | AN_Brúará:SB_TH | 1.000 | 0.000 | 0.000 | -0.07 | 0.227 |
| | Gufudalsá to lake morphs | AN_Gufudalsá:Hest | 1.000 | 0.000 | 0.000 | 1.65 | 0.250 |
| | | AN_Gufudalsá:LB_TH | 1.000 | 0.000 | 0.000 | -0.56 | 1.000 |
| | | AN_Gufudalsá:LB_UL | 1.000 | 0.000 | 0.000 | -0.58 | 1.000 |
| | | AN_Gufudalsá:PI_TH | 1.000 | 0.000 | 0.000 | -0.59 | 1.000 |
| | | AN_Gufudalsá:PI_UL | 1.000 | 0.000 | 0.000 | -0.60 | 1.000 |
| | | AN_Gufudalsá:PL_TH | 1.000 | 0.000 | 0.000 | -0.55 | 1.000 |
| | | AN_Gufudalsá:PL_UL | 1.000 | 0.000 | 0.000 | -0.58 | 1.000 |
| | | AN_Gufudalsá:SB_TH | 1.000 | 0.000 | 0.000 | 1.72 | 0.230 |
| | Lambeyrarkvísl to lake morphs | AN_Lambeyrarkvísl:Hest | 1.000 | 0.000 | 0.000 | 1.74 | 0.232 |
| | | AN_Lambeyrarkvísl:LB_TH | 1.000 | 0.000 | 0.000 | -0.52 | 1.000 |
| | | AN_Lambeyrarkvísl:LB_UL | 1.000 | 0.000 | 0.000 | -0.56 | 1.000 |
| | | AN_Lambeyrarkvísl:PI_TH | 1.000 | 0.000 | 0.000 | -0.55 | 1.000 |
| | | AN_Lambeyrarkvísl:PI_UL | 1.000 | 0.000 | 0.000 | -0.56 | 1.000 |
| | | AN_Lambeyrarkvísl:PL_TH | 1.000 | 0.000 | 0.000 | -0.52 | 1.000 |
| | | AN_Lambeyrarkvísl:PL_UL | 1.000 | 0.000 | 0.000 | -0.57 | 1.000 |
| | | AN_Lambeyrarkvísl:SB_TH | 1.000 | 0.000 | 0.000 | 1.82 | 0.211 |
| | Víðidalsá to lake morphs | AN_Víðidalsá:Hest | 1.000 | 0.000 | 0.000 | 1.76 | 0.230 |
| | | AN_Víðidalsá:LB_TH | 1.000 | 0.000 | 0.000 | -0.53 | 1.000 |
| | | AN_Víðidalsá:LB_UL | 1.000 | 0.000 | 0.000 | -0.55 | 1.000 |
| | | AN_Víðidalsá:PI_TH | 1.000 | 0.000 | 0.000 | -0.56 | 1.000 |
| | | AN_Víðidalsá:PI_UL | 1.000 | 0.000 | 0.000 | -0.55 | 1.000 |
| | | AN_Víðidalsá:PL_TH | 1.000 | 0.000 | 0.000 | -0.53 | 1.000 |
| | | AN_Víðidalsá:PL_UL | 1.000 | 0.000 | 0.000 | -0.57 | 1.000 |
| | | AN_Víðidalsá:SB_TH | 1.000 | 0.000 | 0.000 | 1.84 | 0.215 |
| | Vífisdalsá to lake morphs | AN_Vífilsdalsá:Hest | 1.000 | 0.000 | 0.000 | 2.63 | 0.016 |
| | | AN_Vífilsdalsá:LB_TH | 1.000 | 0.000 | 0.000 | 1.73 | 0.240 |
| | | AN_Vífilsdalsá:LB_UL | 1.000 | 0.000 | 0.000 | 1.66 | 0.247 |
| | | AN_Vífilsdalsá:PI_TH | 1.000 | 0.000 | 0.000 | 1.66 | 0.245 |
| | | AN_Vífilsdalsá:PI_UL | 1.000 | 0.000 | 0.000 | 1.64 | 0.259 |
| | | AN_Vífilsdalsá:PL_TH | 1.000 | 0.000 | 0.000 | 1.77 | 0.225 |
| | | AN_Vífilsdalsá:PL_UL | 1.000 | 0.000 | 0.000 | 1.65 | 0.258 |
| | | AN_Vífilsdalsá:SB_TH | 1.000 | 0.000 | 0.000 | 2.72 | 0.017 |
| | Hestvatn | Hest:LB_TH | 1.000 | 0.000 | 0.000 | 1.81 | 0.223 |
| | | Hest:LB_UL | 1.000 | 0.000 | 0.000 | 1.73 | 0.236 |
| | | Hest:PI_TH | 1.000 | 0.000 | 0.000 | 1.73 | 0.236 |
| | | Hest:PI_UL | 1.000 | 0.000 | 0.000 | 1.71 | 0.241 |
| | | Hest:PL_TH | 1.000 | 0.000 | 0.000 | 1.85 | 0.216 |
| | | Hest:PL_UL | 1.000 | 0.000 | 0.000 | 1.72 | 0.238 |
| | | Hest:SB_TH | 1.000 | 0.000 | 0.000 | 2.83 | 0.017 |
| | Þingvallavatn | LB_TH:PI_TH | 1.000 | 0.000 | 0.000 | -0.54 | 1.000 |
| | | LB_TH:PL_TH | 1.000 | 0.000 | 0.000 | -0.50 | 1.000 |
| | | LB_TH:SB_TH | 1.000 | 0.000 | 0.000 | 1.89 | 0.209 |

| | | | | | | | |
|---|---|---|---|---|---|---|---|
| | | PI_TH:PL_TH | 1.000 | 0.000 | 0.000 | -0.52 | 1.000 |
| | | PI_TH:SB_TH | 1.000 | 0.000 | 0.000 | 1.81 | 0.222 |
| | | PL_TH:SB_TH | 1.000 | 0.000 | 0.000 | 1.94 | 0.198 |
| | Úlfljótsvatn | LB_UL:PI_UL | 1.000 | 0.000 | 0.000 | -0.57 | 1.000 |
| | | LB_UL:PL_UL | 1.000 | 0.000 | 0.000 | -0.57 | 1.000 |
| | | PI_UL:PL_UL | 1.000 | 0.000 | 0.000 | -0.57 | 1.000 |
| | Across Þingvallavatn and Úlfljótsvatn | LB_TH:LB_UL | 1.000 | 0.000 | 0.000 | -0.54 | 1.000 |
| | | LB_TH:PI_UL | 1.000 | 0.000 | 0.000 | -0.54 | 1.000 |
| | | LB_TH:PL_UL | 1.000 | 0.000 | 0.000 | -0.54 | 1.000 |
| | | LB_UL:PI_TH | 1.000 | 0.000 | 0.000 | -0.57 | 1.000 |
| | | LB_UL:PL_TH | 1.000 | 0.000 | 0.000 | -0.53 | 1.000 |
| | | LB_UL:SB_TH | 1.000 | 0.000 | 0.000 | 1.80 | 0.220 |
| | | PI_TH:PI_UL | 1.000 | 0.000 | 0.000 | -0.56 | 1.000 |
| | | PI_TH:PL_UL | 1.000 | 0.000 | 0.000 | -0.57 | 1.000 |
| | | PI_UL:PL_TH | 1.000 | 0.000 | 0.000 | -0.54 | 1.000 |
| | | PI_UL:SB_TH | 1.000 | 0.000 | 0.000 | 1.78 | 0.224 |
| | | PL_TH:PL_UL | 1.000 | 0.000 | 0.000 | -0.53 | 1.000 |
| | | PL_UL:SB_TH | 1.000 | 0.000 | 0.000 | 1.79 | 0.221 |
| Premaxilla | AN populations | AN_Brúará:AN_Gufudalsá | 1.000 | 0.000 | 0.000 | -0.10 | 1.000 |
| | | AN_Brúará:AN_Lambeyrarkvísl | 1.000 | 0.000 | 0.000 | -0.10 | 1.000 |
| | | AN_Brúará:AN_Víðidalsá | 1.000 | 0.000 | 0.000 | -0.09 | 0.275 |
| | | AN_Brúará:AN_Vífilsdalsá | 1.000 | 0.000 | 0.000 | -0.10 | 1.000 |
| | | AN_Gufudalsá:AN_Lambeyrarkvísl | 1.000 | 0.000 | 0.000 | -0.55 | 1.000 |
| | | AN_Gufudalsá:AN_Víðidalsá | 1.000 | 0.000 | 0.000 | 1.80 | 0.220 |
| | | AN_Gufudalsá:AN_Vífilsdalsá | 1.000 | 0.000 | 0.000 | -0.54 | 1.000 |
| | | AN_Lambeyrarkvísl:AN_Víðidalsá | 1.000 | 0.000 | 0.000 | 1.69 | 0.241 |
| | | AN_Lambeyrarkvísl:AN_Vífilsdalsá | 1.000 | 0.000 | 0.000 | -0.55 | 1.000 |
| | | AN_Víðidalsá:AN_Vífilsdalsá | 1.000 | 0.000 | 0.000 | 1.73 | 0.233 |
| | Brúará to lake morphs | AN_Brúará:Hest | 1.000 | 0.000 | 0.000 | -0.10 | 1.000 |
| | | AN_Brúará:LB_TH | 1.000 | 0.000 | 0.000 | -0.10 | 1.000 |
| | | AN_Brúará:LB_UL | 1.000 | 0.000 | 0.000 | -0.10 | 1.000 |
| | | AN_Brúará:PI_TH | 1.000 | 0.000 | 0.000 | -0.10 | 1.000 |
| | | AN_Brúará:PI_UL | 1.000 | 0.000 | 0.000 | -0.10 | 1.000 |
| | | AN_Brúará:PL_TH | 1.000 | 0.000 | 0.000 | -0.10 | 1.000 |
| | | AN_Brúará:PL_UL | 1.000 | 0.000 | 0.000 | -0.10 | 1.000 |
| | | AN_Brúará:SB_TH | 1.000 | 0.000 | 0.000 | -0.09 | 0.202 |
| | Gufudalsá to lake morphs | AN_Gufudalsá:Hest | 1.000 | 0.000 | 0.000 | -0.44 | 1.000 |
| | | AN_Gufudalsá:LB_TH | 1.000 | 0.000 | 0.000 | -0.46 | 1.000 |
| | | AN_Gufudalsá:LB_UL | 1.000 | 0.000 | 0.000 | -0.47 | 1.000 |
| | | AN_Gufudalsá:PI_TH | 1.000 | 0.000 | 0.000 | -0.47 | 1.000 |
| | | AN_Gufudalsá:PI_UL | 1.000 | 0.000 | 0.000 | -0.54 | 1.000 |
| | | AN_Gufudalsá:PL_TH | 1.000 | 0.000 | 0.000 | -0.43 | 1.000 |
| | | AN_Gufudalsá:PL_UL | 1.000 | 0.000 | 0.000 | -0.51 | 1.000 |
| | | AN_Gufudalsá:SB_TH | 1.000 | 0.000 | 0.000 | 2.38 | 0.149 |
| | Lambeyrarkvísl to lake morphs | AN_Lambeyrarkvísl:Hest | 1.000 | 0.000 | 0.000 | -0.48 | 1.000 |
| | | AN_Lambeyrarkvísl:LB_TH | 1.000 | 0.000 | 0.000 | -0.49 | 1.000 |
| | | AN_Lambeyrarkvísl:LB_UL | 1.000 | 0.000 | 0.000 | -0.50 | 1.000 |
| | | AN_Lambeyrarkvísl:PI_TH | 1.000 | 0.000 | 0.000 | -0.51 | 1.000 |
| | | AN_Lambeyrarkvísl:PI_UL | 1.000 | 0.000 | 0.000 | -0.56 | 1.000 |
| | | AN_Lambeyrarkvísl:PL_TH | 1.000 | 0.000 | 0.000 | -0.47 | 1.000 |
| | | AN_Lambeyrarkvísl:PL_UL | 1.000 | 0.000 | 0.000 | -0.55 | 1.000 |
| | | AN_Lambeyrarkvísl:SB_TH | 1.000 | 0.000 | 0.000 | 2.19 | 0.169 |
| | Víðidalsá to lake morphs | AN_Víðidalsá:Hest | 1.000 | 0.000 | 0.000 | 2.09 | 0.177 |
| | | AN_Víðidalsá:LB_TH | 1.000 | 0.000 | 0.000 | 2.01 | 0.189 |
| | | AN_Víðidalsá:LB_UL | 1.000 | 0.000 | 0.000 | 1.99 | 0.186 |
| | | AN_Víðidalsá:PI_TH | 1.000 | 0.000 | 0.000 | 1.96 | 0.199 |
| | | AN_Víðidalsá:PI_UL | 1.000 | 0.000 | 0.000 | 1.73 | 0.239 |
| | | AN_Víðidalsá:PL_TH | 1.000 | 0.000 | 0.000 | 2.12 | 0.175 |
| | | AN_Víðidalsá:PL_UL | 1.000 | 0.000 | 0.000 | 1.80 | 0.224 |
| | | AN_Víðidalsá:SB_TH | 1.000 | 0.000 | 0.000 | 3.37 | ***0.006** |
| | Vífisdalsá to lake morphs | AN_Vífilsdalsá:Hest | 1.000 | 0.000 | 0.000 | -0.47 | 1.000 |

| | | | | | | | |
|---|---|---|---|---|---|---|---|
| | | AN_Vífilsdalsá:LB_TH | 1.000 | 0.000 | 0.000 | -0.48 | 1.000 |
| | | AN_Vífilsdalsá:LB_UL | 1.000 | 0.000 | 0.000 | -0.48 | 1.000 |
| | | AN_Vífilsdalsá:PI_TH | 1.000 | 0.000 | 0.000 | -0.49 | 1.000 |
| | | AN_Vífilsdalsá:PI_UL | 1.000 | 0.000 | 0.000 | -0.56 | 1.000 |
| | | AN_Vífilsdalsá:PL_TH | 1.000 | 0.000 | 0.000 | -0.46 | 1.000 |
| | | AN_Vífilsdalsá:PL_UL | 1.000 | 0.000 | 0.000 | -0.53 | 1.000 |
| | | AN_Vífilsdalsá:SB_TH | 1.000 | 0.000 | 0.000 | 2.26 | 0.160 |
| | Hestvatn | Hest:LB_TH | 1.000 | 0.000 | 0.000 | -0.38 | 1.000 |
| | | Hest:LB_UL | 1.000 | 0.000 | 0.000 | -0.39 | 1.000 |
| | | Hest:PI_TH | 1.000 | 0.000 | 0.000 | -0.40 | 1.000 |
| | | Hest:PI_UL | 1.000 | 0.000 | 0.000 | -0.46 | 1.000 |
| | | Hest:PL_TH | 1.000 | 0.000 | 0.000 | -0.36 | 1.000 |
| | | Hest:PL_UL | 1.000 | 0.000 | 0.000 | -0.44 | 1.000 |
| | | Hest:SB_TH | 1.000 | 0.000 | 0.000 | 3.00 | 0.098 |
| | Þingvallavatn | LB_TH:PI_TH | 1.000 | 0.000 | 0.000 | -0.42 | 1.000 |
| | | LB_TH:PL_TH | 1.000 | 0.000 | 0.000 | -0.38 | 1.000 |
| | | LB_TH:SB_TH | 1.000 | 0.000 | 0.000 | 2.82 | 0.111 |
| | | PI_TH:PL_TH | 1.000 | 0.000 | 0.000 | -0.39 | 1.000 |
| | | PI_TH:SB_TH | 1.000 | 0.000 | 0.000 | 2.70 | 0.118 |
| | | PL_TH:SB_TH | 1.000 | 0.000 | 0.000 | 3.07 | 0.093 |
| | Úlfljótsvatn | LB_UL:PI_UL | 1.000 | 0.000 | 0.000 | -0.50 | 1.000 |
| | | LB_UL:PL_UL | 1.000 | 0.000 | 0.000 | -0.47 | 1.000 |
| | | PI_UL:PL_UL | 1.000 | 0.000 | 0.000 | -0.53 | 1.000 |
| | Across Þingvallavatn and Úlfljótsvatn | LB_TH:LB_UL | 1.000 | 0.000 | 0.000 | -0.41 | 1.000 |
| | | LB_TH:PI_UL | 1.000 | 0.000 | 0.000 | -0.48 | 1.000 |
| | | LB_TH:PL_UL | 1.000 | 0.000 | 0.000 | -0.46 | 1.000 |
| | | LB_UL:PI_TH | 1.000 | 0.000 | 0.000 | -0.42 | 1.000 |
| | | LB_UL:PL_TH | 1.000 | 0.000 | 0.000 | -0.38 | 1.000 |
| | | LB_UL:SB_TH | 1.000 | 0.000 | 0.000 | 2.76 | 0.114 |
| | | PI_TH:PI_UL | 1.000 | 0.000 | 0.000 | -0.49 | 1.000 |
| | | PI_TH:PL_UL | 1.000 | 0.000 | 0.000 | -0.46 | 1.000 |
| | | PI_UL:PL_TH | 1.000 | 0.000 | 0.000 | -0.46 | 1.000 |
| | | PI_UL:SB_TH | 1.000 | 0.000 | 0.000 | 2.25 | 0.162 |
| | | PL_TH:PL_UL | 1.000 | 0.000 | 0.000 | -0.44 | 1.000 |
| | | PL_UL:SB_TH | 1.000 | 0.000 | 0.000 | 2.39 | 0.143 |
| Maxilla | AN populations | AN_Brúará:AN_Gufudalsá | 1.000 | 0.000 | 0.000 | -0.07 | 1.000 |
| | | AN_Brúará:AN_Lambeyrarkvísl | 1.000 | 0.000 | 0.000 | -0.07 | 1.000 |
| | | AN_Brúará:AN_Víðidalsá | 1.000 | 0.000 | 0.000 | -0.05 | 0.245 |
| | | AN_Brúará:AN_Vífilsdalsá | 1.000 | 0.000 | 0.000 | -0.07 | 1.000 |
| | | AN_Gufudalsá:AN_Lambeyrarkvísl | 1.000 | 0.000 | 0.000 | -0.59 | 1.000 |
| | | AN_Gufudalsá:AN_Víðidalsá | 1.000 | 0.000 | 0.000 | 1.73 | 0.232 |
| | | AN_Gufudalsá:AN_Vífilsdalsá | 1.000 | 0.000 | 0.000 | -0.58 | 1.000 |
| | | AN_Lambeyrarkvísl:AN_Víðidalsá | 1.000 | 0.000 | 0.000 | 1.67 | 0.247 |
| | | AN_Lambeyrarkvísl:AN_Vífilsdalsá | 1.000 | 0.000 | 0.000 | -0.60 | 1.000 |
| | | AN_Víðidalsá:AN_Vífilsdalsá | 1.000 | 0.000 | 0.000 | 1.68 | 0.236 |
| | Brúará to lake morphs | AN_Brúará:Hest | 1.000 | 0.000 | 0.000 | -0.07 | 1.000 |
| | | AN_Brúará:LB_TH | 1.000 | 0.000 | 0.000 | -0.07 | 1.000 |
| | | AN_Brúará:LB_UL | 1.000 | 0.000 | 0.000 | -0.05 | 0.239 |
| | | AN_Brúará:PI_TH | 1.000 | 0.000 | 0.000 | -0.07 | 1.000 |
| | | AN_Brúará:PI_UL | 1.000 | 0.000 | 0.000 | -0.07 | 1.000 |
| | | AN_Brúará:PL_TH | 1.000 | 0.000 | 0.000 | -0.05 | 0.234 |
| | | AN_Brúará:PL_UL | 1.000 | 0.000 | 0.000 | -0.07 | 1.000 |
| | | AN_Brúará:SB_TH | 1.000 | 0.000 | 0.000 | -0.07 | 1.000 |
| | Gufudalsá to lake morphs | AN_Gufudalsá:Hest | 1.000 | 0.000 | 0.000 | -0.53 | 1.000 |
| | | AN_Gufudalsá:LB_TH | 1.000 | 0.000 | 0.000 | -0.54 | 1.000 |
| | | AN_Gufudalsá:LB_UL | 1.000 | 0.000 | 0.000 | 1.81 | 0.220 |
| | | AN_Gufudalsá:PI_TH | 1.000 | 0.000 | 0.000 | -0.52 | 1.000 |
| | | AN_Gufudalsá:PI_UL | 1.000 | 0.000 | 0.000 | -0.56 | 1.000 |
| | | AN_Gufudalsá:PL_TH | 1.000 | 0.000 | 0.000 | 1.84 | 0.213 |
| | | AN_Gufudalsá:PL_UL | 1.000 | 0.000 | 0.000 | -0.56 | 1.000 |
| | | AN_Gufudalsá:SB_TH | 1.000 | 0.000 | 0.000 | -0.48 | 1.000 |

| | | | | | | | |
|---|---|---|---|---|---|---|---|
| | Lambeyrarkvísl to lake morphs | AN_Lambeyrarkvísl:Hest | 1.000 | 0.000 | 0.000 | -0.53 | 1.000 |
| | | AN_Lambeyrarkvísl:LB_TH | 1.000 | 0.000 | 0.000 | -0.56 | 1.000 |
| | | AN_Lambeyrarkvísl:LB_UL | 1.000 | 0.000 | 0.000 | 1.75 | 0.231 |
| | | AN_Lambeyrarkvísl:PI_TH | 1.000 | 0.000 | 0.000 | -0.55 | 1.000 |
| | | AN_Lambeyrarkvísl:PI_UL | 1.000 | 0.000 | 0.000 | -0.59 | 1.000 |
| | | AN_Lambeyrarkvísl:PL_TH | 1.000 | 0.000 | 0.000 | 1.78 | 0.225 |
| | | AN_Lambeyrarkvísl:PL_UL | 1.000 | 0.000 | 0.000 | -0.59 | 1.000 |
| | | AN_Lambeyrarkvísl:SB_TH | 1.000 | 0.000 | 0.000 | -0.50 | 1.000 |
| | Víðidalsá to lake morphs | AN_Víðidalsá:Hest | 1.000 | 0.000 | 0.000 | 1.88 | 0.208 |
| | | AN_Víðidalsá:LB_TH | 1.000 | 0.000 | 0.000 | 1.86 | 0.211 |
| | | AN_Víðidalsá:LB_UL | 1.000 | 0.000 | 0.000 | 2.86 | 0.011 |
| | | AN_Víðidalsá:PI_TH | 1.000 | 0.000 | 0.000 | 1.88 | 0.203 |
| | | AN_Víðidalsá:PI_UL | 1.000 | 0.000 | 0.000 | 1.74 | 0.232 |
| | | AN_Víðidalsá:PL_TH | 1.000 | 0.000 | 0.000 | 2.90 | 0.014 |
| | | AN_Víðidalsá:PL_UL | 1.000 | 0.000 | 0.000 | 1.73 | 0.237 |
| | | AN_Víðidalsá:SB_TH | 1.000 | 0.000 | 0.000 | 2.07 | 0.178 |
| | Vífisdalsá to lake morphs | AN_Vífilsdalsá:Hest | 1.000 | 0.000 | 0.000 | -0.55 | 1.000 |
| | | AN_Vífilsdalsá:LB_TH | 1.000 | 0.000 | 0.000 | -0.55 | 1.000 |
| | | AN_Vífilsdalsá:LB_UL | 1.000 | 0.000 | 0.000 | 1.75 | 0.226 |
| | | AN_Vífilsdalsá:PI_TH | 1.000 | 0.000 | 0.000 | -0.53 | 1.000 |
| | | AN_Vífilsdalsá:PI_UL | 1.000 | 0.000 | 0.000 | -0.57 | 1.000 |
| | | AN_Vífilsdalsá:PL_TH | 1.000 | 0.000 | 0.000 | 1.78 | 0.229 |
| | | AN_Vífilsdalsá:PL_UL | 1.000 | 0.000 | 0.000 | -0.58 | 1.000 |
| | | AN_Vífilsdalsá:SB_TH | 1.000 | 0.000 | 0.000 | -0.51 | 1.000 |
| | Hestvatn | Hest:LB_TH | 1.000 | 0.000 | 0.000 | -0.49 | 1.000 |
| | | Hest:LB_UL | 1.000 | 0.000 | 0.000 | 1.98 | 0.192 |
| | | Hest:PI_TH | 1.000 | 0.000 | 0.000 | -0.48 | 1.000 |
| | | Hest:PI_UL | 1.000 | 0.000 | 0.000 | -0.52 | 1.000 |
| | | Hest:PL_TH | 1.000 | 0.000 | 0.000 | 2.01 | 0.187 |
| | | Hest:PL_UL | 1.000 | 0.000 | 0.000 | -0.52 | 1.000 |
| | | Hest:SB_TH | 1.000 | 0.000 | 0.000 | -0.45 | 1.000 |
| | Þingvallavatn | LB_TH:PI_TH | 1.000 | 0.000 | 0.000 | -0.48 | 1.000 |
| | | LB_TH:PL_TH | 1.000 | 0.000 | 0.000 | 1.99 | 0.196 |
| | | LB_TH:SB_TH | 1.000 | 0.000 | 0.000 | -0.45 | 1.000 |
| | | PI_TH:PL_TH | 1.000 | 0.000 | 0.000 | 2.01 | 0.187 |
| | | PI_TH:SB_TH | 1.000 | 0.000 | 0.000 | -0.44 | 1.000 |
| | | PL_TH:SB_TH | 1.000 | 0.000 | 0.000 | 2.23 | 0.161 |
| | Úlfljótsvatn | LB_UL:PI_UL | 1.000 | 0.000 | 0.000 | 1.82 | 0.212 |
| | | LB_UL:PL_UL | 1.000 | 0.000 | 0.000 | 1.81 | 0.223 |
| | | PI_UL:PL_UL | 1.000 | 0.000 | 0.000 | -0.57 | 1.000 |
| | Across Þingvallavatn and Úlfljótsvatn | LB_TH:LB_UL | 1.000 | 0.000 | 0.000 | 1.95 | 0.196 |
| | | LB_TH:PI_UL | 1.000 | 0.000 | 0.000 | -0.53 | 1.000 |
| | | LB_TH:PL_UL | 1.000 | 0.000 | 0.000 | -0.52 | 1.000 |
| | | LB_UL:PI_TH | 1.000 | 0.000 | 0.000 | 1.97 | 0.196 |
| | | LB_UL:PL_TH | 1.000 | 0.000 | 0.000 | 3.03 | 0.014 |
| | | LB_UL:SB_TH | 1.000 | 0.000 | 0.000 | 2.18 | 0.165 |
| | | PI_TH:PI_UL | 1.000 | 0.000 | 0.000 | -0.52 | 1.000 |
| | | PI_TH:PL_UL | 1.000 | 0.000 | 0.000 | -0.52 | 1.000 |
| | | PI_UL:PL_TH | 1.000 | 0.000 | 0.000 | 1.85 | 0.214 |
| | | PI_UL:SB_TH | 1.000 | 0.000 | 0.000 | -0.49 | 1.000 |
| | | PL_TH:PL_UL | 1.000 | 0.000 | 0.000 | 1.84 | 0.218 |
| | | PL_UL:SB_TH | 1.000 | 0.000 | 0.000 | -0.48 | 1.000 |
| Supramaxilla | AN populations | AN_Brúará:AN_Gufudalsá | 1.000 | 0.000 | 0.000 | -0.05 | 1.000 |
| | | AN_Brúará:AN_Lambeyrarkvísl | 1.000 | 0.000 | 0.000 | -0.06 | 1.000 |
| | | AN_Brúará:AN_Víðidalsá | 1.000 | 0.000 | 0.000 | -0.03 | 0.232 |
| | | AN_Brúará:AN_Vífilsdalsá | 1.000 | 0.000 | 0.000 | -0.05 | 1.000 |
| | | AN_Gufudalsá:AN_Lambeyrarkvísl | 1.000 | 0.000 | 0.000 | -0.04 | 1.000 |
| | | AN_Gufudalsá:AN_Víðidalsá | 1.000 | 0.000 | 0.000 | 1.67 | 0.245 |
| | | AN_Gufudalsá:AN_Vífilsdalsá | 1.000 | 0.000 | 0.000 | -0.54 | 1.000 |
| | | AN_Lambeyrarkvísl:AN_Víðidalsá | 1.000 | 0.000 | 0.000 | -0.01 | 0.234 |
| | | AN_Lambeyrarkvísl:AN_Vífilsdalsá | 1.000 | 0.000 | 0.000 | -0.04 | 1.000 |

| | | | | | | | |
|---|---|---|---|---|---|---|---|
| | | AN_Víðidalsá:AN_Vífilsdalsá | 1.000 | 0.000 | 0.000 | 1.75 | 0.232 |
| | Brúará to lake morphs | AN_Brúará:Hest | 1.000 | 0.000 | 0.000 | -0.05 | 1.000 |
| | | AN_Brúará:LB_TH | 1.000 | 0.000 | 0.000 | -0.05 | 1.000 |
| | | AN_Brúará:LB_UL | 1.000 | 0.000 | 0.000 | -0.05 | 1.000 |
| | | AN_Brúará:PI_TH | 1.000 | 0.000 | 0.000 | -0.05 | 1.000 |
| | | AN_Brúará:PI_UL | 1.000 | 0.000 | 0.000 | -0.05 | 1.000 |
| | | AN_Brúará:PL_TH | 1.000 | 0.000 | 0.000 | -0.05 | 1.000 |
| | | AN_Brúará:PL_UL | 1.000 | 0.000 | 0.000 | -0.05 | 1.000 |
| | | AN_Brúará:SB_TH | 1.000 | 0.000 | 0.000 | -0.05 | 1.000 |
| | Gufudalsá to lake morphs | AN_Gufudalsá:Hest | 1.000 | 0.000 | 0.000 | -0.54 | 1.000 |
| | | AN_Gufudalsá:LB_TH | 1.000 | 0.000 | 0.000 | -0.54 | 1.000 |
| | | AN_Gufudalsá:LB_UL | 1.000 | 0.000 | 0.000 | -0.54 | 1.000 |
| | | AN_Gufudalsá:PI_TH | 1.000 | 0.000 | 0.000 | -0.54 | 1.000 |
| | | AN_Gufudalsá:PI_UL | 1.000 | 0.000 | 0.000 | -0.56 | 1.000 |
| | | AN_Gufudalsá:PL_TH | 1.000 | 0.000 | 0.000 | -0.55 | 1.000 |
| | | AN_Gufudalsá:PL_UL | 1.000 | 0.000 | 0.000 | -0.57 | 1.000 |
| | | AN_Gufudalsá:SB_TH | 1.000 | 0.000 | 0.000 | -0.53 | 1.000 |
| | Lambeyrarkvísl to lake morphs | AN_Lambeyrarkvísl:Hest | 1.000 | 0.000 | 0.000 | -0.04 | 1.000 |
| | | AN_Lambeyrarkvísl:LB_TH | 1.000 | 0.000 | 0.000 | -0.04 | 1.000 |
| | | AN_Lambeyrarkvísl:LB_UL | 1.000 | 0.000 | 0.000 | -0.04 | 1.000 |
| | | AN_Lambeyrarkvísl:PI_TH | 1.000 | 0.000 | 0.000 | -0.04 | 1.000 |
| | | AN_Lambeyrarkvísl:PI_UL | 1.000 | 0.000 | 0.000 | -0.04 | 1.000 |
| | | AN_Lambeyrarkvísl:PL_TH | 1.000 | 0.000 | 0.000 | -0.04 | 1.000 |
| | | AN_Lambeyrarkvísl:PL_UL | 1.000 | 0.000 | 0.000 | -0.04 | 1.000 |
| | | AN_Lambeyrarkvísl:SB_TH | 1.000 | 0.000 | 0.000 | -0.04 | 1.000 |
| | Víðidalsá to lake morphs | AN_Víðidalsá:Hest | 1.000 | 0.000 | 0.000 | 1.73 | 0.235 |
| | | AN_Víðidalsá:LB_TH | 1.000 | 0.000 | 0.000 | 1.73 | 0.234 |
| | | AN_Víðidalsá:LB_UL | 1.000 | 0.000 | 0.000 | 1.76 | 0.226 |
| | | AN_Víðidalsá:PI_TH | 1.000 | 0.000 | 0.000 | 1.72 | 0.236 |
| | | AN_Víðidalsá:PI_UL | 1.000 | 0.000 | 0.000 | 1.64 | 0.258 |
| | | AN_Víðidalsá:PL_TH | 1.000 | 0.000 | 0.000 | 1.71 | 0.236 |
| | | AN_Víðidalsá:PL_UL | 1.000 | 0.000 | 0.000 | 1.66 | 0.254 |
| | | AN_Víðidalsá:SB_TH | 1.000 | 0.000 | 0.000 | 1.84 | 0.213 |
| | Vífisdalsá to lake morphs | AN_Vífilsdalsá:Hest | 1.000 | 0.000 | 0.000 | -0.53 | 1.000 |
| | | AN_Vífilsdalsá:LB_TH | 1.000 | 0.000 | 0.000 | -0.53 | 1.000 |
| | | AN_Vífilsdalsá:LB_UL | 1.000 | 0.000 | 0.000 | -0.52 | 1.000 |
| | | AN_Vífilsdalsá:PI_TH | 1.000 | 0.000 | 0.000 | -0.52 | 1.000 |
| | | AN_Vífilsdalsá:PI_UL | 1.000 | 0.000 | 0.000 | -0.55 | 1.000 |
| | | AN_Vífilsdalsá:PL_TH | 1.000 | 0.000 | 0.000 | -0.53 | 1.000 |
| | | AN_Vífilsdalsá:PL_UL | 1.000 | 0.000 | 0.000 | -0.54 | 1.000 |
| | | AN_Vífilsdalsá:SB_TH | 1.000 | 0.000 | 0.000 | -0.50 | 1.000 |
| | Hestvatn | Hest:LB_TH | 1.000 | 0.000 | 0.000 | -0.52 | 1.000 |
| | | Hest:LB_UL | 1.000 | 0.000 | 0.000 | -0.51 | 1.000 |
| | | Hest:PI_TH | 1.000 | 0.000 | 0.000 | -0.53 | 1.000 |
| | | Hest:PI_UL | 1.000 | 0.000 | 0.000 | -0.55 | 1.000 |
| | | Hest:PL_TH | 1.000 | 0.000 | 0.000 | -0.53 | 1.000 |
| | | Hest:PL_UL | 1.000 | 0.000 | 0.000 | -0.55 | 1.000 |
| | | Hest:SB_TH | 1.000 | 0.000 | 0.000 | -0.50 | 1.000 |
| | Þingvallavatn | LB_TH:PI_TH | 1.000 | 0.000 | 0.000 | -0.53 | 1.000 |
| | | LB_TH:PL_TH | 1.000 | 0.000 | 0.000 | -0.54 | 1.000 |
| | | LB_TH:SB_TH | 1.000 | 0.000 | 0.000 | -0.50 | 1.000 |
| | | PI_TH:PL_TH | 1.000 | 0.000 | 0.000 | -0.54 | 1.000 |
| | | PI_TH:SB_TH | 1.000 | 0.000 | 0.000 | -0.50 | 1.000 |
| | | PL_TH:SB_TH | 1.000 | 0.000 | 0.000 | -0.49 | 1.000 |
| | Úlfljótsvatn | LB_UL:PI_UL | 1.000 | 0.000 | 0.000 | -0.55 | 1.000 |
| | | LB_UL:PL_UL | 1.000 | 0.000 | 0.000 | -0.56 | 1.000 |
| | | PI_UL:PL_UL | 1.000 | 0.000 | 0.000 | -0.58 | 1.000 |
| | Across Þingvallavatn and Úlfljótsvatn | LB_TH:LB_UL | 1.000 | 0.000 | 0.000 | -0.52 | 1.000 |
| | | LB_TH:PI_UL | 1.000 | 0.000 | 0.000 | -0.56 | 1.000 |
| | | LB_TH:PL_UL | 1.000 | 0.000 | 0.000 | -0.55 | 1.000 |
| | | LB_UL:PI_TH | 1.000 | 0.000 | 0.000 | -0.52 | 1.000 |

| | | | | | | | |
|---|---|---|---|---|---|---|---|
| | | LB_UL:PL_TH | 1.000 | 0.000 | 0.000 | -0.53 | 1.000 |
| | | LB_UL:SB_TH | 1.000 | 0.000 | 0.000 | -0.49 | 1.000 |
| | | PI_TH:PI_UL | 1.000 | 0.000 | 0.000 | -0.57 | 1.000 |
| | | PI_TH:PL_UL | 1.000 | 0.000 | 0.000 | -0.56 | 1.000 |
| | | PI_UL:PL_TH | 1.000 | 0.000 | 0.000 | -0.56 | 1.000 |
| | | PI_UL:SB_TH | 1.000 | 0.000 | 0.000 | -0.53 | 1.000 |
| | | PL_TH:PL_UL | 1.000 | 0.000 | 0.000 | -0.56 | 1.000 |
| | | PL_UL:SB_TH | 1.000 | 0.000 | 0.000 | -0.53 | 1.000 |

R = slope of vector correlation; UCL = upper confidence level, Z = Z-statistic; P = P-value, ≤ 0.008 in bold with asterisk.

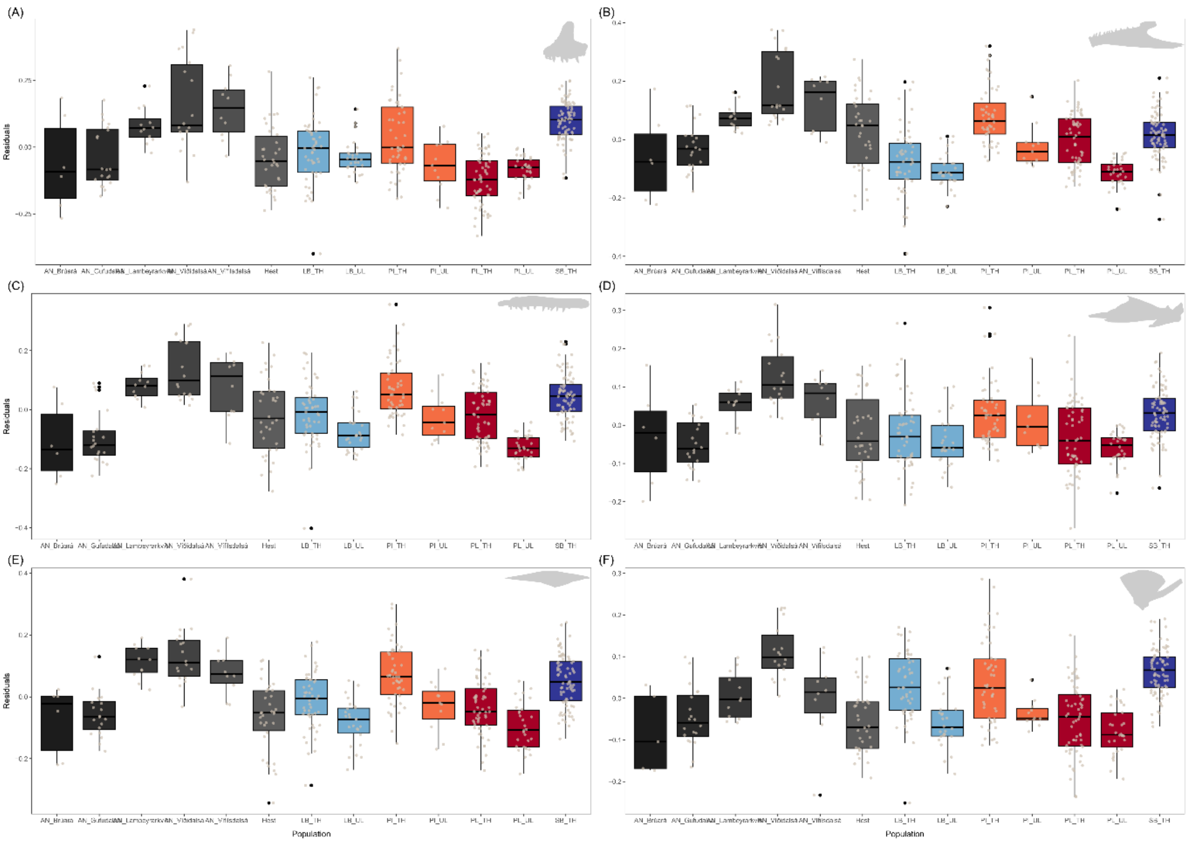


***Supplementary material 11****: Boxplots showing the differences of residuals between the populations, when the AN-charr populations are treated as separate populations. Upper jaw bones on left side, from top A) premaxilla, C) maxilla and E) supramaxilla. Lower jaw bones on right side, from top: B) dentary, D) articular-angular and F) quadrate. Upper right side of each graph shows illustration of the bone.*

***Supplementary material 12****: Mean log CS ($CS_{bone}$) size for each group and mean residuals with standard deviation. Note Hítarvatn individuals included in this table however excluded from any analysis of mean bone size.*

| Bones | Population | Mean Log Bone size | Mean Residuals |
|---|---|---|---|
| Dentary | AN_Brúará | 1.60 ± 0.16 | -0.06 ± 0.15 |
| | AN_Gufudalsá | 1.34 ± 0.18 | -0.03 ± 0.08 |
| | AN_Lambeyrarkvísl | 1.56 ± 0.13 | 0.08 ± 0.04 |
| | AN_Víðidalsá | 1.87 ± 0.22 | 0.19 ± 0.12 |
| | AN_Vífilsdalsá | 1.63 ± 0.16 | 0.12 ± 0.09 |
| | AN | 1.59 ± 0.27 | 0.07 ± 0.13 |

| | | | |
|---|---|---|---|
| | Hestvatn | 1.43 ± 0.26 | 0.02 ± 0.13 |
| | Hítarvatn | 1.29 ± 0.18 | -0.07 ± 0.10 |
| | LB_TH | 1.56 ± 0.23 | -0.08 ± 0.12 |
| | LB_UL | 1.41 ± 0.24 | -0.10 ± 0.06 |
| | PI_TH | 1.62 ± 0.27 | 0.08 ± 0.10 |
| | PI_UL | 1.68 ± 0.19 | -0.02 ± 0.07 |
| | PL_TH | 0.78 ± 0.21 | 0.00 ± 0.09 |
| | PL_UL | 0.80 ± 0.17 | -0.11 ± 0.04 |
| | SB_TH | 0.57 ± 0.22 | 0.01 ± 0.08 |
| Articular - Angular | AN_Brúará | 1.35 ± 0.15 | -0.03 ± 0.13 |
| | AN_Gufudalsá | 1.05 ± 0.17 | -0.05 ± 0.06 |
| | AN_Lambeyrarkvísl | 1.27 ± 0.11 | 0.05 ± 0.05 |
| | AN_Víðidalsá | 1.52 ± 0.19 | 0.13 ± 0.08 |
| | AN_Vífilsdalsá | 1.29 ± 0.14 | 0.06 ± 0.07 |
| | AN | 1.28 ± 0.24 | 0.04 ± 0.10 |
| | Hestvatn | 1.12 ± 0.26 | -0.02 ± 0.10 |
| | Hítarvatn | 1.02 ± 0.16 | -0.07 ± 0.09 |
| | LB_TH | 1.31 ± 0.21 | -0.04 ± 0.09 |
| | LB_UL | 1.20 ± 0.23 | -0.04 ± 0.07 |
| | PI_TH | 1.31 ± 0.25 | 0.05 ± 0.10 |
| | PI_UL | 1.43 ± 0.18 | 0.01 ± 0.08 |
| | PL_TH | 0.53 ± 0.20 | -0.03 ± 0.08 |
| | PL_UL | 0.61 ± 0.16 | -0.06 ± 0.04 |
| | SB_TH | 0.37 ± 0.22 | 0.03 ± 0.07 |
| Quadrate | AN_Brúará | 0.41 ± 0.08 | -0.08 ± 0.10 |
| | AN_Gufudalsá | 0.20 ± 0.16 | -0.05 ± 0.07 |
| | AN_Lambeyrarkvísl | 0.37 ± 0.13 | 0.01 ± 0.06 |
| | AN_Víðidalsá | 0.65 ± 0.16 | 0.11 ± 0.07 |
| | AN_Vífilsdalsá | 0.38 ± 0.19 | 0.00 ± 0.10 |
| | AN | 0.39 ± 0.23 | 0.01 ± 0.10 |
| | Hestvatn | 0.23 ± 0.25 | -0.06 ± 0.08 |
| | Hítarvatn | 0.18 ± 0.13 | -0.06 ± 0.06 |
| | LB_TH | 0.52 ± 0.19 | 0.03 ± 0.08 |
| | LB_UL | 0.33 ± 0.24 | -0.06 ± 0.06 |
| | PI_TH | 0.45 ± 0.27 | 0.04 ± 0.10 |
| | PI_UL | 0.53 ± 0.13 | -0.04 ± 0.03 |
| | PL_TH | -0.33 ± 0.21 | -0.05 ± 0.09 |
| | PL_UL | -0.25 ± 0.16 | -0.08 ± 0.06 |
| | SB_TH | -0.43 ± 0.24 | 0.06 ± 0.06 |
| Premaxilla | AN_Brúará | 0.11 ± 0.17 | -0.06 ± 0.18 |
| | AN_Gufudalsá | -0.17 ± 0.21 | -0.04 ± 0.11 |
| | AN_Lambeyrarkvísl | 0.08 ± 0.15 | 0.08 ± 0.07 |
| | AN_Víðidalsá | 0.35 ± 0.28 | 0.16 ± 0.17 |
| | AN_Vífilsdalsá | 0.16 ± 0.19 | 0.14 ± 0.11 |
| | AN | 0.09 ± 0.29 | 0.06 ± 0.16 |
| | Hestvatn | -0.13 ± 0.30 | -0.04 ± 0.12 |
| | Hítarvatn | -0.24 ± 0.16 | -0.11 ± 0.08 |
| | LB_TH | 0.13 ± 0.23 | -0.01 ± 0.12 |
| | LB_UL | -0.01 ± 0.24 | -0.04 ± 0.06 |
| | PI_TH | 0.09 ± 0.31 | 0.04 ± 0.14 |
| | PI_UL | 0.15 ± 0.21 | -0.07 ± 0.10 |
| | PL_TH | -0.84 ± 0.23 | -0.12 ± 0.10 |
| | PL_UL | -0.67 ± 0.18 | -0.08 ± 0.05 |
| | SB_TH | -0.85 ± 0.24 | 0.10 ± 0.07 |
| Maxilla | AN_Brúará | 1.54 ± 0.12 | -0.11 ± 0.13 |

| | | | |
|---|---|---|---|
| | AN_Gufudalsá | 1.26 ± 0.19 | -0.10 ± 0.09 |
| | AN_Lambeyrarkvísl | 1.56 ± 0.13 | 0.08 ± 0.04 |
| | AN_Víðidalsá | 1.80 ± 0.20 | 0.14 ± 0.10 |
| | AN_Vífilsdalsá | 1.57 ± 0.18 | 0.08 ± 0.11 |
| | AN | 1.54 ± 0.27 | 0.03 ± 0.14 |
| | Hestvatn | 1.38 ± 0.28 | -0.02 ± 0.13 |
| | Hítarvatn | 1.26 ± 0.19 | -0.10 ± 0.10 |
| | LB_TH | 1.61 ± 0.22 | -0.01 ± 0.11 |
| | LB_UL | 1.43 ± 0.25 | -0.08 ± 0.06 |
| | PI_TH | 1.60 ± 0.26 | 0.07 ± 0.10 |
| | PI_UL | 1.67 ± 0.19 | -0.03 ± 0.08 |
| | PL_TH | 0.77 ± 0.20 | -0.02 ± 0.09 |
| | PL_UL | 0.79 ± 0.17 | -0.13 ± 0.04 |
| | SB_TH | 0.61 ± 0.24 | 0.05 ± 0.08 |
| Supramaxilla | AN_Brúará | 0.32 ± 0.15 | -0.08 ± 0.11 |
| | AN_Gufudalsá | 0.07 ± 0.17 | -0.06 ± 0.07 |
| | AN_Lambeyrarkvísl | 0.35 ± 0.14 | 0.11 ± 0.05 |
| | AN_Víðidalsá | 0.55 ± 0.20 | 0.13 ± 0.09 |
| | AN_Vífilsdalsá | 0.33 ± 0.15 | 0.08 ± 0.07 |
| | AN | 0.31 ± 0.25 | 0.04 ± 0.12 |
| | Hestvatn | 0.10 ± 0.26 | -0.06 ± 0.11 |
| | Hítarvatn | 0.06 ± 0.14 | -0.06 ± 0.07 |
| | LB_TH | 0.36 ± 0.21 | -0.01 ± 0.09 |
| | LB_UL | 0.18 ± 0.24 | -0.08 ± 0.07 |
| | PI_TH | 0.36 ± 0.24 | 0.07 ± 0.10 |
| | PI_UL | 0.41 ± 0.18 | -0.03 ± 0.08 |
| | PL_TH | -0.46 ± 0.22 | -0.03 ± 0.09 |
| | PL_UL | -0.40 ± 0.18 | -0.10 ± 0.08 |
| | SB_TH | -0.58 ± 0.24 | 0.05 ± 0.08 |

***Supplementary material 13****: Anova results when testing whether residuals of bone size differ between populations. Hítarvatn fish excluded from this analysis. AN-charr treated as five separate populations.*

| Bone | Df | Sum Sq | Mean Sq | F-value | P-value |
|---|---|---|---|---|---|
| Dentary | 12 | 2.206 | 0.184 | 21.27 | ***< 0.001** |
| Maxilla | 12 | 1.855 | 0.155 | 18.88 | ***< 0.001** |
| Articular - Angular | 12 | 0.901 | 0.075 | 12.06 | ***< 0.001** |
| Quadrate | 12 | 1.293 | 0.108 | 18.95 | ***< 0.001** |
| Premaxilla | 12 | 2.934 | 0.244 | 22.80 | ***< 0.001** |
| Supramaxilla | 12 | 1.743 | 0.145 | 18.54 | ***< 0.001** |

Df = degrees of freedom, Sum sq= Sum of squares, Mean sq= Mean square F = F-statistic; P = P-value, ≤ 0.008 in bold with asterisk.

***Supplementary material 14****: Results from post hoc Tukey test of variation in residuals (representing bone size) between populations for the six bones (dentary, articular-angular, quadrate, premaxilla, maxilla and supramaxilla). AN-charr treated as five separate populations.*

| Bone | System | Pairs | Diff | LCL | UCL | P-value |
|---|---|---|---|---|---|---|
| Dentary | AN populations | AN_Gufudalsá-AN_Brúará | 0.032 | -0.112 | 0.176 | 1.000 |
| | | AN_Lambeyrarkvísl-AN_Brúará | 0.137 | -0.018 | 0.292 | 0.149 |
| | | AN_Víðidalsá-AN_Brúará | 0.247 | 0.101 | 0.393 | ***<0.001** |
| | | AN_Vífilsdalsá-AN_Brúará | 0.181 | 0.021 | 0.341 | 0.012 |
| | | AN_Lambeyrarkvísl-AN_Gufudalsá | 0.104 | -0.009 | 0.218 | 0.105 |
| | | AN_Víðidalsá-AN_Gufudalsá | 0.214 | 0.114 | 0.315 | ***<0.001** |
| | | AN_Vífilsdalsá-AN_Gufudalsá | 0.149 | 0.029 | 0.269 | ***0.003** |
| | | AN_Víðidalsá-AN_Lambeyrarkvísl | 0.110 | -0.005 | 0.226 | 0.079 |
| | | AN_Vífilsdalsá-AN_Lambeyrarkvísl | 0.044 | -0.088 | 0.177 | 0.996 |
| | | AN_Vífilsdalsá-AN_Víðidalsá | -0.066 | -0.188 | 0.056 | 0.851 |
| | Brúará to lake morphs | Hest-AN_Brúará | 0.084 | -0.053 | 0.221 | 0.703 |
| | | LB_TH-AN_Brúará | -0.019 | -0.153 | 0.115 | 1.000 |
| | | LB_UL-AN_Brúará | -0.045 | -0.185 | 0.096 | 0.998 |
| | | PI_TH-AN_Brúará | 0.141 | 0.006 | 0.275 | 0.031 |
| | | PI_UL-AN_Brúará | 0.036 | -0.121 | 0.193 | 1.000 |
| | | PL_TH-AN_Brúará | 0.057 | -0.076 | 0.190 | 0.969 |
| | | PL_UL-AN_Brúará | -0.054 | -0.192 | 0.085 | 0.987 |
| | | SB_TH-AN_Brúará | 0.075 | -0.056 | 0.206 | 0.792 |
| | Gufudalsá to lake morphs | Hest-AN_Gufudalsá | 0.052 | -0.035 | 0.139 | 0.743 |
| | | LB_TH-AN_Gufudalsá | -0.051 | -0.133 | 0.031 | 0.694 |
| | | LB_UL-AN_Gufudalsá | -0.077 | -0.169 | 0.015 | 0.215 |
| | | PI_TH-AN_Gufudalsá | 0.108 | 0.026 | 0.191 | ***0.001** |
| | | PI_UL-AN_Gufudalsá | 0.004 | -0.113 | 0.120 | 1.000 |
| | | PL_TH-AN_Gufudalsá | 0.025 | -0.055 | 0.105 | 0.998 |
| | | PL_UL-AN_Gufudalsá | -0.086 | -0.175 | 0.004 | 0.074 |
| | | SB_TH-AN_Gufudalsá | 0.043 | -0.035 | 0.120 | 0.828 |
| | Lambeyrarkvísl to lake morphs | Hest-AN_Lambeyrarkvísl | -0.052 | -0.156 | 0.052 | 0.902 |
| | | LB_TH-AN_Lambeyrarkvísl | -0.155 | -0.255 | -0.055 | ***<0.001** |
| | | LB_UL-AN_Lambeyrarkvísl | -0.181 | -0.289 | -0.073 | ***<0.001** |
| | | PI_TH-AN_Lambeyrarkvísl | 0.004 | -0.096 | 0.104 | 1.000 |
| | | PI_UL-AN_Lambeyrarkvísl | -0.101 | -0.230 | 0.029 | 0.319 |
| | | PL_TH-AN_Lambeyrarkvísl | -0.080 | -0.178 | 0.018 | 0.253 |
| | | PL_UL-AN_Lambeyrarkvísl | -0.190 | -0.296 | -0.084 | ***<0.001** |
| | | SB_TH-AN_Lambeyrarkvísl | -0.062 | -0.158 | 0.034 | 0.629 |
| | Víðidalsá to lake morphs | Hest-AN_Víðidalsá | -0.163 | -0.253 | -0.072 | ***<0.001** |
| | | LB_TH-AN_Víðidalsá | -0.265 | -0.351 | -0.180 | ***<0.001** |
| | | LB_UL-AN_Víðidalsá | -0.291 | -0.386 | -0.196 | ***<0.001** |
| | | PI_TH-AN_Víðidalsá | -0.106 | -0.192 | -0.020 | ***0.003** |
| | | PI_UL-AN_Víðidalsá | -0.211 | -0.329 | -0.092 | ***<0.001** |
| | | PL_TH-AN_Víðidalsá | -0.190 | -0.273 | -0.106 | ***<0.001** |
| | | PL_UL-AN_Víðidalsá | -0.300 | -0.393 | -0.208 | ***<0.001** |
| | | SB_TH-AN_Víðidalsá | -0.172 | -0.253 | -0.091 | ***<0.001** |
| | Vífisdalsá to lake morphs | Hest-AN_Vífilsdalsá | -0.097 | -0.208 | 0.015 | 0.167 |
| | | LB_TH-AN_Vífilsdalsá | -0.199 | -0.307 | -0.092 | ***<0.001** |
| | | LB_UL-AN_Vífilsdalsá | -0.226 | -0.341 | -0.110 | ***<0.001** |
| | | PI_TH-AN_Vífilsdalsá | -0.040 | -0.148 | 0.068 | 0.990 |
| | | PI_UL-AN_Vífilsdalsá | -0.145 | -0.280 | -0.009 | 0.024 |
| | | PL_TH-AN_Vífilsdalsá | -0.124 | -0.230 | -0.018 | ***0.007** |
| | | PL_UL-AN_Vífilsdalsá | -0.235 | -0.348 | -0.121 | ***<0.001** |
| | | SB_TH-AN_Vífilsdalsá | -0.106 | -0.210 | -0.002 | 0.040 |
| | Hestvatn | LB_TH-Hest | -0.103 | -0.172 | -0.034 | ***<0.001** |
| | | LB_UL-Hest | -0.129 | -0.210 | -0.048 | ***<0.001** |
| | | PI_TH-Hest | 0.056 | -0.013 | 0.126 | 0.259 |
| | | PI_UL-Hest | -0.048 | -0.156 | 0.059 | 0.957 |

| | | | | | | |
|---|---|---|---|---|---|---|
| | | PL_TH-Hest | -0.027 | -0.094 | 0.039 | 0.979 |
| | | PL_UL-Hest | -0.138 | -0.215 | -0.060 | ***<0.001** |
| | | SB_TH-Hest | -0.009 | -0.073 | 0.054 | 1.000 |
| | Þingvallavatn | PI_TH-LB_TH | 0.159 | 0.096 | 0.222 | ***<0.001** |
| | | PL_TH-LB_TH | 0.076 | 0.016 | 0.135 | ***0.002** |
| | | SB_TH-LB_TH | 0.093 | 0.037 | 0.149 | ***<0.001** |
| | | PL_TH-PI_TH | -0.084 | -0.144 | -0.023 | ***<0.001** |
| | | SB_TH-PI_TH | -0.066 | -0.122 | -0.009 | 0.008 |
| | | SB_TH-PL_TH | 0.018 | -0.035 | 0.070 | 0.996 |
| | Úlfljótsvatn | PI_UL-LB_UL | 0.081 | -0.031 | 0.192 | 0.437 |
| | | PL_UL-LB_UL | -0.009 | -0.092 | 0.074 | 1.000 |
| | | PL_UL-PI_UL | -0.090 | -0.199 | 0.020 | 0.237 |
| | Across Þingvallavatn and Úlfljótsvatn | LB_UL-LB_TH | -0.026 | -0.101 | 0.049 | 0.995 |
| | | PI_UL-LB_TH | 0.055 | -0.049 | 0.158 | 0.869 |
| | | PL_UL-LB_TH | -0.035 | -0.107 | 0.037 | 0.919 |
| | | PI_TH-LB_UL | 0.185 | 0.110 | 0.261 | ***<0.001** |
| | | PL_TH-LB_UL | 0.102 | 0.029 | 0.174 | ***<0.001** |
| | | SB_TH-LB_UL | 0.120 | 0.050 | 0.189 | ***<0.001** |
| | | PI_UL-PI_TH | -0.105 | -0.208 | -0.001 | 0.046 |
| | | PL_UL-PI_TH | -0.194 | -0.267 | -0.122 | ***<0.001** |
| | | PL_TH-PI_UL | 0.021 | -0.081 | 0.123 | 1.000 |
| | | SB_TH-PI_UL | 0.039 | -0.061 | 0.138 | 0.986 |
| | | PL_UL-PL_TH | -0.111 | -0.180 | -0.041 | ***<0.001** |
| | | SB_TH-PL_UL | 0.129 | 0.062 | 0.195 | ***<0.001** |
| Articular angular | AN populations | AN_Gufudalsá-AN_Brúará | -0.020 | -0.142 | 0.103 | 1.000 |
| | | AN_Lambeyrarkvísl-AN_Brúará | 0.082 | -0.050 | 0.213 | 0.683 |
| | | AN_Víðidalsá-AN_Brúará | 0.158 | 0.034 | 0.282 | ***0.002** |
| | | AN_Vífilsdalsá-AN_Brúará | 0.091 | -0.044 | 0.227 | 0.559 |
| | | AN_Lambeyrarkvísl-AN_Gufudalsá | 0.102 | 0.006 | 0.198 | 0.027 |
| | | AN_Víðidalsá-AN_Gufudalsá | 0.178 | 0.093 | 0.264 | ***<0.001** |
| | | AN_Vífilsdalsá-AN_Gufudalsá | 0.111 | 0.009 | 0.213 | 0.018 |
| | | AN_Víðidalsá-AN_Lambeyrarkvísl | 0.077 | -0.021 | 0.175 | 0.309 |
| | | AN_Vífilsdalsá-AN_Lambeyrarkvísl | 0.010 | -0.103 | 0.122 | 1.000 |
| | | AN_Vífilsdalsá-AN_Víðidalsá | -0.067 | -0.171 | 0.037 | 0.626 |
| | Brúará to lake morphs | Hest-AN_Brúará | 0.010 | -0.106 | 0.127 | 1.000 |
| | | LB_TH-AN_Brúará | -0.009 | -0.123 | 0.104 | 1.000 |
| | | LB_UL-AN_Brúará | -0.012 | -0.131 | 0.108 | 1.000 |
| | | PI_TH-AN_Brúará | 0.076 | -0.038 | 0.190 | 0.571 |
| | | PI_UL-AN_Brúará | 0.044 | -0.090 | 0.177 | 0.997 |
| | | PL_TH-AN_Brúará | 0.000 | -0.113 | 0.113 | 1.000 |
| | | PL_UL-AN_Brúará | -0.030 | -0.148 | 0.088 | 1.000 |
| | | SB_TH-AN_Brúará | 0.061 | -0.050 | 0.173 | 0.831 |
| | Gufudalsá to lake morphs | Hest-AN_Gufudalsá | 0.030 | -0.044 | 0.104 | 0.979 |
| | | LB_TH-AN_Gufudalsá | 0.010 | -0.059 | 0.080 | 1.000 |
| | | LB_UL-AN_Gufudalsá | 0.008 | -0.070 | 0.086 | 1.000 |
| | | PI_TH-AN_Gufudalsá | 0.096 | 0.026 | 0.166 | ***<0.001** |
| | | PI_UL-AN_Gufudalsá | 0.063 | -0.035 | 0.162 | 0.633 |
| | | PL_TH-AN_Gufudalsá | 0.020 | -0.048 | 0.088 | 0.999 |
| | | PL_UL-AN_Gufudalsá | -0.010 | -0.086 | 0.066 | 1.000 |
| | | SB_TH-AN_Gufudalsá | 0.081 | 0.015 | 0.147 | ***0.003** |
| | Lambeyrarkvísl to lake morphs | Hest-AN_Lambeyrarkvísl | -0.071 | -0.160 | 0.017 | 0.262 |
| | | LB_TH-AN_Lambeyrarkvísl | -0.091 | -0.176 | -0.006 | 0.022 |
| | | LB_UL-AN_Lambeyrarkvísl | -0.093 | -0.185 | -0.002 | 0.042 |
| | | PI_TH-AN_Lambeyrarkvísl | -0.006 | -0.091 | 0.079 | 1.000 |
| | | PI_UL-AN_Lambeyrarkvísl | -0.038 | -0.148 | 0.072 | 0.995 |
| | | PL_TH-AN_Lambeyrarkvísl | -0.082 | -0.165 | 0.002 | 0.060 |
| | | PL_UL-AN_Lambeyrarkvísl | -0.112 | -0.202 | -0.022 | ***0.003** |
| | | SB_TH-AN_Lambeyrarkvísl | -0.021 | -0.102 | 0.061 | 1.000 |
| | Víðidalsá to lake morphs | Hest-AN_Víðidalsá | -0.148 | -0.225 | -0.071 | ***<0.001** |
| | | LB_TH-AN_Víðidalsá | -0.168 | -0.240 | -0.095 | ***<0.001** |
| | | LB_UL-AN_Víðidalsá | -0.170 | -0.251 | -0.089 | ***<0.001** |

| | | | | | | |
|---|---|---|---|---|---|---|
| | | PI_TH-AN_Víðidalsá | -0.082 | -0.155 | -0.010 | 0.012 |
| | | PI_UL-AN_Víðidalsá | -0.115 | -0.216 | -0.014 | 0.010 |
| | | PL_TH-AN_Víðidalsá | -0.158 | -0.229 | -0.088 | ***<0.001** |
| | | PL_UL-AN_Víðidalsá | -0.188 | -0.267 | -0.110 | ***<0.001** |
| | | SB_TH-AN_Víðidalsá | -0.097 | -0.166 | -0.029 | ***<0.001** |
| | Vífisdalsá to lake morphs | Hest-AN_Vífilsdalsá | -0.081 | -0.176 | 0.014 | 0.183 |
| | | LB_TH-AN_Vífilsdalsá | -0.101 | -0.192 | -0.010 | 0.016 |
| | | LB_UL-AN_Vífilsdalsá | -0.103 | -0.201 | -0.005 | 0.029 |
| | | PI_TH-AN_Vífilsdalsá | -0.015 | -0.107 | 0.076 | 1.000 |
| | | PI_UL-AN_Vífilsdalsá | -0.048 | -0.163 | 0.067 | 0.976 |
| | | PL_TH-AN_Vífilsdalsá | -0.091 | -0.181 | -0.001 | 0.042 |
| | | PL_UL-AN_Vífilsdalsá | -0.121 | -0.217 | -0.025 | ***0.002** |
| | | SB_TH-AN_Vífilsdalsá | -0.030 | -0.118 | 0.058 | 0.995 |
| | Hestvatn | LB_TH-Hest | -0.020 | -0.079 | 0.039 | 0.996 |
| | | LB_UL-Hest | -0.022 | -0.091 | 0.046 | 0.997 |
| | | PI_TH-Hest | 0.066 | 0.007 | 0.125 | 0.015 |
| | | PI_UL-Hest | 0.033 | -0.058 | 0.124 | 0.992 |
| | | PL_TH-Hest | -0.010 | -0.067 | 0.046 | 1.000 |
| | | PL_UL-Hest | -0.040 | -0.106 | 0.025 | 0.703 |
| | | SB_TH-Hest | 0.051 | -0.003 | 0.104 | 0.085 |
| | Þingvallavatn | PI_TH-LB_TH | 0.086 | 0.032 | 0.139 | ***<0.001** |
| | | PL_TH-LB_TH | 0.009 | -0.041 | 0.060 | 1.000 |
| | | SB_TH-LB_TH | 0.071 | 0.023 | 0.118 | ***<0.001** |
| | | PL_TH-PI_TH | -0.076 | -0.127 | -0.025 | ***<0.001** |
| | | SB_TH-PI_TH | -0.015 | -0.063 | 0.033 | 0.998 |
| | | SB_TH-PL_TH | 0.061 | 0.016 | 0.106 | ***0.001** |
| | Úlfljótsvatn | PI_UL-LB_UL | 0.055 | -0.039 | 0.150 | 0.766 |
| | | PL_UL-LB_UL | -0.018 | -0.089 | 0.052 | 1.000 |
| | | PL_UL-PI_UL | -0.074 | -0.166 | 0.019 | 0.288 |
| | Across Þingvallavatn and Úlfljótsvatn | LB_UL-LB_TH | -0.002 | -0.066 | 0.062 | 1.000 |
| | | PI_UL-LB_TH | 0.053 | -0.035 | 0.141 | 0.723 |
| | | PL_UL-LB_TH | -0.021 | -0.081 | 0.040 | 0.996 |
| | | PI_TH-LB_UL | 0.088 | 0.024 | 0.152 | ***<0.001** |
| | | PL_TH-LB_UL | 0.012 | -0.050 | 0.074 | 1.000 |
| | | SB_TH-LB_UL | 0.073 | 0.014 | 0.132 | ***0.003** |
| | | PI_UL-PI_TH | -0.033 | -0.120 | 0.055 | 0.991 |
| | | PL_UL-PI_TH | -0.106 | -0.167 | -0.045 | ***<0.001** |
| | | PL_TH-PI_UL | -0.044 | -0.130 | 0.043 | 0.901 |
| | | SB_TH-PI_UL | 0.018 | -0.067 | 0.102 | 1.000 |
| | | PL_UL-PL_TH | -0.030 | -0.089 | 0.029 | 0.897 |
| | | SB_TH-PL_UL | 0.091 | 0.035 | 0.147 | ***<0.001** |
| Qudrate | AN populations | AN_Gufudalsá-AN_Brúará | 0.035 | -0.091 | 0.161 | 0.999 |
| | | AN_Lambeyrarkvísl-AN_Brúará | 0.088 | -0.045 | 0.222 | 0.590 |
| | | AN_Víðidalsá-AN_Brúará | 0.194 | 0.066 | 0.321 | ***<0.001** |
| | | AN_Vífilsdalsá-AN_Brúará | 0.083 | -0.055 | 0.220 | 0.731 |
| | | AN_Lambeyrarkvísl-AN_Gufudalsá | 0.054 | -0.038 | 0.145 | 0.768 |
| | | AN_Víðidalsá-AN_Gufudalsá | 0.159 | 0.077 | 0.240 | ***<0.001** |
| | | AN_Vífilsdalsá-AN_Gufudalsá | 0.048 | -0.050 | 0.145 | 0.917 |
| | | AN_Víðidalsá-AN_Lambeyrarkvísl | 0.105 | 0.011 | 0.199 | 0.013 |
| | | AN_Vífilsdalsá-AN_Lambeyrarkvísl | -0.006 | -0.113 | 0.102 | 1.000 |
| | | AN_Vífilsdalsá-AN_Víðidalsá | -0.111 | -0.210 | -0.012 | 0.014 |
| | Brúará to lake morphs | Hest-AN_Brúará | 0.026 | -0.094 | 0.146 | 1.000 |
| | | LB_TH-AN_Brúará | 0.113 | -0.005 | 0.231 | 0.078 |
| | | LB_UL-AN_Brúará | 0.025 | -0.098 | 0.148 | 1.000 |
| | | PI_TH-AN_Brúará | 0.121 | 0.002 | 0.239 | 0.040 |
| | | PI_UL-AN_Brúará | 0.046 | -0.089 | 0.182 | 0.996 |
| | | PL_TH-AN_Brúará | 0.030 | -0.087 | 0.147 | 1.000 |
| | | PL_UL-AN_Brúará | 0.003 | -0.118 | 0.124 | 1.000 |
| | | SB_TH-AN_Brúará | 0.146 | 0.030 | 0.262 | ***0.002** |
| | Gufudalsá to lake morphs | Hest-AN_Gufudalsá | -0.009 | -0.080 | 0.062 | 1.000 |
| | | LB_TH-AN_Gufudalsá | 0.078 | 0.011 | 0.144 | 0.008 |

| | | | | | | |
|---|---|---|---|---|---|---|
| | | LB_UL-AN_Gufudalsá | -0.010 | -0.084 | 0.065 | 1.000 |
| | | PI_TH-AN_Gufudalsá | 0.086 | 0.019 | 0.153 | ***0.002** |
| | | PI_UL-AN_Gufudalsá | 0.011 | -0.083 | 0.106 | 1.000 |
| | | PL_TH-AN_Gufudalsá | -0.005 | -0.070 | 0.060 | 1.000 |
| | | PL_UL-AN_Gufudalsá | -0.032 | -0.104 | 0.041 | 0.962 |
| | | SB_TH-AN_Gufudalsá | 0.111 | 0.048 | 0.174 | ***<0.001** |
| | Lambeyrarkvísl to lake morphs | Hest-AN_Lambeyrarkvísl | -0.062 | -0.147 | 0.022 | 0.402 |
| | | LB_TH-AN_Lambeyrarkvísl | 0.024 | -0.057 | 0.105 | 0.999 |
| | | LB_UL-AN_Lambeyrarkvísl | -0.063 | -0.151 | 0.025 | 0.445 |
| | | PI_TH-AN_Lambeyrarkvísl | 0.032 | -0.049 | 0.113 | 0.983 |
| | | PI_UL-AN_Lambeyrarkvísl | -0.042 | -0.147 | 0.063 | 0.982 |
| | | PL_TH-AN_Lambeyrarkvísl | -0.058 | -0.138 | 0.021 | 0.415 |
| | | PL_UL-AN_Lambeyrarkvísl | -0.085 | -0.171 | 0.000 | 0.052 |
| | | SB_TH-AN_Lambeyrarkvísl | 0.057 | -0.020 | 0.135 | 0.406 |
| | Víðidalsá to lake morphs | Hest-AN_Víðidalsá | -0.168 | -0.241 | -0.094 | ***<0.001** |
| | | LB_TH-AN_Víðidalsá | -0.081 | -0.150 | -0.012 | 0.008 |
| | | LB_UL-AN_Víðidalsá | -0.168 | -0.245 | -0.091 | ***<0.001** |
| | | PI_TH-AN_Víðidalsá | -0.073 | -0.142 | -0.003 | 0.030 |
| | | PI_UL-AN_Víðidalsá | -0.147 | -0.244 | -0.051 | ***<0.001** |
| | | PL_TH-AN_Víðidalsá | -0.164 | -0.231 | -0.096 | ***<0.001** |
| | | PL_UL-AN_Víðidalsá | -0.191 | -0.266 | -0.116 | ***<0.001** |
| | | SB_TH-AN_Víðidalsá | -0.048 | -0.113 | 0.018 | 0.431 |
| | Vífisdalsá to lake morphs | Hest-AN_Vífilsdalsá | -0.057 | -0.147 | 0.034 | 0.671 |
| | | LB_TH-AN_Vífilsdalsá | 0.030 | -0.057 | 0.117 | 0.995 |
| | | LB_UL-AN_Vífilsdalsá | -0.058 | -0.151 | 0.036 | 0.699 |
| | | PI_TH-AN_Vífilsdalsá | 0.038 | -0.049 | 0.125 | 0.966 |
| | | PI_UL-AN_Vífilsdalsá | -0.037 | -0.146 | 0.073 | 0.997 |
| | | PL_TH-AN_Vífilsdalsá | -0.053 | -0.139 | 0.033 | 0.703 |
| | | PL_UL-AN_Vífilsdalsá | -0.080 | -0.172 | 0.012 | 0.165 |
| | | SB_TH-AN_Vífilsdalsá | 0.063 | -0.021 | 0.147 | 0.381 |
| | Hestvatn | LB_TH-Hest | 0.087 | 0.031 | 0.143 | ***<0.001** |
| | | LB_UL-Hest | -0.001 | -0.066 | 0.065 | 1.000 |
| | | PI_TH-Hest | 0.095 | 0.038 | 0.151 | ***<0.001** |
| | | PI_UL-Hest | 0.020 | -0.067 | 0.107 | 1.000 |
| | | PL_TH-Hest | 0.004 | -0.050 | 0.058 | 1.000 |
| | | PL_UL-Hest | -0.023 | -0.086 | 0.040 | 0.992 |
| | | SB_TH-Hest | 0.120 | 0.068 | 0.171 | ***<0.001** |
| | Þingvallavatn | PI_TH-LB_TH | 0.008 | -0.043 | 0.059 | 1.000 |
| | | PL_TH-LB_TH | -0.083 | -0.131 | -0.034 | ***<0.001** |
| | | SB_TH-LB_TH | 0.033 | -0.013 | 0.079 | 0.433 |
| | | PL_TH-PI_TH | -0.091 | -0.139 | -0.042 | ***<0.001** |
| | | SB_TH-PI_TH | 0.025 | -0.021 | 0.071 | 0.835 |
| | | SB_TH-PL_TH | 0.116 | 0.073 | 0.159 | ***<0.001** |
| | Úlfljótsvatn | PI_UL-LB_UL | 0.021 | -0.069 | 0.111 | 1.000 |
| | | PL_UL-LB_UL | -0.022 | -0.090 | 0.045 | 0.997 |
| | | PL_UL-PI_UL | -0.043 | -0.132 | 0.045 | 0.921 |
| | Across Þingvallavatn and Úlfljótsvatn | LB_UL-LB_TH | -0.087 | -0.148 | -0.026 | ***<0.001** |
| | | PI_UL-LB_TH | -0.066 | -0.150 | 0.017 | 0.289 |
| | | PL_UL-LB_TH | -0.110 | -0.168 | -0.051 | ***<0.001** |
| | | PI_TH-LB_UL | 0.095 | 0.034 | 0.157 | ***<0.001** |
| | | PL_TH-LB_UL | 0.005 | -0.054 | 0.064 | 1.000 |
| | | SB_TH-LB_UL | 0.121 | 0.064 | 0.177 | ***<0.001** |
| | | PI_UL-PI_TH | -0.074 | -0.158 | 0.010 | 0.144 |
| | | PL_UL-PI_TH | -0.118 | -0.176 | -0.059 | ***<0.001** |
| | | PL_TH-PI_UL | -0.016 | -0.099 | 0.066 | 1.000 |
| | | SB_TH-PI_UL | 0.100 | 0.019 | 0.181 | ***0.003** |
| | | PL_UL-PL_TH | -0.027 | -0.083 | 0.029 | 0.929 |
| | | SB_TH-PL_UL | 0.143 | 0.089 | 0.197 | ***<0.001** |
| Premaxilla | AN populations | AN_Gufudalsá-AN_Brúará | 0.022 | -0.140 | 0.185 | 1.000 |
| | | AN_Lambeyrarkvísl-AN_Brúará | 0.142 | -0.033 | 0.317 | 0.258 |
| | | AN_Víðidalsá-AN_Brúará | 0.219 | 0.054 | 0.384 | ***0.001** |

| | | | | | | |
|---|---|---|---|---|---|---|
| | | AN_Vífilsdalsá-AN_Brúará | 0.202 | 0.022 | 0.383 | 0.013 |
| | | AN_Lambeyrarkvísl-AN_Gufudalsá | 0.119 | -0.009 | 0.247 | 0.096 |
| | | AN_Víðidalsá-AN_Gufudalsá | 0.197 | 0.083 | 0.310 | ***<0.001** |
| | | AN_Vífilsdalsá-AN_Gufudalsá | 0.180 | 0.044 | 0.316 | ***0.001** |
| | | AN_Víðidalsá-AN_Lambeyrarkvísl | 0.077 | -0.053 | 0.208 | 0.747 |
| | | AN_Vífilsdalsá-AN_Lambeyrarkvísl | 0.061 | -0.089 | 0.211 | 0.980 |
| | | AN_Vífilsdalsá-AN_Víðidalsá | -0.017 | -0.155 | 0.121 | 1.000 |
| | Brúará to lake morphs | Hest-AN_Brúará | 0.019 | -0.136 | 0.174 | 1.000 |
| | | LB_TH-AN_Brúará | 0.047 | -0.105 | 0.198 | 0.998 |
| | | LB_UL-AN_Brúará | 0.024 | -0.134 | 0.183 | 1.000 |
| | | PI_TH-AN_Brúará | 0.098 | -0.053 | 0.250 | 0.619 |
| | | PI_UL-AN_Brúará | -0.008 | -0.186 | 0.170 | 1.000 |
| | | PL_TH-AN_Brúará | -0.062 | -0.212 | 0.088 | 0.976 |
| | | PL_UL-AN_Brúará | -0.022 | -0.179 | 0.134 | 1.000 |
| | | SB_TH-AN_Brúará | 0.159 | 0.011 | 0.307 | 0.022 |
| | Gufudalsá to lake morphs | Hest-AN_Gufudalsá | -0.004 | -0.102 | 0.095 | 1.000 |
| | | LB_TH-AN_Gufudalsá | 0.024 | -0.069 | 0.117 | 1.000 |
| | | LB_UL-AN_Gufudalsá | 0.002 | -0.102 | 0.106 | 1.000 |
| | | PI_TH-AN_Gufudalsá | 0.076 | -0.017 | 0.169 | 0.248 |
| | | PI_UL-AN_Gufudalsá | -0.030 | -0.162 | 0.101 | 1.000 |
| | | PL_TH-AN_Gufudalsá | -0.085 | -0.175 | 0.005 | 0.090 |
| | | PL_UL-AN_Gufudalsá | -0.045 | -0.146 | 0.056 | 0.961 |
| | | SB_TH-AN_Gufudalsá | 0.137 | 0.050 | 0.224 | ***<0.001** |
| | Lambeyrarkvísl to lake morphs | Hest-AN_Lambeyrarkvísl | -0.123 | -0.240 | -0.005 | 0.031 |
| | | LB_TH-AN_Lambeyrarkvísl | -0.095 | -0.208 | 0.018 | 0.201 |
| | | LB_UL-AN_Lambeyrarkvísl | -0.118 | -0.240 | 0.005 | 0.073 |
| | | PI_TH-AN_Lambeyrarkvísl | -0.043 | -0.156 | 0.070 | 0.988 |
| | | PI_UL-AN_Lambeyrarkvísl | -0.150 | -0.296 | -0.003 | 0.039 |
| | | PL_TH-AN_Lambeyrarkvísl | -0.204 | -0.314 | -0.093 | ***<0.001** |
| | | PL_UL-AN_Lambeyrarkvísl | -0.164 | -0.283 | -0.044 | ***<0.001** |
| | | SB_TH-AN_Lambeyrarkvísl | 0.018 | -0.090 | 0.126 | 1.000 |
| | Víðidalsá to lake morphs | Hest-AN_Víðidalsá | -0.200 | -0.302 | -0.098 | ***<0.001** |
| | | LB_TH-AN_Víðidalsá | -0.172 | -0.269 | -0.076 | ***<0.001** |
| | | LB_UL-AN_Víðidalsá | -0.195 | -0.302 | -0.088 | ***<0.001** |
| | | PI_TH-AN_Víðidalsá | -0.121 | -0.217 | -0.024 | ***0.003** |
| | | PI_UL-AN_Víðidalsá | -0.227 | -0.361 | -0.093 | ***<0.001** |
| | | PL_TH-AN_Víðidalsá | -0.281 | -0.375 | -0.187 | ***<0.001** |
| | | PL_UL-AN_Víðidalsá | -0.241 | -0.346 | -0.137 | ***<0.001** |
| | | SB_TH-AN_Víðidalsá | -0.060 | -0.151 | 0.032 | 0.606 |
| | Vífisdalsá to lake morphs | Hest-AN_Vífilsdalsá | -0.184 | -0.310 | -0.058 | ***<0.001** |
| | | LB_TH-AN_Vífilsdalsá | -0.156 | -0.277 | -0.034 | ***0.002** |
| | | LB_UL-AN_Vífilsdalsá | -0.178 | -0.309 | -0.048 | ***<0.001** |
| | | PI_TH-AN_Vífilsdalsá | -0.104 | -0.226 | 0.018 | 0.183 |
| | | PI_UL-AN_Vífilsdalsá | -0.210 | -0.363 | -0.057 | ***<0.001** |
| | | PL_TH-AN_Vífilsdalsá | -0.265 | -0.384 | -0.145 | ***<0.001** |
| | | PL_UL-AN_Vífilsdalsá | -0.225 | -0.352 | -0.097 | ***<0.001** |
| | | SB_TH-AN_Vífilsdalsá | -0.043 | -0.160 | 0.074 | 0.992 |
| | Hestvatn | LB_TH-Hest | 0.028 | -0.050 | 0.106 | 0.994 |
| | | LB_UL-Hest | 0.005 | -0.086 | 0.097 | 1.000 |
| | | PI_TH-Hest | 0.080 | 0.001 | 0.158 | 0.043 |
| | | PI_UL-Hest | -0.027 | -0.148 | 0.095 | 1.000 |
| | | PL_TH-Hest | -0.081 | -0.156 | -0.006 | 0.021 |
| | | PL_UL-Hest | -0.041 | -0.129 | 0.047 | 0.941 |
| | | SB_TH-Hest | 0.141 | 0.069 | 0.212 | ***<0.001** |
| | Þingvallavatn | PI_TH-LB_TH | 0.052 | -0.019 | 0.123 | 0.426 |
| | | PL_TH-LB_TH | -0.109 | -0.176 | -0.042 | ***<0.001** |
| | | SB_TH-LB_TH | 0.113 | 0.050 | 0.176 | ***<0.001** |
| | | PL_TH-PI_TH | -0.161 | -0.228 | -0.093 | ***<0.001** |
| | | SB_TH-PI_TH | 0.061 | -0.003 | 0.125 | 0.074 |
| | | SB_TH-PL_TH | 0.222 | 0.162 | 0.281 | ***<0.001** |
| | Úlfljótsvatn | PI_UL-LB_UL | -0.032 | -0.158 | 0.094 | 1.000 |

| | | | | | | |
|---|---|---|---|---|---|---|
| | | PL_UL-LB_UL | -0.046 | -0.140 | 0.047 | 0.914 |
| | | PL_UL-PI_UL | -0.014 | -0.138 | 0.109 | 1.000 |
| | Across Þingvallavatn and Úlfljótsvatn | LB_UL-LB_TH | -0.022 | -0.107 | 0.062 | 1.000 |
| | | PI_UL-LB_TH | -0.054 | -0.171 | 0.062 | 0.942 |
| | | PL_UL-LB_TH | -0.069 | -0.150 | 0.012 | 0.195 |
| | | PI_TH-LB_UL | 0.074 | -0.011 | 0.159 | 0.162 |
| | | PL_TH-LB_UL | -0.086 | -0.168 | -0.004 | 0.029 |
| | | SB_TH-LB_UL | 0.135 | 0.057 | 0.214 | *<0.001 |
| | | PI_UL-PI_TH | -0.106 | -0.223 | 0.011 | 0.118 |
| | | PL_UL-PI_TH | -0.121 | -0.202 | -0.039 | *<0.001 |
| | | PL_TH-PI_UL | -0.054 | -0.169 | 0.060 | 0.936 |
| | | SB_TH-PI_UL | 0.167 | 0.055 | 0.280 | *<0.001 |
| | | PL_UL-PL_TH | 0.040 | -0.038 | 0.118 | 0.891 |
| | | SB_TH-PL_UL | 0.182 | 0.107 | 0.256 | *<0.001 |
| Maxilla | AN populations | AN_Gufudalsá-AN_Brúará | 0.012 | -0.128 | 0.153 | 1.000 |
| | | AN_Lambeyrarkvísl-AN_Brúará | 0.190 | 0.039 | 0.341 | *0.002 |
| | | AN_Víðidalsá-AN_Brúará | 0.247 | 0.104 | 0.389 | *<0.001 |
| | | AN_Vífilsdalsá-AN_Brúará | 0.185 | 0.029 | 0.340 | *0.006 |
| | | AN_Lambeyrarkvísl-AN_Gufudalsá | 0.177 | 0.067 | 0.288 | *<0.001 |
| | | AN_Víðidalsá-AN_Gufudalsá | 0.234 | 0.136 | 0.332 | *<0.001 |
| | | AN_Vífilsdalsá-AN_Gufudalsá | 0.172 | 0.055 | 0.289 | *<0.001 |
| | | AN_Víðidalsá-AN_Lambeyrarkvísl | 0.057 | -0.056 | 0.169 | 0.901 |
| | | AN_Vífilsdalsá-AN_Lambeyrarkvísl | -0.005 | -0.134 | 0.124 | 1.000 |
| | | AN_Vífilsdalsá-AN_Víðidalsá | -0.062 | -0.181 | 0.057 | 0.878 |
| | Brúará to lake morphs | Hest-AN_Brúará | 0.088 | -0.046 | 0.221 | 0.602 |
| | | LB_TH-AN_Brúará | 0.095 | -0.035 | 0.226 | 0.421 |
| | | LB_UL-AN_Brúará | 0.031 | -0.105 | 0.168 | 1.000 |
| | | PI_TH-AN_Brúará | 0.181 | 0.050 | 0.311 | *<0.001 |
| | | PI_UL-AN_Brúará | 0.082 | -0.071 | 0.235 | 0.858 |
| | | PL_TH-AN_Brúará | 0.089 | -0.040 | 0.218 | 0.519 |
| | | PL_UL-AN_Brúará | -0.020 | -0.155 | 0.115 | 1.000 |
| | | SB_TH-AN_Brúará | 0.153 | 0.026 | 0.281 | *0.005 |
| | Gufudalsá to lake morphs | Hest-AN_Gufudalsá | 0.075 | -0.010 | 0.160 | 0.145 |
| | | LB_TH-AN_Gufudalsá | 0.083 | 0.003 | 0.163 | 0.034 |
| | | LB_UL-AN_Gufudalsá | 0.019 | -0.071 | 0.109 | 1.000 |
| | | PI_TH-AN_Gufudalsá | 0.168 | 0.088 | 0.249 | *<0.001 |
| | | PI_UL-AN_Gufudalsá | 0.069 | -0.044 | 0.183 | 0.704 |
| | | PL_TH-AN_Gufudalsá | 0.077 | -0.001 | 0.155 | 0.058 |
| | | PL_UL-AN_Gufudalsá | -0.033 | -0.120 | 0.055 | 0.990 |
| | | SB_TH-AN_Gufudalsá | 0.141 | 0.066 | 0.217 | *<0.001 |
| | Lambeyrarkvísl to lake morphs | Hest-AN_Lambeyrarkvísl | -0.102 | -0.204 | -0.001 | 0.045 |
| | | LB_TH-AN_Lambeyrarkvísl | -0.095 | -0.192 | 0.003 | 0.065 |
| | | LB_UL-AN_Lambeyrarkvísl | -0.159 | -0.264 | -0.053 | *<0.001 |
| | | PI_TH-AN_Lambeyrarkvísl | -0.009 | -0.106 | 0.088 | 1.000 |
| | | PI_UL-AN_Lambeyrarkvísl | -0.108 | -0.234 | 0.018 | 0.179 |
| | | PL_TH-AN_Lambeyrarkvísl | -0.101 | -0.196 | -0.005 | 0.028 |
| | | PL_UL-AN_Lambeyrarkvísl | -0.210 | -0.313 | -0.107 | *<0.001 |
| | | SB_TH-AN_Lambeyrarkvísl | -0.036 | -0.130 | 0.057 | 0.986 |
| | Víðidalsá to lake morphs | Hest-AN_Víðidalsá | -0.159 | -0.247 | -0.071 | *<0.001 |
| | | LB_TH-AN_Víðidalsá | -0.151 | -0.234 | -0.068 | *<0.001 |
| | | LB_UL-AN_Víðidalsá | -0.215 | -0.308 | -0.123 | *<0.001 |
| | | PI_TH-AN_Víðidalsá | -0.066 | -0.149 | 0.018 | 0.297 |
| | | PI_UL-AN_Víðidalsá | -0.165 | -0.280 | -0.049 | *<0.001 |
| | | PL_TH-AN_Víðidalsá | -0.157 | -0.239 | -0.076 | *<0.001 |
| | | PL_UL-AN_Víðidalsá | -0.267 | -0.357 | -0.177 | *<0.001 |
| | | SB_TH-AN_Víðidalsá | -0.093 | -0.172 | -0.014 | *0.006 |
| | Vífisdalsá to lake morphs | Hest-AN_Vífilsdalsá | -0.097 | -0.205 | 0.012 | 0.134 |
| | | LB_TH-AN_Vífilsdalsá | -0.089 | -0.194 | 0.015 | 0.188 |
| | | LB_UL-AN_Vífilsdalsá | -0.153 | -0.266 | -0.041 | *0.001 |
| | | PI_TH-AN_Vífilsdalsá | -0.004 | -0.109 | 0.101 | 1.000 |
| | | PI_UL-AN_Vífilsdalsá | -0.103 | -0.235 | 0.029 | 0.314 |

| | | | | | | |
|---|---|---|---|---|---|---|
| | | PL_TH-AN_Vífilsdalsá | -0.095 | -0.198 | 0.008 | 0.101 |
| | | PL_UL-AN_Vífilsdalsá | -0.205 | -0.315 | -0.095 | ***<0.001** |
| | | SB_TH-AN_Vífilsdalsá | -0.031 | -0.132 | 0.070 | 0.998 |
| | Hestvatn | LB_TH-Hest | 0.008 | -0.060 | 0.075 | 1.000 |
| | | LB_UL-Hest | -0.056 | -0.135 | 0.022 | 0.452 |
| | | PI_TH-Hest | 0.093 | 0.026 | 0.161 | ***<0.001** |
| | | PI_UL-Hest | -0.006 | -0.110 | 0.099 | 1.000 |
| | | PL_TH-Hest | 0.002 | -0.063 | 0.066 | 1.000 |
| | | PL_UL-Hest | -0.108 | -0.183 | -0.032 | ***<0.001** |
| | | SB_TH-Hest | 0.066 | 0.004 | 0.128 | 0.024 |
| | Þingvallavatn | PI_TH-LB_TH | 0.085 | 0.024 | 0.147 | ***<0.001** |
| | | PL_TH-LB_TH | -0.006 | -0.064 | 0.052 | 1.000 |
| | | SB_TH-LB_TH | 0.058 | 0.003 | 0.113 | 0.026 |
| | | PL_TH-PI_TH | -0.092 | -0.150 | -0.033 | ***<0.001** |
| | | SB_TH-PI_TH | -0.027 | -0.082 | 0.028 | 0.911 |
| | | SB_TH-PL_TH | 0.064 | 0.013 | 0.116 | ***0.003** |
| | Úlfljótsvatn | PI_UL-LB_UL | 0.051 | -0.058 | 0.159 | 0.942 |
| | | PL_UL-LB_UL | -0.051 | -0.132 | 0.029 | 0.651 |
| | | PL_UL-PI_UL | -0.102 | -0.208 | 0.004 | 0.075 |
| | Across Þingvallavatn and Úlfljótsvatn | LB_UL-LB_TH | -0.064 | -0.137 | 0.009 | 0.155 |
| | | PI_UL-LB_TH | -0.014 | -0.114 | 0.087 | 1.000 |
| | | PL_UL-LB_TH | -0.116 | -0.185 | -0.046 | ***<0.001** |
| | | PI_TH-LB_UL | 0.150 | 0.076 | 0.223 | ***<0.001** |
| | | PL_TH-LB_UL | 0.058 | -0.013 | 0.129 | 0.244 |
| | | SB_TH-LB_UL | 0.122 | 0.054 | 0.190 | ***<0.001** |
| | | PI_UL-PI_TH | -0.099 | -0.200 | 0.002 | 0.060 |
| | | PL_UL-PI_TH | -0.201 | -0.271 | -0.131 | ***<0.001** |
| | | PL_TH-PI_UL | 0.007 | -0.092 | 0.106 | 1.000 |
| | | SB_TH-PI_UL | 0.072 | -0.025 | 0.169 | 0.401 |
| | | PL_UL-PL_TH | -0.109 | -0.177 | -0.042 | ***<0.001** |
| | | SB_TH-PL_UL | 0.174 | 0.109 | 0.238 | ***<0.001** |
| Supramaxilla | AN populations | AN_Gufudalsá-AN_Brúará | 0.019 | -0.118 | 0.157 | 1.000 |
| | | AN_Lambeyrarkvísl-AN_Brúará | 0.188 | 0.038 | 0.337 | ***0.002** |
| | | AN_Víðidalsá-AN_Brúará | 0.207 | 0.068 | 0.346 | ***<0.001** |
| | | AN_Vífilsdalsá-AN_Brúará | 0.154 | 0.002 | 0.306 | 0.045 |
| | | AN_Lambeyrarkvísl-AN_Gufudalsá | 0.168 | 0.058 | 0.279 | ***<0.001** |
| | | AN_Víðidalsá-AN_Gufudalsá | 0.187 | 0.092 | 0.283 | ***<0.001** |
| | | AN_Vífilsdalsá-AN_Gufudalsá | 0.135 | 0.020 | 0.249 | ***0.007** |
| | | AN_Víðidalsá-AN_Lambeyrarkvísl | 0.019 | -0.094 | 0.132 | 1.000 |
| | | AN_Vífilsdalsá-AN_Lambeyrarkvísl | -0.034 | -0.163 | 0.095 | 1.000 |
| | | AN_Vífilsdalsá-AN_Víðidalsá | -0.053 | -0.169 | 0.064 | 0.953 |
| | Brúará to lake morphs | Hest-AN_Brúará | 0.012 | -0.119 | 0.142 | 1.000 |
| | | LB_TH-AN_Brúará | 0.062 | -0.066 | 0.189 | 0.925 |
| | | LB_UL-AN_Brúará | -0.007 | -0.141 | 0.126 | 1.000 |
| | | PI_TH-AN_Brúará | 0.147 | 0.019 | 0.274 | 0.010 |
| | | PI_UL-AN_Brúará | 0.046 | -0.104 | 0.195 | 0.999 |
| | | PL_TH-AN_Brúará | 0.041 | -0.086 | 0.168 | 0.997 |
| | | PL_UL-AN_Brúará | -0.020 | -0.152 | 0.112 | 1.000 |
| | | SB_TH-AN_Brúará | 0.128 | 0.003 | 0.253 | 0.039 |
| | Gufudalsá to lake morphs | Hest-AN_Gufudalsá | -0.008 | -0.091 | 0.076 | 1.000 |
| | | LB_TH-AN_Gufudalsá | 0.043 | -0.036 | 0.121 | 0.844 |
| | | LB_UL-AN_Gufudalsá | -0.026 | -0.114 | 0.061 | 0.999 |
| | | PI_TH-AN_Gufudalsá | 0.127 | 0.049 | 0.206 | ***<0.001** |
| | | PI_UL-AN_Gufudalsá | 0.026 | -0.084 | 0.137 | 1.000 |
| | | PL_TH-AN_Gufudalsá | 0.022 | -0.055 | 0.099 | 0.999 |
| | | PL_UL-AN_Gufudalsá | -0.039 | -0.125 | 0.046 | 0.946 |
| | | SB_TH-AN_Gufudalsá | 0.109 | 0.035 | 0.183 | ***<0.001** |
| | Lambeyrarkvísl to lake morphs | Hest-AN_Lambeyrarkvísl | -0.176 | -0.278 | -0.074 | ***<0.001** |
| | | LB_TH-AN_Lambeyrarkvísl | -0.126 | -0.224 | -0.028 | ***0.002** |
| | | LB_UL-AN_Lambeyrarkvísl | -0.195 | -0.301 | -0.089 | ***<0.001** |
| | | PI_TH-AN_Lambeyrarkvísl | -0.041 | -0.140 | 0.058 | 0.975 |

| | | | | | | |
|---|---|---|---|---|---|---|
| | | PI_UL-AN_Lambeyrarkvísl | -0.142 | -0.268 | -0.016 | 0.012 |
| | | PL_TH-AN_Lambeyrarkvísl | -0.147 | -0.244 | -0.049 | ***<0.001** |
| | | PL_UL-AN_Lambeyrarkvísl | -0.208 | -0.312 | -0.104 | ***<0.001** |
| | | SB_TH-AN_Lambeyrarkvísl | -0.060 | -0.155 | 0.035 | 0.669 |
| | Víðidalsá to lake morphs | Hest-AN_Víðidalsá | -0.195 | -0.281 | -0.109 | ***<0.001** |
| | | LB_TH-AN_Víðidalsá | -0.145 | -0.226 | -0.064 | ***<0.001** |
| | | LB_UL-AN_Víðidalsá | -0.214 | -0.304 | -0.123 | ***<0.001** |
| | | PI_TH-AN_Víðidalsá | -0.060 | -0.142 | 0.021 | 0.406 |
| | | PI_UL-AN_Víðidalsá | -0.161 | -0.274 | -0.048 | ***<0.001** |
| | | PL_TH-AN_Víðidalsá | -0.166 | -0.245 | -0.086 | ***<0.001** |
| | | PL_UL-AN_Víðidalsá | -0.227 | -0.315 | -0.139 | ***<0.001** |
| | | SB_TH-AN_Víðidalsá | -0.079 | -0.156 | -0.001 | 0.041 |
| | Vífisdalsá to lake morphs | Hest-AN_Vífilsdalsá | -0.142 | -0.248 | -0.036 | ***0.001** |
| | | LB_TH-AN_Vífilsdalsá | -0.092 | -0.195 | 0.010 | 0.127 |
| | | LB_UL-AN_Vífilsdalsá | -0.161 | -0.271 | -0.051 | ***<0.001** |
| | | PI_TH-AN_Vífilsdalsá | -0.007 | -0.110 | 0.095 | 1.000 |
| | | PI_UL-AN_Vífilsdalsá | -0.108 | -0.237 | 0.021 | 0.205 |
| | | PL_TH-AN_Vífilsdalsá | -0.113 | -0.214 | -0.012 | 0.014 |
| | | PL_UL-AN_Vífilsdalsá | -0.174 | -0.282 | -0.066 | ***<0.001** |
| | | SB_TH-AN_Vífilsdalsá | -0.026 | -0.125 | 0.073 | 1.000 |
| | Hestvatn | LB_TH-Hest | 0.050 | -0.016 | 0.116 | 0.354 |
| | | LB_UL-Hest | -0.019 | -0.096 | 0.058 | 1.000 |
| | | PI_TH-Hest | 0.135 | 0.069 | 0.201 | ***<0.001** |
| | | PI_UL-Hest | 0.034 | -0.068 | 0.136 | 0.997 |
| | | PL_TH-Hest | 0.029 | -0.034 | 0.093 | 0.947 |
| | | PL_UL-Hest | -0.032 | -0.106 | 0.042 | 0.969 |
| | | SB_TH-Hest | 0.116 | 0.056 | 0.177 | ***<0.001** |
| | Þingvallavatn | PI_TH-LB_TH | 0.085 | 0.025 | 0.145 | ***<0.001** |
| | | PL_TH-LB_TH | -0.021 | -0.078 | 0.037 | 0.993 |
| | | SB_TH-LB_TH | 0.066 | 0.013 | 0.120 | ***0.003** |
| | | PL_TH-PI_TH | -0.105 | -0.163 | -0.048 | ***<0.001** |
| | | SB_TH-PI_TH | -0.019 | -0.073 | 0.036 | 0.995 |
| | | SB_TH-PL_TH | 0.087 | 0.036 | 0.138 | ***<0.001** |
| | Úlfljótsvatn | PI_UL-LB_UL | 0.053 | -0.053 | 0.159 | 0.909 |
| | | PL_UL-LB_UL | -0.013 | -0.092 | 0.066 | 1.000 |
| | | PL_UL-PI_UL | -0.066 | -0.170 | 0.038 | 0.660 |
| | Across Þingvallavatn and Úlfljótsvatn | LB_UL-LB_TH | -0.069 | -0.141 | 0.003 | 0.071 |
| | | PI_UL-LB_TH | -0.016 | -0.115 | 0.082 | 1.000 |
| | | PL_UL-LB_TH | -0.082 | -0.150 | -0.013 | ***0.005** |
| | | PI_TH-LB_UL | 0.154 | 0.082 | 0.226 | ***<0.001** |
| | | PL_TH-LB_UL | 0.048 | -0.021 | 0.118 | 0.512 |
| | | SB_TH-LB_UL | 0.135 | 0.068 | 0.202 | ***<0.001** |
| | | PI_UL-PI_TH | -0.101 | -0.200 | -0.002 | 0.039 |
| | | PL_UL-PI_TH | -0.167 | -0.235 | -0.098 | ***<0.001** |
| | | PL_TH-PI_UL | -0.004 | -0.102 | 0.093 | 1.000 |
| | | SB_TH-PI_UL | 0.082 | -0.013 | 0.177 | 0.166 |
| | | PL_UL-PL_TH | -0.061 | -0.128 | 0.005 | 0.107 |
| | | SB_TH-PL_UL | 0.148 | 0.085 | 0.212 | ***<0.001** |

Diff = difference in means; LCL = lower confidence level; UCL = upper confidence level; P = P-value, ≤ 0.008 in bold with asterisk

***Supplementary material 15****: Results from post hoc Tukey test of variation in residuals (representing bone size) between populations for the six bones (dentary, articular-angular, quadrate, premaxilla, maxilla and supramaxilla). AN-charr pooled as one population.*

| Bone | System | Pairs | Diff | LCL | UCL | P-value |
|---|---|---|---|---|---|---|
| Dentary | AN to lake morphs | Hest-AN | -0.045 | -0.111 | 0.021 | 0.448 |
| | | LB_TH-AN | -0.148 | -0.207 | -0.089 | ***<0.001** |
| | | LB_UL-AN | -0.174 | -0.246 | -0.102 | ***<0.001** |
| | | PI_TH-AN | 0.011 | -0.048 | 0.071 | 1.000 |
| | | PI_UL-AN | -0.093 | -0.195 | 0.008 | 0.100 |

| | | | | | | |
|---|---|---|---|---|---|---|
| | | PL_TH-AN | -0.072 | -0.128 | -0.017 | ***0.002** |
| | | PL_UL-AN | -0.183 | -0.252 | -0.114 | ***<0.001** |
| | | SB_TH-AN | -0.054 | -0.106 | -0.003 | 0.029 |
| | Hestvatn | LB_TH-Hest | -0.103 | -0.172 | -0.033 | ***<0.001** |
| | | LB_UL-Hest | -0.129 | -0.210 | -0.048 | ***<0.001** |
| | | PI_TH-Hest | 0.056 | -0.014 | 0.127 | 0.231 |
| | | PI_UL-Hest | -0.048 | -0.156 | 0.060 | 0.901 |
| | | PL_TH-Hest | -0.027 | -0.094 | 0.040 | 0.940 |
| | | PL_UL-Hest | -0.138 | -0.216 | -0.060 | ***<0.001** |
| | | SB_TH-Hest | -0.009 | -0.073 | 0.054 | 1.000 |
| | Þingvallavatn | PI_TH-LB_TH | 0.159 | 0.096 | 0.223 | ***<0.001** |
| | | PL_TH-LB_TH | 0.076 | 0.016 | 0.136 | ***0.003** |
| | | SB_TH-LB_TH | 0.093 | 0.037 | 0.150 | ***<0.001** |
| | | PL_TH-PI_TH | -0.084 | -0.144 | -0.023 | ***0.001** |
| | | SB_TH-PI_TH | -0.066 | -0.123 | -0.009 | 0.011 |
| | | SB_TH-PL_TH | 0.018 | -0.035 | 0.071 | 0.980 |
| | Úlfljótsvatn | PI_UL-LB_UL | 0.081 | -0.031 | 0.193 | 0.378 |
| | | PL_UL-LB_UL | -0.009 | -0.092 | 0.074 | 1.000 |
| | | PL_UL-PI_UL | -0.090 | -0.200 | 0.020 | 0.213 |
| | Across Þingvallavatn and Úlfljótsvatn | LB_UL-LB_TH | -0.026 | -0.102 | 0.049 | 0.977 |
| | | PI_UL-LB_TH | 0.055 | -0.049 | 0.158 | 0.784 |
| | | PL_UL-LB_TH | -0.035 | -0.107 | 0.037 | 0.846 |
| | | PI_TH-LB_UL | 0.185 | 0.109 | 0.261 | ***<0.001** |
| | | PL_TH-LB_UL | 0.102 | 0.029 | 0.175 | ***0.001** |
| | | SB_TH-LB_UL | 0.120 | 0.049 | 0.190 | ***<0.001** |
| | | PI_UL-PI_TH | -0.105 | -0.209 | 0.000 | 0.049 |
| | | PL_UL-PI_TH | -0.194 | -0.267 | -0.121 | ***<0.001** |
| | | PL_TH-PI_UL | 0.021 | -0.081 | 0.123 | 0.999 |
| | | SB_TH-PI_UL | 0.039 | -0.061 | 0.139 | 0.954 |
| | | PL_UL-PL_TH | -0.111 | -0.180 | -0.041 | ***<0.001** |
| | | SB_TH-PL_UL | 0.129 | 0.062 | 0.195 | ***<0.001** |
| Articular-angular | AN to lake morphs | Hest-AN | -0.055 | -0.111 | 0.000 | 0.048 |
| | | LB_TH-AN | -0.075 | -0.125 | -0.026 | ***<0.001** |
| | | LB_UL-AN | -0.078 | -0.138 | -0.017 | ***0.002** |
| | | PI_TH-AN | 0.010 | -0.039 | 0.060 | 0.999 |
| | | PI_UL-AN | -0.022 | -0.107 | 0.063 | 0.996 |
| | | PL_TH-AN | -0.066 | -0.113 | -0.019 | ***<0.001** |
| | | PL_UL-AN | -0.096 | -0.153 | -0.038 | ***<0.001** |
| | | SB_TH-AN | -0.005 | -0.048 | 0.039 | 1.000 |
| | Hestvatn | LB_TH-Hest | -0.020 | -0.078 | 0.038 | 0.979 |
| | | LB_UL-Hest | -0.022 | -0.090 | 0.046 | 0.984 |
| | | PI_TH-Hest | 0.066 | 0.007 | 0.124 | 0.015 |
| | | PI_UL-Hest | 0.033 | -0.057 | 0.124 | 0.968 |
| | | PL_TH-Hest | -0.010 | -0.067 | 0.046 | 1.000 |
| | | PL_UL-Hest | -0.040 | -0.106 | 0.025 | 0.596 |
| | | SB_TH-Hest | 0.051 | -0.003 | 0.104 | 0.076 |
| | Þingvallavatn | PI_TH-LB_TH | 0.086 | 0.032 | 0.139 | ***<0.001** |
| | | PL_TH-LB_TH | 0.009 | -0.041 | 0.060 | 1.000 |
| | | SB_TH-LB_TH | 0.071 | 0.023 | 0.118 | ***<0.001** |
| | | PL_TH-PI_TH | -0.076 | -0.127 | -0.025 | ***<0.001** |
| | | SB_TH-PI_TH | -0.015 | -0.062 | 0.033 | 0.988 |
| | | SB_TH-PL_TH | 0.061 | 0.017 | 0.106 | ***0.001** |
| | Úlfljótsvatn | PI_UL-LB_UL | 0.055 | -0.039 | 0.149 | 0.659 |
| | | PL_UL-LB_UL | -0.018 | -0.088 | 0.052 | 0.996 |
| | | PL_UL-PI_UL | -0.074 | -0.166 | 0.019 | 0.240 |
| | Across Þingvallavatn and Úlfljótsvatn | LB_UL-LB_TH | -0.002 | -0.066 | 0.061 | 1.000 |
| | | PI_UL-LB_TH | 0.053 | -0.034 | 0.140 | 0.616 |
| | | PL_UL-LB_TH | -0.021 | -0.081 | 0.040 | 0.980 |
| | | PI_TH-LB_UL | 0.088 | 0.024 | 0.151 | ***0.001** |
| | | PL_TH-LB_UL | 0.012 | -0.050 | 0.073 | 1.000 |
| | | SB_TH-LB_UL | 0.073 | 0.014 | 0.132 | ***0.004** |

| | | | | | | |
|---|---|---|---|---|---|---|
| | | PI_UL-PI_TH | -0.033 | -0.120 | 0.055 | 0.964 |
| | | PL_UL-PI_TH | -0.106 | -0.167 | -0.045 | ***<0.001** |
| | | PL_TH-PI_UL | -0.044 | -0.129 | 0.042 | 0.813 |
| | | SB_TH-PI_UL | 0.018 | -0.066 | 0.102 | 0.999 |
| | | PL_UL-PL_TH | -0.030 | -0.089 | 0.029 | 0.807 |
| | | SB_TH-PL_UL | 0.091 | 0.035 | 0.147 | ***<0.001** |
| Quadrate | AN to lake morphs | Hest-AN | -0.067 | -0.120 | -0.015 | ***0.003** |
| | | LB_TH-AN | 0.019 | -0.028 | 0.066 | 0.937 |
| | | LB_UL-AN | -0.068 | -0.126 | -0.010 | 0.008 |
| | | PI_TH-AN | 0.027 | -0.020 | 0.075 | 0.687 |
| | | PI_UL-AN | -0.047 | -0.128 | 0.034 | 0.674 |
| | | PL_TH-AN | -0.063 | -0.108 | -0.019 | ***<0.001** |
| | | PL_UL-AN | -0.090 | -0.145 | -0.035 | ***<0.001** |
| | | SB_TH-AN | 0.052 | 0.011 | 0.094 | ***0.003** |
| | Hestvatn | LB_TH-Hest | 0.087 | 0.031 | 0.142 | ***<0.001** |
| | | LB_UL-Hest | -0.001 | -0.066 | 0.064 | 1.000 |
| | | PI_TH-Hest | 0.095 | 0.039 | 0.150 | ***<0.001** |
| | | PI_UL-Hest | 0.020 | -0.066 | 0.107 | 0.998 |
| | | PL_TH-Hest | 0.004 | -0.050 | 0.058 | 1.000 |
| | | PL_UL-Hest | -0.023 | -0.085 | 0.039 | 0.966 |
| | | SB_TH-Hest | 0.120 | 0.069 | 0.171 | ***<0.001** |
| | Þingvallavatn | PI_TH-LB_TH | 0.008 | -0.043 | 0.059 | 1.000 |
| | | PL_TH-LB_TH | -0.083 | -0.131 | -0.035 | ***<0.001** |
| | | SB_TH-LB_TH | 0.033 | -0.012 | 0.078 | 0.355 |
| | | PL_TH-PI_TH | -0.091 | -0.139 | -0.042 | ***<0.001** |
| | | SB_TH-PI_TH | 0.025 | -0.020 | 0.071 | 0.731 |
| | | SB_TH-PL_TH | 0.116 | 0.073 | 0.159 | ***<0.001** |
| | Úlfljótsvatn | PI_UL-LB_UL | 0.021 | -0.069 | 0.111 | 0.998 |
| | | PL_UL-LB_UL | -0.022 | -0.089 | 0.044 | 0.982 |
| | | PL_UL-PI_UL | -0.043 | -0.131 | 0.045 | 0.838 |
| | Across Þingvallavatn and Úlfljótsvatn | LB_UL-LB_TH | -0.087 | -0.148 | -0.027 | ***<0.001** |
| | | PI_UL-LB_TH | -0.066 | -0.150 | 0.017 | 0.237 |
| | | PL_UL-LB_TH | -0.110 | -0.167 | -0.052 | ***<0.001** |
| | | PI_TH-LB_UL | 0.095 | 0.035 | 0.156 | ***<0.001** |
| | | PL_TH-LB_UL | 0.005 | -0.054 | 0.063 | 1.000 |
| | | SB_TH-LB_UL | 0.121 | 0.064 | 0.177 | ***<0.001** |
| | | PI_UL-PI_TH | -0.074 | -0.158 | 0.009 | 0.122 |
| | | PL_UL-PI_TH | -0.118 | -0.176 | -0.060 | ***<0.001** |
| | | PL_TH-PI_UL | -0.016 | -0.098 | 0.066 | 1.000 |
| | | SB_TH-PI_UL | 0.100 | 0.019 | 0.180 | ***0.004** |
| | | PL_UL-PL_TH | -0.027 | -0.083 | 0.029 | 0.850 |
| | | SB_TH-PL_UL | 0.143 | 0.089 | 0.196 | ***<0.001** |
| Premaxilla | AN to lake morphs | Hest-AN | -0.104 | -0.177 | -0.031 | ***<0.001** |
| | | LB_TH-AN | -0.076 | -0.141 | -0.011 | 0.009 |
| | | LB_UL-AN | -0.099 | -0.179 | -0.019 | ***0.004** |
| | | PI_TH-AN | -0.025 | -0.090 | 0.041 | 0.962 |
| | | PI_UL-AN | -0.131 | -0.243 | -0.019 | 0.010 |
| | | PL_TH-AN | -0.185 | -0.246 | -0.124 | ***<0.001** |
| | | PL_UL-AN | -0.145 | -0.221 | -0.069 | ***<0.001** |
| | | SB_TH-AN | 0.036 | -0.021 | 0.094 | 0.548 |
| | Hestvatn | LB_TH-Hest | 0.028 | -0.049 | 0.105 | 0.970 |
| | | LB_UL-Hest | 0.005 | -0.084 | 0.095 | 1.000 |
| | | PI_TH-Hest | 0.080 | 0.002 | 0.157 | 0.038 |
| | | PI_UL-Hest | -0.027 | -0.146 | 0.093 | 0.999 |
| | | PL_TH-Hest | -0.081 | -0.155 | -0.007 | 0.020 |
| | | PL_UL-Hest | -0.041 | -0.127 | 0.045 | 0.865 |
| | | SB_TH-Hest | 0.141 | 0.070 | 0.211 | ***<0.001** |
| | Þingvallavatn | PI_TH-LB_TH | 0.052 | -0.018 | 0.122 | 0.341 |
| | | PL_TH-LB_TH | -0.109 | -0.175 | -0.043 | ***<0.001** |
| | | SB_TH-LB_TH | 0.113 | 0.051 | 0.175 | ***<0.001** |
| | | PL_TH-PI_TH | -0.161 | -0.227 | -0.094 | ***<0.001** |

| | | | | | | |
|---|---|---|---|---|---|---|
| | | SB_TH-PI_TH | 0.061 | -0.002 | 0.124 | 0.063 |
| | | SB_TH-PL_TH | 0.222 | 0.163 | 0.280 | ***<0.001** |
| | Úlfljótsvatn | PI_UL-LB_UL | -0.032 | -0.156 | 0.092 | 0.997 |
| | | PL_UL-LB_UL | -0.046 | -0.139 | 0.046 | 0.824 |
| | | PL_UL-PI_UL | -0.014 | -0.136 | 0.107 | 1.000 |
| | Across Þingvallavatn and Úlfljótsvatn | LB_UL-LB_TH | -0.022 | -0.106 | 0.061 | 0.996 |
| | | PI_UL-LB_TH | -0.054 | -0.170 | 0.061 | 0.866 |
| | | PL_UL-LB_TH | -0.069 | -0.149 | 0.011 | 0.158 |
| | | PI_TH-LB_UL | 0.074 | -0.010 | 0.158 | 0.132 |
| | | PL_TH-LB_UL | -0.086 | -0.167 | -0.006 | 0.026 |
| | | SB_TH-LB_UL | 0.135 | 0.058 | 0.213 | ***<0.001** |
| | | PI_UL-PI_TH | -0.106 | -0.222 | 0.009 | 0.098 |
| | | PL_UL-PI_TH | -0.121 | -0.201 | -0.040 | ***<0.001** |
| | | PL_TH-PI_UL | -0.054 | -0.167 | 0.059 | 0.856 |
| | | SB_TH-PI_UL | 0.167 | 0.057 | 0.278 | ***<0.001** |
| | | PL_UL-PL_TH | 0.040 | -0.037 | 0.117 | 0.793 |
| | | SB_TH-PL_UL | 0.182 | 0.108 | 0.255 | ***<0.001** |
| Maxilla | AN to lake morphs | Hest-AN | -0.046 | -0.111 | 0.019 | 0.415 |
| | | LB_TH-AN | -0.038 | -0.097 | 0.020 | 0.519 |
| | | LB_UL-AN | -0.102 | -0.174 | -0.031 | ***<0.001** |
| | | PI_TH-AN | 0.047 | -0.011 | 0.106 | 0.229 |
| | | PI_UL-AN | -0.052 | -0.153 | 0.049 | 0.805 |
| | | PL_TH-AN | -0.044 | -0.100 | 0.011 | 0.232 |
| | | PL_UL-AN | -0.154 | -0.222 | -0.085 | ***<0.001** |
| | | SB_TH-AN | 0.020 | -0.032 | 0.072 | 0.954 |
| | Hestvatn | LB_TH-Hest | 0.008 | -0.061 | 0.077 | 1.000 |
| | | LB_UL-Hest | -0.056 | -0.137 | 0.024 | 0.421 |
| | | PI_TH-Hest | 0.093 | 0.024 | 0.163 | ***0.001** |
| | | PI_UL-Hest | -0.006 | -0.113 | 0.102 | 1.000 |
| | | PL_TH-Hest | 0.002 | -0.065 | 0.068 | 1.000 |
| | | PL_UL-Hest | -0.108 | -0.185 | -0.030 | ***0.001** |
| | | SB_TH-Hest | 0.066 | 0.002 | 0.129 | 0.035 |
| | Þingvallavatn | PI_TH-LB_TH | 0.085 | 0.023 | 0.148 | ***0.001** |
| | | PL_TH-LB_TH | -0.006 | -0.066 | 0.053 | 1.000 |
| | | SB_TH-LB_TH | 0.058 | 0.002 | 0.114 | 0.036 |
| | | PL_TH-PI_TH | -0.092 | -0.152 | -0.032 | ***<0.001** |
| | | SB_TH-PI_TH | -0.027 | -0.084 | 0.029 | 0.851 |
| | | SB_TH-PL_TH | 0.064 | 0.011 | 0.117 | ***0.005** |
| | Úlfljótsvatn | PI_UL-LB_UL | 0.051 | -0.061 | 0.162 | 0.891 |
| | | PL_UL-LB_UL | -0.051 | -0.134 | 0.032 | 0.593 |
| | | PL_UL-PI_UL | -0.102 | -0.211 | 0.007 | 0.089 |
| | Across Þingvallavatn and Úlfljótsvatn | LB_UL-LB_TH | -0.064 | -0.139 | 0.011 | 0.165 |
| | | PI_UL-LB_TH | -0.014 | -0.117 | 0.090 | 1.000 |
| | | PL_UL-LB_TH | -0.116 | -0.187 | -0.044 | ***<0.001** |
| | | PI_TH-LB_UL | 0.150 | 0.074 | 0.225 | ***<0.001** |
| | | PL_TH-LB_UL | 0.058 | -0.015 | 0.131 | 0.244 |
| | | SB_TH-LB_UL | 0.122 | 0.052 | 0.192 | ***<0.001** |
| | | PI_UL-PI_TH | -0.099 | -0.203 | 0.004 | 0.073 |
| | | PL_UL-PI_TH | -0.201 | -0.273 | -0.129 | ***<0.001** |
| | | PL_TH-PI_UL | 0.007 | -0.094 | 0.109 | 1.000 |
| | | SB_TH-PI_UL | 0.072 | -0.028 | 0.171 | 0.378 |
| | | PL_UL-PL_TH | -0.109 | -0.179 | -0.040 | ***<0.001** |
| | | SB_TH-PL_UL | 0.174 | 0.107 | 0.240 | ***<0.001** |
| Supramaxilla | AN to lake morphs | Hest-AN | -0.107 | -0.170 | -0.044 | ***<0.001** |
| | | LB_TH-AN | -0.057 | -0.113 | -0.001 | 0.043 |
| | | LB_UL-AN | -0.126 | -0.195 | -0.057 | ***<0.001** |
| | | PI_TH-AN | 0.028 | -0.028 | 0.084 | 0.831 |
| | | PI_UL-AN | -0.073 | -0.170 | 0.023 | 0.309 |
| | | PL_TH-AN | -0.078 | -0.131 | -0.024 | ***<0.001** |
| | | PL_UL-AN | -0.139 | -0.204 | -0.073 | ***<0.001** |
| | | SB_TH-AN | 0.009 | -0.040 | 0.059 | 1.000 |

| | | | | | | |
|---|---|---|---|---|---|---|
| | Hestvatn | LB_TH-Hest | 0.050 | -0.016 | 0.116 | 0.306 |
| | | LB_UL-Hest | -0.019 | -0.096 | 0.058 | 0.998 |
| | | PI_TH-Hest | 0.135 | 0.069 | 0.201 | ***<0.001** |
| | | PI_UL-Hest | 0.034 | -0.069 | 0.137 | 0.983 |
| | | PL_TH-Hest | 0.029 | -0.035 | 0.094 | 0.885 |
| | | PL_UL-Hest | -0.032 | -0.106 | 0.042 | 0.920 |
| | | SB_TH-Hest | 0.116 | 0.056 | 0.177 | ***<0.001** |
| | Þingvallavatn | PI_TH-LB_TH | 0.085 | 0.025 | 0.145 | ***<0.001** |
| | | PL_TH-LB_TH | -0.021 | -0.078 | 0.037 | 0.971 |
| | | SB_TH-LB_TH | 0.066 | 0.012 | 0.120 | ***0.005** |
| | | PL_TH-PI_TH | -0.105 | -0.163 | -0.048 | ***<0.001** |
| | | SB_TH-PI_TH | -0.019 | -0.073 | 0.036 | 0.979 |
| | | SB_TH-PL_TH | 0.087 | 0.035 | 0.139 | ***<0.001** |
| | Úlfljótsvatn | PI_UL-LB_UL | 0.053 | -0.054 | 0.159 | 0.832 |
| | | PL_UL-LB_UL | -0.013 | -0.092 | 0.066 | 1.000 |
| | | PL_UL-PI_UL | -0.066 | -0.170 | 0.039 | 0.570 |
| | Across Þingvallavatn and Úlfljótsvatn | LB_UL-LB_TH | -0.069 | -0.141 | 0.003 | 0.071 |
| | | PI_UL-LB_TH | -0.016 | -0.115 | 0.083 | 1.000 |
| | | PL_UL-LB_TH | -0.082 | -0.150 | -0.013 | ***0.007** |
| | | PI_TH-LB_UL | 0.154 | 0.082 | 0.226 | ***<0.001** |
| | | PL_TH-LB_UL | 0.048 | -0.022 | 0.118 | 0.438 |
| | | SB_TH-LB_UL | 0.135 | 0.068 | 0.202 | ***<0.001** |
| | | PI_UL-PI_TH | -0.101 | -0.200 | -0.002 | 0.041 |
| | | PL_UL-PI_TH | -0.167 | -0.236 | -0.098 | ***<0.001** |
| | | PL_TH-PI_UL | -0.004 | -0.102 | 0.093 | 1.000 |
| | | SB_TH-PI_UL | 0.082 | -0.013 | 0.178 | 0.152 |
| | | PL_UL-PL_TH | -0.061 | -0.128 | 0.006 | 0.102 |
| | | SB_TH-PL_UL | 0.148 | 0.085 | 0.212 | ***<0.001** |

Diff = difference in means; LCL = lower confidence level; UCL = upper confidence level; P = P-value, ≤ 0.008 in bold with asterisk

***Supplementary material 16**: Results from pairwise tests of shape by population, for differences between absolute vector lengths (D) used to examine possible shape differences by population in five bones. These bones had significant shape differences according to ANOVA model with main effects and interaction of population by bone size. Used to verify results from ANOVA and test for true allometry (i.e., different group slopes). AN-charr treated as five separate populations.*

| Bone | System | Pairs | D-value | UCL | Z | P-value |
|---|---|---|---|---|---|---|
| Dentary | AN populations | AN_Brúará:AN_Gufudalsá | 0.110 | 0.297 | -2.33 | 0.992 |
| | | AN_Brúará:AN_Lambeyrarkvísl | 0.202 | 0.332 | -0.52 | 0.710 |
| | | AN_Brúará:AN_Víðidalsá | 0.218 | 0.298 | 0.51 | 0.307 |
| | | AN_Brúará:AN_Vífilsdalsá | 0.158 | 0.328 | -1.33 | 0.903 |
| | | AN_Gufudalsá:AN_Lambeyrarkvísl | 0.255 | 0.244 | 1.86 | 0.033 |
| | | AN_Gufudalsá:AN_Víðidalsá | 0.259 | 0.155 | 3.96 | ***0.001** |
| | | AN_Gufudalsá:AN_Vífilsdalsá | 0.208 | 0.226 | 1.26 | 0.106 |
| | | AN_Lambeyrarkvísl:AN_Víðidalsá | 0.231 | 0.235 | 1.57 | 0.058 |
| | | AN_Lambeyrarkvísl:AN_Vífilsdalsá | 0.164 | 0.284 | -0.64 | 0.730 |
| | | AN_Víðidalsá:AN_Vífilsdalsá | 0.150 | 0.219 | 0.12 | 0.448 |
| | Brúará to lake morphs | AN_Brúará:Hest | 0.139 | 0.282 | -1.15 | 0.879 |
| | | AN_Brúará:Hit | 0.153 | 0.341 | -1.51 | 0.937 |
| | | AN_Brúará:LB_TH | 0.109 | 0.288 | -2.05 | 0.981 |
| | | AN_Brúará:LB_UL | 0.139 | 0.287 | -1.20 | 0.887 |
| | | AN_Brúará:PI_TH | 0.174 | 0.277 | -0.19 | 0.579 |
| | | AN_Brúará:PI_UL | 0.166 | 0.309 | -0.91 | 0.824 |
| | | AN_Brúará:PL_TH | 0.108 | 0.286 | -2.09 | 0.981 |
| | | AN_Brúará:PL_UL | 0.120 | 0.297 | -1.85 | 0.970 |
| | | AN_Brúará:SB_TH | 0.141 | 0.280 | -1.02 | 0.862 |

| | | | | | | |
|---|---|---|---|---|---|---|
| | Gufudalsá to lake morphs | AN_Gufudalsá:Hest | 0.152 | 0.132 | 2.29 | 0.010 |
| | | AN_Gufudalsá:Hit | 0.147 | 0.232 | -0.30 | 0.621 |
| | | AN_Gufudalsá:LB_TH | 0.126 | 0.133 | 1.43 | 0.080 |
| | | AN_Gufudalsá:LB_UL | 0.166 | 0.142 | 2.46 | ***0.004** |
| | | AN_Gufudalsá:PI_TH | 0.215 | 0.126 | 4.20 | ***0.001** |
| | | AN_Gufudalsá:PI_UL | 0.182 | 0.200 | 1.21 | 0.113 |
| | | AN_Gufudalsá:PL_TH | 0.143 | 0.132 | 1.99 | 0.023 |
| | | AN_Gufudalsá:PL_UL | 0.121 | 0.155 | 0.59 | 0.285 |
| | | AN_Gufudalsá:SB_TH | 0.181 | 0.128 | 3.34 | ***0.001** |
| | Lambeyrarkvísl to lake morphs | AN_Lambeyrarkvísl:Hest | 0.156 | 0.218 | 0.19 | 0.423 |
| | | AN_Lambeyrarkvísl:Hit | 0.203 | 0.288 | 0.17 | 0.443 |
| | | AN_Lambeyrarkvísl:LB_TH | 0.200 | 0.223 | 1.21 | 0.127 |
| | | AN_Lambeyrarkvísl:LB_UL | 0.144 | 0.229 | -0.19 | 0.582 |
| | | AN_Lambeyrarkvísl:PI_TH | 0.152 | 0.224 | 0.18 | 0.426 |
| | | AN_Lambeyrarkvísl:PI_UL | 0.174 | 0.260 | -0.13 | 0.541 |
| | | AN_Lambeyrarkvísl:PL_TH | 0.146 | 0.225 | -0.01 | 0.502 |
| | | AN_Lambeyrarkvísl:PL_UL | 0.193 | 0.233 | 0.85 | 0.198 |
| | | AN_Lambeyrarkvísl:SB_TH | 0.116 | 0.219 | -0.92 | 0.817 |
| | Víðidalsá to lake morphs | AN_Víðidalsá:Hest | 0.194 | 0.116 | 3.88 | ***0.001** |
| | | AN_Víðidalsá:Hit | 0.282 | 0.225 | 2.48 | 0.011 |
| | | AN_Víðidalsá:LB_TH | 0.181 | 0.118 | 3.81 | ***0.001** |
| | | AN_Víðidalsá:LB_UL | 0.192 | 0.129 | 3.31 | ***0.001** |
| | | AN_Víðidalsá:PI_TH | 0.168 | 0.113 | 3.29 | ***0.001** |
| | | AN_Víðidalsá:PI_UL | 0.210 | 0.188 | 2.22 | 0.018 |
| | | AN_Víðidalsá:PL_TH | 0.215 | 0.116 | 4.65 | ***0.001** |
| | | AN_Víðidalsá:PL_UL | 0.204 | 0.145 | 3.09 | ***0.001** |
| | | AN_Víðidalsá:SB_TH | 0.212 | 0.112 | 4.49 | ***0.001** |
| | Vífisdalsá to lake morphs | AN_Vífilsdalsá:Hest | 0.143 | 0.203 | 0.23 | 0.410 |
| | | AN_Vífilsdalsá:Hit | 0.192 | 0.271 | 0.16 | 0.439 |
| | | AN_Vífilsdalsá:LB_TH | 0.135 | 0.205 | 0.03 | 0.485 |
| | | AN_Vífilsdalsá:LB_UL | 0.132 | 0.210 | -0.18 | 0.576 |
| | | AN_Vífilsdalsá:PI_TH | 0.160 | 0.198 | 0.73 | 0.246 |
| | | AN_Vífilsdalsá:PI_UL | 0.181 | 0.245 | 0.35 | 0.372 |
| | | AN_Vífilsdalsá:PL_TH | 0.147 | 0.203 | 0.35 | 0.376 |
| | | AN_Vífilsdalsá:PL_UL | 0.140 | 0.220 | -0.19 | 0.577 |
| | | AN_Vífilsdalsá:SB_TH | 0.147 | 0.198 | 0.42 | 0.334 |
| | Hestvatn and Hítarvatn | Hest:Hit | 0.118 | 0.217 | -0.67 | 0.751 |
| | | Hest:LB_TH | 0.099 | 0.087 | 2.25 | 0.009 |
| | | Hest:LB_UL | 0.062 | 0.098 | -0.28 | 0.610 |
| | | Hest:PI_TH | 0.103 | 0.079 | 2.85 | ***0.003** |
| | | Hest:PI_UL | 0.101 | 0.171 | -0.57 | 0.712 |
| | | Hest:PL_TH | 0.070 | 0.084 | 0.82 | 0.203 |
| | | Hest:PL_UL | 0.086 | 0.117 | 0.28 | 0.384 |
| | | Hest:SB_TH | 0.077 | 0.077 | 1.78 | 0.042 |
| | | Hit:LB_TH | 0.154 | 0.216 | 0.40 | 0.341 |
| | | Hit:LB_UL | 0.130 | 0.216 | -0.41 | 0.656 |
| | | Hit:PI_TH | 0.189 | 0.213 | 1.22 | 0.104 |
| | | Hit:PI_UL | 0.154 | 0.256 | -0.48 | 0.688 |
| | | Hit:PL_TH | 0.105 | 0.210 | -1.09 | 0.867 |
| | | Hit:PL_UL | 0.103 | 0.226 | -1.50 | 0.946 |
| | | Hit:SB_TH | 0.134 | 0.207 | -0.09 | 0.538 |
| | Þingvallavatn | LB_TH:PI_TH | 0.157 | 0.079 | 4.68 | ***0.001** |
| | | LB_TH:PL_TH | 0.107 | 0.083 | 2.77 | ***0.004** |
| | | LB_TH:SB_TH | 0.135 | 0.077 | 4.13 | ***0.001** |
| | | PI_TH:PL_TH | 0.111 | 0.078 | 3.17 | ***0.003** |
| | | PI_TH:SB_TH | 0.089 | 0.069 | 2.70 | ***0.006** |
| | | PL_TH:SB_TH | 0.052 | 0.074 | 0.13 | 0.457 |
| | Úlfljótsvatn | LB_UL:PI_UL | 0.107 | 0.181 | -0.50 | 0.690 |
| | | LB_UL:PL_UL | 0.093 | 0.127 | 0.32 | 0.367 |
| | | PI_UL:PL_UL | 0.138 | 0.190 | 0.25 | 0.403 |
| | | LB_TH:LB_UL | 0.109 | 0.098 | 2.18 | 0.011 |

| | | | | | | |
|---|---|---|---|---|---|---|
| | Across Þingvallavatn and Úlfljótsvatn | LB_TH:PI_UL | 0.135 | 0.173 | 0.64 | 0.267 |
| | | LB_TH:PL_UL | 0.095 | 0.116 | 0.74 | 0.219 |
| | | LB_UL:PI_TH | 0.096 | 0.097 | 1.74 | 0.052 |
| | | LB_UL:PL_TH | 0.069 | 0.093 | 0.27 | 0.387 |
| | | LB_UL:SB_TH | 0.069 | 0.090 | 0.48 | 0.317 |
| | | PI_TH:PI_UL | 0.119 | 0.169 | 0.20 | 0.429 |
| | | PI_TH:PL_UL | 0.146 | 0.114 | 2.73 | ***0.005** |
| | | PI_UL:PL_TH | 0.106 | 0.170 | -0.35 | 0.634 |
| | | PI_UL:SB_TH | 0.102 | 0.168 | -0.39 | 0.660 |
| | | PL_TH:PL_UL | 0.088 | 0.118 | 0.42 | 0.341 |
| | | PL_UL:SB_TH | 0.118 | 0.112 | 1.85 | 0.031 |
| Articular-angular | AN populations | AN_Brúará:AN_Gufudalsá | 0.236 | 0.328 | 0.36 | 0.351 |
| | | AN_Brúará:AN_Lambeyrarkvísl | 0.364 | 0.387 | 1.37 | 0.082 |
| | | AN_Brúará:AN_Víðidalsá | 0.304 | 0.322 | 1.39 | 0.079 |
| | | AN_Brúará:AN_Vífilsdalsá | 0.267 | 0.376 | 0.34 | 0.367 |
| | | AN_Gufudalsá:AN_Lambeyrarkvísl | 0.301 | 0.295 | 1.70 | 0.041 |
| | | AN_Gufudalsá:AN_Víðidalsá | 0.144 | 0.182 | 0.62 | 0.264 |
| | | AN_Gufudalsá:AN_Vífilsdalsá | 0.119 | 0.264 | -1.55 | 0.939 |
| | | AN_Lambeyrarkvísl:AN_Víðidalsá | 0.316 | 0.286 | 1.98 | 0.027 |
| | | AN_Lambeyrarkvísl:AN_Vífilsdalsá | 0.274 | 0.346 | 0.71 | 0.226 |
| | | AN_Víðidalsá:AN_Vífilsdalsá | 0.109 | 0.261 | -1.83 | 0.972 |
| | Brúará to lake morphs | AN_Brúará:Hest | 0.230 | 0.299 | 0.57 | 0.276 |
| | | AN_Brúará:Hit | 0.349 | 0.372 | 1.39 | 0.091 |
| | | AN_Brúará:LB_TH | 0.269 | 0.299 | 1.15 | 0.115 |
| | | AN_Brúará:LB_UL | 0.258 | 0.304 | 0.94 | 0.171 |
| | | AN_Brúará:PI_TH | 0.280 | 0.302 | 1.33 | 0.083 |
| | | AN_Brúará:PI_UL | 0.260 | 0.344 | 0.54 | 0.298 |
| | | AN_Brúará:PL_TH | 0.294 | 0.301 | 1.49 | 0.060 |
| | | AN_Brúará:PL_UL | 0.210 | 0.313 | 0.02 | 0.496 |
| | | AN_Brúará:SB_TH | 0.276 | 0.296 | 1.29 | 0.101 |
| | Gufudalsá to lake morphs | AN_Gufudalsá:Hest | 0.084 | 0.150 | -0.73 | 0.763 |
| | | AN_Gufudalsá:Hit | 0.220 | 0.262 | 0.92 | 0.181 |
| | | AN_Gufudalsá:LB_TH | 0.074 | 0.150 | -1.23 | 0.887 |
| | | AN_Gufudalsá:LB_UL | 0.119 | 0.153 | 0.45 | 0.332 |
| | | AN_Gufudalsá:PI_TH | 0.108 | 0.143 | 0.41 | 0.352 |
| | | AN_Gufudalsá:PI_UL | 0.146 | 0.216 | -0.09 | 0.531 |
| | | AN_Gufudalsá:PL_TH | 0.148 | 0.149 | 1.62 | 0.052 |
| | | AN_Gufudalsá:PL_UL | 0.105 | 0.174 | -0.53 | 0.702 |
| | | AN_Gufudalsá:SB_TH | 0.135 | 0.143 | 1.38 | 0.081 |
| | Lambeyrarkvísl to lake morphs | AN_Lambeyrarkvísl:Hest | 0.259 | 0.270 | 1.49 | 0.068 |
| | | AN_Lambeyrarkvísl:Hit | 0.422 | 0.349 | 2.53 | ***0.007** |
| | | AN_Lambeyrarkvísl:LB_TH | 0.292 | 0.268 | 1.94 | 0.033 |
| | | AN_Lambeyrarkvísl:LB_UL | 0.218 | 0.281 | 0.71 | 0.237 |
| | | AN_Lambeyrarkvísl:PI_TH | 0.244 | 0.269 | 1.28 | 0.094 |
| | | AN_Lambeyrarkvísl:PI_UL | 0.236 | 0.312 | 0.54 | 0.296 |
| | | AN_Lambeyrarkvísl:PL_TH | 0.212 | 0.269 | 0.72 | 0.245 |
| | | AN_Lambeyrarkvísl:PL_UL | 0.260 | 0.284 | 1.28 | 0.101 |
| | | AN_Lambeyrarkvísl:SB_TH | 0.209 | 0.268 | 0.71 | 0.240 |
| | Víðidalsá to lake morphs | AN_Víðidalsá:Hest | 0.124 | 0.137 | 1.14 | 0.133 |
| | | AN_Víðidalsá:Hit | 0.189 | 0.252 | 0.42 | 0.337 |
| | | AN_Víðidalsá:LB_TH | 0.107 | 0.142 | 0.44 | 0.330 |
| | | AN_Víðidalsá:LB_UL | 0.167 | 0.153 | 2.03 | 0.016 |
| | | AN_Víðidalsá:PI_TH | 0.153 | 0.137 | 2.17 | 0.017 |
| | | AN_Víðidalsá:PI_UL | 0.155 | 0.211 | 0.32 | 0.370 |
| | | AN_Víðidalsá:PL_TH | 0.158 | 0.140 | 2.15 | 0.016 |
| | | AN_Víðidalsá:PL_UL | 0.143 | 0.168 | 0.91 | 0.186 |
| | | AN_Víðidalsá:SB_TH | 0.167 | 0.132 | 2.60 | ***0.004** |
| | Vífisdalsá to lake morphs | AN_Vífilsdalsá:Hest | 0.075 | 0.238 | -2.88 | 0.999 |
| | | AN_Vífilsdalsá:Hit | 0.229 | 0.315 | 0.27 | 0.395 |
| | | AN_Vífilsdalsá:LB_TH | 0.087 | 0.239 | -2.27 | 0.989 |
| | | AN_Vífilsdalsá:LB_UL | 0.101 | 0.247 | -1.84 | 0.969 |

| | | | | | | |
|---|---|---|---|---|---|---|
| | | AN_Vífilsdalsá:PI_TH | 0.116 | 0.238 | -1.16 | 0.875 |
| | | AN_Vífilsdalsá:PI_UL | 0.147 | 0.293 | -1.01 | 0.852 |
| | | AN_Vífilsdalsá:PL_TH | 0.105 | 0.241 | -1.55 | 0.935 |
| | | AN_Vífilsdalsá:PL_UL | 0.093 | 0.261 | -2.22 | 0.986 |
| | | AN_Vífilsdalsá:SB_TH | 0.115 | 0.238 | -1.13 | 0.864 |
| | Hestvatn and Hítarvatn | Hest:Hit | 0.225 | 0.233 | 1.50 | 0.071 |
| | | Hest:LB_TH | 0.086 | 0.090 | 1.40 | 0.077 |
| | | Hest:LB_UL | 0.090 | 0.106 | 1.00 | 0.160 |
| | | Hest:PI_TH | 0.077 | 0.087 | 1.17 | 0.121 |
| | | Hest:PI_UL | 0.112 | 0.183 | -0.40 | 0.666 |
| | | Hest:PL_TH | 0.098 | 0.092 | 1.99 | 0.019 |
| | | Hest:PL_UL | 0.071 | 0.130 | -0.90 | 0.822 |
| | | Hest:SB_TH | 0.091 | 0.080 | 2.20 | 0.014 |
| | | Hit:LB_TH | 0.212 | 0.233 | 1.24 | 0.111 |
| | | Hit:LB_UL | 0.281 | 0.243 | 2.33 | 0.008 |
| | | Hit:PI_TH | 0.223 | 0.232 | 1.49 | 0.070 |
| | | Hit:PI_UL | 0.278 | 0.279 | 1.64 | 0.052 |
| | | Hit:PL_TH | 0.249 | 0.234 | 1.91 | 0.027 |
| | | Hit:PL_UL | 0.236 | 0.248 | 1.42 | 0.090 |
| | | Hit:SB_TH | 0.246 | 0.226 | 1.95 | 0.028 |
| | Þingvallavatn | LB_TH:PI_TH | 0.109 | 0.084 | 2.85 | ***0.002** |
| | | LB_TH:PL_TH | 0.136 | 0.091 | 3.52 | ***0.001** |
| | | LB_TH:SB_TH | 0.136 | 0.081 | 4.03 | ***0.001** |
| | | PI_TH:PL_TH | 0.079 | 0.083 | 1.47 | 0.072 |
| | | PI_TH:SB_TH | 0.066 | 0.074 | 1.19 | 0.127 |
| | | PL_TH:SB_TH | 0.042 | 0.080 | -1.12 | 0.865 |
| | Úlfljótsvatn | LB_UL:PI_UL | 0.113 | 0.192 | -0.54 | 0.706 |
| | | LB_UL:PL_UL | 0.106 | 0.143 | 0.46 | 0.320 |
| | | PI_UL:PL_UL | 0.138 | 0.204 | -0.05 | 0.511 |
| | Across Þingvallavatn and Úlfljótsvatn | LB_TH:LB_UL | 0.117 | 0.108 | 1.96 | 0.027 |
| | | LB_TH:PI_UL | 0.149 | 0.183 | 0.75 | 0.215 |
| | | LB_TH:PL_UL | 0.114 | 0.130 | 1.05 | 0.153 |
| | | LB_UL:PI_TH | 0.104 | 0.099 | 1.75 | 0.037 |
| | | LB_UL:PL_TH | 0.092 | 0.103 | 1.11 | 0.144 |
| | | LB_UL:SB_TH | 0.090 | 0.098 | 1.31 | 0.098 |
| | | PI_TH:PI_UL | 0.127 | 0.180 | 0.21 | 0.411 |
| | | PI_TH:PL_UL | 0.122 | 0.128 | 1.45 | 0.065 |
| | | PI_UL:PL_TH | 0.141 | 0.184 | 0.54 | 0.302 |
| | | PI_UL:SB_TH | 0.130 | 0.176 | 0.31 | 0.377 |
| | | PL_TH:PL_UL | 0.118 | 0.131 | 1.24 | 0.109 |
| | | PL_UL:SB_TH | 0.106 | 0.126 | 0.94 | 0.184 |
| Quadrate | AN populations | AN_Brúará:AN_Gufudalsá | 0.446 | 0.780 | -0.19 | 0.580 |
| | | AN_Brúará:AN_Lambeyrarkvísl | 0.474 | 0.800 | -0.09 | 0.531 |
| | | AN_Brúará:AN_Víðidalsá | 0.432 | 0.771 | -0.29 | 0.621 |
| | | AN_Brúará:AN_Vífilsdalsá | 0.604 | 0.797 | 0.83 | 0.198 |
| | | AN_Gufudalsá:AN_Lambeyrarkvísl | 0.171 | 0.307 | -0.58 | 0.707 |
| | | AN_Gufudalsá:AN_Víðidalsá | 0.174 | 0.238 | 0.39 | 0.342 |
| | | AN_Gufudalsá:AN_Vífilsdalsá | 0.284 | 0.264 | 1.92 | 0.024 |
| | | AN_Lambeyrarkvísl:AN_Víðidalsá | 0.232 | 0.322 | 0.48 | 0.292 |
| | | AN_Lambeyrarkvísl:AN_Vífilsdalsá | 0.316 | 0.333 | 1.42 | 0.072 |
| | | AN_Víðidalsá:AN_Vífilsdalsá | 0.389 | 0.269 | 2.94 | ***0.002** |
| | Brúará to lake morphs | AN_Brúará:Hest | 0.435 | 0.767 | -0.19 | 0.579 |
| | | AN_Brúará:Hit | 0.361 | 0.828 | -1.23 | 0.893 |
| | | AN_Brúará:LB_TH | 0.431 | 0.780 | -0.23 | 0.605 |
| | | AN_Brúará:LB_UL | 0.453 | 0.775 | -0.05 | 0.516 |
| | | AN_Brúará:PI_TH | 0.477 | 0.765 | 0.14 | 0.437 |
| | | AN_Brúará:PI_UL | 0.372 | 0.798 | -0.98 | 0.847 |
| | | AN_Brúará:PL_TH | 0.476 | 0.767 | 0.13 | 0.439 |
| | | AN_Brúará:PL_UL | 0.501 | 0.757 | 0.26 | 0.385 |
| | | AN_Brúará:SB_TH | 0.440 | 0.765 | -0.13 | 0.557 |
| | | AN_Gufudalsá:Hest | 0.128 | 0.178 | 0.32 | 0.375 |

| | | | | | | |
|---|---|---|---|---|---|---|
| | Gufudalsá to lake morphs | AN_Gufudalsá:Hit | 0.196 | 0.376 | -0.80 | 0.793 |
| | | AN_Gufudalsá:LB_TH | 0.123 | 0.184 | 0.13 | 0.454 |
| | | AN_Gufudalsá:LB_UL | 0.117 | 0.185 | -0.16 | 0.567 |
| | | AN_Gufudalsá:PI_TH | 0.116 | 0.170 | 0.10 | 0.457 |
| | | AN_Gufudalsá:PI_UL | 0.264 | 0.320 | 0.94 | 0.169 |
| | | AN_Gufudalsá:PL_TH | 0.102 | 0.178 | -0.47 | 0.673 |
| | | AN_Gufudalsá:PL_UL | 0.177 | 0.202 | 1.09 | 0.141 |
| | | AN_Gufudalsá:SB_TH | 0.130 | 0.173 | 0.61 | 0.269 |
| | Lambeyrarkvísl to lake morphs | AN_Lambeyrarkvísl:Hest | 0.189 | 0.274 | 0.28 | 0.392 |
| | | AN_Lambeyrarkvísl:Hit | 0.237 | 0.430 | -0.59 | 0.721 |
| | | AN_Lambeyrarkvísl:LB_TH | 0.175 | 0.281 | -0.04 | 0.522 |
| | | AN_Lambeyrarkvísl:LB_UL | 0.160 | 0.285 | -0.43 | 0.665 |
| | | AN_Lambeyrarkvísl:PI_TH | 0.163 | 0.274 | -0.24 | 0.579 |
| | | AN_Lambeyrarkvísl:PI_UL | 0.309 | 0.373 | 0.92 | 0.180 |
| | | AN_Lambeyrarkvísl:PL_TH | 0.152 | 0.278 | -0.54 | 0.712 |
| | | AN_Lambeyrarkvísl:PL_UL | 0.182 | 0.290 | -0.14 | 0.565 |
| | | AN_Lambeyrarkvísl:SB_TH | 0.173 | 0.281 | 0.02 | 0.496 |
| | Víðidalsá to lake morphs | AN_Víðidalsá:Hest | 0.141 | 0.194 | 0.44 | 0.336 |
| | | AN_Víðidalsá:Hit | 0.196 | 0.383 | -0.86 | 0.797 |
| | | AN_Víðidalsá:LB_TH | 0.151 | 0.197 | 0.62 | 0.259 |
| | | AN_Víðidalsá:LB_UL | 0.156 | 0.204 | 0.69 | 0.245 |
| | | AN_Víðidalsá:PI_TH | 0.189 | 0.189 | 1.71 | 0.049 |
| | | AN_Víðidalsá:PI_UL | 0.217 | 0.320 | 0.14 | 0.451 |
| | | AN_Víðidalsá:PL_TH | 0.208 | 0.189 | 1.99 | 0.026 |
| | | AN_Víðidalsá:PL_UL | 0.273 | 0.217 | 2.64 | ***0.004** |
| | | AN_Víðidalsá:SB_TH | 0.199 | 0.187 | 1.92 | 0.028 |
| | Vífisdalsá to lake morphs | AN_Vífilsdalsá:Hest | 0.369 | 0.222 | 3.47 | ***0.002** |
| | | AN_Vífilsdalsá:Hit | 0.403 | 0.400 | 1.72 | 0.047 |
| | | AN_Vífilsdalsá:LB_TH | 0.343 | 0.231 | 3.13 | ***0.003** |
| | | AN_Vífilsdalsá:LB_UL | 0.313 | 0.227 | 2.70 | ***0.003** |
| | | AN_Vífilsdalsá:PI_TH | 0.285 | 0.217 | 2.67 | ***0.003** |
| | | AN_Vífilsdalsá:PI_UL | 0.489 | 0.341 | 2.92 | ***0.003** |
| | | AN_Vífilsdalsá:PL_TH | 0.289 | 0.220 | 2.64 | ***0.003** |
| | | AN_Vífilsdalsá:PL_UL | 0.260 | 0.243 | 1.90 | 0.030 |
| | | AN_Vífilsdalsá:SB_TH | 0.334 | 0.220 | 3.03 | ***0.002** |
| | Hestvatn and Hítarvatn | Hest:Hit | 0.138 | 0.355 | -1.68 | 0.949 |
| | | Hest:LB_TH | 0.064 | 0.114 | -0.68 | 0.757 |
| | | Hest:LB_UL | 0.086 | 0.124 | 0.25 | 0.388 |
| | | Hest:PI_TH | 0.121 | 0.100 | 2.41 | 0.010 |
| | | Hest:PI_UL | 0.199 | 0.279 | 0.33 | 0.378 |
| | | Hest:PL_TH | 0.121 | 0.106 | 2.13 | 0.018 |
| | | Hest:PL_UL | 0.190 | 0.152 | 2.50 | 0.008 |
| | | Hest:SB_TH | 0.099 | 0.096 | 1.75 | 0.042 |
| | | Hit:LB_TH | 0.135 | 0.356 | -1.74 | 0.961 |
| | | Hit:LB_UL | 0.171 | 0.350 | -1.00 | 0.839 |
| | | Hit:PI_TH | 0.205 | 0.352 | -0.22 | 0.575 |
| | | Hit:PI_UL | 0.206 | 0.432 | -1.22 | 0.887 |
| | | Hit:PL_TH | 0.192 | 0.358 | -0.48 | 0.683 |
| | | Hit:PL_UL | 0.234 | 0.366 | 0.06 | 0.487 |
| | | Hit:SB_TH | 0.150 | 0.350 | -1.31 | 0.913 |
| | Þingvallavatn | LB_TH:PI_TH | 0.097 | 0.105 | 1.37 | 0.096 |
| | | LB_TH:PL_TH | 0.098 | 0.111 | 1.18 | 0.124 |
| | | LB_TH:SB_TH | 0.071 | 0.099 | 0.27 | 0.401 |
| | | PI_TH:PL_TH | 0.068 | 0.096 | 0.39 | 0.355 |
| | | PI_TH:SB_TH | 0.087 | 0.085 | 1.72 | 0.044 |
| | | PL_TH:SB_TH | 0.074 | 0.091 | 0.92 | 0.165 |
| | Úlfljótsvatn | LB_UL:PI_UL | 0.246 | 0.283 | 1.12 | 0.127 |
| | | LB_UL:PL_UL | 0.148 | 0.163 | 1.33 | 0.089 |
| | | PI_UL:PL_UL | 0.323 | 0.298 | 1.98 | 0.023 |
| | Across Þingvallavatn and Úlfljótsvatn | LB_TH:LB_UL | 0.078 | 0.127 | -0.31 | 0.627 |
| | | LB_TH:PI_UL | 0.215 | 0.281 | 0.60 | 0.279 |

| | | | | | | |
|---|---|---|---|---|---|---|
| | | LB_TH:PL_UL | 0.157 | 0.152 | 1.74 | 0.041 |
| | | LB_UL:PI_TH | 0.081 | 0.113 | 0.37 | 0.353 |
| | | LB_UL:PL_TH | 0.090 | 0.120 | 0.58 | 0.275 |
| | | LB_UL:SB_TH | 0.093 | 0.112 | 0.99 | 0.156 |
| | | PI_TH:PI_UL | 0.277 | 0.276 | 1.66 | 0.050 |
| | | PI_TH:PL_UL | 0.113 | 0.142 | 0.72 | 0.232 |
| | | PI_UL:PL_TH | 0.283 | 0.278 | 1.71 | 0.039 |
| | | PI_UL:SB_TH | 0.249 | 0.267 | 1.27 | 0.096 |
| | | PL_TH:PL_UL | 0.105 | 0.146 | 0.36 | 0.361 |
| | | PL_UL:SB_TH | 0.121 | 0.141 | 1.07 | 0.145 |
| Premaxilla | AN populations | AN_Brúará:AN_Gufudalsá | 0.243 | 0.428 | -0.84 | 0.792 |
| | | AN_Brúará:AN_Lambeyrarkvísl | 0.218 | 0.500 | -1.88 | 0.968 |
| | | AN_Brúará:AN_Víðidalsá | 0.244 | 0.421 | -0.70 | 0.738 |
| | | AN_Brúará:AN_Vífilsdalsá | 0.322 | 0.487 | -0.15 | 0.553 |
| | | AN_Gufudalsá:AN_Lambeyrarkvísl | 0.227 | 0.342 | -0.23 | 0.586 |
| | | AN_Gufudalsá:AN_Víðidalsá | 0.172 | 0.213 | 0.68 | 0.246 |
| | | AN_Gufudalsá:AN_Vífilsdalsá | 0.178 | 0.308 | -0.93 | 0.827 |
| | | AN_Lambeyrarkvísl:AN_Víðidalsá | 0.269 | 0.325 | 0.72 | 0.245 |
| | | AN_Lambeyrarkvísl:AN_Vífilsdalsá | 0.311 | 0.405 | 0.49 | 0.310 |
| | | AN_Víðidalsá:AN_Vífilsdalsá | 0.207 | 0.296 | 0.02 | 0.492 |
| | Brúará to lake morphs | AN_Brúará:Hest | 0.231 | 0.408 | -0.83 | 0.791 |
| | | AN_Brúará:Hit | 0.510 | 0.529 | 1.40 | 0.077 |
| | | AN_Brúará:LB_TH | 0.263 | 0.412 | -0.30 | 0.619 |
| | | AN_Brúará:LB_UL | 0.263 | 0.412 | -0.40 | 0.651 |
| | | AN_Brúará:PI_TH | 0.254 | 0.408 | -0.40 | 0.658 |
| | | AN_Brúará:PI_UL | 0.301 | 0.466 | -0.26 | 0.596 |
| | | AN_Brúará:PL_TH | 0.330 | 0.410 | 0.69 | 0.251 |
| | | AN_Brúará:PL_UL | 0.361 | 0.438 | 0.84 | 0.207 |
| | | AN_Brúará:SB_TH | 0.376 | 0.409 | 1.27 | 0.113 |
| | Gufudalsá to lake morphs | AN_Gufudalsá:Hest | 0.122 | 0.186 | -0.38 | 0.644 |
| | | AN_Gufudalsá:Hit | 0.377 | 0.394 | 1.39 | 0.081 |
| | | AN_Gufudalsá:LB_TH | 0.081 | 0.188 | -2.17 | 0.986 |
| | | AN_Gufudalsá:LB_UL | 0.065 | 0.203 | -3.43 | 1.000 |
| | | AN_Gufudalsá:PI_TH | 0.131 | 0.181 | 0.13 | 0.447 |
| | | AN_Gufudalsá:PI_UL | 0.155 | 0.277 | -1.07 | 0.853 |
| | | AN_Gufudalsá:PL_TH | 0.172 | 0.186 | 1.27 | 0.109 |
| | | AN_Gufudalsá:PL_UL | 0.188 | 0.224 | 0.77 | 0.210 |
| | | AN_Gufudalsá:SB_TH | 0.209 | 0.179 | 2.45 | ***0.006** |
| | Lambeyrarkvísl to lake morphs | AN_Lambeyrarkvísl:Hest | 0.270 | 0.314 | 0.96 | 0.170 |
| | | AN_Lambeyrarkvísl:Hit | 0.522 | 0.471 | 2.20 | 0.017 |
| | | AN_Lambeyrarkvísl:LB_TH | 0.221 | 0.312 | 0.04 | 0.480 |
| | | AN_Lambeyrarkvísl:LB_UL | 0.237 | 0.323 | 0.19 | 0.412 |
| | | AN_Lambeyrarkvísl:PI_TH | 0.273 | 0.312 | 1.06 | 0.148 |
| | | AN_Lambeyrarkvísl:PI_UL | 0.284 | 0.368 | 0.34 | 0.367 |
| | | AN_Lambeyrarkvísl:PL_TH | 0.321 | 0.311 | 1.77 | 0.032 |
| | | AN_Lambeyrarkvísl:PL_UL | 0.324 | 0.340 | 1.40 | 0.082 |
| | | AN_Lambeyrarkvísl:SB_TH | 0.376 | 0.308 | 2.65 | ***0.004** |
| | Víðidalsá to lake morphs | AN_Víðidalsá:Hest | 0.111 | 0.157 | -0.04 | 0.523 |
| | | AN_Víðidalsá:Hit | 0.384 | 0.400 | 1.54 | 0.069 |
| | | AN_Víðidalsá:LB_TH | 0.175 | 0.159 | 2.09 | 0.017 |
| | | AN_Víðidalsá:LB_UL | 0.178 | 0.180 | 1.63 | 0.058 |
| | | AN_Víðidalsá:PI_TH | 0.134 | 0.147 | 1.17 | 0.127 |
| | | AN_Víðidalsá:PI_UL | 0.225 | 0.257 | 0.92 | 0.202 |
| | | AN_Víðidalsá:PL_TH | 0.208 | 0.157 | 3.15 | ***0.001** |
| | | AN_Víðidalsá:PL_UL | 0.224 | 0.202 | 2.16 | 0.015 |
| | | AN_Víðidalsá:SB_TH | 0.226 | 0.149 | 3.58 | ***0.001** |
| | Vífisdalsá to lake morphs | AN_Vífilsdalsá:Hest | 0.148 | 0.281 | -1.20 | 0.883 |
| | | AN_Vífilsdalsá:Hit | 0.401 | 0.449 | 1.11 | 0.140 |
| | | AN_Vífilsdalsá:LB_TH | 0.173 | 0.278 | -0.58 | 0.711 |
| | | AN_Vífilsdalsá:LB_UL | 0.179 | 0.297 | -0.57 | 0.723 |
| | | AN_Vífilsdalsá:PI_TH | 0.188 | 0.279 | -0.07 | 0.532 |

| | | | | | | |
|---|---|---|---|---|---|---|
| | | AN_Vífilsdalsá:PI_UL | 0.255 | 0.345 | 0.23 | 0.399 |
| | | AN_Vífilsdalsá:PL_TH | 0.173 | 0.280 | -0.47 | 0.672 |
| | | AN_Vífilsdalsá:PL_UL | 0.210 | 0.305 | -0.08 | 0.530 |
| | | AN_Vífilsdalsá:SB_TH | 0.218 | 0.277 | 0.57 | 0.305 |
| | Hestvatn and Hítarvatn | Hest:Hit | 0.364 | 0.379 | 1.42 | 0.080 |
| | | Hest:LB_TH | 0.136 | 0.127 | 2.00 | 0.023 |
| | | Hest:LB_UL | 0.121 | 0.150 | 0.53 | 0.294 |
| | | Hest:PI_TH | 0.088 | 0.114 | 0.44 | 0.337 |
| | | Hest:PI_UL | 0.192 | 0.243 | 0.49 | 0.317 |
| | | Hest:PL_TH | 0.142 | 0.122 | 2.44 | 0.009 |
| | | Hest:PL_UL | 0.197 | 0.177 | 2.24 | 0.013 |
| | | Hest:SB_TH | 0.180 | 0.111 | 4.11 | ***0.001** |
| | | Hit:LB_TH | 0.383 | 0.374 | 1.70 | 0.040 |
| | | Hit:LB_UL | 0.373 | 0.388 | 1.43 | 0.078 |
| | | Hit:PI_TH | 0.400 | 0.373 | 1.92 | 0.020 |
| | | Hit:PI_UL | 0.395 | 0.436 | 1.26 | 0.115 |
| | | Hit:PL_TH | 0.358 | 0.380 | 1.39 | 0.085 |
| | | Hit:PL_UL | 0.330 | 0.396 | 0.76 | 0.243 |
| | | Hit:SB_TH | 0.319 | 0.373 | 0.92 | 0.184 |
| | Þingvallavatn | LB_TH:PI_TH | 0.149 | 0.115 | 2.99 | ***0.003** |
| | | LB_TH:PL_TH | 0.160 | 0.127 | 2.72 | ***0.003** |
| | | LB_TH:SB_TH | 0.203 | 0.114 | 4.88 | ***0.001** |
| | | PI_TH:PL_TH | 0.142 | 0.111 | 3.02 | ***0.001** |
| | | PI_TH:SB_TH | 0.175 | 0.100 | 4.52 | ***0.001** |
| | | PL_TH:SB_TH | 0.086 | 0.107 | 0.52 | 0.302 |
| | Úlfljótsvatn | LB_UL:PI_UL | 0.164 | 0.253 | -0.50 | 0.706 |
| | | LB_UL:PL_UL | 0.163 | 0.201 | 0.64 | 0.266 |
| | | PI_UL:PL_UL | 0.221 | 0.272 | 0.56 | 0.289 |
| | Across Þingvallavatn and Úlfljótsvatn | LB_TH:LB_UL | 0.088 | 0.153 | -0.99 | 0.847 |
| | | LB_TH:PI_UL | 0.156 | 0.240 | -0.46 | 0.674 |
| | | LB_TH:PL_UL | 0.165 | 0.181 | 1.15 | 0.130 |
| | | LB_UL:PI_TH | 0.115 | 0.143 | 0.56 | 0.297 |
| | | LB_UL:PL_TH | 0.141 | 0.149 | 1.38 | 0.094 |
| | | LB_UL:SB_TH | 0.178 | 0.141 | 2.89 | ***0.002** |
| | | PI_TH:PI_UL | 0.169 | 0.236 | 0.02 | 0.496 |
| | | PI_TH:PL_UL | 0.210 | 0.172 | 2.76 | ***0.002** |
| | | PI_UL:PL_TH | 0.176 | 0.243 | 0.11 | 0.452 |
| | | PI_UL:SB_TH | 0.208 | 0.236 | 1.01 | 0.145 |
| | | PL_TH:PL_UL | 0.141 | 0.177 | 0.53 | 0.308 |
| | | PL_UL:SB_TH | 0.145 | 0.171 | 0.82 | 0.213 |
| Maxilla | AN populations | AN_Brúará:AN_Gufudalsá | 0.305 | 0.345 | 1.31 | 0.106 |
| | | AN_Brúará:AN_Lambeyrarkvísl | 0.343 | 0.371 | 1.34 | 0.091 |
| | | AN_Brúará:AN_Víðidalsá | 0.452 | 0.340 | 2.65 | ***0.004** |
| | | AN_Brúará:AN_Vífilsdalsá | 0.375 | 0.359 | 1.79 | 0.035 |
| | | AN_Gufudalsá:AN_Lambeyrarkvísl | 0.114 | 0.229 | -0.75 | 0.784 |
| | | AN_Gufudalsá:AN_Víðidalsá | 0.246 | 0.148 | 3.26 | ***0.002** |
| | | AN_Gufudalsá:AN_Vífilsdalsá | 0.140 | 0.192 | 0.57 | 0.290 |
| | | AN_Lambeyrarkvísl:AN_Víðidalsá | 0.210 | 0.231 | 1.35 | 0.097 |
| | | AN_Lambeyrarkvísl:AN_Vífilsdalsá | 0.131 | 0.261 | -0.68 | 0.761 |
| | | AN_Víðidalsá:AN_Vífilsdalsá | 0.176 | 0.195 | 1.31 | 0.104 |
| | Brúará to lake morphs | AN_Brúará:Hest | 0.358 | 0.327 | 1.95 | 0.024 |
| | | AN_Brúará:Hit | 0.365 | 0.370 | 1.62 | 0.057 |
| | | AN_Brúará:LB_TH | 0.302 | 0.329 | 1.40 | 0.085 |
| | | AN_Brúará:LB_UL | 0.417 | 0.331 | 2.49 | ***0.005** |
| | | AN_Brúará:PI_TH | 0.385 | 0.326 | 2.17 | 0.015 |
| | | AN_Brúará:PI_UL | 0.375 | 0.355 | 1.85 | 0.031 |
| | | AN_Brúará:PL_TH | 0.367 | 0.326 | 2.04 | 0.019 |
| | | AN_Brúará:PL_UL | 0.403 | 0.329 | 2.34 | 0.010 |
| | | AN_Brúará:SB_TH | 0.370 | 0.326 | 2.07 | 0.018 |
| | Gufudalsá to lake morphs | AN_Gufudalsá:Hest | 0.120 | 0.117 | 1.74 | 0.037 |
| | | AN_Gufudalsá:Hit | 0.102 | 0.202 | -0.75 | 0.768 |

| | | | | | | |
|---|---|---|---|---|---|---|
| | | AN_Gufudalsá:LB_TH | 0.123 | 0.120 | 1.75 | 0.040 |
| | | AN_Gufudalsá:LB_UL | 0.149 | 0.126 | 2.25 | 0.014 |
| | | AN_Gufudalsá:PI_TH | 0.160 | 0.114 | 2.69 | ***0.005** |
| | | AN_Gufudalsá:PI_UL | 0.129 | 0.178 | 0.56 | 0.298 |
| | | AN_Gufudalsá:PL_TH | 0.132 | 0.119 | 2.01 | 0.026 |
| | | AN_Gufudalsá:PL_UL | 0.130 | 0.138 | 1.42 | 0.086 |
| | | AN_Gufudalsá:SB_TH | 0.113 | 0.110 | 1.71 | 0.043 |
| | Lambeyrarkvísl to lake morphs | AN_Lambeyrarkvísl:Hest | 0.094 | 0.206 | -1.13 | 0.881 |
| | | AN_Lambeyrarkvísl:Hit | 0.104 | 0.271 | -1.73 | 0.954 |
| | | AN_Lambeyrarkvísl:LB_TH | 0.110 | 0.210 | -0.58 | 0.716 |
| | | AN_Lambeyrarkvísl:LB_UL | 0.120 | 0.214 | -0.37 | 0.639 |
| | | AN_Lambeyrarkvísl:PI_TH | 0.117 | 0.209 | -0.31 | 0.626 |
| | | AN_Lambeyrarkvísl:PI_UL | 0.109 | 0.241 | -1.25 | 0.890 |
| | | AN_Lambeyrarkvísl:PL_TH | 0.101 | 0.211 | -0.88 | 0.820 |
| | | AN_Lambeyrarkvísl:PL_UL | 0.137 | 0.221 | -0.05 | 0.513 |
| | | AN_Lambeyrarkvísl:SB_TH | 0.100 | 0.207 | -0.84 | 0.802 |
| | Víðidalsá to lake morphs | AN_Víðidalsá:Hest | 0.137 | 0.119 | 2.12 | 0.014 |
| | | AN_Víðidalsá:Hit | 0.201 | 0.211 | 1.54 | 0.074 |
| | | AN_Víðidalsá:LB_TH | 0.181 | 0.124 | 2.96 | ***0.002** |
| | | AN_Víðidalsá:LB_UL | 0.137 | 0.129 | 1.86 | 0.040 |
| | | AN_Víðidalsá:PI_TH | 0.114 | 0.120 | 1.50 | 0.072 |
| | | AN_Víðidalsá:PI_UL | 0.128 | 0.178 | 0.47 | 0.321 |
| | | AN_Víðidalsá:PL_TH | 0.154 | 0.123 | 2.43 | ***0.005** |
| | | AN_Víðidalsá:PL_UL | 0.162 | 0.144 | 2.14 | 0.018 |
| | | AN_Víðidalsá:SB_TH | 0.175 | 0.114 | 3.01 | ***0.002** |
| | Vífisdalsá to lake morphs | AN_Vífilsdalsá:Hest | 0.094 | 0.170 | -0.34 | 0.628 |
| | | AN_Vífilsdalsá:Hit | 0.092 | 0.242 | -1.59 | 0.950 |
| | | AN_Vífilsdalsá:LB_TH | 0.111 | 0.172 | 0.19 | 0.424 |
| | | AN_Vífilsdalsá:LB_UL | 0.106 | 0.181 | -0.10 | 0.544 |
| | | AN_Vífilsdalsá:PI_TH | 0.109 | 0.168 | 0.21 | 0.417 |
| | | AN_Vífilsdalsá:PI_UL | 0.090 | 0.217 | -1.35 | 0.912 |
| | | AN_Vífilsdalsá:PL_TH | 0.069 | 0.171 | -1.40 | 0.914 |
| | | AN_Vífilsdalsá:PL_UL | 0.104 | 0.190 | -0.33 | 0.627 |
| | | AN_Vífilsdalsá:SB_TH | 0.085 | 0.165 | -0.62 | 0.728 |
| | Hestvatn and Hítarvatn | Hest:Hit | 0.095 | 0.187 | -0.58 | 0.720 |
| | | Hest:LB_TH | 0.075 | 0.076 | 1.60 | 0.060 |
| | | Hest:LB_UL | 0.071 | 0.088 | 0.85 | 0.200 |
| | | Hest:PI_TH | 0.053 | 0.072 | 0.57 | 0.277 |
| | | Hest:PI_UL | 0.044 | 0.150 | -2.75 | 0.998 |
| | | Hest:PL_TH | 0.052 | 0.077 | 0.26 | 0.394 |
| | | Hest:PL_UL | 0.082 | 0.108 | 0.64 | 0.268 |
| | | Hest:SB_TH | 0.065 | 0.065 | 1.56 | 0.054 |
| | | Hit:LB_TH | 0.118 | 0.184 | 0.14 | 0.449 |
| | | Hit:LB_UL | 0.109 | 0.187 | -0.23 | 0.592 |
| | | Hit:PI_TH | 0.121 | 0.179 | 0.27 | 0.398 |
| | | Hit:PI_UL | 0.100 | 0.224 | -1.20 | 0.880 |
| | | Hit:PL_TH | 0.087 | 0.187 | -0.92 | 0.812 |
| | | Hit:PL_UL | 0.098 | 0.199 | -0.78 | 0.780 |
| | | Hit:SB_TH | 0.072 | 0.179 | -1.47 | 0.928 |
| | Þingvallavatn | LB_TH:PI_TH | 0.090 | 0.071 | 2.51 | ***0.003** |
| | | LB_TH:PL_TH | 0.077 | 0.078 | 1.61 | 0.056 |
| | | LB_TH:SB_TH | 0.098 | 0.069 | 2.97 | ***0.001** |
| | | PI_TH:PL_TH | 0.063 | 0.073 | 1.09 | 0.140 |
| | | PI_TH:SB_TH | 0.087 | 0.062 | 2.79 | ***0.002** |
| | | PL_TH:SB_TH | 0.052 | 0.068 | 0.70 | 0.258 |
| | Úlfljótsvatn | LB_UL:PI_UL | 0.063 | 0.158 | -1.52 | 0.941 |
| | | LB_UL:PL_UL | 0.058 | 0.117 | -0.84 | 0.796 |
| | | PI_UL:PL_UL | 0.078 | 0.167 | -1.07 | 0.857 |
| | Across Þingvallavatn and Úlfljótsvatn | LB_TH:LB_UL | 0.135 | 0.088 | 3.10 | ***0.002** |
| | | LB_TH:PI_UL | 0.095 | 0.155 | 0.02 | 0.489 |
| | | LB_TH:PL_UL | 0.135 | 0.108 | 2.46 | 0.009 |

| | | | | | | |
|---|---|---|---|---|---|---|
| | | LB_UL:PI_TH | 0.082 | 0.085 | 1.49 | 0.068 |
| | | LB_UL:PL_TH | 0.081 | 0.087 | 1.33 | 0.092 |
| | | LB_UL:SB_TH | 0.075 | 0.082 | 1.37 | 0.090 |
| | | PI_TH:PI_UL | 0.058 | 0.149 | -1.58 | 0.939 |
| | | PI_TH:PL_UL | 0.106 | 0.103 | 1.75 | 0.042 |
| | | PI_UL:PL_TH | 0.068 | 0.150 | -1.17 | 0.874 |
| | | PI_UL:SB_TH | 0.079 | 0.149 | -0.49 | 0.689 |
| | | PL_TH:PL_UL | 0.087 | 0.107 | 0.85 | 0.200 |
| | | PL_UL:SB_TH | 0.070 | 0.099 | 0.35 | 0.370 |
| Supramaxilla | AN populations | AN_Brúará:AN_Gufudalsá | 0.599 | 0.588 | 1.77 | 0.044 |
| | | AN_Brúará:AN_Lambeyrarkvísl | 0.504 | 0.682 | 0.75 | 0.226 |
| | | AN_Brúará:AN_Víðidalsá | 0.444 | 0.567 | 0.92 | 0.188 |
| | | AN_Brúará:AN_Vífilsdalsá | 0.478 | 0.675 | 0.67 | 0.272 |
| | | AN_Gufudalsá:AN_Lambeyrarkvísl | 0.446 | 0.465 | 1.44 | 0.072 |
| | | AN_Gufudalsá:AN_Víðidalsá | 0.293 | 0.318 | 1.41 | 0.079 |
| | | AN_Gufudalsá:AN_Vífilsdalsá | 0.213 | 0.454 | -0.52 | 0.695 |
| | | AN_Lambeyrarkvísl:AN_Víðidalsá | 0.302 | 0.472 | 0.34 | 0.370 |
| | | AN_Lambeyrarkvísl:AN_Vífilsdalsá | 0.465 | 0.557 | 1.05 | 0.151 |
| | | AN_Víðidalsá:AN_Vífilsdalsá | 0.235 | 0.433 | -0.17 | 0.574 |
| | Brúará to lake morphs | AN_Brúará:Hest | 0.578 | 0.544 | 1.82 | 0.032 |
| | | AN_Brúará:Hit | 0.740 | 0.712 | 1.74 | 0.043 |
| | | AN_Brúará:LB_TH | 0.436 | 0.542 | 0.97 | 0.169 |
| | | AN_Brúará:LB_UL | 0.593 | 0.559 | 1.86 | 0.028 |
| | | AN_Brúará:PI_TH | 0.483 | 0.545 | 1.29 | 0.110 |
| | | AN_Brúará:PI_UL | 0.530 | 0.625 | 1.15 | 0.134 |
| | | AN_Brúará:PL_TH | 0.544 | 0.548 | 1.65 | 0.054 |
| | | AN_Brúará:PL_UL | 0.520 | 0.564 | 1.40 | 0.086 |
| | | AN_Brúará:SB_TH | 0.615 | 0.537 | 2.04 | 0.016 |
| | Gufudalsá to lake morphs | AN_Gufudalsá:Hest | 0.186 | 0.260 | 0.54 | 0.301 |
| | | AN_Gufudalsá:Hit | 0.500 | 0.547 | 1.41 | 0.083 |
| | | AN_Gufudalsá:LB_TH | 0.240 | 0.256 | 1.35 | 0.082 |
| | | AN_Gufudalsá:LB_UL | 0.172 | 0.279 | 0.17 | 0.433 |
| | | AN_Gufudalsá:PI_TH | 0.164 | 0.255 | 0.25 | 0.413 |
| | | AN_Gufudalsá:PI_UL | 0.224 | 0.402 | -0.05 | 0.502 |
| | | AN_Gufudalsá:PL_TH | 0.075 | 0.253 | -1.78 | 0.961 |
| | | AN_Gufudalsá:PL_UL | 0.107 | 0.308 | -1.23 | 0.897 |
| | | AN_Gufudalsá:SB_TH | 0.145 | 0.249 | -0.03 | 0.514 |
| | Lambeyrarkvísl to lake morphs | AN_Lambeyrarkvísl:Hest | 0.331 | 0.432 | 0.82 | 0.211 |
| | | AN_Lambeyrarkvísl:Hit | 0.800 | 0.629 | 2.52 | ***0.007** |
| | | AN_Lambeyrarkvísl:LB_TH | 0.264 | 0.430 | 0.15 | 0.449 |
| | | AN_Lambeyrarkvísl:LB_UL | 0.403 | 0.451 | 1.33 | 0.090 |
| | | AN_Lambeyrarkvísl:PI_TH | 0.321 | 0.437 | 0.75 | 0.237 |
| | | AN_Lambeyrarkvísl:PI_UL | 0.537 | 0.532 | 1.67 | 0.047 |
| | | AN_Lambeyrarkvísl:PL_TH | 0.400 | 0.430 | 1.43 | 0.078 |
| | | AN_Lambeyrarkvísl:PL_UL | 0.412 | 0.459 | 1.31 | 0.103 |
| | | AN_Lambeyrarkvísl:SB_TH | 0.389 | 0.426 | 1.33 | 0.097 |
| | Víðidalsá to lake morphs | AN_Víðidalsá:Hest | 0.197 | 0.249 | 0.94 | 0.186 |
| | | AN_Víðidalsá:Hit | 0.538 | 0.526 | 1.71 | 0.045 |
| | | AN_Víðidalsá:LB_TH | 0.116 | 0.243 | -0.56 | 0.708 |
| | | AN_Víðidalsá:LB_UL | 0.194 | 0.253 | 0.74 | 0.233 |
| | | AN_Víðidalsá:PI_TH | 0.149 | 0.245 | 0.14 | 0.448 |
| | | AN_Víðidalsá:PI_UL | 0.405 | 0.386 | 1.82 | 0.033 |
| | | AN_Víðidalsá:PL_TH | 0.235 | 0.240 | 1.55 | 0.057 |
| | | AN_Víðidalsá:PL_UL | 0.220 | 0.291 | 0.81 | 0.213 |
| | | AN_Víðidalsá:SB_TH | 0.236 | 0.225 | 1.73 | 0.035 |
| | Vífisdalsá to lake morphs | AN_Vífilsdalsá:Hest | 0.239 | 0.402 | 0.09 | 0.473 |
| | | AN_Vífilsdalsá:Hit | 0.399 | 0.623 | 0.35 | 0.355 |
| | | AN_Vífilsdalsá:LB_TH | 0.228 | 0.395 | -0.08 | 0.538 |
| | | AN_Vífilsdalsá:LB_UL | 0.221 | 0.407 | -0.19 | 0.568 |
| | | AN_Vífilsdalsá:PI_TH | 0.177 | 0.394 | -0.68 | 0.751 |
| | | AN_Vífilsdalsá:PI_UL | 0.282 | 0.508 | -0.05 | 0.511 |

| | | | | | | |
|---|---|---|---|---|---|---|
| | | AN_Vífilsdalsá:PL_TH | 0.170 | 0.391 | -0.75 | 0.763 |
| | | AN_Vífilsdalsá:PL_UL | 0.140 | 0.415 | -1.46 | 0.926 |
| | | AN_Vífilsdalsá:SB_TH | 0.225 | 0.382 | -0.02 | 0.524 |
| | Hestvatn and Hítarvatn | Hest:Hit | 0.564 | 0.503 | 2.04 | 0.020 |
| | | Hest:LB_TH | 0.157 | 0.169 | 1.42 | 0.079 |
| | | Hest:LB_UL | 0.120 | 0.188 | 0.21 | 0.413 |
| | | Hest:PI_TH | 0.113 | 0.163 | 0.54 | 0.301 |
| | | Hest:PI_UL | 0.360 | 0.340 | 1.87 | 0.035 |
| | | Hest:PL_TH | 0.156 | 0.160 | 1.59 | 0.062 |
| | | Hest:PL_UL | 0.176 | 0.217 | 0.94 | 0.170 |
| | | Hest:SB_TH | 0.099 | 0.146 | 0.44 | 0.335 |
| | | Hit:LB_TH | 0.588 | 0.500 | 2.17 | 0.014 |
| | | Hit:LB_UL | 0.479 | 0.510 | 1.45 | 0.077 |
| | | Hit:PI_TH | 0.530 | 0.488 | 1.85 | 0.028 |
| | | Hit:PI_UL | 0.569 | 0.562 | 1.62 | 0.047 |
| | | Hit:PL_TH | 0.486 | 0.490 | 1.58 | 0.059 |
| | | Hit:PL_UL | 0.448 | 0.510 | 1.18 | 0.137 |
| | | Hit:SB_TH | 0.501 | 0.496 | 1.68 | 0.047 |
| | Þingvallavatn | LB_TH:PI_TH | 0.092 | 0.164 | -0.09 | 0.528 |
| | | LB_TH:PL_TH | 0.190 | 0.162 | 2.12 | 0.014 |
| | | LB_TH:SB_TH | 0.217 | 0.150 | 2.80 | ***0.003** |
| | | PI_TH:PL_TH | 0.105 | 0.147 | 0.55 | 0.303 |
| | | PI_TH:SB_TH | 0.142 | 0.137 | 1.78 | 0.040 |
| | | PL_TH:SB_TH | 0.120 | 0.135 | 1.27 | 0.118 |
| | Úlfljótsvatn | LB_UL:PI_UL | 0.359 | 0.361 | 1.63 | 0.055 |
| | | LB_UL:PL_UL | 0.143 | 0.242 | 0.10 | 0.453 |
| | | PI_UL:PL_UL | 0.248 | 0.375 | 0.41 | 0.341 |
| | Across Þingvallavatn and Úlfljótsvatn | LB_TH:LB_UL | 0.204 | 0.190 | 1.86 | 0.033 |
| | | LB_TH:PI_UL | 0.332 | 0.340 | 1.56 | 0.063 |
| | | LB_TH:PL_UL | 0.190 | 0.230 | 1.09 | 0.142 |
| | | LB_UL:PI_TH | 0.138 | 0.184 | 0.75 | 0.240 |
| | | LB_UL:PL_TH | 0.142 | 0.182 | 0.85 | 0.211 |
| | | LB_UL:SB_TH | 0.095 | 0.169 | -0.11 | 0.549 |
| | | PI_TH:PI_UL | 0.290 | 0.338 | 1.18 | 0.119 |
| | | PI_TH:PL_UL | 0.110 | 0.212 | -0.25 | 0.582 |
| | | PI_UL:PL_TH | 0.239 | 0.332 | 0.63 | 0.259 |
| | | PI_UL:SB_TH | 0.352 | 0.331 | 1.80 | 0.041 |
| | | PL_TH:PL_UL | 0.057 | 0.216 | -1.81 | 0.966 |
| | | PL_UL:SB_TH | 0.141 | 0.208 | 0.50 | 0.319 |

D = pairwise distances between mean; UCL = upper confidence level, Z = Z-statistic; P = P-value, ≤ 0.008 in bold with asterisk.

***Supplementary material 17****: Results from "Significant Vector Angle test" examining possibly allometric shape differences by population for five bones, that showed significant allometric differences according to ANOVA model with main and interaction effects (population x bone size). Used to verify results from ANOVA and test for true allometry (i.e., different group slopes). AN-charr treated as five separate populations.*

| Bone | System | Pairs | R | Angle | UCL | Z | P-value |
|---|---|---|---|---|---|---|---|
| Dentary | AN populations | AN_Brúará:AN_Gufudalsá | 0.885 | 27.689 | 102.885 | -2.70 | 0.999 |
| | | AN_Brúará:AN_Lambeyrarkvísl | 0.038 | 87.831 | 107.396 | 0.73 | 0.240 |
| | | AN_Brúará:AN_Víðidalsá | 0.331 | 70.660 | 100.481 | 0.30 | 0.381 |
| | | AN_Brúará:AN_Vífilsdalsá | 0.574 | 54.972 | 104.802 | -0.94 | 0.811 |
| | | AN_Gufudalsá:AN_Lambeyrarkvísl | -0.055 | 93.173 | 97.469 | 1.49 | 0.069 |
| | | AN_Gufudalsá:AN_Víðidalsá | 0.270 | 74.314 | 75.722 | 1.61 | 0.061 |
| | | AN_Gufudalsá:AN_Vífilsdalsá | 0.482 | 61.170 | 93.577 | 0.06 | 0.482 |
| | | AN_Lambeyrarkvísl:AN_Víðidalsá | -0.179 | 100.306 | 94.480 | 1.84 | 0.033 |
| | | AN_Lambeyrarkvísl:AN_Vífilsdalsá | 0.233 | 76.536 | 103.703 | 0.35 | 0.374 |

| | | | | | | | |
|---|---|---|---|---|---|---|---|
| | | AN_Víðidalsá:AN_Vífilsdalsá | 0.652 | 49.340 | 91.580 | -0.59 | 0.707 |
| | Brúará to lake morphs | AN_Brúará:Hest | 0.651 | 49.408 | 98.869 | -0.70 | 0.736 |
| | | AN_Brúará:Hit | 0.677 | 47.410 | 107.308 | -1.42 | 0.922 |
| | | AN_Brúará:LB_TH | 0.813 | 35.581 | 100.490 | -1.67 | 0.959 |
| | | AN_Brúará:LB_UL | 0.652 | 49.332 | 99.035 | -0.79 | 0.783 |
| | | AN_Brúará:PI_TH | 0.340 | 70.111 | 96.782 | 0.50 | 0.313 |
| | | AN_Brúará:PI_UL | 0.511 | 59.297 | 102.523 | -0.56 | 0.693 |
| | | AN_Brúará:PL_TH | 0.827 | 34.221 | 99.298 | -1.86 | 0.975 |
| | | AN_Brúará:PL_UL | 0.769 | 39.743 | 100.829 | -1.59 | 0.947 |
| | | AN_Brúará:SB_TH | 0.680 | 47.139 | 98.941 | -0.75 | 0.760 |
| | Gufudalsá to lake morphs | AN_Gufudalsá:Hest | 0.795 | 37.357 | 66.786 | -0.40 | 0.658 |
| | | AN_Gufudalsá:Hit | 0.776 | 39.105 | 93.384 | -1.56 | 0.937 |
| | | AN_Gufudalsá:LB_TH | 0.845 | 32.349 | 67.029 | -0.86 | 0.796 |
| | | AN_Gufudalsá:LB_UL | 0.762 | 40.384 | 69.692 | -0.34 | 0.632 |
| | | AN_Gufudalsá:PI_TH | 0.366 | 68.505 | 66.096 | 1.84 | 0.038 |
| | | AN_Gufudalsá:PI_UL | 0.620 | 51.658 | 87.881 | -0.36 | 0.635 |
| | | AN_Gufudalsá:PL_TH | 0.861 | 30.571 | 67.176 | -1.01 | 0.846 |
| | | AN_Gufudalsá:PL_UL | 0.864 | 30.192 | 73.354 | -1.64 | 0.947 |
| | | AN_Gufudalsá:SB_TH | 0.733 | 42.898 | 65.754 | 0.31 | 0.388 |
| | Lambeyrarkvísl to lake morphs | AN_Lambeyrarkvísl:Hest | 0.015 | 89.126 | 92.404 | 1.52 | 0.065 |
| | | AN_Lambeyrarkvísl:Hit | 0.180 | 79.622 | 103.794 | 0.53 | 0.299 |
| | | AN_Lambeyrarkvísl:LB_TH | -0.016 | 90.912 | 93.666 | 1.59 | 0.058 |
| | | AN_Lambeyrarkvísl:LB_UL | -0.042 | 92.434 | 92.128 | 1.61 | 0.049 |
| | | AN_Lambeyrarkvísl:PI_TH | -0.345 | 110.196 | 92.667 | 2.24 | 0.012 |
| | | AN_Lambeyrarkvísl:PI_UL | 0.010 | 89.407 | 99.930 | 1.14 | 0.135 |
| | | AN_Lambeyrarkvísl:PL_TH | 0.068 | 86.080 | 92.470 | 1.42 | 0.083 |
| | | AN_Lambeyrarkvísl:PL_UL | -0.016 | 90.909 | 94.641 | 1.49 | 0.070 |
| | | AN_Lambeyrarkvísl:SB_TH | 0.121 | 83.034 | 92.441 | 1.30 | 0.100 |
| | Víðidalsá to lake morphs | AN_Víðidalsá:Hest | 0.318 | 71.450 | 59.869 | 2.25 | 0.013 |
| | | AN_Víðidalsá:Hit | -0.039 | 92.231 | 92.796 | 1.66 | 0.053 |
| | | AN_Víðidalsá:LB_TH | 0.523 | 58.467 | 58.869 | 1.60 | 0.054 |
| | | AN_Víðidalsá:LB_UL | 0.287 | 73.348 | 62.844 | 2.22 | 0.014 |
| | | AN_Víðidalsá:PI_TH | 0.506 | 59.589 | 58.616 | 1.69 | 0.043 |
| | | AN_Víðidalsá:PI_UL | 0.264 | 74.712 | 85.642 | 1.15 | 0.131 |
| | | AN_Víðidalsá:PL_TH | 0.118 | 83.209 | 59.219 | 2.77 | ***0.004** |
| | | AN_Víðidalsá:PL_UL | 0.373 | 68.115 | 72.648 | 1.49 | 0.070 |
| | | AN_Víðidalsá:SB_TH | -0.023 | 91.342 | 57.989 | 2.97 | ***0.002** |
| | Vífisdalsá to lake morphs | AN_Vífilsdalsá:Hest | 0.509 | 59.423 | 90.677 | 0.35 | 0.372 |
| | | AN_Vífilsdalsá:Hit | 0.428 | 64.630 | 101.110 | -0.14 | 0.552 |
| | | AN_Vífilsdalsá:LB_TH | 0.673 | 47.680 | 88.296 | -0.34 | 0.618 |
| | | AN_Vífilsdalsá:LB_UL | 0.557 | 56.118 | 89.713 | 0.06 | 0.462 |
| | | AN_Vífilsdalsá:PI_TH | 0.255 | 75.203 | 89.443 | 1.15 | 0.130 |
| | | AN_Vífilsdalsá:PI_UL | 0.293 | 72.951 | 98.637 | 0.48 | 0.320 |
| | | AN_Vífilsdalsá:PL_TH | 0.463 | 62.420 | 89.840 | 0.53 | 0.304 |
| | | AN_Vífilsdalsá:PL_UL | 0.630 | 50.924 | 92.310 | -0.48 | 0.668 |
| | | AN_Vífilsdalsá:SB_TH | 0.379 | 67.737 | 88.973 | 0.84 | 0.200 |
| | Hestvatn and Hítarvatn | Hest:Hit | 0.824 | 34.481 | 88.943 | -1.43 | 0.923 |
| | | Hest:LB_TH | 0.833 | 33.578 | 44.376 | 0.56 | 0.292 |
| | | Hest:LB_UL | 0.867 | 29.895 | 49.065 | -0.33 | 0.620 |
| | | Hest:PI_TH | 0.586 | 54.113 | 39.845 | 2.90 | ***0.002** |
| | | Hest:PI_UL | 0.734 | 42.770 | 79.902 | -0.41 | 0.649 |
| | | Hest:PL_TH | 0.837 | 33.149 | 43.230 | 0.60 | 0.283 |
| | | Hest:PL_UL | 0.866 | 30.046 | 59.128 | -0.84 | 0.794 |
| | | Hest:SB_TH | 0.800 | 36.840 | 39.571 | 1.43 | 0.074 |
| | | Hit:LB_TH | 0.661 | 48.586 | 89.039 | -0.27 | 0.600 |
| | | Hit:LB_UL | 0.801 | 36.799 | 91.553 | -1.36 | 0.903 |
| | | Hit:PI_TH | 0.314 | 71.699 | 87.675 | 1.04 | 0.168 |
| | | Hit:PI_UL | 0.636 | 50.500 | 97.521 | -0.87 | 0.811 |
| | | Hit:PL_TH | 0.899 | 25.959 | 88.060 | -2.32 | 0.993 |
| | | Hit:PL_UL | 0.852 | 31.577 | 91.296 | -2.01 | 0.975 |
| | | Hit:SB_TH | 0.868 | 29.754 | 87.069 | -1.77 | 0.963 |

| | | | | | | | |
|---|---|---|---|---|---|---|---|
| | Þingvallavatn | LB_TH:PI_TH | 0.447 | 63.430 | 39.937 | 3.57 | ***0.001** |
| | | LB_TH:PL_TH | 0.798 | 37.068 | 41.979 | 1.15 | 0.121 |
| | | LB_TH:SB_TH | 0.672 | 47.797 | 39.319 | 2.50 | 0.008 |
| | | PI_TH:PL_TH | 0.464 | 62.329 | 39.638 | 3.51 | ***0.001** |
| | | PI_TH:SB_TH | 0.467 | 62.156 | 34.885 | 4.10 | ***0.001** |
| | | PL_TH:SB_TH | 0.936 | 20.678 | 37.464 | -0.70 | 0.765 |
| | Úlfljótsvatn | LB_UL:PI_UL | 0.689 | 46.466 | 83.826 | -0.30 | 0.613 |
| | | LB_UL:PL_UL | 0.865 | 30.104 | 63.132 | -1.04 | 0.841 |
| | | PI_UL:PL_UL | 0.621 | 51.592 | 83.755 | -0.21 | 0.584 |
| | Across Þingvallavatn and Úlfljótsvatn | LB_TH:LB_UL | 0.808 | 36.129 | 48.702 | 0.42 | 0.337 |
| | | LB_TH:PI_UL | 0.660 | 48.675 | 81.492 | 0.05 | 0.474 |
| | | LB_TH:PL_UL | 0.846 | 32.173 | 60.109 | -0.56 | 0.710 |
| | | LB_UL:PI_TH | 0.521 | 58.633 | 45.371 | 2.63 | ***0.007** |
| | | LB_UL:PL_TH | 0.814 | 35.481 | 46.330 | 0.44 | 0.330 |
| | | LB_UL:SB_TH | 0.750 | 41.423 | 45.308 | 1.23 | 0.111 |
| | | PI_TH:PI_UL | 0.587 | 54.086 | 79.946 | 0.43 | 0.332 |
| | | PI_TH:PL_UL | 0.489 | 60.731 | 58.324 | 1.79 | 0.035 |
| | | PI_UL:PL_TH | 0.702 | 45.415 | 81.018 | -0.19 | 0.572 |
| | | PI_UL:SB_TH | 0.749 | 41.476 | 79.639 | -0.36 | 0.629 |
| | | PL_TH:PL_UL | 0.863 | 30.289 | 58.678 | -0.77 | 0.775 |
| | | PL_UL:SB_TH | 0.764 | 40.219 | 58.740 | 0.40 | 0.343 |
| Articular-angular | AN populations | AN_Brúará:AN_Gufudalsá | 0.618 | 51.815 | 100.533 | -0.92 | 0.818 |
| | | AN_Brúará:AN_Lambeyrarkvísl | -0.072 | 94.135 | 107.439 | 0.98 | 0.172 |
| | | AN_Brúará:AN_Víðidalsá | 0.339 | 70.184 | 101.553 | 0.18 | 0.435 |
| | | AN_Brúará:AN_Vífilsdalsá | 0.465 | 62.280 | 107.527 | -0.56 | 0.703 |
| | | AN_Gufudalsá:AN_Lambeyrarkvísl | -0.359 | 111.059 | 98.014 | 2.20 | 0.010 |
| | | AN_Gufudalsá:AN_Víðidalsá | 0.748 | 41.554 | 78.605 | -0.76 | 0.780 |
| | | AN_Gufudalsá:AN_Vífilsdalsá | 0.762 | 40.339 | 96.561 | -1.54 | 0.933 |
| | | AN_Lambeyrarkvísl:AN_Víðidalsá | -0.249 | 104.390 | 100.469 | 1.84 | 0.036 |
| | | AN_Lambeyrarkvísl:AN_Vífilsdalsá | -0.231 | 103.371 | 104.734 | 1.57 | 0.062 |
| | | AN_Víðidalsá:AN_Vífilsdalsá | 0.868 | 29.819 | 94.012 | -2.38 | 0.995 |
| | Brúará to lake morphs | AN_Brúará:Hest | 0.675 | 47.541 | 95.869 | -0.86 | 0.800 |
| | | AN_Brúará:Hit | 0.302 | 72.427 | 105.528 | -0.03 | 0.513 |
| | | AN_Brúará:LB_TH | 0.467 | 62.179 | 97.660 | -0.01 | 0.490 |
| | | AN_Brúará:LB_UL | 0.533 | 57.766 | 97.071 | -0.28 | 0.594 |
| | | AN_Brúará:PI_TH | 0.359 | 68.965 | 96.034 | 0.41 | 0.340 |
| | | AN_Brúará:PI_UL | 0.500 | 60.013 | 104.550 | -0.56 | 0.702 |
| | | AN_Brúará:PL_TH | 0.222 | 77.177 | 97.280 | 0.78 | 0.235 |
| | | AN_Brúará:PL_UL | 0.761 | 40.416 | 97.834 | -1.60 | 0.951 |
| | | AN_Brúará:SB_TH | 0.448 | 63.372 | 96.499 | 0.16 | 0.443 |
| | Gufudalsá to lake morphs | AN_Gufudalsá:Hest | 0.891 | 26.990 | 67.697 | -1.59 | 0.947 |
| | | AN_Gufudalsá:Hit | 0.653 | 49.214 | 94.810 | -0.83 | 0.798 |
| | | AN_Gufudalsá:LB_TH | 0.917 | 23.502 | 67.149 | -2.06 | 0.980 |
| | | AN_Gufudalsá:LB_UL | 0.757 | 40.831 | 70.462 | -0.34 | 0.631 |
| | | AN_Gufudalsá:PI_TH | 0.834 | 33.508 | 65.050 | -0.73 | 0.760 |
| | | AN_Gufudalsá:PI_UL | 0.618 | 51.862 | 89.278 | -0.41 | 0.645 |
| | | AN_Gufudalsá:PL_TH | 0.564 | 55.669 | 67.669 | 0.94 | 0.174 |
| | | AN_Gufudalsá:PL_UL | 0.813 | 35.612 | 75.994 | -1.15 | 0.869 |
| | | AN_Gufudalsá:SB_TH | 0.748 | 41.606 | 63.825 | 0.08 | 0.466 |
| | Lambeyrarkvísl to lake morphs | AN_Lambeyrarkvísl:Hest | -0.255 | 104.799 | 95.243 | 2.14 | 0.017 |
| | | AN_Lambeyrarkvísl:Hit | -0.560 | 124.023 | 103.751 | 2.52 | ***0.003** |
| | | AN_Lambeyrarkvísl:LB_TH | -0.255 | 104.800 | 93.819 | 2.09 | 0.015 |
| | | AN_Lambeyrarkvísl:LB_UL | 0.014 | 89.179 | 98.099 | 1.35 | 0.103 |
| | | AN_Lambeyrarkvísl:PI_TH | -0.355 | 110.774 | 94.347 | 2.35 | 0.008 |
| | | AN_Lambeyrarkvísl:PI_UL | -0.006 | 90.361 | 101.853 | 1.12 | 0.127 |
| | | AN_Lambeyrarkvísl:PL_TH | -0.041 | 92.360 | 95.552 | 1.57 | 0.062 |
| | | AN_Lambeyrarkvísl:PL_UL | -0.196 | 101.274 | 95.962 | 1.83 | 0.033 |
| | | AN_Lambeyrarkvísl:SB_TH | -0.154 | 98.833 | 93.115 | 1.89 | 0.031 |
| | Víðidalsá to lake morphs | AN_Víðidalsá:Hest | 0.836 | 33.281 | 66.192 | -0.77 | 0.771 |
| | | AN_Víðidalsá:Hit | 0.757 | 40.806 | 93.136 | -1.32 | 0.904 |
| | | AN_Víðidalsá:LB_TH | 0.868 | 29.762 | 65.589 | -1.20 | 0.880 |

| | | | | | | | |
|---|---|---|---|---|---|---|---|
| | | AN_Víðidalsá:LB_UL | 0.633 | 50.716 | 69.886 | 0.53 | 0.305 |
| | | AN_Víðidalsá:PI_TH | 0.742 | 42.069 | 65.877 | 0.16 | 0.433 |
| | | AN_Víðidalsá:PI_UL | 0.694 | 46.013 | 85.779 | -0.72 | 0.762 |
| | | AN_Víðidalsá:PL_TH | 0.734 | 42.780 | 66.064 | 0.14 | 0.451 |
| | | AN_Víðidalsá:PL_UL | 0.747 | 41.701 | 75.055 | -0.58 | 0.714 |
| | | AN_Víðidalsá:SB_TH | 0.759 | 40.629 | 62.049 | 0.11 | 0.456 |
| | Vífisdalsá to lake morphs | AN_Vífilsdalsá:Hest | 0.891 | 26.949 | 88.777 | -2.34 | 0.992 |
| | | AN_Vífilsdalsá:Hit | 0.613 | 52.170 | 101.935 | -1.06 | 0.865 |
| | | AN_Vífilsdalsá:LB_TH | 0.881 | 28.220 | 89.358 | -2.23 | 0.987 |
| | | AN_Vífilsdalsá:LB_UL | 0.791 | 37.700 | 91.682 | -1.40 | 0.920 |
| | | AN_Vífilsdalsá:PI_TH | 0.712 | 44.595 | 89.791 | -0.71 | 0.760 |
| | | AN_Vífilsdalsá:PI_UL | 0.553 | 56.405 | 101.547 | -0.61 | 0.727 |
| | | AN_Vífilsdalsá:PL_TH | 0.797 | 37.190 | 89.667 | -1.35 | 0.914 |
| | | AN_Vífilsdalsá:PL_UL | 0.827 | 34.204 | 94.574 | -1.85 | 0.963 |
| | | AN_Vífilsdalsá:SB_TH | 0.795 | 37.369 | 89.957 | -1.26 | 0.899 |
| | Hestvatn and Hítarvatn | Hest:Hit | 0.652 | 49.291 | 89.041 | -0.32 | 0.617 |
| | | Hest:LB_TH | 0.894 | 26.562 | 44.373 | -0.35 | 0.639 |
| | | Hest:LB_UL | 0.768 | 39.840 | 50.135 | 0.75 | 0.227 |
| | | Hest:PI_TH | 0.842 | 32.618 | 41.707 | 0.70 | 0.238 |
| | | Hest:PI_UL | 0.695 | 45.953 | 79.139 | -0.23 | 0.589 |
| | | Hest:PL_TH | 0.714 | 44.402 | 44.086 | 1.68 | 0.047 |
| | | Hest:PL_UL | 0.882 | 28.144 | 62.118 | -1.19 | 0.880 |
| | | Hest:SB_TH | 0.822 | 34.742 | 38.102 | 1.19 | 0.111 |
| | | Hit:LB_TH | 0.685 | 46.795 | 89.189 | -0.52 | 0.690 |
| | | Hit:LB_UL | 0.274 | 74.116 | 90.974 | 0.95 | 0.170 |
| | | Hit:PI_TH | 0.738 | 42.436 | 88.287 | -0.74 | 0.768 |
| | | Hit:PI_UL | 0.332 | 70.639 | 99.625 | 0.30 | 0.368 |
| | | Hit:PL_TH | 0.557 | 56.137 | 88.932 | 0.13 | 0.439 |
| | | Hit:PL_UL | 0.585 | 54.177 | 92.887 | -0.33 | 0.634 |
| | | Hit:SB_TH | 0.678 | 47.300 | 87.571 | -0.38 | 0.644 |
| | Þingvallavatn | LB_TH:PI_TH | 0.851 | 31.703 | 40.305 | 0.59 | 0.277 |
| | | LB_TH:PL_TH | 0.699 | 45.647 | 44.074 | 1.77 | 0.041 |
| | | LB_TH:SB_TH | 0.770 | 39.634 | 39.308 | 1.66 | 0.047 |
| | | PI_TH:PL_TH | 0.697 | 45.809 | 40.216 | 2.16 | 0.013 |
| | | PI_TH:SB_TH | 0.792 | 37.653 | 35.822 | 1.86 | 0.032 |
| | | PL_TH:SB_TH | 0.917 | 23.516 | 38.325 | -0.31 | 0.621 |
| | Úlfljótsvatn | LB_UL:PI_UL | 0.653 | 49.202 | 81.080 | -0.16 | 0.567 |
| | | LB_UL:PL_UL | 0.712 | 44.640 | 64.508 | 0.26 | 0.397 |
| | | PI_UL:PL_UL | 0.569 | 55.304 | 86.211 | -0.01 | 0.505 |
| | Across Þingvallavatn and Úlfljótsvatn | LB_TH:LB_UL | 0.787 | 38.107 | 50.640 | 0.53 | 0.306 |
| | | LB_TH:PI_UL | 0.615 | 52.079 | 80.574 | 0.17 | 0.437 |
| | | LB_TH:PL_UL | 0.788 | 38.045 | 62.633 | -0.12 | 0.542 |
| | | LB_UL:PI_TH | 0.572 | 55.102 | 47.282 | 2.27 | 0.010 |
| | | LB_UL:PL_TH | 0.646 | 49.769 | 48.948 | 1.63 | 0.045 |
| | | LB_UL:SB_TH | 0.646 | 49.771 | 46.181 | 1.91 | 0.026 |
| | | PI_TH:PI_UL | 0.527 | 58.195 | 79.325 | 0.63 | 0.257 |
| | | PI_TH:PL_UL | 0.596 | 53.432 | 59.234 | 1.28 | 0.111 |
| | | PI_UL:PL_TH | 0.381 | 67.588 | 79.745 | 1.08 | 0.137 |
| | | PI_UL:SB_TH | 0.458 | 62.759 | 79.099 | 0.92 | 0.167 |
| | | PL_TH:PL_UL | 0.617 | 51.900 | 61.150 | 1.08 | 0.141 |
| | | PL_UL:SB_TH | 0.765 | 40.068 | 59.102 | 0.28 | 0.391 |
| Quadrate | AN populations | AN_Brúará:AN_Gufudalsá | 0.403 | 66.254 | 118.053 | -0.71 | 0.753 |
| | | AN_Brúará:AN_Lambeyrarkvísl | 0.248 | 75.665 | 117.990 | -0.36 | 0.622 |
| | | AN_Brúará:AN_Víðidalsá | 0.464 | 62.361 | 115.044 | -0.90 | 0.804 |
| | | AN_Brúará:AN_Vífilsdalsá | -0.231 | 103.382 | 118.357 | 0.99 | 0.170 |
| | | AN_Gufudalsá:AN_Lambeyrarkvísl | 0.475 | 61.655 | 104.417 | -0.39 | 0.627 |
| | | AN_Gufudalsá:AN_Víðidalsá | 0.777 | 39.048 | 94.109 | -1.45 | 0.920 |
| | | AN_Gufudalsá:AN_Vífilsdalsá | 0.159 | 80.870 | 101.180 | 0.83 | 0.211 |
| | | AN_Lambeyrarkvísl:AN_Víðidalsá | 0.507 | 59.543 | 104.135 | -0.56 | 0.712 |
| | | AN_Lambeyrarkvísl:AN_Vífilsdalsá | 0.017 | 88.997 | 106.076 | 0.84 | 0.210 |
| | | AN_Víðidalsá:AN_Vífilsdalsá | -0.085 | 94.850 | 98.852 | 1.42 | 0.080 |

| | | | | | | | |
|---|---|---|---|---|---|---|---|
| | Brúará to lake morphs | AN_Brúará:Hest | 0.452 | 63.140 | 116.375 | -0.67 | 0.725 |
| | | AN_Brúará:Hit | 0.719 | 44.032 | 120.915 | -2.03 | 0.984 |
| | | AN_Brúará:LB_TH | 0.508 | 59.475 | 118.428 | -0.88 | 0.789 |
| | | AN_Brúará:LB_UL | 0.366 | 68.537 | 115.982 | -0.46 | 0.665 |
| | | AN_Brúará:PI_TH | 0.181 | 79.600 | 117.629 | 0.17 | 0.450 |
| | | AN_Brúará:PI_UL | 0.643 | 50.010 | 118.031 | -1.59 | 0.943 |
| | | AN_Brúará:PL_TH | 0.195 | 78.772 | 116.840 | 0.10 | 0.468 |
| | | AN_Brúará:PL_UL | -0.119 | 96.858 | 116.193 | 0.81 | 0.222 |
| | | AN_Brúará:SB_TH | 0.527 | 58.223 | 116.586 | -0.85 | 0.780 |
| | Gufudalsá to lake morphs | AN_Gufudalsá:Hest | 0.719 | 44.034 | 82.547 | -0.38 | 0.656 |
| | | AN_Gufudalsá:Hit | 0.501 | 59.914 | 107.002 | -0.68 | 0.740 |
| | | AN_Gufudalsá:LB_TH | 0.658 | 48.862 | 82.164 | -0.08 | 0.522 |
| | | AN_Gufudalsá:LB_UL | 0.684 | 46.812 | 85.396 | -0.29 | 0.599 |
| | | AN_Gufudalsá:PI_TH | 0.655 | 49.116 | 79.434 | 0.15 | 0.433 |
| | | AN_Gufudalsá:PI_UL | 0.615 | 52.038 | 104.264 | -0.95 | 0.834 |
| | | AN_Gufudalsá:PL_TH | 0.771 | 39.538 | 80.970 | -0.65 | 0.762 |
| | | AN_Gufudalsá:PL_UL | -0.041 | 92.338 | 86.852 | 1.91 | 0.022 |
| | | AN_Gufudalsá:SB_TH | 0.546 | 56.899 | 80.068 | 0.65 | 0.243 |
| | Lambeyrarkvísl to lake morphs | AN_Lambeyrarkvísl:Hest | 0.439 | 63.983 | 96.354 | 0.18 | 0.419 |
| | | AN_Lambeyrarkvísl:Hit | 0.305 | 72.217 | 113.634 | -0.26 | 0.605 |
| | | AN_Lambeyrarkvísl:LB_TH | 0.415 | 65.460 | 97.309 | 0.22 | 0.400 |
| | | AN_Lambeyrarkvísl:LB_UL | 0.506 | 59.593 | 98.334 | -0.13 | 0.538 |
| | | AN_Lambeyrarkvísl:PI_TH | 0.408 | 65.939 | 96.557 | 0.34 | 0.362 |
| | | AN_Lambeyrarkvísl:PI_UL | 0.377 | 67.853 | 108.661 | -0.35 | 0.634 |
| | | AN_Lambeyrarkvísl:PL_TH | 0.508 | 59.494 | 96.706 | -0.04 | 0.501 |
| | | AN_Lambeyrarkvísl:PL_UL | 0.161 | 80.746 | 98.631 | 0.78 | 0.227 |
| | | AN_Lambeyrarkvísl:SB_TH | 0.324 | 71.108 | 97.452 | 0.61 | 0.285 |
| | Víðidalsá to lake morphs | AN_Víðidalsá:Hest | 0.866 | 30.037 | 83.882 | -1.73 | 0.956 |
| | | AN_Víðidalsá:Hit | 0.685 | 46.744 | 108.555 | -1.50 | 0.931 |
| | | AN_Víðidalsá:LB_TH | 0.892 | 26.896 | 82.660 | -2.09 | 0.982 |
| | | AN_Víðidalsá:LB_UL | 0.881 | 28.246 | 84.862 | -1.97 | 0.976 |
| | | AN_Víðidalsá:PI_TH | 0.845 | 32.330 | 82.402 | -1.31 | 0.903 |
| | | AN_Víðidalsá:PI_UL | 0.752 | 41.260 | 104.584 | -1.66 | 0.952 |
| | | AN_Víðidalsá:PL_TH | 0.721 | 43.882 | 82.371 | -0.40 | 0.653 |
| | | AN_Víðidalsá:PL_UL | 0.037 | 87.899 | 89.820 | 1.62 | 0.061 |
| | | AN_Víðidalsá:SB_TH | 0.748 | 41.551 | 81.062 | -0.46 | 0.666 |
| | Vífisdalsá to lake morphs | AN_Vífilsdalsá:Hest | -0.351 | 110.535 | 88.220 | 2.44 | 0.008 |
| | | AN_Vífilsdalsá:Hit | -0.376 | 112.059 | 109.478 | 1.72 | 0.037 |
| | | AN_Vífilsdalsá:LB_TH | -0.361 | 111.165 | 91.567 | 2.29 | ***0.007** |
| | | AN_Vífilsdalsá:LB_UL | -0.113 | 96.488 | 93.792 | 1.79 | 0.041 |
| | | AN_Vífilsdalsá:PI_TH | -0.031 | 91.757 | 87.705 | 1.75 | 0.039 |
| | | AN_Vífilsdalsá:PI_UL | -0.349 | 110.451 | 109.177 | 1.69 | 0.045 |
| | | AN_Vífilsdalsá:PL_TH | -0.104 | 95.965 | 89.696 | 1.94 | 0.029 |
| | | AN_Vífilsdalsá:PL_UL | 0.214 | 77.662 | 96.489 | 0.88 | 0.189 |
| | | AN_Vífilsdalsá:SB_TH | -0.574 | 125.009 | 89.880 | 2.57 | ***0.003** |
| | Hestvatn and Hítarvatn | Hest:Hit | 0.779 | 38.797 | 103.037 | -1.63 | 0.953 |
| | | Hest:LB_TH | 0.947 | 18.655 | 54.807 | -2.23 | 0.985 |
| | | Hest:LB_UL | 0.885 | 27.736 | 58.632 | -1.04 | 0.850 |
| | | Hest:PI_TH | 0.778 | 38.912 | 48.577 | 0.79 | 0.214 |
| | | Hest:PI_UL | 0.853 | 31.489 | 96.283 | -2.00 | 0.982 |
| | | Hest:PL_TH | 0.795 | 37.343 | 50.857 | 0.48 | 0.322 |
| | | Hest:PL_UL | 0.096 | 84.503 | 72.734 | 2.11 | 0.012 |
| | | Hest:SB_TH | 0.896 | 26.325 | 47.236 | -0.49 | 0.683 |
| | | Hit:LB_TH | 0.804 | 36.465 | 103.926 | -1.74 | 0.963 |
| | | Hit:LB_UL | 0.629 | 51.040 | 103.252 | -0.88 | 0.800 |
| | | Hit:PI_TH | 0.372 | 68.183 | 104.246 | 0.20 | 0.417 |
| | | Hit:PI_UL | 0.789 | 37.879 | 112.274 | -2.20 | 0.991 |
| | | Hit:PL_TH | 0.501 | 59.947 | 103.605 | -0.21 | 0.584 |
| | | Hit:PL_UL | 0.008 | 89.568 | 104.224 | 0.99 | 0.182 |
| | | Hit:SB_TH | 0.813 | 35.653 | 104.664 | -1.72 | 0.960 |
| | Þingvallavatn | LB_TH:PI_TH | 0.738 | 42.472 | 51.315 | 1.01 | 0.161 |

| | | | | | | | |
|---|---|---|---|---|---|---|---|
| | | LB_TH:PL_TH | 0.738 | 42.456 | 53.082 | 0.81 | 0.214 |
| | | LB_TH:SB_TH | 0.883 | 28.007 | 49.471 | -0.42 | 0.660 |
| | | PI_TH:PL_TH | 0.742 | 42.130 | 45.478 | 1.32 | 0.090 |
| | | PI_TH:SB_TH | 0.622 | 51.514 | 40.734 | 2.46 | 0.011 |
| | | PL_TH:SB_TH | 0.717 | 44.232 | 42.450 | 1.74 | 0.038 |
| | Úlfljótsvatn | LB_UL:PI_UL | 0.729 | 43.168 | 98.676 | -1.15 | 0.869 |
| | | LB_UL:PL_UL | 0.176 | 79.841 | 76.103 | 1.82 | 0.032 |
| | | PI_UL:PL_UL | 0.191 | 78.982 | 101.056 | 0.69 | 0.266 |
| | Across Þingvallavatn and Úlfljótsvatn | LB_TH:LB_UL | 0.848 | 32.008 | 60.436 | -0.72 | 0.764 |
| | | LB_TH:PI_UL | 0.874 | 29.110 | 100.396 | -2.18 | 0.991 |
| | | LB_TH:PL_UL | 0.108 | 83.814 | 72.648 | 2.17 | 0.014 |
| | | LB_UL:PI_TH | 0.822 | 34.721 | 55.007 | 0.05 | 0.479 |
| | | LB_UL:PL_TH | 0.773 | 39.352 | 57.132 | 0.30 | 0.381 |
| | | LB_UL:SB_TH | 0.748 | 41.585 | 53.532 | 0.77 | 0.211 |
| | | PI_TH:PI_UL | 0.638 | 50.338 | 97.208 | -0.50 | 0.682 |
| | | PI_TH:PL_UL | 0.204 | 78.214 | 70.675 | 2.01 | 0.019 |
| | | PI_UL:PL_TH | 0.610 | 52.386 | 97.857 | -0.41 | 0.654 |
| | | PI_UL:SB_TH | 0.841 | 32.769 | 97.835 | -1.79 | 0.970 |
| | | PL_TH:PL_UL | 0.261 | 74.875 | 70.849 | 1.82 | 0.035 |
| | | PL_UL:SB_TH | 0.143 | 81.757 | 69.607 | 2.13 | 0.014 |
| Premaxilla | AN populations | AN_Brúará:AN_Gufudalsá | 0.753 | 41.110 | 104.602 | -2.13 | 0.985 |
| | | AN_Brúará:AN_Lambeyrarkvísl | 0.792 | 37.584 | 109.368 | -2.59 | 0.995 |
| | | AN_Brúará:AN_Víðidalsá | 0.715 | 44.349 | 103.968 | -1.66 | 0.950 |
| | | AN_Brúará:AN_Vífilsdalsá | 0.446 | 63.539 | 108.246 | -0.87 | 0.803 |
| | | AN_Gufudalsá:AN_Lambeyrarkvísl | 0.742 | 42.080 | 100.057 | -1.82 | 0.965 |
| | | AN_Gufudalsá:AN_Víðidalsá | 0.645 | 49.856 | 81.175 | -0.43 | 0.662 |
| | | AN_Gufudalsá:AN_Vífilsdalsá | 0.674 | 47.619 | 97.088 | -1.39 | 0.914 |
| | | AN_Lambeyrarkvísl:AN_Víðidalsá | 0.575 | 54.931 | 96.743 | -0.76 | 0.769 |
| | | AN_Lambeyrarkvísl:AN_Vífilsdalsá | 0.428 | 64.669 | 106.603 | -0.60 | 0.718 |
| | | AN_Víðidalsá:AN_Vífilsdalsá | 0.595 | 53.482 | 94.954 | -0.73 | 0.760 |
| | Brúará to lake morphs | AN_Brúará:Hest | 0.791 | 37.741 | 101.608 | -2.03 | 0.980 |
| | | AN_Brúará:Hit | 0.023 | 88.699 | 111.075 | 0.47 | 0.312 |
| | | AN_Brúará:LB_TH | 0.667 | 48.182 | 101.432 | -1.29 | 0.903 |
| | | AN_Brúará:LB_UL | 0.737 | 42.491 | 102.864 | -1.80 | 0.964 |
| | | AN_Brúará:PI_TH | 0.752 | 41.279 | 101.733 | -1.65 | 0.960 |
| | | AN_Brúará:PI_UL | 0.501 | 59.966 | 106.745 | -0.98 | 0.828 |
| | | AN_Brúará:PL_TH | 0.309 | 71.993 | 101.804 | 0.21 | 0.425 |
| | | AN_Brúará:PL_UL | 0.135 | 82.224 | 104.448 | 0.50 | 0.314 |
| | | AN_Brúará:SB_TH | -0.126 | 97.212 | 100.927 | 1.47 | 0.075 |
| | Gufudalsá to lake morphs | AN_Gufudalsá:Hest | 0.774 | 39.254 | 73.827 | -0.97 | 0.832 |
| | | AN_Gufudalsá:Hit | 0.260 | 74.959 | 100.776 | 0.18 | 0.428 |
| | | AN_Gufudalsá:LB_TH | 0.900 | 25.872 | 73.700 | -2.42 | 0.996 |
| | | AN_Gufudalsá:LB_UL | 0.950 | 18.137 | 78.560 | -3.92 | 1.000 |
| | | AN_Gufudalsá:PI_TH | 0.698 | 45.721 | 72.969 | -0.22 | 0.578 |
| | | AN_Gufudalsá:PI_UL | 0.655 | 49.090 | 93.041 | -1.04 | 0.863 |
| | | AN_Gufudalsá:PL_TH | 0.378 | 67.813 | 76.121 | 1.23 | 0.112 |
| | | AN_Gufudalsá:PL_UL | 0.380 | 67.642 | 85.236 | 0.62 | 0.262 |
| | | AN_Gufudalsá:SB_TH | -0.003 | 90.165 | 71.906 | 2.64 | ***0.006** |
| | Lambeyrarkvísl to lake morphs | AN_Lambeyrarkvísl:Hest | 0.561 | 55.904 | 93.471 | -0.46 | 0.667 |
| | | AN_Lambeyrarkvísl:Hit | -0.086 | 94.957 | 109.156 | 0.93 | 0.177 |
| | | AN_Lambeyrarkvísl:LB_TH | 0.764 | 40.221 | 93.975 | -1.65 | 0.954 |
| | | AN_Lambeyrarkvísl:LB_UL | 0.771 | 39.578 | 98.911 | -1.88 | 0.971 |
| | | AN_Lambeyrarkvísl:PI_TH | 0.554 | 56.347 | 93.392 | -0.37 | 0.636 |
| | | AN_Lambeyrarkvísl:PI_UL | 0.498 | 60.150 | 103.475 | -0.78 | 0.765 |
| | | AN_Lambeyrarkvísl:PL_TH | 0.214 | 77.653 | 96.444 | 0.79 | 0.216 |
| | | AN_Lambeyrarkvísl:PL_UL | 0.255 | 75.227 | 99.147 | 0.33 | 0.369 |
| | | AN_Lambeyrarkvísl:SB_TH | -0.339 | 109.786 | 94.275 | 2.35 | ***0.006** |
| | Víðidalsá to lake morphs | AN_Víðidalsá:Hest | 0.862 | 30.485 | 66.157 | -1.57 | 0.938 |
| | | AN_Víðidalsá:Hit | 0.273 | 74.174 | 103.505 | 0.18 | 0.426 |
| | | AN_Víðidalsá:LB_TH | 0.631 | 50.846 | 67.795 | 0.42 | 0.350 |
| | | AN_Víðidalsá:LB_UL | 0.581 | 54.486 | 73.486 | 0.35 | 0.361 |

| | | | | | | | |
|---|---|---|---|---|---|---|---|
| | | AN_Víðidalsá:PI_TH | 0.797 | 37.181 | 64.258 | -0.49 | 0.688 |
| | | AN_Víðidalsá:PI_UL | 0.409 | 65.883 | 92.371 | 0.33 | 0.370 |
| | | AN_Víðidalsá:PL_TH | 0.352 | 69.364 | 67.692 | 1.78 | 0.041 |
| | | AN_Víðidalsá:PL_UL | 0.325 | 71.063 | 83.244 | 1.09 | 0.144 |
| | | AN_Víðidalsá:SB_TH | 0.174 | 79.978 | 64.960 | 2.52 | 0.009 |
| | Vífisdalsá to lake morphs | AN_Vífilsdalsá:Hest | 0.789 | 37.953 | 91.663 | -1.78 | 0.964 |
| | | AN_Vífilsdalsá:Hit | 0.230 | 76.682 | 109.892 | -0.01 | 0.500 |
| | | AN_Vífilsdalsá:LB_TH | 0.696 | 45.903 | 91.023 | -1.12 | 0.863 |
| | | AN_Vífilsdalsá:LB_UL | 0.672 | 47.772 | 93.529 | -1.12 | 0.864 |
| | | AN_Vífilsdalsá:PI_TH | 0.621 | 51.625 | 90.087 | -0.54 | 0.704 |
| | | AN_Vífilsdalsá:PI_UL | 0.321 | 71.284 | 102.703 | 0.04 | 0.475 |
| | | AN_Vífilsdalsá:PL_TH | 0.744 | 41.943 | 89.476 | -1.38 | 0.917 |
| | | AN_Vífilsdalsá:PL_UL | 0.505 | 59.639 | 95.465 | -0.46 | 0.660 |
| | | AN_Vífilsdalsá:SB_TH | 0.419 | 65.197 | 89.887 | 0.32 | 0.375 |
| | Hestvatn and Hítarvatn | Hest:Hit | 0.332 | 70.604 | 100.683 | 0.17 | 0.442 |
| | | Hest:LB_TH | 0.725 | 43.557 | 56.154 | 0.52 | 0.314 |
| | | Hest:LB_UL | 0.752 | 41.206 | 63.396 | -0.25 | 0.600 |
| | | Hest:PI_TH | 0.882 | 28.121 | 50.035 | -0.77 | 0.776 |
| | | Hest:PI_UL | 0.483 | 61.106 | 88.076 | 0.29 | 0.374 |
| | | Hest:PL_TH | 0.636 | 50.533 | 52.763 | 1.41 | 0.082 |
| | | Hest:PL_UL | 0.346 | 69.769 | 74.432 | 1.42 | 0.075 |
| | | Hest:SB_TH | 0.328 | 70.826 | 49.022 | 3.27 | ***0.001** |
| | | Hit:LB_TH | 0.227 | 76.887 | 100.208 | 0.52 | 0.306 |
| | | Hit:LB_UL | 0.240 | 76.128 | 103.730 | 0.37 | 0.357 |
| | | Hit:PI_TH | 0.074 | 85.784 | 100.210 | 1.01 | 0.164 |
| | | Hit:PI_UL | 0.181 | 79.588 | 106.558 | 0.26 | 0.400 |
| | | Hit:PL_TH | 0.345 | 69.834 | 101.595 | 0.19 | 0.422 |
| | | Hit:PL_UL | 0.507 | 59.554 | 103.695 | -0.82 | 0.788 |
| | | Hit:SB_TH | 0.674 | 47.608 | 100.878 | -1.18 | 0.885 |
| | Þingvallavatn | LB_TH:PI_TH | 0.615 | 52.071 | 51.109 | 1.72 | 0.041 |
| | | LB_TH:PL_TH | 0.491 | 60.585 | 56.892 | 1.97 | 0.023 |
| | | LB_TH:SB_TH | 0.080 | 85.415 | 50.713 | 4.04 | ***0.001** |
| | | PI_TH:PL_TH | 0.452 | 63.159 | 49.292 | 2.75 | ***0.004** |
| | | PI_TH:SB_TH | 0.115 | 83.405 | 45.548 | 4.30 | ***0.001** |
| | | PL_TH:SB_TH | 0.689 | 46.473 | 48.464 | 1.44 | 0.080 |
| | Úlfljótsvatn | LB_UL:PI_UL | 0.552 | 56.500 | 87.825 | -0.25 | 0.606 |
| | | LB_UL:PL_UL | 0.395 | 66.707 | 79.296 | 0.88 | 0.182 |
| | | PI_UL:PL_UL | 0.217 | 77.450 | 93.206 | 0.82 | 0.209 |
| | Across Þingvallavatn and Úlfljótsvatn | LB_TH:LB_UL | 0.884 | 27.830 | 64.257 | -1.75 | 0.965 |
| | | LB_TH:PI_UL | 0.653 | 49.218 | 87.336 | -0.53 | 0.701 |
| | | LB_TH:PL_UL | 0.538 | 57.485 | 76.057 | 0.57 | 0.280 |
| | | LB_UL:PI_TH | 0.686 | 46.678 | 61.732 | 0.50 | 0.305 |
| | | LB_UL:PL_TH | 0.368 | 68.396 | 63.650 | 1.91 | 0.032 |
| | | LB_UL:SB_TH | -0.053 | 93.040 | 61.576 | 3.46 | ***0.002** |
| | | PI_TH:PI_UL | 0.542 | 57.164 | 84.689 | 0.16 | 0.439 |
| | | PI_TH:PL_UL | 0.079 | 85.490 | 71.108 | 2.43 | 0.009 |
| | | PI_UL:PL_TH | 0.432 | 64.416 | 85.524 | 0.52 | 0.307 |
| | | PI_UL:SB_TH | 0.134 | 82.286 | 84.632 | 1.56 | 0.063 |
| | | PL_TH:PL_UL | 0.495 | 60.332 | 74.811 | 0.82 | 0.201 |
| | | PL_UL:SB_TH | 0.451 | 63.189 | 71.426 | 1.16 | 0.124 |
| Maxilla | AN populations | AN_Brúará:AN_Gufudalsá | 0.723 | 43.700 | 118.425 | -1.90 | 0.970 |
| | | AN_Brúará:AN_Lambeyrarkvísl | 0.542 | 57.178 | 122.111 | -1.23 | 0.872 |
| | | AN_Brúará:AN_Víðidalsá | -0.241 | 103.936 | 117.668 | 1.08 | 0.158 |
| | | AN_Brúará:AN_Vífilsdalsá | 0.242 | 76.011 | 121.356 | -0.32 | 0.622 |
| | | AN_Gufudalsá:AN_Lambeyrarkvísl | 0.538 | 57.464 | 111.096 | -0.93 | 0.810 |
| | | AN_Gufudalsá:AN_Víðidalsá | -0.426 | 115.204 | 98.459 | 2.28 | 0.009 |
| | | AN_Gufudalsá:AN_Vífilsdalsá | 0.276 | 73.967 | 106.315 | 0.13 | 0.468 |
| | | AN_Lambeyrarkvísl:AN_Víðidalsá | -0.289 | 106.773 | 113.765 | 1.42 | 0.096 |
| | | AN_Lambeyrarkvísl:AN_Vífilsdalsá | 0.083 | 85.234 | 116.747 | 0.30 | 0.404 |
| | | AN_Víðidalsá:AN_Vífilsdalsá | 0.110 | 83.657 | 108.633 | 0.55 | 0.296 |
| | | AN_Brúará:Hest | 0.535 | 57.645 | 113.035 | -1.07 | 0.856 |

| | | | | | | | |
|---|---|---|---|---|---|---|---|
| | Brúará to lake morphs | AN_Brúará:Hit | 0.351 | 69.428 | 120.387 | -0.61 | 0.725 |
| | | AN_Brúará:LB_TH | 0.841 | 32.707 | 111.085 | -2.63 | 0.999 |
| | | AN_Brúará:LB_UL | -0.313 | 108.249 | 112.764 | 1.44 | 0.082 |
| | | AN_Brúará:PI_TH | 0.111 | 83.613 | 114.265 | 0.29 | 0.379 |
| | | AN_Brúará:PI_UL | 0.243 | 75.908 | 121.396 | -0.35 | 0.624 |
| | | AN_Brúará:PL_TH | 0.392 | 66.904 | 111.412 | -0.54 | 0.697 |
| | | AN_Brúará:PL_UL | -0.093 | 95.326 | 115.223 | 0.73 | 0.236 |
| | | AN_Brúará:SB_TH | 0.360 | 68.883 | 109.768 | -0.42 | 0.665 |
| | Gufudalsá to lake morphs | AN_Gufudalsá:Hest | 0.411 | 65.752 | 84.874 | 0.59 | 0.284 |
| | | AN_Gufudalsá:Hit | 0.633 | 50.735 | 108.225 | -1.20 | 0.883 |
| | | AN_Gufudalsá:LB_TH | 0.483 | 61.110 | 89.017 | 0.23 | 0.413 |
| | | AN_Gufudalsá:LB_UL | 0.045 | 87.431 | 92.497 | 1.44 | 0.078 |
| | | AN_Gufudalsá:PI_TH | -0.168 | 99.693 | 86.417 | 2.20 | 0.009 |
| | | AN_Gufudalsá:PI_UL | 0.317 | 71.517 | 104.869 | 0.05 | 0.493 |
| | | AN_Gufudalsá:PL_TH | 0.224 | 77.080 | 88.667 | 1.10 | 0.142 |
| | | AN_Gufudalsá:PL_UL | 0.347 | 69.684 | 97.293 | 0.28 | 0.395 |
| | | AN_Gufudalsá:SB_TH | 0.519 | 58.728 | 83.103 | 0.32 | 0.379 |
| | Lambeyrarkvísl to lake morphs | AN_Lambeyrarkvísl:Hest | 0.373 | 68.071 | 101.997 | -0.11 | 0.543 |
| | | AN_Lambeyrarkvísl:Hit | 0.353 | 69.328 | 115.027 | -0.49 | 0.673 |
| | | AN_Lambeyrarkvísl:LB_TH | 0.430 | 64.561 | 104.334 | -0.32 | 0.625 |
| | | AN_Lambeyrarkvísl:LB_UL | 0.076 | 85.620 | 107.403 | 0.67 | 0.276 |
| | | AN_Lambeyrarkvísl:PI_TH | 0.077 | 85.599 | 102.633 | 0.84 | 0.212 |
| | | AN_Lambeyrarkvísl:PI_UL | 0.239 | 76.200 | 114.515 | -0.10 | 0.539 |
| | | AN_Lambeyrarkvísl:PL_TH | 0.271 | 74.302 | 105.117 | 0.22 | 0.418 |
| | | AN_Lambeyrarkvísl:PL_UL | -0.110 | 96.331 | 110.438 | 1.02 | 0.166 |
| | | AN_Lambeyrarkvísl:SB_TH | 0.263 | 74.779 | 100.825 | 0.36 | 0.364 |
| | Víðidalsá to lake morphs | AN_Víðidalsá:Hest | 0.526 | 58.258 | 89.603 | 0.06 | 0.480 |
| | | AN_Víðidalsá:Hit | -0.301 | 107.502 | 108.225 | 1.57 | 0.055 |
| | | AN_Víðidalsá:LB_TH | 0.118 | 83.230 | 90.632 | 1.31 | 0.108 |
| | | AN_Víðidalsá:LB_UL | 0.502 | 59.850 | 92.251 | -0.03 | 0.509 |
| | | AN_Víðidalsá:PI_TH | 0.764 | 40.216 | 88.511 | -1.18 | 0.868 |
| | | AN_Víðidalsá:PI_UL | 0.608 | 52.557 | 104.629 | -1.07 | 0.839 |
| | | AN_Víðidalsá:PL_TH | 0.268 | 74.451 | 90.425 | 0.92 | 0.189 |
| | | AN_Víðidalsá:PL_UL | 0.230 | 76.715 | 99.842 | 0.64 | 0.272 |
| | | AN_Víðidalsá:SB_TH | -0.120 | 96.916 | 85.375 | 2.15 | 0.019 |
| | Vífisdalsá to lake morphs | AN_Vífilsdalsá:Hest | 0.331 | 70.641 | 99.180 | 0.29 | 0.389 |
| | | AN_Vífilsdalsá:Hit | 0.480 | 61.319 | 113.693 | -0.75 | 0.762 |
| | | AN_Vífilsdalsá:LB_TH | 0.404 | 66.156 | 99.229 | 0.01 | 0.495 |
| | | AN_Vífilsdalsá:LB_UL | 0.263 | 74.780 | 103.893 | 0.33 | 0.372 |
| | | AN_Vífilsdalsá:PI_TH | 0.178 | 79.770 | 96.749 | 0.86 | 0.210 |
| | | AN_Vífilsdalsá:PI_UL | 0.469 | 62.024 | 113.074 | -0.72 | 0.761 |
| | | AN_Vífilsdalsá:PL_TH | 0.694 | 46.028 | 99.542 | -1.23 | 0.887 |
| | | AN_Vífilsdalsá:PL_UL | 0.348 | 69.632 | 105.825 | -0.07 | 0.525 |
| | | AN_Vífilsdalsá:SB_TH | 0.477 | 61.532 | 95.149 | -0.13 | 0.548 |
| | Hestvatn and Hítarvatn | Hest:Hit | 0.153 | 81.170 | 99.535 | 0.80 | 0.218 |
| | | Hest:LB_TH | 0.736 | 42.632 | 70.467 | -0.13 | 0.531 |
| | | Hest:LB_UL | 0.471 | 61.925 | 76.298 | 0.84 | 0.201 |
| | | Hest:PI_TH | 0.691 | 46.320 | 66.664 | 0.34 | 0.364 |
| | | Hest:PI_UL | 0.822 | 34.683 | 96.310 | -1.99 | 0.981 |
| | | Hest:PL_TH | 0.632 | 50.819 | 70.452 | 0.49 | 0.305 |
| | | Hest:PL_UL | 0.412 | 65.651 | 86.919 | 0.59 | 0.296 |
| | | Hest:SB_TH | 0.369 | 68.354 | 61.634 | 2.01 | 0.016 |
| | | Hit:LB_TH | 0.254 | 75.299 | 99.139 | 0.48 | 0.325 |
| | | Hit:LB_UL | 0.077 | 85.566 | 101.614 | 0.88 | 0.203 |
| | | Hit:PI_TH | -0.204 | 101.789 | 99.791 | 1.76 | 0.043 |
| | | Hit:PI_UL | 0.225 | 76.977 | 110.871 | 0.02 | 0.492 |
| | | Hit:PL_TH | 0.314 | 71.682 | 100.965 | 0.23 | 0.412 |
| | | Hit:PL_UL | 0.338 | 70.229 | 106.878 | -0.07 | 0.542 |
| | | Hit:SB_TH | 0.531 | 57.937 | 96.807 | -0.40 | 0.650 |
| | Þingvallavatn | LB_TH:PI_TH | 0.549 | 56.670 | 66.306 | 1.02 | 0.179 |
| | | LB_TH:PL_TH | 0.709 | 44.828 | 71.106 | 0.00 | 0.496 |

| | | | | | | | |
|---|---|---|---|---|---|---|---|
| | | LB_TH:SB_TH | 0.419 | 65.204 | 63.121 | 1.76 | 0.034 |
| | | PI_TH:PL_TH | 0.570 | 55.251 | 65.621 | 0.94 | 0.185 |
| | | PI_TH:SB_TH | 0.102 | 84.146 | 58.924 | 3.10 | ***0.001** |
| | | PL_TH:SB_TH | 0.621 | 51.592 | 61.371 | 0.95 | 0.179 |
| | Úlfljótsvatn | LB_UL:PI_UL | 0.663 | 48.434 | 98.273 | -1.03 | 0.839 |
| | | LB_UL:PL_UL | 0.752 | 41.204 | 89.505 | -1.14 | 0.870 |
| | | PI_UL:PL_UL | 0.539 | 57.414 | 102.903 | -0.67 | 0.729 |
| | Across Þingvallavatn and Úlfljótsvatn | LB_TH:LB_UL | -0.063 | 93.615 | 77.924 | 2.40 | ***0.007** |
| | | LB_TH:PI_UL | 0.500 | 60.019 | 96.891 | -0.28 | 0.624 |
| | | LB_TH:PL_UL | 0.039 | 87.791 | 86.254 | 1.78 | 0.041 |
| | | LB_UL:PI_TH | 0.391 | 66.981 | 75.827 | 1.24 | 0.121 |
| | | LB_UL:PL_TH | 0.324 | 71.110 | 77.342 | 1.32 | 0.102 |
| | | LB_UL:SB_TH | 0.399 | 66.485 | 71.996 | 1.40 | 0.081 |
| | | PI_TH:PI_UL | 0.692 | 46.229 | 95.081 | -1.08 | 0.847 |
| | | PI_TH:PL_UL | 0.109 | 83.718 | 83.088 | 1.66 | 0.048 |
| | | PI_UL:PL_TH | 0.530 | 57.995 | 95.794 | -0.35 | 0.638 |
| | | PI_UL:SB_TH | 0.313 | 71.766 | 93.144 | 0.56 | 0.291 |
| | | PL_TH:PL_UL | 0.340 | 70.147 | 87.258 | 0.84 | 0.213 |
| | | PL_UL:SB_TH | 0.580 | 54.542 | 81.136 | 0.19 | 0.423 |
| Supramaxilla | AN populations | AN_Brúará:AN_Gufudalsá | -0.121 | 96.973 | 132.539 | 0.50 | 0.330 |
| | | AN_Brúará:AN_Lambeyrarkvísl | 0.496 | 60.267 | 136.249 | -0.87 | 0.793 |
| | | AN_Brúará:AN_Víðidalsá | 0.699 | 45.653 | 129.309 | -1.35 | 0.905 |
| | | AN_Brúará:AN_Vífilsdalsá | 0.606 | 52.669 | 131.936 | -1.18 | 0.859 |
| | | AN_Gufudalsá:AN_Lambeyrarkvísl | -0.552 | 123.534 | 129.534 | 1.44 | 0.078 |
| | | AN_Gufudalsá:AN_Víðidalsá | -0.354 | 110.703 | 119.545 | 1.32 | 0.111 |
| | | AN_Gufudalsá:AN_Vífilsdalsá | 0.205 | 78.164 | 130.106 | -0.08 | 0.541 |
| | | AN_Lambeyrarkvísl:AN_Víðidalsá | 0.572 | 55.094 | 127.620 | -0.85 | 0.780 |
| | | AN_Lambeyrarkvísl:AN_Vífilsdalsá | -0.273 | 105.828 | 133.874 | 0.75 | 0.251 |
| | | AN_Víðidalsá:AN_Vífilsdalsá | 0.406 | 66.053 | 127.613 | -0.46 | 0.666 |
| | Brúará to lake morphs | AN_Brúará:Hest | 0.059 | 86.600 | 127.111 | 0.32 | 0.384 |
| | | AN_Brúará:Hit | 0.058 | 86.693 | 136.726 | 0.04 | 0.479 |
| | | AN_Brúará:LB_TH | 0.789 | 37.866 | 125.921 | -1.67 | 0.944 |
| | | AN_Brúará:LB_UL | -0.053 | 93.017 | 128.510 | 0.50 | 0.346 |
| | | AN_Brúará:PI_TH | 0.822 | 34.684 | 123.962 | -1.80 | 0.964 |
| | | AN_Brúará:PI_UL | 0.403 | 66.213 | 131.203 | -0.63 | 0.723 |
| | | AN_Brúará:PL_TH | 0.434 | 64.301 | 124.563 | -0.53 | 0.694 |
| | | AN_Brúará:PL_UL | 0.563 | 55.737 | 127.060 | -0.99 | 0.820 |
| | | AN_Brúará:SB_TH | -0.601 | 126.941 | 123.780 | 1.71 | 0.040 |
| | Gufudalsá to lake morphs | AN_Gufudalsá:Hest | -0.168 | 99.643 | 112.845 | 1.21 | 0.117 |
| | | AN_Gufudalsá:Hit | 0.149 | 81.449 | 130.962 | 0.03 | 0.491 |
| | | AN_Gufudalsá:LB_TH | -0.124 | 97.106 | 112.496 | 1.09 | 0.152 |
| | | AN_Gufudalsá:LB_UL | 0.049 | 87.181 | 117.328 | 0.63 | 0.270 |
| | | AN_Gufudalsá:PI_TH | 0.026 | 88.483 | 110.619 | 0.87 | 0.202 |
| | | AN_Gufudalsá:PI_UL | 0.804 | 36.473 | 125.594 | -1.68 | 0.960 |
| | | AN_Gufudalsá:PL_TH | 0.803 | 36.580 | 110.193 | -1.44 | 0.914 |
| | | AN_Gufudalsá:PL_UL | 0.562 | 55.797 | 121.939 | -0.64 | 0.729 |
| | | AN_Gufudalsá:SB_TH | -0.109 | 96.234 | 108.363 | 1.19 | 0.133 |
| | Lambeyrarkvísl to lake morphs | AN_Lambeyrarkvísl:Hest | 0.450 | 63.273 | 119.963 | -0.47 | 0.681 |
| | | AN_Lambeyrarkvísl:Hit | -0.682 | 132.988 | 132.500 | 1.62 | 0.047 |
| | | AN_Lambeyrarkvísl:LB_TH | 0.723 | 43.689 | 123.872 | -1.35 | 0.903 |
| | | AN_Lambeyrarkvísl:LB_UL | -0.115 | 96.622 | 123.960 | 0.70 | 0.265 |
| | | AN_Lambeyrarkvísl:PI_TH | 0.534 | 57.711 | 118.788 | -0.69 | 0.726 |
| | | AN_Lambeyrarkvísl:PI_UL | -0.253 | 104.668 | 133.419 | 0.67 | 0.269 |
| | | AN_Lambeyrarkvísl:PL_TH | -0.345 | 110.159 | 119.058 | 1.32 | 0.095 |
| | | AN_Lambeyrarkvísl:PL_UL | -0.303 | 107.643 | 128.892 | 1.01 | 0.163 |
| | | AN_Lambeyrarkvísl:SB_TH | -0.232 | 103.402 | 115.886 | 1.11 | 0.136 |
| | Víðidalsá to lake morphs | AN_Víðidalsá:Hest | 0.503 | 59.784 | 109.995 | -0.24 | 0.604 |
| | | AN_Víðidalsá:Hit | 0.070 | 85.967 | 130.175 | 0.20 | 0.428 |
| | | AN_Víðidalsá:LB_TH | 0.859 | 30.764 | 108.520 | -1.68 | 0.952 |
| | | AN_Víðidalsá:LB_UL | 0.525 | 58.314 | 114.154 | -0.40 | 0.657 |
| | | AN_Víðidalsá:PI_TH | 0.821 | 34.786 | 108.389 | -1.41 | 0.910 |

| | | | | | | | |
|---|---|---|---|---|---|---|---|
| | | AN_Víðidalsá:PI_UL | -0.119 | 96.860 | 125.366 | 0.64 | 0.286 |
| | | AN_Víðidalsá:PL_TH | 0.061 | 86.476 | 108.454 | 0.86 | 0.197 |
| | | AN_Víðidalsá:PL_UL | 0.306 | 72.190 | 118.948 | 0.05 | 0.489 |
| | | AN_Víðidalsá:SB_TH | 0.020 | 88.879 | 105.302 | 1.08 | 0.156 |
| | Vífisdalsá to lake morphs | AN_Vífilsdalsá:Hest | -0.025 | 91.418 | 122.401 | 0.57 | 0.302 |
| | | AN_Vífilsdalsá:Hit | 0.661 | 48.591 | 134.699 | -1.28 | 0.894 |
| | | AN_Vífilsdalsá:LB_TH | 0.339 | 70.160 | 122.542 | -0.24 | 0.589 |
| | | AN_Vífilsdalsá:LB_UL | 0.161 | 80.738 | 124.442 | 0.13 | 0.451 |
| | | AN_Vífilsdalsá:PI_TH | 0.493 | 60.473 | 119.360 | -0.58 | 0.708 |
| | | AN_Vífilsdalsá:PI_UL | 0.455 | 62.957 | 133.446 | -0.68 | 0.745 |
| | | AN_Vífilsdalsá:PL_TH | 0.578 | 54.669 | 123.029 | -0.77 | 0.778 |
| | | AN_Vífilsdalsá:PL_UL | 0.757 | 40.820 | 122.341 | -1.47 | 0.929 |
| | | AN_Vífilsdalsá:SB_TH | -0.185 | 100.681 | 118.193 | 1.03 | 0.163 |
| | Hestvatn and Hítarvatn | Hest:Hit | -0.405 | 113.900 | 124.395 | 1.25 | 0.117 |
| | | Hest:LB_TH | 0.584 | 54.253 | 96.250 | -0.05 | 0.523 |
| | | Hest:LB_UL | 0.545 | 57.008 | 105.302 | -0.10 | 0.539 |
| | | Hest:PI_TH | 0.550 | 56.603 | 93.181 | 0.16 | 0.439 |
| | | Hest:PI_UL | -0.252 | 104.586 | 117.196 | 1.21 | 0.137 |
| | | Hest:PL_TH | -0.184 | 100.588 | 95.172 | 1.81 | 0.034 |
| | | Hest:PL_UL | -0.171 | 99.874 | 105.611 | 1.47 | 0.078 |
| | | Hest:SB_TH | 0.595 | 53.502 | 87.435 | 0.23 | 0.410 |
| | | Hit:LB_TH | -0.284 | 106.528 | 124.305 | 1.02 | 0.173 |
| | | Hit:LB_UL | 0.315 | 71.663 | 125.880 | -0.32 | 0.602 |
| | | Hit:PI_TH | -0.130 | 97.485 | 124.420 | 0.74 | 0.261 |
| | | Hit:PI_UL | 0.080 | 85.427 | 132.044 | 0.09 | 0.472 |
| | | Hit:PL_TH | 0.296 | 72.773 | 119.699 | -0.14 | 0.563 |
| | | Hit:PL_UL | 0.597 | 53.326 | 128.264 | -1.02 | 0.835 |
| | | Hit:SB_TH | 0.098 | 84.387 | 121.032 | 0.31 | 0.397 |
| | Þingvallavatn | LB_TH:PI_TH | 0.951 | 17.975 | 97.213 | -2.23 | 0.993 |
| | | LB_TH:PL_TH | 0.238 | 76.239 | 95.394 | 0.94 | 0.182 |
| | | LB_TH:SB_TH | -0.194 | 101.195 | 90.549 | 2.02 | 0.017 |
| | | PI_TH:PL_TH | 0.447 | 63.429 | 90.733 | 0.56 | 0.307 |
| | | PI_TH:SB_TH | -0.170 | 99.783 | 83.920 | 2.18 | 0.012 |
| | | PL_TH:SB_TH | -0.397 | 113.394 | 83.945 | 2.77 | ***0.001** |
| | Úlfljótsvatn | LB_UL:PI_UL | -0.209 | 102.064 | 125.143 | 0.95 | 0.184 |
| | | LB_UL:PL_UL | 0.269 | 74.369 | 111.949 | 0.32 | 0.380 |
| | | PI_UL:PL_UL | 0.689 | 46.470 | 125.728 | -1.22 | 0.885 |
| | Across Þingvallavatn and Úlfljótsvatn | LB_TH:LB_UL | 0.248 | 75.626 | 106.520 | 0.66 | 0.252 |
| | | LB_TH:PI_UL | 0.190 | 79.019 | 118.967 | 0.23 | 0.418 |
| | | LB_TH:PL_UL | 0.308 | 72.038 | 106.027 | 0.33 | 0.383 |
| | | LB_UL:PI_TH | 0.355 | 69.214 | 98.078 | 0.51 | 0.321 |
| | | LB_UL:PL_TH | 0.106 | 83.897 | 99.959 | 1.06 | 0.151 |
| | | LB_UL:SB_TH | 0.690 | 46.404 | 95.115 | -0.42 | 0.669 |
| | | PI_TH:PI_UL | 0.348 | 69.661 | 114.334 | -0.08 | 0.530 |
| | | PI_TH:PL_UL | 0.504 | 59.767 | 104.340 | 0.00 | 0.503 |
| | | PI_UL:PL_TH | 0.935 | 20.845 | 115.061 | -2.53 | 0.999 |
| | | PI_UL:SB_TH | -0.575 | 125.122 | 115.274 | 1.98 | 0.024 |
| | | PL_TH:PL_UL | 0.854 | 31.361 | 105.990 | -1.53 | 0.934 |
| | | PL_UL:SB_TH | -0.277 | 106.078 | 99.630 | 1.85 | 0.029 |

R = slope of vector correlation; UCL = upper confidence level, Z = Z-statistic; P = P-value, ≤ 0.008 in bold with asterisk.

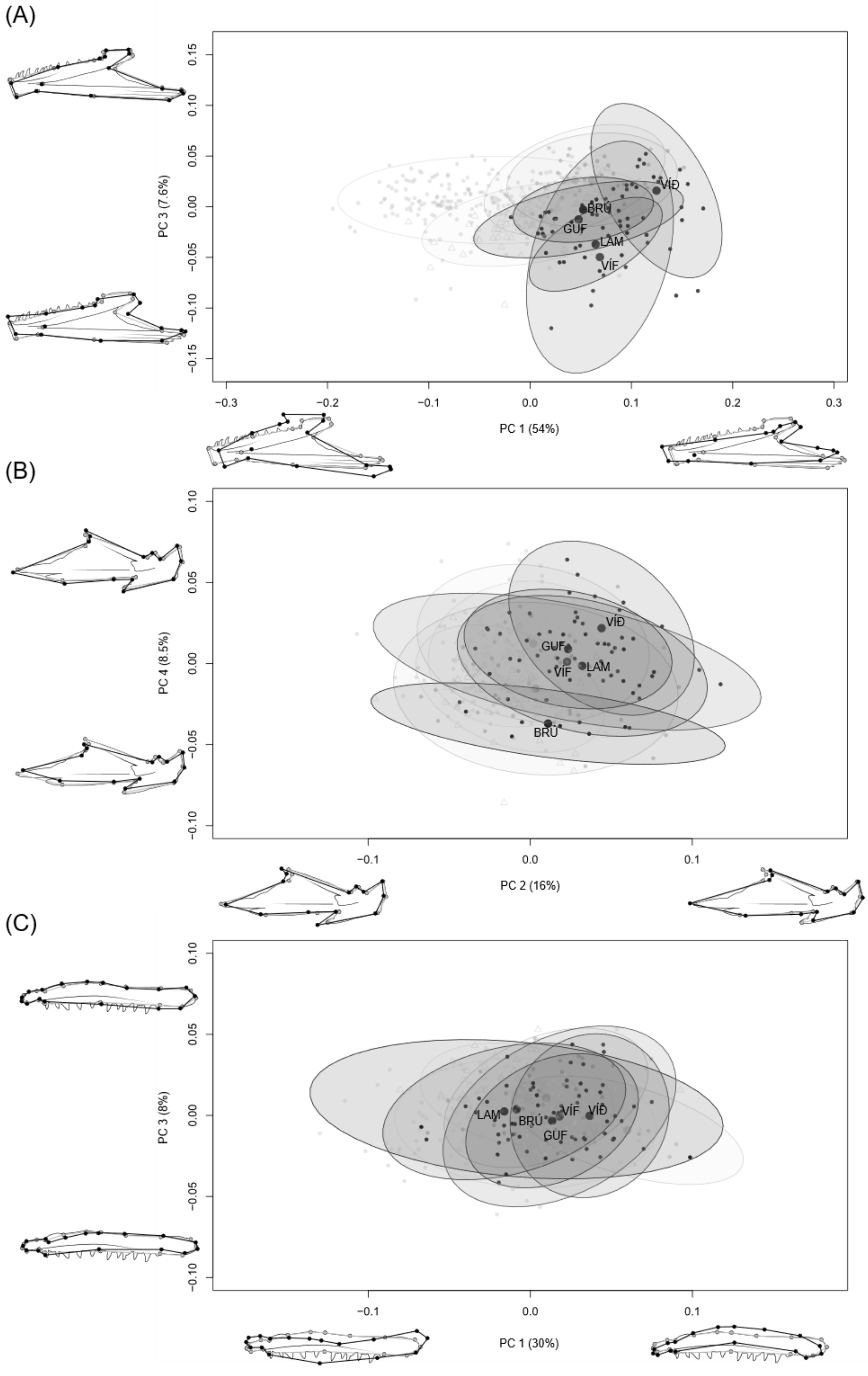


***Supplementary material 18****: Variation within the AN-charr populations. PC-plot with deformation grids showing variation in shape for three bones A) dentary, B) articular-angular, C) maxilla within the five AN-charr populations. Note that the effects of size have not been removed for the A) dentary due to differences in allometric trajectory between the*

*groups. Size effects have been removed for the B) articular-angular and C) maxilla plots. None of the deformation grids are magnified, grey outline represents mean shape and black lines the two extremes for each PC. Each dot represents an individual and the ellipses 95% CI for the distribution by group (large dot represents the mean shape of each group in these dimensions). The percentage of total variation which each PC explains is written next to the PC.*

***Supplementary material 19****: Results from pairwise tests on population differences in mean bone shape, from the best fit ANCOVA models for each bone. AN-charr treated as five separate populations.*

| Bone | System | Pairs | D | UCL | Z | P-value |
|---|---|---|---|---|---|---|
| Dentary[1] | AN populations | AN_Brúará:AN_Gufudalsá | 0.110 | 0.297 | -2.33 | 0.992 |
| | | AN_Brúará:AN_Lambeyrarkvísl | 0.202 | 0.332 | -0.52 | 0.710 |
| | | AN_Brúará:AN_Víðidalsá | 0.218 | 0.298 | 0.51 | 0.307 |
| | | AN_Brúará:AN_Vífilsdalsá | 0.158 | 0.328 | -1.33 | 0.903 |
| | | AN_Gufudalsá:AN_Lambeyrarkvísl | 0.255 | 0.244 | 1.86 | 0.033 |
| | | AN_Gufudalsá:AN_Víðidalsá | 0.259 | 0.155 | 3.96 | ***0.001** |
| | | AN_Gufudalsá:AN_Vífilsdalsá | 0.208 | 0.226 | 1.26 | 0.106 |
| | | AN_Lambeyrarkvísl:AN_Víðidalsá | 0.231 | 0.235 | 1.57 | 0.058 |
| | | AN_Lambeyrarkvísl:AN_Vífilsdalsá | 0.164 | 0.284 | -0.64 | 0.730 |
| | | AN_Víðidalsá:AN_Vífilsdalsá | 0.150 | 0.219 | 0.12 | 0.448 |
| | Brúará to lake morphs | AN_Brúará:Hest | 0.139 | 0.282 | -1.15 | 0.879 |
| | | AN_Brúará:Hit | 0.153 | 0.341 | -1.51 | 0.937 |
| | | AN_Brúará:LB_TH | 0.109 | 0.288 | -2.05 | 0.981 |
| | | AN_Brúará:LB_UL | 0.139 | 0.287 | -1.20 | 0.887 |
| | | AN_Brúará:PI_TH | 0.174 | 0.277 | -0.19 | 0.579 |
| | | AN_Brúará:PI_UL | 0.166 | 0.309 | -0.91 | 0.824 |
| | | AN_Brúará:PL_TH | 0.108 | 0.286 | -2.09 | 0.981 |
| | | AN_Brúará:PL_UL | 0.120 | 0.297 | -1.85 | 0.970 |
| | | AN_Brúará:SB_TH | 0.141 | 0.280 | -1.02 | 0.862 |
| | Gufudalsá to lake morphs | AN_Gufudalsá:Hest | 0.152 | 0.132 | 2.29 | 0.010 |
| | | AN_Gufudalsá:Hit | 0.147 | 0.232 | -0.30 | 0.621 |
| | | AN_Gufudalsá:LB_TH | 0.126 | 0.133 | 1.43 | 0.080 |
| | | AN_Gufudalsá:LB_UL | 0.166 | 0.142 | 2.46 | ***0.004** |
| | | AN_Gufudalsá:PI_TH | 0.215 | 0.126 | 4.20 | ***0.001** |
| | | AN_Gufudalsá:PI_UL | 0.182 | 0.200 | 1.21 | 0.113 |
| | | AN_Gufudalsá:PL_TH | 0.143 | 0.132 | 1.99 | 0.023 |
| | | AN_Gufudalsá:PL_UL | 0.121 | 0.155 | 0.59 | 0.285 |
| | | AN_Gufudalsá:SB_TH | 0.181 | 0.128 | 3.34 | ***0.001** |
| | Lambeyrarkvísl to lake morphs | AN_Lambeyrarkvísl:Hest | 0.156 | 0.218 | 0.19 | 0.423 |
| | | AN_Lambeyrarkvísl:Hit | 0.203 | 0.288 | 0.17 | 0.443 |
| | | AN_Lambeyrarkvísl:LB_TH | 0.200 | 0.223 | 1.21 | 0.127 |
| | | AN_Lambeyrarkvísl:LB_UL | 0.144 | 0.229 | -0.19 | 0.582 |
| | | AN_Lambeyrarkvísl:PI_TH | 0.152 | 0.224 | 0.18 | 0.426 |
| | | AN_Lambeyrarkvísl:PI_UL | 0.174 | 0.260 | -0.13 | 0.541 |
| | | AN_Lambeyrarkvísl:PL_TH | 0.146 | 0.225 | -0.01 | 0.502 |
| | | AN_Lambeyrarkvísl:PL_UL | 0.193 | 0.233 | 0.85 | 0.198 |
| | | AN_Lambeyrarkvísl:SB_TH | 0.116 | 0.219 | -0.92 | 0.817 |
| | Víðidalsá to lake morphs | AN_Víðidalsá:Hest | 0.194 | 0.116 | 3.88 | ***0.001** |
| | | AN_Víðidalsá:Hit | 0.282 | 0.225 | 2.48 | 0.011 |
| | | AN_Víðidalsá:LB_TH | 0.181 | 0.118 | 3.81 | ***0.001** |
| | | AN_Víðidalsá:LB_UL | 0.192 | 0.129 | 3.31 | ***0.001** |
| | | AN_Víðidalsá:PI_TH | 0.168 | 0.113 | 3.29 | ***0.001** |
| | | AN_Víðidalsá:PI_UL | 0.210 | 0.188 | 2.22 | 0.018 |
| | | AN_Víðidalsá:PL_TH | 0.215 | 0.116 | 4.65 | ***0.001** |

| | | | | | | |
|---|---|---|---|---|---|---|
| | | AN_Víðidalsá:PL_UL | 0.204 | 0.145 | 3.09 | ***0.001** |
| | | AN_Víðidalsá:SB_TH | 0.212 | 0.112 | 4.49 | ***0.001** |
| | Vífisdalsá to lake morphs | AN_Vífilsdalsá:Hest | 0.143 | 0.203 | 0.23 | 0.410 |
| | | AN_Vífilsdalsá:Hit | 0.192 | 0.271 | 0.16 | 0.439 |
| | | AN_Vífilsdalsá:LB_TH | 0.135 | 0.205 | 0.03 | 0.485 |
| | | AN_Vífilsdalsá:LB_UL | 0.132 | 0.210 | -0.18 | 0.576 |
| | | AN_Vífilsdalsá:PI_TH | 0.160 | 0.198 | 0.73 | 0.246 |
| | | AN_Vífilsdalsá:PI_UL | 0.181 | 0.245 | 0.35 | 0.372 |
| | | AN_Vífilsdalsá:PL_TH | 0.147 | 0.203 | 0.35 | 0.376 |
| | | AN_Vífilsdalsá:PL_UL | 0.140 | 0.220 | -0.19 | 0.577 |
| | | AN_Vífilsdalsá:SB_TH | 0.147 | 0.198 | 0.42 | 0.334 |
| | Hestvatn and Hítarvatn | Hest:Hit | 0.118 | 0.217 | -0.67 | 0.751 |
| | | Hest:LB_TH | 0.099 | 0.087 | 2.25 | 0.009 |
| | | Hest:LB_UL | 0.062 | 0.098 | -0.28 | 0.610 |
| | | Hest:PI_TH | 0.103 | 0.079 | 2.85 | ***0.003** |
| | | Hest:PI_UL | 0.101 | 0.171 | -0.57 | 0.712 |
| | | Hest:PL_TH | 0.070 | 0.084 | 0.82 | 0.203 |
| | | Hest:PL_UL | 0.086 | 0.117 | 0.28 | 0.384 |
| | | Hest:SB_TH | 0.077 | 0.077 | 1.78 | 0.042 |
| | | Hit:LB_TH | 0.154 | 0.216 | 0.40 | 0.341 |
| | | Hit:LB_UL | 0.130 | 0.216 | -0.41 | 0.656 |
| | | Hit:PI_TH | 0.189 | 0.213 | 1.22 | 0.104 |
| | | Hit:PI_UL | 0.154 | 0.256 | -0.48 | 0.688 |
| | | Hit:PL_TH | 0.105 | 0.210 | -1.09 | 0.867 |
| | | Hit:PL_UL | 0.103 | 0.226 | -1.50 | 0.946 |
| | | Hit:SB_TH | 0.134 | 0.207 | -0.09 | 0.538 |
| | Þingvallavatn | LB_TH:PI_TH | 0.157 | 0.079 | 4.68 | ***0.001** |
| | | LB_TH:PL_TH | 0.107 | 0.083 | 2.77 | ***0.004** |
| | | LB_TH:SB_TH | 0.135 | 0.077 | 4.13 | ***0.001** |
| | | PI_TH:PL_TH | 0.111 | 0.078 | 3.17 | ***0.003** |
| | | PI_TH:SB_TH | 0.089 | 0.069 | 2.70 | ***0.006** |
| | | PL_TH:SB_TH | 0.052 | 0.074 | 0.13 | 0.457 |
| | Úlfljótsvatn | LB_UL:PI_UL | 0.107 | 0.181 | -0.50 | 0.690 |
| | | LB_UL:PL_UL | 0.093 | 0.127 | 0.32 | 0.367 |
| | | PI_UL:PL_UL | 0.138 | 0.190 | 0.25 | 0.403 |
| | Across Þingvallavatn and Úlfljótsvatn | LB_TH:LB_UL | 0.109 | 0.098 | 2.18 | 0.011 |
| | | LB_TH:PI_UL | 0.135 | 0.173 | 0.64 | 0.267 |
| | | LB_TH:PL_UL | 0.095 | 0.116 | 0.74 | 0.219 |
| | | LB_UL:PI_TH | 0.096 | 0.097 | 1.74 | 0.052 |
| | | LB_UL:PL_TH | 0.069 | 0.093 | 0.27 | 0.387 |
| | | LB_UL:SB_TH | 0.069 | 0.090 | 0.48 | 0.317 |
| | | PI_TH:PI_UL | 0.119 | 0.169 | 0.20 | 0.429 |
| | | PI_TH:PL_UL | 0.146 | 0.114 | 2.73 | ***0.005** |
| | | PI_UL:PL_TH | 0.106 | 0.170 | -0.35 | 0.634 |
| | | PI_UL:SB_TH | 0.102 | 0.168 | -0.39 | 0.660 |
| | | PL_TH:PL_UL | 0.088 | 0.118 | 0.42 | 0.341 |
| | | PL_UL:SB_TH | 0.118 | 0.112 | 1.85 | 0.031 |
| Articular-angular[2] | AN populations | AN_Brúará:AN_Gufudalsá | 0.057 | 0.057 | 1.72 | 0.044 |
| | | AN_Brúará:AN_Lambeyrarkvísl | 0.061 | 0.059 | 1.72 | 0.043 |
| | | AN_Brúará:AN_Víðidalsá | 0.074 | 0.056 | 2.62 | ***0.007** |
| | | AN_Brúará:AN_Vífilsdalsá | 0.056 | 0.061 | 1.32 | 0.101 |
| | | AN_Gufudalsá:AN_Lambeyrarkvísl | 0.043 | 0.043 | 1.63 | 0.056 |
| | | AN_Gufudalsá:AN_Víðidalsá | 0.044 | 0.041 | 1.99 | 0.020 |
| | | AN_Gufudalsá:AN_Vífilsdalsá | 0.024 | 0.047 | -0.83 | 0.797 |
| | | AN_Lambeyrarkvísl:AN_Víðidalsá | 0.044 | 0.044 | 1.61 | 0.056 |
| | | AN_Lambeyrarkvísl:AN_Vífilsdalsá | 0.035 | 0.049 | 0.33 | 0.390 |
| | | AN_Víðidalsá:AN_Vífilsdalsá | 0.050 | 0.048 | 1.83 | 0.040 |
| | Brúará to lake morphs | AN_Brúará:Hest | 0.055 | 0.053 | 1.76 | 0.033 |
| | | AN_Brúará:Hit | 0.061 | 0.066 | 1.36 | 0.084 |
| | | AN_Brúará:LB_TH | 0.111 | 0.051 | 4.45 | ***0.001** |
| | | AN_Brúará:LB_UL | 0.054 | 0.054 | 1.72 | 0.046 |

| | | | | | | |
|---|---|---|---|---|---|---|
| | | AN_Brúará:PI_TH | 0.064 | 0.051 | 2.38 | 0.008 |
| | | AN_Brúará:PI_UL | 0.052 | 0.061 | 1.08 | 0.130 |
| | | AN_Brúará:PL_TH | 0.047 | 0.056 | 1.02 | 0.154 |
| | | AN_Brúará:PL_UL | 0.054 | 0.057 | 1.43 | 0.078 |
| | | AN_Brúará:SB_TH | 0.068 | 0.058 | 2.19 | 0.018 |
| | Gufudalsá to lake morphs | AN_Gufudalsá:Hest | 0.051 | 0.033 | 3.36 | ***0.001** |
| | | AN_Gufudalsá:Hit | 0.042 | 0.050 | 1.03 | 0.155 |
| | | AN_Gufudalsá:LB_TH | 0.120 | 0.032 | 6.03 | ***0.001** |
| | | AN_Gufudalsá:LB_UL | 0.068 | 0.036 | 4.28 | ***0.001** |
| | | AN_Gufudalsá:PI_TH | 0.046 | 0.033 | 2.98 | ***0.003** |
| | | AN_Gufudalsá:PI_UL | 0.057 | 0.045 | 2.57 | ***0.004** |
| | | AN_Gufudalsá:PL_TH | 0.045 | 0.034 | 2.72 | ***0.007** |
| | | AN_Gufudalsá:PL_UL | 0.069 | 0.037 | 3.89 | ***0.001** |
| | | AN_Gufudalsá:SB_TH | 0.076 | 0.035 | 4.69 | ***0.001** |
| | Lambeyrarkvísl to lake morphs | AN_Lambeyrarkvísl:Hest | 0.068 | 0.039 | 3.59 | ***0.001** |
| | | AN_Lambeyrarkvísl:Hit | 0.047 | 0.054 | 1.08 | 0.140 |
| | | AN_Lambeyrarkvísl:LB_TH | 0.122 | 0.037 | 5.91 | ***0.001** |
| | | AN_Lambeyrarkvísl:LB_UL | 0.071 | 0.041 | 3.89 | ***0.001** |
| | | AN_Lambeyrarkvísl:PI_TH | 0.068 | 0.037 | 3.98 | ***0.001** |
| | | AN_Lambeyrarkvísl:PI_UL | 0.067 | 0.049 | 2.83 | **0.005** |
| | | AN_Lambeyrarkvísl:PL_TH | 0.061 | 0.041 | 3.16 | ***0.001** |
| | | AN_Lambeyrarkvísl:PL_UL | 0.062 | 0.044 | 2.95 | ***0.002** |
| | | AN_Lambeyrarkvísl:SB_TH | 0.072 | 0.043 | 4.08 | ***0.001** |
| | Víðidalsá to lake morphs | AN_Víðidalsá:Hest | 0.067 | 0.036 | 4.16 | ***0.001** |
| | | AN_Víðidalsá:Hit | 0.051 | 0.051 | 1.56 | 0.056 |
| | | AN_Víðidalsá:LB_TH | 0.113 | 0.034 | 6.37 | ***0.001** |
| | | AN_Víðidalsá:LB_UL | 0.081 | 0.036 | 4.64 | ***0.001** |
| | | AN_Víðidalsá:PI_TH | 0.065 | 0.033 | 4.19 | ***0.001** |
| | | AN_Víðidalsá:PI_UL | 0.074 | 0.045 | 3.61 | ***0.001** |
| | | AN_Víðidalsá:PL_TH | 0.070 | 0.042 | 3.74 | ***0.001** |
| | | AN_Víðidalsá:PL_UL | 0.083 | 0.044 | 3.84 | ***0.001** |
| | | AN_Víðidalsá:SB_TH | 0.071 | 0.044 | 3.46 | ***0.001** |
| | Vífisdalsá to lake morphs | AN_Vífilsdalsá:Hest | 0.054 | 0.043 | 2.57 | ***0.005** |
| | | AN_Vífilsdalsá:Hit | 0.048 | 0.058 | 0.97 | 0.169 |
| | | AN_Vífilsdalsá:LB_TH | 0.125 | 0.041 | 5.40 | ***0.001** |
| | | AN_Vífilsdalsá:LB_UL | 0.068 | 0.045 | 3.07 | ***0.002** |
| | | AN_Vífilsdalsá:PI_TH | 0.053 | 0.040 | 2.54 | ***0.006** |
| | | AN_Vífilsdalsá:PI_UL | 0.059 | 0.052 | 2.22 | 0.021 |
| | | AN_Vífilsdalsá:PL_TH | 0.045 | 0.046 | 1.58 | 0.061 |
| | | AN_Vífilsdalsá:PL_UL | 0.064 | 0.048 | 2.78 | ***0.002** |
| | | AN_Vífilsdalsá:SB_TH | 0.077 | 0.048 | 3.50 | ***0.001** |
| | Hestvatn and Hítarvatn | Hest:Hit | 0.058 | 0.046 | 2.47 | ***0.006** |
| | | Hest:LB_TH | 0.096 | 0.027 | 6.31 | ***0.001** |
| | | Hest:LB_UL | 0.048 | 0.031 | 3.27 | ***0.002** |
| | | Hest:PI_TH | 0.041 | 0.027 | 3.16 | ***0.001** |
| | | Hest:PI_UL | 0.035 | 0.042 | 1.01 | 0.153 |
| | | Hest:PL_TH | 0.054 | 0.030 | 4.06 | ***0.001** |
| | | Hest:PL_UL | 0.067 | 0.033 | 4.24 | ***0.001** |
| | | Hest:SB_TH | 0.064 | 0.031 | 4.88 | ***0.001** |
| | | Hit:LB_TH | 0.101 | 0.046 | 4.86 | ***0.001** |
| | | Hit:LB_UL | 0.062 | 0.048 | 2.67 | ***0.005** |
| | | Hit:PI_TH | 0.048 | 0.046 | 1.85 | 0.029 |
| | | Hit:PI_UL | 0.064 | 0.056 | 2.23 | 0.015 |
| | | Hit:PL_TH | 0.047 | 0.047 | 1.66 | 0.050 |
| | | Hit:PL_UL | 0.061 | 0.048 | 2.62 | ***0.005** |
| | | Hit:SB_TH | 0.062 | 0.047 | 2.66 | ***0.004** |
| | Þingvallavatn | LB_TH:PI_TH | 0.089 | 0.024 | 7.01 | ***0.001** |
| | | LB_TH:PL_TH | 0.111 | 0.032 | 7.02 | ***0.001** |
| | | LB_TH:SB_TH | 0.063 | 0.033 | 4.34 | ***0.001** |
| | | PI_TH:PL_TH | 0.040 | 0.031 | 2.44 | ***0.006** |
| | | PI_TH:SB_TH | 0.054 | 0.034 | 3.48 | ***0.001** |

| | | | | | | |
|---|---|---|---|---|---|---|
| | | PL_TH:SB_TH | 0.061 | 0.021 | 6.12 | ***0.001** |
| | Úlfljótsvatn | LB_UL:PI_UL | 0.031 | 0.041 | 0.49 | 0.312 |
| | | LB_UL:PL_UL | 0.032 | 0.035 | 1.23 | 0.118 |
| | | PI_UL:PL_UL | 0.051 | 0.047 | 1.90 | 0.031 |
| | Across Þingvallavatn and Úlfljótsvatn | LB_TH:LB_UL | 0.076 | 0.029 | 5.19 | ***0.001** |
| | | LB_TH:PI_UL | 0.092 | 0.039 | 4.50 | ***0.001** |
| | | LB_TH:PL_UL | 0.094 | 0.034 | 5.48 | ***0.001** |
| | | LB_UL:PI_TH | 0.045 | 0.029 | 3.22 | ***0.001** |
| | | LB_UL:PL_TH | 0.051 | 0.033 | 3.46 | ***0.001** |
| | | LB_UL:SB_TH | 0.040 | 0.035 | 2.21 | 0.013 |
| | | PI_TH:PI_UL | 0.042 | 0.039 | 1.96 | 0.024 |
| | | PI_TH:PL_UL | 0.057 | 0.034 | 3.74 | ***0.001** |
| | | PI_UL:PL_TH | 0.052 | 0.046 | 2.16 | 0.016 |
| | | PI_UL:SB_TH | 0.060 | 0.048 | 2.54 | 0.008 |
| | | PL_TH:PL_UL | 0.045 | 0.026 | 3.62 | ***0.001** |
| | | PL_UL:SB_TH | 0.046 | 0.025 | 4.19 | ***0.001** |
| Quadrate[2] | AN populations | AN_Brúará:AN_Gufudalsá | 0.052 | 0.063 | 0.98 | 0.174 |
| | | AN_Brúará:AN_Lambeyrarkvísl | 0.061 | 0.065 | 1.35 | 0.092 |
| | | AN_Brúará:AN_Víðidalsá | 0.068 | 0.063 | 1.96 | 0.025 |
| | | AN_Brúará:AN_Vífilsdalsá | 0.069 | 0.069 | 1.64 | 0.054 |
| | | AN_Gufudalsá:AN_Lambeyrarkvísl | 0.037 | 0.045 | 0.85 | 0.208 |
| | | AN_Gufudalsá:AN_Víðidalsá | 0.045 | 0.043 | 1.89 | 0.038 |
| | | AN_Gufudalsá:AN_Vífilsdalsá | 0.050 | 0.048 | 1.81 | 0.042 |
| | | AN_Lambeyrarkvísl:AN_Víðidalsá | 0.046 | 0.046 | 1.65 | 0.051 |
| | | AN_Lambeyrarkvísl:AN_Vífilsdalsá | 0.053 | 0.053 | 1.63 | 0.051 |
| | | AN_Víðidalsá:AN_Vífilsdalsá | 0.055 | 0.050 | 2.08 | 0.019 |
| | Brúará to lake morphs | AN_Brúará:Hest | 0.035 | 0.060 | -0.27 | 0.607 |
| | | AN_Brúará:Hit | 0.071 | 0.069 | 1.73 | 0.042 |
| | | AN_Brúará:LB_TH | 0.065 | 0.059 | 1.97 | 0.021 |
| | | AN_Brúará:LB_UL | 0.069 | 0.061 | 2.13 | 0.012 |
| | | AN_Brúará:PI_TH | 0.072 | 0.061 | 2.25 | 0.011 |
| | | AN_Brúará:PI_UL | 0.078 | 0.067 | 2.18 | 0.012 |
| | | AN_Brúará:PL_TH | 0.053 | 0.062 | 1.11 | 0.125 |
| | | AN_Brúará:PL_UL | 0.047 | 0.062 | 0.59 | 0.282 |
| | | AN_Brúará:SB_TH | 0.053 | 0.062 | 1.05 | 0.141 |
| | Gufudalsá to lake morphs | AN_Gufudalsá:Hest | 0.042 | 0.034 | 2.29 | 0.013 |
| | | AN_Gufudalsá:Hit | 0.042 | 0.053 | 0.90 | 0.178 |
| | | AN_Gufudalsá:LB_TH | 0.081 | 0.034 | 5.15 | ***0.001** |
| | | AN_Gufudalsá:LB_UL | 0.079 | 0.038 | 3.90 | ***0.001** |
| | | AN_Gufudalsá:PI_TH | 0.083 | 0.034 | 4.68 | ***0.001** |
| | | AN_Gufudalsá:PI_UL | 0.084 | 0.048 | 3.75 | ***0.001** |
| | | AN_Gufudalsá:PL_TH | 0.059 | 0.035 | 3.56 | ***0.001** |
| | | AN_Gufudalsá:PL_UL | 0.052 | 0.038 | 2.87 | ***0.001** |
| | | AN_Gufudalsá:SB_TH | 0.058 | 0.036 | 3.50 | ***0.002** |
| | Lambeyrarkvísl to lake morphs | AN_Lambeyrarkvísl:Hest | 0.057 | 0.042 | 2.62 | ***0.007** |
| | | AN_Lambeyrarkvísl:Hit | 0.048 | 0.057 | 1.03 | 0.150 |
| | | AN_Lambeyrarkvísl:LB_TH | 0.084 | 0.040 | 4.05 | ***0.001** |
| | | AN_Lambeyrarkvísl:LB_UL | 0.081 | 0.042 | 3.84 | ***0.001** |
| | | AN_Lambeyrarkvísl:PI_TH | 0.088 | 0.041 | 4.22 | ***0.001** |
| | | AN_Lambeyrarkvísl:PI_UL | 0.088 | 0.053 | 3.25 | ***0.002** |
| | | AN_Lambeyrarkvísl:PL_TH | 0.069 | 0.043 | 3.37 | ***0.001** |
| | | AN_Lambeyrarkvísl:PL_UL | 0.060 | 0.046 | 2.60 | ***0.005** |
| | | AN_Lambeyrarkvísl:SB_TH | 0.066 | 0.045 | 2.99 | ***0.001** |
| | Víðidalsá to lake morphs | AN_Víðidalsá:Hest | 0.054 | 0.037 | 2.91 | ***0.002** |
| | | AN_Víðidalsá:Hit | 0.074 | 0.054 | 2.83 | ***0.003** |
| | | AN_Víðidalsá:LB_TH | 0.086 | 0.034 | 5.14 | ***0.001** |
| | | AN_Víðidalsá:LB_UL | 0.067 | 0.040 | 3.49 | ***0.001** |
| | | AN_Víðidalsá:PI_TH | 0.082 | 0.034 | 5.17 | ***0.001** |
| | | AN_Víðidalsá:PI_UL | 0.068 | 0.048 | 2.90 | ***0.003** |
| | | AN_Víðidalsá:PL_TH | 0.070 | 0.044 | 3.45 | ***0.001** |
| | | AN_Víðidalsá:PL_UL | 0.060 | 0.046 | 2.70 | ***0.005** |

| | | | | | | |
|---|---|---|---|---|---|---|
| | | AN_Víðidalsá:SB_TH | 0.066 | 0.044 | 3.09 | ***0.001** |
| | Vífisdalsá to lake morphs | AN_Vífilsdalsá:Hest | 0.047 | 0.046 | 1.83 | 0.041 |
| | | AN_Vífilsdalsá:Hit | 0.067 | 0.059 | 2.12 | 0.017 |
| | | AN_Vífilsdalsá:LB_TH | 0.072 | 0.044 | 3.56 | ***0.001** |
| | | AN_Vífilsdalsá:LB_UL | 0.067 | 0.046 | 3.06 | ***0.002** |
| | | AN_Vífilsdalsá:PI_TH | 0.065 | 0.043 | 3.22 | ***0.001** |
| | | AN_Vífilsdalsá:PI_UL | 0.068 | 0.055 | 2.45 | ***0.007** |
| | | AN_Vífilsdalsá:PL_TH | 0.059 | 0.048 | 2.39 | 0.009 |
| | | AN_Vífilsdalsá:PL_UL | 0.060 | 0.051 | 2.31 | 0.008 |
| | | AN_Vífilsdalsá:SB_TH | 0.056 | 0.049 | 2.09 | 0.018 |
| | Hestvatn and Hítarvatn | Hest:Hit | 0.064 | 0.049 | 2.84 | ***0.002** |
| | | Hest:LB_TH | 0.059 | 0.028 | 4.74 | ***0.001** |
| | | Hest:LB_UL | 0.057 | 0.032 | 3.76 | ***0.001** |
| | | Hest:PI_TH | 0.059 | 0.028 | 4.48 | ***0.001** |
| | | Hest:PI_UL | 0.063 | 0.045 | 2.99 | ***0.003** |
| | | Hest:PL_TH | 0.045 | 0.030 | 2.88 | ***0.005** |
| | | Hest:PL_UL | 0.040 | 0.034 | 2.25 | 0.013 |
| | | Hest:SB_TH | 0.046 | 0.031 | 3.05 | ***0.002** |
| | | Hit:LB_TH | 0.098 | 0.049 | 4.41 | ***0.001** |
| | | Hit:LB_UL | 0.105 | 0.051 | 4.05 | ***0.001** |
| | | Hit:PI_TH | 0.104 | 0.048 | 4.67 | ***0.001** |
| | | Hit:PI_UL | 0.111 | 0.059 | 4.26 | ***0.001** |
| | | Hit:PL_TH | 0.081 | 0.049 | 3.43 | ***0.001** |
| | | Hit:PL_UL | 0.077 | 0.052 | 2.99 | ***0.002** |
| | | Hit:SB_TH | 0.081 | 0.052 | 3.24 | ***0.001** |
| | Þingvallavatn | LB_TH:PI_TH | 0.026 | 0.026 | 1.71 | 0.049 |
| | | LB_TH:PL_TH | 0.033 | 0.034 | 1.52 | 0.067 |
| | | LB_TH:SB_TH | 0.037 | 0.036 | 1.72 | 0.040 |
| | | PI_TH:PL_TH | 0.037 | 0.033 | 2.10 | 0.018 |
| | | PI_TH:SB_TH | 0.038 | 0.034 | 1.94 | 0.025 |
| | | PL_TH:SB_TH | 0.023 | 0.021 | 1.96 | 0.024 |
| | Úlfljótsvatn | LB_UL:PI_UL | 0.020 | 0.046 | -1.33 | 0.901 |
| | | LB_UL:PL_UL | 0.041 | 0.037 | 2.02 | 0.025 |
| | | PI_UL:PL_UL | 0.052 | 0.049 | 1.89 | 0.029 |
| | Across Þingvallavatn and Úlfljótsvatn | LB_TH:LB_UL | 0.044 | 0.031 | 3.05 | ***0.002** |
| | | LB_TH:PI_UL | 0.057 | 0.041 | 2.84 | ***0.005** |
| | | LB_TH:PL_UL | 0.037 | 0.036 | 1.65 | 0.047 |
| | | LB_UL:PI_TH | 0.035 | 0.030 | 2.18 | 0.018 |
| | | LB_UL:PL_TH | 0.052 | 0.033 | 3.11 | ***0.003** |
| | | LB_UL:SB_TH | 0.046 | 0.035 | 2.62 | ***0.005** |
| | | PI_TH:PI_UL | 0.041 | 0.041 | 1.63 | 0.060 |
| | | PI_TH:PL_UL | 0.043 | 0.035 | 2.55 | ***0.004** |
| | | PI_UL:PL_TH | 0.059 | 0.047 | 2.49 | ***0.005** |
| | | PI_UL:SB_TH | 0.054 | 0.049 | 2.08 | 0.023 |
| | | PL_TH:PL_UL | 0.024 | 0.028 | 1.15 | 0.126 |
| | | PL_UL:SB_TH | 0.029 | 0.027 | 1.96 | 0.027 |
| Premaxilla[1] | AN populations | AN_Brúará:AN_Gufudalsá | 0.243 | 0.428 | -0.84 | 0.792 |
| | | AN_Brúará:AN_Lambeyrarkvísl | 0.218 | 0.500 | -1.88 | 0.968 |
| | | AN_Brúará:AN_Víðidalsá | 0.244 | 0.421 | -0.70 | 0.738 |
| | | AN_Brúará:AN_Vífilsdalsá | 0.322 | 0.487 | -0.15 | 0.553 |
| | | AN_Gufudalsá:AN_Lambeyrarkvísl | 0.227 | 0.342 | -0.23 | 0.586 |
| | | AN_Gufudalsá:AN_Víðidalsá | 0.172 | 0.213 | 0.68 | 0.246 |
| | | AN_Gufudalsá:AN_Vífilsdalsá | 0.178 | 0.308 | -0.93 | 0.827 |
| | | AN_Lambeyrarkvísl:AN_Víðidalsá | 0.269 | 0.325 | 0.72 | 0.245 |
| | | AN_Lambeyrarkvísl:AN_Vífilsdalsá | 0.311 | 0.405 | 0.49 | 0.310 |
| | | AN_Víðidalsá:AN_Vífilsdalsá | 0.207 | 0.296 | 0.02 | 0.492 |
| | Brúará to lake morphs | AN_Brúará:Hest | 0.231 | 0.408 | -0.83 | 0.791 |
| | | AN_Brúará:Hit | 0.510 | 0.529 | 1.40 | 0.077 |
| | | AN_Brúará:LB_TH | 0.263 | 0.412 | -0.30 | 0.619 |
| | | AN_Brúará:LB_UL | 0.263 | 0.412 | -0.40 | 0.651 |
| | | AN_Brúará:PI_TH | 0.254 | 0.408 | -0.40 | 0.658 |

| | | | | | | |
|---|---|---|---|---|---|---|
| | | AN_Brúará:PI_UL | 0.301 | 0.466 | -0.26 | 0.596 |
| | | AN_Brúará:PL_TH | 0.330 | 0.410 | 0.69 | 0.251 |
| | | AN_Brúará:PL_UL | 0.361 | 0.438 | 0.84 | 0.207 |
| | | AN_Brúará:SB_TH | 0.376 | 0.409 | 1.27 | 0.113 |
| | Gufudalsá to lake morphs | AN_Gufudalsá:Hest | 0.122 | 0.186 | -0.38 | 0.644 |
| | | AN_Gufudalsá:Hit | 0.377 | 0.394 | 1.39 | 0.081 |
| | | AN_Gufudalsá:LB_TH | 0.081 | 0.188 | -2.17 | 0.986 |
| | | AN_Gufudalsá:LB_UL | 0.065 | 0.203 | -3.43 | 1.000 |
| | | AN_Gufudalsá:PI_TH | 0.131 | 0.181 | 0.13 | 0.447 |
| | | AN_Gufudalsá:PI_UL | 0.155 | 0.277 | -1.07 | 0.853 |
| | | AN_Gufudalsá:PL_TH | 0.172 | 0.186 | 1.27 | 0.109 |
| | | AN_Gufudalsá:PL_UL | 0.188 | 0.224 | 0.77 | 0.210 |
| | | AN_Gufudalsá:SB_TH | 0.209 | 0.179 | 2.45 | ***0.006** |
| | Lambeyrarkvísl to lake morphs | AN_Lambeyrarkvísl:Hest | 0.270 | 0.314 | 0.96 | 0.170 |
| | | AN_Lambeyrarkvísl:Hit | 0.522 | 0.471 | 2.20 | 0.017 |
| | | AN_Lambeyrarkvísl:LB_TH | 0.221 | 0.312 | 0.04 | 0.480 |
| | | AN_Lambeyrarkvísl:LB_UL | 0.237 | 0.323 | 0.19 | 0.412 |
| | | AN_Lambeyrarkvísl:PI_TH | 0.273 | 0.312 | 1.06 | 0.148 |
| | | AN_Lambeyrarkvísl:PI_UL | 0.284 | 0.368 | 0.34 | 0.367 |
| | | AN_Lambeyrarkvísl:PL_TH | 0.321 | 0.311 | 1.77 | 0.032 |
| | | AN_Lambeyrarkvísl:PL_UL | 0.324 | 0.340 | 1.40 | 0.082 |
| | | AN_Lambeyrarkvísl:SB_TH | 0.376 | 0.308 | 2.65 | ***0.004** |
| | Víðidalsá to lake morphs | AN_Víðidalsá:Hest | 0.111 | 0.157 | -0.04 | 0.523 |
| | | AN_Víðidalsá:Hit | 0.384 | 0.400 | 1.54 | 0.069 |
| | | AN_Víðidalsá:LB_TH | 0.175 | 0.159 | 2.09 | 0.017 |
| | | AN_Víðidalsá:LB_UL | 0.178 | 0.180 | 1.63 | 0.058 |
| | | AN_Víðidalsá:PI_TH | 0.134 | 0.147 | 1.17 | 0.127 |
| | | AN_Víðidalsá:PI_UL | 0.225 | 0.257 | 0.92 | 0.202 |
| | | AN_Víðidalsá:PL_TH | 0.208 | 0.157 | 3.15 | ***0.001** |
| | | AN_Víðidalsá:PL_UL | 0.224 | 0.202 | 2.16 | 0.015 |
| | | AN_Víðidalsá:SB_TH | 0.226 | 0.149 | 3.58 | ***0.001** |
| | Vífisdalsá to lake morphs | AN_Vífilsdalsá:Hest | 0.148 | 0.281 | -1.20 | 0.883 |
| | | AN_Vífilsdalsá:Hit | 0.401 | 0.449 | 1.11 | 0.140 |
| | | AN_Vífilsdalsá:LB_TH | 0.173 | 0.278 | -0.58 | 0.711 |
| | | AN_Vífilsdalsá:LB_UL | 0.179 | 0.297 | -0.57 | 0.723 |
| | | AN_Vífilsdalsá:PI_TH | 0.188 | 0.279 | -0.07 | 0.532 |
| | | AN_Vífilsdalsá:PI_UL | 0.255 | 0.345 | 0.23 | 0.399 |
| | | AN_Vífilsdalsá:PL_TH | 0.173 | 0.280 | -0.47 | 0.672 |
| | | AN_Vífilsdalsá:PL_UL | 0.210 | 0.305 | -0.08 | 0.530 |
| | | AN_Vífilsdalsá:SB_TH | 0.218 | 0.277 | 0.57 | 0.305 |
| | Hestvatn and Hítarvatn | Hest:Hit | 0.364 | 0.379 | 1.42 | 0.080 |
| | | Hest:LB_TH | 0.136 | 0.127 | 2.00 | 0.023 |
| | | Hest:LB_UL | 0.121 | 0.150 | 0.53 | 0.294 |
| | | Hest:PI_TH | 0.088 | 0.114 | 0.44 | 0.337 |
| | | Hest:PI_UL | 0.192 | 0.243 | 0.49 | 0.317 |
| | | Hest:PL_TH | 0.142 | 0.122 | 2.44 | 0.009 |
| | | Hest:PL_UL | 0.197 | 0.177 | 2.24 | 0.013 |
| | | Hest:SB_TH | 0.180 | 0.111 | 4.11 | ***0.001** |
| | | Hit:LB_TH | 0.383 | 0.374 | 1.70 | 0.040 |
| | | Hit:LB_UL | 0.373 | 0.388 | 1.43 | 0.078 |
| | | Hit:PI_TH | 0.400 | 0.373 | 1.92 | 0.020 |
| | | Hit:PI_UL | 0.395 | 0.436 | 1.26 | 0.115 |
| | | Hit:PL_TH | 0.358 | 0.380 | 1.39 | 0.085 |
| | | Hit:PL_UL | 0.330 | 0.396 | 0.76 | 0.243 |
| | | Hit:SB_TH | 0.319 | 0.373 | 0.92 | 0.184 |
| | Þingvallavatn | LB_TH:PI_TH | 0.149 | 0.115 | 2.99 | ***0.003** |
| | | LB_TH:PL_TH | 0.160 | 0.127 | 2.72 | ***0.003** |
| | | LB_TH:SB_TH | 0.203 | 0.114 | 4.88 | ***0.001** |
| | | PI_TH:PL_TH | 0.142 | 0.111 | 3.02 | ***0.001** |
| | | PI_TH:SB_TH | 0.175 | 0.100 | 4.52 | ***0.001** |
| | | PL_TH:SB_TH | 0.086 | 0.107 | 0.52 | 0.302 |

| | | | | | | |
|---|---|---|---|---|---|---|
| | Úlfljótsvatn | LB_UL:PI_UL | 0.164 | 0.253 | -0.50 | 0.706 |
| | | LB_UL:PL_UL | 0.163 | 0.201 | 0.64 | 0.266 |
| | | PI_UL:PL_UL | 0.221 | 0.272 | 0.56 | 0.289 |
| | Across Þingvallavatn and Úlfljótsvatn | LB_TH:LB_UL | 0.088 | 0.153 | -0.99 | 0.847 |
| | | LB_TH:PI_UL | 0.156 | 0.240 | -0.46 | 0.674 |
| | | LB_TH:PL_UL | 0.165 | 0.181 | 1.15 | 0.130 |
| | | LB_UL:PI_TH | 0.115 | 0.143 | 0.56 | 0.297 |
| | | LB_UL:PL_TH | 0.141 | 0.149 | 1.38 | 0.094 |
| | | LB_UL:SB_TH | 0.178 | 0.141 | 2.89 | ***0.002** |
| | | PI_TH:PI_UL | 0.169 | 0.236 | 0.02 | 0.496 |
| | | PI_TH:PL_UL | 0.210 | 0.172 | 2.76 | ***0.002** |
| | | PI_UL:PL_TH | 0.176 | 0.243 | 0.11 | 0.452 |
| | | PI_UL:SB_TH | 0.208 | 0.236 | 1.01 | 0.145 |
| | | PL_TH:PL_UL | 0.141 | 0.177 | 0.53 | 0.308 |
| | | PL_UL:SB_TH | 0.145 | 0.171 | 0.82 | 0.213 |
| Maxilla[2] | AN populations | AN_Brúará:AN_Gufudalsá | 0.031 | 0.047 | 0.26 | 0.394 |
| | | AN_Brúará:AN_Lambeyrarkvísl | 0.030 | 0.049 | -0.10 | 0.529 |
| | | AN_Brúará:AN_Víðidalsá | 0.039 | 0.048 | 1.00 | 0.160 |
| | | AN_Brúará:AN_Vífilsdalsá | 0.039 | 0.051 | 0.71 | 0.241 |
| | | AN_Gufudalsá:AN_Lambeyrarkvísl | 0.042 | 0.036 | 2.23 | 0.010 |
| | | AN_Gufudalsá:AN_Víðidalsá | 0.034 | 0.034 | 1.67 | 0.056 |
| | | AN_Gufudalsá:AN_Vífilsdalsá | 0.029 | 0.039 | 0.63 | 0.271 |
| | | AN_Lambeyrarkvísl:AN_Víðidalsá | 0.052 | 0.036 | 2.86 | ***0.001** |
| | | AN_Lambeyrarkvísl:AN_Vífilsdalsá | 0.044 | 0.044 | 1.73 | 0.051 |
| | | AN_Víðidalsá:AN_Vífilsdalsá | 0.027 | 0.039 | 0.35 | 0.370 |
| | Brúará to lake morphs | AN_Brúará:Hest | 0.033 | 0.044 | 0.68 | 0.247 |
| | | AN_Brúará:Hit | 0.069 | 0.053 | 2.56 | ***0.006** |
| | | AN_Brúará:LB_TH | 0.039 | 0.042 | 1.32 | 0.097 |
| | | AN_Brúará:LB_UL | 0.036 | 0.046 | 0.86 | 0.201 |
| | | AN_Brúará:PI_TH | 0.029 | 0.044 | 0.31 | 0.388 |
| | | AN_Brúará:PI_UL | 0.032 | 0.050 | 0.01 | 0.512 |
| | | AN_Brúará:PL_TH | 0.052 | 0.045 | 2.11 | 0.018 |
| | | AN_Brúará:PL_UL | 0.043 | 0.047 | 1.35 | 0.093 |
| | | AN_Brúará:SB_TH | 0.054 | 0.047 | 2.11 | 0.011 |
| | Gufudalsá to lake morphs | AN_Gufudalsá:Hest | 0.038 | 0.028 | 2.84 | ***0.002** |
| | | AN_Gufudalsá:Hit | 0.062 | 0.041 | 3.01 | ***0.002** |
| | | AN_Gufudalsá:LB_TH | 0.058 | 0.027 | 4.43 | ***0.001** |
| | | AN_Gufudalsá:LB_UL | 0.045 | 0.029 | 3.33 | ***0.001** |
| | | AN_Gufudalsá:PI_TH | 0.046 | 0.027 | 3.85 | ***0.001** |
| | | AN_Gufudalsá:PI_UL | 0.045 | 0.038 | 2.26 | **0.007** |
| | | AN_Gufudalsá:PL_TH | 0.055 | 0.028 | 4.01 | ***0.001** |
| | | AN_Gufudalsá:PL_UL | 0.048 | 0.030 | 3.28 | ***0.001** |
| | | AN_Gufudalsá:SB_TH | 0.067 | 0.029 | 4.77 | ***0.001** |
| | Lambeyrarkvísl to lake morphs | AN_Lambeyrarkvísl:Hest | 0.051 | 0.033 | 3.22 | ***0.001** |
| | | AN_Lambeyrarkvísl:Hit | 0.084 | 0.045 | 3.97 | ***0.001** |
| | | AN_Lambeyrarkvísl:LB_TH | 0.051 | 0.032 | 3.18 | ***0.001** |
| | | AN_Lambeyrarkvísl:LB_UL | 0.029 | 0.034 | 1.07 | 0.144 |
| | | AN_Lambeyrarkvísl:PI_TH | 0.037 | 0.032 | 2.18 | 0.014 |
| | | AN_Lambeyrarkvísl:PI_UL | 0.036 | 0.041 | 1.21 | 0.126 |
| | | AN_Lambeyrarkvísl:PL_TH | 0.073 | 0.035 | 4.43 | ***0.001** |
| | | AN_Lambeyrarkvísl:PL_UL | 0.041 | 0.038 | 1.96 | 0.026 |
| | | AN_Lambeyrarkvísl:SB_TH | 0.071 | 0.036 | 3.96 | ***0.001** |
| | Víðidalsá to lake morphs | AN_Víðidalsá:Hest | 0.046 | 0.029 | 3.26 | ***0.001** |
| | | AN_Víðidalsá:Hit | 0.040 | 0.043 | 1.38 | 0.077 |
| | | AN_Víðidalsá:LB_TH | 0.064 | 0.027 | 4.81 | ***0.001** |
| | | AN_Víðidalsá:LB_UL | 0.054 | 0.031 | 3.78 | ***0.001** |
| | | AN_Víðidalsá:PI_TH | 0.053 | 0.027 | 4.23 | ***0.001** |
| | | AN_Víðidalsá:PI_UL | 0.051 | 0.038 | 2.83 | ***0.002** |
| | | AN_Víðidalsá:PL_TH | 0.053 | 0.035 | 3.23 | ***0.001** |
| | | AN_Víðidalsá:PL_UL | 0.049 | 0.037 | 2.84 | ***0.003** |
| | | AN_Víðidalsá:SB_TH | 0.073 | 0.037 | 4.00 | ***0.001** |

| | | | | | | |
|---|---|---|---|---|---|---|
| | Vífisdalsá to lake morphs | AN_Vífilsdalsá:Hest | 0.043 | 0.036 | 2.24 | 0.011 |
| | | AN_Vífilsdalsá:Hit | 0.050 | 0.048 | 1.78 | 0.033 |
| | | AN_Vífilsdalsá:LB_TH | 0.059 | 0.035 | 3.30 | ***0.001** |
| | | AN_Vífilsdalsá:LB_UL | 0.045 | 0.037 | 2.41 | ***0.007** |
| | | AN_Vífilsdalsá:PI_TH | 0.047 | 0.034 | 2.83 | ***0.001** |
| | | AN_Vífilsdalsá:PI_UL | 0.043 | 0.044 | 1.65 | 0.052 |
| | | AN_Vífilsdalsá:PL_TH | 0.056 | 0.038 | 3.02 | ***0.003** |
| | | AN_Vífilsdalsá:PL_UL | 0.042 | 0.040 | 1.84 | 0.034 |
| | | AN_Vífilsdalsá:SB_TH | 0.069 | 0.040 | 3.72 | ***0.001** |
| | Hestvatn and Hítarvatn | Hest:Hit | 0.065 | 0.039 | 3.60 | ***0.001** |
| | | Hest:LB_TH | 0.048 | 0.023 | 4.29 | ***0.001** |
| | | Hest:LB_UL | 0.055 | 0.026 | 4.29 | ***0.001** |
| | | Hest:PI_TH | 0.042 | 0.022 | 3.72 | ***0.001** |
| | | Hest:PI_UL | 0.035 | 0.034 | 1.66 | 0.047 |
| | | Hest:PL_TH | 0.035 | 0.024 | 3.03 | ***0.001** |
| | | Hest:PL_UL | 0.059 | 0.028 | 4.40 | ***0.001** |
| | | Hest:SB_TH | 0.045 | 0.026 | 3.71 | ***0.001** |
| | | Hit:LB_TH | 0.083 | 0.038 | 4.46 | ***0.001** |
| | | Hit:LB_UL | 0.085 | 0.040 | 4.57 | ***0.001** |
| | | Hit:PI_TH | 0.078 | 0.039 | 4.34 | ***0.001** |
| | | Hit:PI_UL | 0.078 | 0.047 | 3.48 | ***0.001** |
| | | Hit:PL_TH | 0.057 | 0.038 | 3.13 | ***0.001** |
| | | Hit:PL_UL | 0.077 | 0.040 | 3.93 | ***0.001** |
| | | Hit:SB_TH | 0.083 | 0.039 | 4.62 | ***0.001** |
| | Þingvallavatn | LB_TH:PI_TH | 0.024 | 0.020 | 2.17 | 0.017 |
| | | LB_TH:PL_TH | 0.054 | 0.026 | 4.55 | ***0.001** |
| | | LB_TH:SB_TH | 0.035 | 0.028 | 2.47 | ***0.003** |
| | | PI_TH:PL_TH | 0.050 | 0.027 | 3.99 | ***0.001** |
| | | PI_TH:SB_TH | 0.050 | 0.028 | 3.56 | ***0.001** |
| | | PL_TH:SB_TH | 0.052 | 0.017 | 5.74 | ***0.001** |
| | Úlfljótsvatn | LB_UL:PI_UL | 0.030 | 0.034 | 1.09 | 0.146 |
| | | LB_UL:PL_UL | 0.021 | 0.031 | 0.32 | 0.377 |
| | | PI_UL:PL_UL | 0.038 | 0.040 | 1.49 | 0.079 |
| | Across Þingvallavatn and Úlfljótsvatn | LB_TH:LB_UL | 0.051 | 0.024 | 4.23 | ***0.001** |
| | | LB_TH:PI_UL | 0.044 | 0.032 | 2.82 | ***0.004** |
| | | LB_TH:PL_UL | 0.055 | 0.029 | 4.02 | ***0.001** |
| | | LB_UL:PI_TH | 0.045 | 0.024 | 3.88 | ***0.001** |
| | | LB_UL:PL_TH | 0.078 | 0.028 | 5.59 | ***0.001** |
| | | LB_UL:SB_TH | 0.065 | 0.029 | 4.46 | ***0.001** |
| | | PI_TH:PI_UL | 0.040 | 0.033 | 2.42 | ***0.003** |
| | | PI_TH:PL_UL | 0.051 | 0.030 | 3.47 | ***0.001** |
| | | PI_UL:PL_TH | 0.061 | 0.037 | 3.43 | ***0.001** |
| | | PI_UL:SB_TH | 0.052 | 0.040 | 2.72 | ***0.002** |
| | | PL_TH:PL_UL | 0.079 | 0.022 | 7.05 | ***0.001** |
| | | PL_UL:SB_TH | 0.068 | 0.021 | 5.68 | ***0.001** |
| Supramaxilla[2] | AN populations | AN_Brúará:AN_Gufudalsá | 0.057 | 0.090 | 0.23 | 0.403 |
| | | AN_Brúará:AN_Lambeyrarkvísl | 0.048 | 0.100 | -0.46 | 0.676 |
| | | AN_Brúará:AN_Víðidalsá | 0.088 | 0.088 | 1.51 | 0.055 |
| | | AN_Brúará:AN_Vífilsdalsá | 0.036 | 0.103 | -1.17 | 0.870 |
| | | AN_Gufudalsá:AN_Lambeyrarkvísl | 0.042 | 0.071 | -0.01 | 0.508 |
| | | AN_Gufudalsá:AN_Víðidalsá | 0.083 | 0.066 | 2.37 | 0.011 |
| | | AN_Gufudalsá:AN_Vífilsdalsá | 0.046 | 0.076 | 0.20 | 0.425 |
| | | AN_Lambeyrarkvísl:AN_Víðidalsá | 0.065 | 0.074 | 1.21 | 0.118 |
| | | AN_Lambeyrarkvísl:AN_Vífilsdalsá | 0.028 | 0.082 | -1.34 | 0.902 |
| | | AN_Víðidalsá:AN_Vífilsdalsá | 0.072 | 0.077 | 1.48 | 0.075 |
| | Brúará to lake morphs | AN_Brúará:Hest | 0.034 | 0.087 | -1.00 | 0.838 |
| | | AN_Brúará:Hit | 0.114 | 0.106 | 1.90 | 0.029 |
| | | AN_Brúará:LB_TH | 0.061 | 0.083 | 0.68 | 0.246 |
| | | AN_Brúará:LB_UL | 0.045 | 0.086 | -0.32 | 0.641 |
| | | AN_Brúará:PI_TH | 0.057 | 0.085 | 0.43 | 0.337 |
| | | AN_Brúará:PI_UL | 0.056 | 0.098 | -0.04 | 0.520 |

| | | | | | | |
|---|---|---|---|---|---|---|
| | | AN_Brúará:PL_TH | 0.084 | 0.089 | 1.37 | 0.074 |
| | | AN_Brúará:PL_UL | 0.059 | 0.093 | 0.24 | 0.392 |
| | | AN_Brúará:SB_TH | 0.140 | 0.092 | 3.12 | ***0.002** |
| | Gufudalsá to lake morphs | AN_Gufudalsá:Hest | 0.060 | 0.054 | 1.96 | 0.021 |
| | | AN_Gufudalsá:Hit | 0.086 | 0.080 | 1.89 | 0.026 |
| | | AN_Gufudalsá:LB_TH | 0.038 | 0.053 | 0.61 | 0.273 |
| | | AN_Gufudalsá:LB_UL | 0.071 | 0.056 | 2.44 | ***0.006** |
| | | AN_Gufudalsá:PI_TH | 0.034 | 0.054 | 0.32 | 0.383 |
| | | AN_Gufudalsá:PI_UL | 0.074 | 0.072 | 1.76 | 0.044 |
| | | AN_Gufudalsá:PL_TH | 0.050 | 0.056 | 1.36 | 0.089 |
| | | AN_Gufudalsá:PL_UL | 0.042 | 0.058 | 0.60 | 0.282 |
| | | AN_Gufudalsá:SB_TH | 0.100 | 0.054 | 4.15 | ***0.001** |
| | Lambeyrarkvísl to lake morphs | AN_Lambeyrarkvísl:Hest | 0.031 | 0.065 | -0.60 | 0.713 |
| | | AN_Lambeyrarkvísl:Hit | 0.090 | 0.087 | 1.76 | 0.039 |
| | | AN_Lambeyrarkvísl:LB_TH | 0.048 | 0.064 | 0.75 | 0.226 |
| | | AN_Lambeyrarkvísl:LB_UL | 0.045 | 0.068 | 0.28 | 0.396 |
| | | AN_Lambeyrarkvísl:PI_TH | 0.035 | 0.062 | -0.27 | 0.599 |
| | | AN_Lambeyrarkvísl:PI_UL | 0.065 | 0.082 | 0.91 | 0.192 |
| | | AN_Lambeyrarkvísl:PL_TH | 0.071 | 0.070 | 1.64 | 0.042 |
| | | AN_Lambeyrarkvísl:PL_UL | 0.065 | 0.074 | 1.20 | 0.119 |
| | | AN_Lambeyrarkvísl:SB_TH | 0.129 | 0.073 | 3.73 | ***0.001** |
| | Víðidalsá to lake morphs | AN_Víðidalsá:Hest | 0.080 | 0.060 | 2.56 | 0.008 |
| | | AN_Víðidalsá:Hit | 0.060 | 0.086 | 0.53 | 0.304 |
| | | AN_Víðidalsá:LB_TH | 0.100 | 0.053 | 3.98 | ***0.001** |
| | | AN_Víðidalsá:LB_UL | 0.100 | 0.061 | 3.25 | ***0.001** |
| | | AN_Víðidalsá:PI_TH | 0.081 | 0.053 | 3.11 | ***0.001** |
| | | AN_Víðidalsá:PI_UL | 0.125 | 0.078 | 3.23 | ***0.001** |
| | | AN_Víðidalsá:PL_TH | 0.079 | 0.070 | 2.03 | 0.024 |
| | | AN_Víðidalsá:PL_UL | 0.109 | 0.073 | 3.14 | ***0.002** |
| | | AN_Víðidalsá:SB_TH | 0.156 | 0.072 | 4.34 | ***0.001** |
| | Vífisdalsá to lake morphs | AN_Vífilsdalsá:Hest | 0.032 | 0.072 | -0.63 | 0.730 |
| | | AN_Vífilsdalsá:Hit | 0.103 | 0.095 | 2.00 | 0.030 |
| | | AN_Vífilsdalsá:LB_TH | 0.061 | 0.069 | 1.40 | 0.096 |
| | | AN_Vífilsdalsá:LB_UL | 0.047 | 0.070 | 0.35 | 0.367 |
| | | AN_Vífilsdalsá:PI_TH | 0.052 | 0.067 | 0.88 | 0.203 |
| | | AN_Vífilsdalsá:PI_UL | 0.066 | 0.082 | 0.97 | 0.174 |
| | | AN_Vífilsdalsá:PL_TH | 0.083 | 0.072 | 2.00 | 0.023 |
| | | AN_Vífilsdalsá:PL_UL | 0.070 | 0.075 | 1.43 | 0.077 |
| | | AN_Vífilsdalsá:SB_TH | 0.142 | 0.075 | 4.01 | ***0.001** |
| | Hestvatn and Hítarvatn | Hest:Hit | 0.114 | 0.075 | 3.21 | ***0.001** |
| | | Hest:LB_TH | 0.056 | 0.044 | 2.40 | 0.011 |
| | | Hest:LB_UL | 0.027 | 0.050 | -0.24 | 0.601 |
| | | Hest:PI_TH | 0.048 | 0.043 | 1.93 | 0.027 |
| | | Hest:PI_UL | 0.050 | 0.068 | 0.73 | 0.234 |
| | | Hest:PL_TH | 0.088 | 0.047 | 3.83 | ***0.001** |
| | | Hest:PL_UL | 0.069 | 0.052 | 2.70 | ***0.003** |
| | | Hest:SB_TH | 0.142 | 0.048 | 6.03 | ***0.001** |
| | | Hit:LB_TH | 0.104 | 0.073 | 3.00 | ***0.002** |
| | | Hit:LB_UL | 0.129 | 0.079 | 3.50 | ***0.001** |
| | | Hit:PI_TH | 0.089 | 0.075 | 2.22 | 0.013 |
| | | Hit:PI_UL | 0.145 | 0.090 | 3.37 | ***0.001** |
| | | Hit:PL_TH | 0.062 | 0.075 | 1.00 | 0.170 |
| | | Hit:PL_UL | 0.107 | 0.080 | 2.68 | ***0.003** |
| | | Hit:SB_TH | 0.130 | 0.074 | 3.77 | ***0.001** |
| | Þingvallavatn | LB_TH:PI_TH | 0.022 | 0.039 | -0.12 | 0.545 |
| | | LB_TH:PL_TH | 0.062 | 0.052 | 2.16 | 0.013 |
| | | LB_TH:SB_TH | 0.093 | 0.054 | 3.48 | ***0.001** |
| | | PI_TH:PL_TH | 0.053 | 0.054 | 1.60 | 0.059 |
| | | PI_TH:SB_TH | 0.099 | 0.056 | 3.76 | ***0.001** |
| | | PL_TH:SB_TH | 0.082 | 0.034 | 4.80 | ***0.001** |
| | Úlfljótsvatn | LB_UL:PI_UL | 0.033 | 0.073 | -0.45 | 0.662 |

| | | | | | | |
|---|---|---|---|---|---|---|
| | | LB_UL:PL_UL | 0.073 | 0.057 | 2.55 | ***0.006** |
| | | PI_UL:PL_UL | 0.062 | 0.076 | 1.03 | 0.160 |
| | Across Þingvallavatn and Úlfljótsvatn | LB_TH:LB_UL | 0.058 | 0.047 | 2.33 | 0.011 |
| | | LB_TH:PI_UL | 0.053 | 0.065 | 1.03 | 0.150 |
| | | LB_TH:PL_UL | 0.032 | 0.055 | -0.08 | 0.529 |
| | | LB_UL:PI_TH | 0.058 | 0.047 | 2.26 | 0.009 |
| | | LB_UL:PL_TH | 0.103 | 0.052 | 4.12 | ***0.001** |
| | | LB_UL:SB_TH | 0.149 | 0.054 | 5.95 | ***0.001** |
| | | PI_TH:PI_UL | 0.063 | 0.064 | 1.63 | 0.053 |
| | | PI_TH:PL_UL | 0.043 | 0.056 | 0.79 | 0.227 |
| | | PI_UL:PL_TH | 0.107 | 0.074 | 2.86 | ***0.002** |
| | | PI_UL:SB_TH | 0.136 | 0.078 | 3.69 | ***0.001** |
| | | PL_TH:PL_UL | 0.056 | 0.044 | 2.50 | ***0.005** |
| | | PL_UL:SB_TH | 0.088 | 0.042 | 4.53 | ***0.001** |

1 For Dentary and Premaxilla the full model (shape = CSbone x population) had the best fit.

2 For the Articular-angular, Quadrate, Maxilla and Supramaxilla the reduced model (shape = CSbone + population) had the best fit

D = pairwise differences between vector lengths; UCL = upper confidence level; Z = Z-statistic; P = P-value, ≤ 0.008 in bold with asterisk.

***Supplementary material 20***: *Results from pairwise tests on population differences in mean bone shape, from the best fit ANCOVA models for each bone. AN-charr pooled as one population.*

| Bone | System | Pairs | D | UCL | Z | P-value |
|---|---|---|---|---|---|---|
| Dentary[2] | AN to lake morphs | AN:Hest | 0.070 | 0.075 | 1.36 | 0.076 |
| | | AN:Hit | 0.143 | 0.213 | 0.10 | 0.461 |
| | | AN:LB_TH | 0.054 | 0.074 | 0.20 | 0.419 |
| | | AN:LB_UL | 0.090 | 0.088 | 1.69 | 0.039 |
| | | AN:PI_TH | 0.130 | 0.068 | 4.34 | ***0.001** |
| | | AN:PI_UL | 0.117 | 0.167 | 0.08 | 0.463 |
| | | AN:PL_TH | 0.085 | 0.071 | 2.32 | 0.010 |
| | | AN:PL_UL | 0.085 | 0.114 | 0.41 | 0.329 |
| | | AN:SB_TH | 0.108 | 0.065 | 3.79 | ***0.001** |
| | Hestvatn and Hítarvatn | Hest:Hit | 0.118 | 0.219 | -0.75 | 0.781 |
| | | Hest:LB_TH | 0.099 | 0.090 | 2.11 | 0.015 |
| | | Hest:LB_UL | 0.062 | 0.099 | -0.36 | 0.649 |
| | | Hest:PI_TH | 0.103 | 0.081 | 2.78 | ***0.004** |
| | | Hest:PI_UL | 0.101 | 0.172 | -0.65 | 0.737 |
| | | Hest:PL_TH | 0.070 | 0.085 | 0.74 | 0.230 |
| | | Hest:PL_UL | 0.086 | 0.120 | 0.19 | 0.421 |
| | | Hest:SB_TH | 0.077 | 0.077 | 1.70 | 0.050 |
| | | Hit:LB_TH | 0.154 | 0.220 | 0.32 | 0.366 |
| | | Hit:LB_UL | 0.130 | 0.223 | -0.48 | 0.685 |
| | | Hit:PI_TH | 0.189 | 0.216 | 1.15 | 0.121 |
| | | Hit:PI_UL | 0.154 | 0.264 | -0.54 | 0.701 |
| | | Hit:PL_TH | 0.105 | 0.216 | -1.17 | 0.878 |
| | | Hit:PL_UL | 0.103 | 0.231 | -1.54 | 0.945 |
| | | Hit:SB_TH | 0.134 | 0.213 | -0.17 | 0.573 |
| | Þingvallavatn | LB_TH:PI_TH | 0.157 | 0.081 | 4.55 | ***0.001** |
| | | LB_TH:PL_TH | 0.107 | 0.084 | 2.73 | ***0.005** |
| | | LB_TH:SB_TH | 0.135 | 0.078 | 4.06 | ***0.001** |
| | | PI_TH:PL_TH | 0.111 | 0.078 | 3.03 | ***0.004** |
| | | PI_TH:SB_TH | 0.089 | 0.070 | 2.64 | ***0.006** |
| | | PL_TH:SB_TH | 0.052 | 0.075 | 0.07 | 0.483 |
| | Úlfljótsvatn | LB_UL:PI_UL | 0.107 | 0.182 | -0.59 | 0.720 |
| | | LB_UL:PL_UL | 0.093 | 0.131 | 0.23 | 0.408 |
| | | PI_UL:PL_UL | 0.138 | 0.192 | 0.15 | 0.439 |
| | Across Þingvallavatn and Úlfljótsvatn | LB_TH:LB_UL | 0.109 | 0.098 | 2.10 | 0.016 |
| | | LB_TH:PI_UL | 0.135 | 0.176 | 0.55 | 0.284 |

| | | | | | | |
|---|---|---|---|---|---|---|
| | | LB_TH:PL_UL | 0.095 | 0.119 | 0.65 | 0.242 |
| | | LB_UL:PI_TH | 0.096 | 0.098 | 1.66 | 0.055 |
| | | LB_UL:PL_TH | 0.069 | 0.094 | 0.21 | 0.416 |
| | | LB_UL:SB_TH | 0.069 | 0.092 | 0.40 | 0.346 |
| | | PI_TH:PI_UL | 0.119 | 0.171 | 0.13 | 0.458 |
| | | PI_TH:PL_UL | 0.146 | 0.116 | 2.66 | ***0.006** |
| | | PI_UL:PL_TH | 0.106 | 0.175 | -0.43 | 0.667 |
| | | PI_UL:SB_TH | 0.102 | 0.170 | -0.46 | 0.679 |
| | | PL_TH:PL_UL | 0.088 | 0.120 | 0.32 | 0.371 |
| | | PL_UL:SB_TH | 0.118 | 0.113 | 1.78 | 0.034 |
| Articular-angular[2] | AN to lake morphs | AN:Hest | 0.052 | 0.025 | 4.29 | ***0.001** |
| | | AN:Hit | 0.038 | 0.044 | 1.03 | 0.154 |
| | | AN:LB_TH | 0.115 | 0.022 | 7.53 | ***0.001** |
| | | AN:LB_UL | 0.065 | 0.027 | 5.04 | ***0.001** |
| | | AN:PI_TH | 0.051 | 0.023 | 4.78 | ***0.001** |
| | | AN:PI_UL | 0.057 | 0.037 | 3.27 | ***0.002** |
| | | AN:PL_TH | 0.045 | 0.029 | 3.33 | ***0.002** |
| | | AN:PL_UL | 0.063 | 0.032 | 4.57 | ***0.001** |
| | | AN:SB_TH | 0.070 | 0.031 | 5.11 | ***0.001** |
| | Hestvatn and Hítarvatn | Hest:Hit | 0.058 | 0.046 | 2.48 | ***0.006** |
| | | Hest:LB_TH | 0.096 | 0.027 | 6.21 | ***0.001** |
| | | Hest:LB_UL | 0.048 | 0.031 | 3.27 | ***0.002** |
| | | Hest:PI_TH | 0.041 | 0.026 | 3.18 | ***0.001** |
| | | Hest:PI_UL | 0.035 | 0.042 | 1.06 | 0.137 |
| | | Hest:PL_TH | 0.053 | 0.030 | 3.99 | ***0.001** |
| | | Hest:PL_UL | 0.067 | 0.033 | 4.38 | ***0.001** |
| | | Hest:SB_TH | 0.067 | 0.030 | 5.04 | ***0.001** |
| | | Hit:LB_TH | 0.100 | 0.046 | 4.89 | ***0.001** |
| | | Hit:LB_UL | 0.062 | 0.048 | 2.69 | ***0.005** |
| | | Hit:PI_TH | 0.049 | 0.045 | 1.92 | 0.024 |
| | | Hit:PI_UL | 0.065 | 0.055 | 2.31 | 0.012 |
| | | Hit:PL_TH | 0.045 | 0.047 | 1.53 | 0.064 |
| | | Hit:PL_UL | 0.061 | 0.048 | 2.58 | ***0.005** |
| | | Hit:SB_TH | 0.064 | 0.047 | 2.82 | ***0.003** |
| | Þingvallavatn | LB_TH:PI_TH | 0.089 | 0.024 | 7.01 | ***0.001** |
| | | LB_TH:PL_TH | 0.107 | 0.031 | 6.74 | ***0.001** |
| | | LB_TH:SB_TH | 0.062 | 0.032 | 4.54 | ***0.001** |
| | | PI_TH:PL_TH | 0.039 | 0.031 | 2.42 | ***0.005** |
| | | PI_TH:SB_TH | 0.060 | 0.033 | 3.98 | ***0.001** |
| | | PL_TH:SB_TH | 0.062 | 0.020 | 6.16 | ***0.001** |
| | Úlfljótsvatn | LB_UL:PI_UL | 0.031 | 0.041 | 0.55 | 0.296 |
| | | LB_UL:PL_UL | 0.032 | 0.035 | 1.27 | 0.116 |
| | | PI_UL:PL_UL | 0.052 | 0.047 | 1.99 | 0.025 |
| | Across Þingvallavatn and Úlfljótsvatn | LB_TH:LB_UL | 0.076 | 0.029 | 5.18 | ***0.001** |
| | | LB_TH:PI_UL | 0.092 | 0.039 | 4.52 | ***0.001** |
| | | LB_TH:PL_UL | 0.091 | 0.033 | 5.76 | ***0.001** |
| | | LB_UL:PI_TH | 0.045 | 0.029 | 3.23 | ***0.001** |
| | | LB_UL:PL_TH | 0.050 | 0.032 | 3.38 | ***0.002** |
| | | LB_UL:SB_TH | 0.046 | 0.034 | 2.78 | ***0.002** |
| | | PI_TH:PI_UL | 0.042 | 0.039 | 1.96 | 0.024 |
| | | PI_TH:PL_UL | 0.057 | 0.033 | 3.79 | ***0.001** |
| | | PI_UL:PL_TH | 0.052 | 0.045 | 2.16 | 0.016 |
| | | PI_UL:SB_TH | 0.066 | 0.046 | 2.93 | ***0.003** |
| | | PL_TH:PL_UL | 0.044 | 0.026 | 3.59 | ***0.001** |
| | | PL_UL:SB_TH | 0.048 | 0.025 | 4.30 | ***0.001** |
| Quadrate[2] | AN to lake morphs | AN:Hest | 0.037 | 0.027 | 2.77 | ***0.003** |
| | | AN:Hit | 0.050 | 0.047 | 1.87 | 0.034 |
| | | AN:LB_TH | 0.075 | 0.024 | 6.07 | ***0.001** |
| | | AN:LB_UL | 0.067 | 0.028 | 4.93 | ***0.001** |
| | | AN:PI_TH | 0.074 | 0.024 | 5.10 | ***0.001** |
| | | AN:PI_UL | 0.071 | 0.042 | 3.42 | ***0.001** |

| | | | | | | |
|---|---|---|---|---|---|---|
| | | AN:PL_TH | 0.055 | 0.029 | 3.99 | ***0.001** |
| | | AN:PL_UL | 0.047 | 0.033 | 3.05 | ***0.004** |
| | | AN:SB_TH | 0.053 | 0.031 | 3.55 | ***0.001** |
| | Hestvatn and Hítarvatn | Hest:Hit | 0.063 | 0.049 | 2.83 | ***0.002** |
| | | Hest:LB_TH | 0.059 | 0.028 | 4.69 | ***0.001** |
| | | Hest:LB_UL | 0.057 | 0.032 | 3.74 | ***0.001** |
| | | Hest:PI_TH | 0.058 | 0.028 | 4.46 | ***0.001** |
| | | Hest:PI_UL | 0.062 | 0.045 | 2.89 | ***0.003** |
| | | Hest:PL_TH | 0.045 | 0.030 | 2.90 | ***0.004** |
| | | Hest:PL_UL | 0.041 | 0.034 | 2.29 | 0.011 |
| | | Hest:SB_TH | 0.046 | 0.031 | 3.15 | ***0.001** |
| | | Hit:LB_TH | 0.097 | 0.049 | 4.49 | ***0.001** |
| | | Hit:LB_UL | 0.104 | 0.051 | 4.02 | ***0.001** |
| | | Hit:PI_TH | 0.103 | 0.049 | 4.66 | ***0.001** |
| | | Hit:PI_UL | 0.109 | 0.059 | 4.17 | ***0.001** |
| | | Hit:PL_TH | 0.083 | 0.049 | 3.54 | ***0.001** |
| | | Hit:PL_UL | 0.079 | 0.051 | 3.11 | ***0.002** |
| | | Hit:SB_TH | 0.084 | 0.051 | 3.33 | ***0.001** |
| | Þingvallavatn | LB_TH:PI_TH | 0.026 | 0.026 | 1.76 | 0.041 |
| | | LB_TH:PL_TH | 0.033 | 0.033 | 1.57 | 0.053 |
| | | LB_TH:SB_TH | 0.038 | 0.035 | 1.94 | 0.026 |
| | | PI_TH:PL_TH | 0.034 | 0.032 | 1.87 | 0.029 |
| | | PI_TH:SB_TH | 0.036 | 0.034 | 1.86 | 0.029 |
| | | PL_TH:SB_TH | 0.023 | 0.021 | 2.02 | 0.020 |
| | Úlfljótsvatn | LB_UL:PI_UL | 0.020 | 0.045 | -1.47 | 0.921 |
| | | LB_UL:PL_UL | 0.038 | 0.037 | 1.74 | 0.038 |
| | | PI_UL:PL_UL | 0.047 | 0.049 | 1.54 | 0.063 |
| | Across Þingvallavatn and Úlfljótsvatn | LB_TH:LB_UL | 0.046 | 0.031 | 3.17 | ***0.002** |
| | | LB_TH:PI_UL | 0.057 | 0.041 | 2.84 | ***0.005** |
| | | LB_TH:PL_UL | 0.038 | 0.036 | 1.81 | 0.040 |
| | | LB_UL:PI_TH | 0.035 | 0.030 | 2.25 | 0.015 |
| | | LB_UL:PL_TH | 0.048 | 0.033 | 2.89 | ***0.003** |
| | | LB_UL:SB_TH | 0.042 | 0.036 | 2.37 | 0.008 |
| | | PI_TH:PI_UL | 0.040 | 0.041 | 1.58 | 0.062 |
| | | PI_TH:PL_UL | 0.042 | 0.035 | 2.47 | ***0.007** |
| | | PI_UL:PL_TH | 0.054 | 0.047 | 2.21 | 0.018 |
| | | PI_UL:SB_TH | 0.048 | 0.048 | 1.68 | 0.052 |
| | | PL_TH:PL_UL | 0.024 | 0.028 | 1.12 | 0.136 |
| | | PL_UL:SB_TH | 0.029 | 0.027 | 1.96 | 0.024 |
| Premaxilla[1] | AN to lake morphs | AN:Hest | 0.111 | 0.110 | 1.70 | 0.045 |
| | | AN:Hit | 0.384 | 0.375 | 1.70 | 0.045 |
| | | AN:LB_TH | 0.108 | 0.114 | 1.30 | 0.096 |
| | | AN:LB_UL | 0.091 | 0.141 | -0.47 | 0.681 |
| | | AN:PI_TH | 0.124 | 0.096 | 3.05 | ***0.001** |
| | | AN:PI_UL | 0.171 | 0.238 | 0.05 | 0.479 |
| | | AN:PL_TH | 0.173 | 0.110 | 3.79 | ***0.001** |
| | | AN:PL_UL | 0.179 | 0.169 | 1.99 | 0.020 |
| | | AN:SB_TH | 0.211 | 0.096 | 5.60 | ***0.001** |
| | Hestvatn and Hítarvatn | Hest:Hit | 0.364 | 0.379 | 1.38 | 0.088 |
| | | Hest:LB_TH | 0.136 | 0.128 | 1.94 | 0.023 |
| | | Hest:LB_UL | 0.121 | 0.151 | 0.47 | 0.324 |
| | | Hest:PI_TH | 0.088 | 0.115 | 0.39 | 0.358 |
| | | Hest:PI_UL | 0.192 | 0.244 | 0.44 | 0.334 |
| | | Hest:PL_TH | 0.142 | 0.125 | 2.38 | 0.009 |
| | | Hest:PL_UL | 0.197 | 0.179 | 2.20 | 0.011 |
| | | Hest:SB_TH | 0.180 | 0.113 | 4.11 | ***0.001** |
| | | Hit:LB_TH | 0.383 | 0.378 | 1.66 | 0.044 |
| | | Hit:LB_UL | 0.373 | 0.394 | 1.40 | 0.078 |
| | | Hit:PI_TH | 0.400 | 0.380 | 1.88 | 0.022 |
| | | Hit:PI_UL | 0.395 | 0.437 | 1.21 | 0.119 |
| | | Hit:PL_TH | 0.358 | 0.386 | 1.35 | 0.095 |

| | | | | | | |
|---|---|---|---|---|---|---|
| | | Hit:PL_UL | 0.330 | 0.402 | 0.70 | 0.260 |
| | | Hit:SB_TH | 0.319 | 0.372 | 0.87 | 0.200 |
| | Þingvallavatn | LB_TH:PI_TH | 0.149 | 0.117 | 2.88 | ***0.003** |
| | | LB_TH:PL_TH | 0.160 | 0.129 | 2.66 | ***0.003** |
| | | LB_TH:SB_TH | 0.203 | 0.115 | 4.63 | ***0.001** |
| | | PI_TH:PL_TH | 0.142 | 0.112 | 2.89 | ***0.001** |
| | | PI_TH:SB_TH | 0.175 | 0.101 | 4.44 | ***0.001** |
| | | PL_TH:SB_TH | 0.086 | 0.109 | 0.48 | 0.315 |
| | Úlfljótsvatn | LB_UL:PI_UL | 0.164 | 0.255 | -0.54 | 0.718 |
| | | LB_UL:PL_UL | 0.163 | 0.202 | 0.59 | 0.283 |
| | | PI_UL:PL_UL | 0.221 | 0.276 | 0.50 | 0.322 |
| | Across Þingvallavatn and Úlfljótsvatn | LB_TH:LB_UL | 0.088 | 0.156 | -1.05 | 0.851 |
| | | LB_TH:PI_UL | 0.156 | 0.247 | -0.51 | 0.689 |
| | | LB_TH:PL_UL | 0.165 | 0.182 | 1.11 | 0.136 |
| | | LB_UL:PI_TH | 0.115 | 0.145 | 0.51 | 0.305 |
| | | LB_UL:PL_TH | 0.141 | 0.151 | 1.33 | 0.102 |
| | | LB_UL:SB_TH | 0.178 | 0.142 | 2.85 | ***0.002** |
| | | PI_TH:PI_UL | 0.169 | 0.238 | -0.03 | 0.521 |
| | | PI_TH:PL_UL | 0.210 | 0.173 | 2.68 | ***0.003** |
| | | PI_UL:PL_TH | 0.176 | 0.245 | 0.06 | 0.475 |
| | | PI_UL:SB_TH | 0.208 | 0.238 | 0.97 | 0.154 |
| | | PL_TH:PL_UL | 0.141 | 0.178 | 0.48 | 0.318 |
| | | PL_UL:SB_TH | 0.145 | 0.172 | 0.77 | 0.232 |
| Maxilla[2] | AN to lake morphs | AN:Hest | 0.036 | 0.021 | 3.92 | ***0.001** |
| | | AN:Hit | 0.058 | 0.037 | 3.07 | ***0.001** |
| | | AN:LB_TH | 0.052 | 0.019 | 5.25 | ***0.001** |
| | | AN:LB_UL | 0.038 | 0.023 | 3.27 | ***0.001** |
| | | AN:PI_TH | 0.039 | 0.019 | 4.38 | ***0.001** |
| | | AN:PI_UL | 0.037 | 0.032 | 2.19 | 0.016 |
| | | AN:PL_TH | 0.054 | 0.023 | 4.91 | ***0.001** |
| | | AN:PL_UL | 0.040 | 0.026 | 3.03 | ***0.001** |
| | | AN:SB_TH | 0.064 | 0.025 | 5.20 | ***0.001** |
| | Hestvatn and Hítarvatn | Hest:Hit | 0.065 | 0.039 | 3.62 | ***0.001** |
| | | Hest:LB_TH | 0.048 | 0.023 | 4.28 | ***0.001** |
| | | Hest:LB_UL | 0.055 | 0.026 | 4.30 | ***0.001** |
| | | Hest:PI_TH | 0.042 | 0.022 | 3.69 | ***0.001** |
| | | Hest:PI_UL | 0.034 | 0.034 | 1.64 | 0.049 |
| | | Hest:PL_TH | 0.037 | 0.024 | 3.27 | ***0.001** |
| | | Hest:PL_UL | 0.060 | 0.027 | 4.42 | ***0.001** |
| | | Hest:SB_TH | 0.045 | 0.025 | 3.91 | ***0.001** |
| | | Hit:LB_TH | 0.085 | 0.038 | 4.49 | ***0.001** |
| | | Hit:LB_UL | 0.085 | 0.040 | 4.60 | ***0.001** |
| | | Hit:PI_TH | 0.080 | 0.038 | 4.43 | ***0.001** |
| | | Hit:PI_UL | 0.080 | 0.047 | 3.52 | ***0.001** |
| | | Hit:PL_TH | 0.055 | 0.038 | 3.06 | ***0.001** |
| | | Hit:PL_UL | 0.076 | 0.041 | 3.82 | ***0.001** |
| | | Hit:SB_TH | 0.080 | 0.038 | 4.48 | ***0.001** |
| | Þingvallavatn | LB_TH:PI_TH | 0.024 | 0.020 | 2.17 | 0.017 |
| | | LB_TH:PL_TH | 0.055 | 0.025 | 4.67 | ***0.001** |
| | | LB_TH:SB_TH | 0.035 | 0.027 | 2.58 | ***0.002** |
| | | PI_TH:PL_TH | 0.052 | 0.026 | 4.28 | ***0.001** |
| | | PI_TH:SB_TH | 0.050 | 0.027 | 3.74 | ***0.001** |
| | | PL_TH:SB_TH | 0.052 | 0.017 | 5.72 | ***0.001** |
| | Úlfljótsvatn | LB_UL:PI_UL | 0.030 | 0.034 | 1.06 | 0.155 |
| | | LB_UL:PL_UL | 0.024 | 0.031 | 0.79 | 0.213 |
| | | PI_UL:PL_UL | 0.039 | 0.039 | 1.69 | 0.047 |
| | Across Þingvallavatn and Úlfljótsvatn | LB_TH:LB_UL | 0.051 | 0.024 | 4.22 | ***0.001** |
| | | LB_TH:PI_UL | 0.044 | 0.032 | 2.82 | ***0.003** |
| | | LB_TH:PL_UL | 0.056 | 0.028 | 4.08 | ***0.001** |
| | | LB_UL:PI_TH | 0.045 | 0.024 | 3.88 | ***0.001** |
| | | LB_UL:PL_TH | 0.079 | 0.027 | 5.80 | ***0.001** |

| | | | | | | |
|---|---|---|---|---|---|---|
| | | LB_UL:SB_TH | 0.066 | 0.028 | 4.47 | ***0.001** |
| | | PI_TH:PI_UL | 0.040 | 0.033 | 2.42 | ***0.003** |
| | | PI_TH:PL_UL | 0.052 | 0.029 | 3.69 | ***0.001** |
| | | PI_UL:PL_TH | 0.062 | 0.036 | 3.50 | ***0.001** |
| | | PI_UL:SB_TH | 0.052 | 0.039 | 2.78 | ***0.002** |
| | | PL_TH:PL_UL | 0.079 | 0.022 | 7.04 | ***0.001** |
| | | PL_UL:SB_TH | 0.068 | 0.021 | 5.76 | ***0.001** |
| Supramaxilla[2] | AN to lake morphs | AN:Hest | 0.041 | 0.042 | 1.59 | 0.063 |
| | | AN:Hit | 0.079 | 0.073 | 1.88 | 0.028 |
| | | AN:LB_TH | 0.055 | 0.037 | 3.14 | ***0.002** |
| | | AN:LB_UL | 0.060 | 0.045 | 2.64 | ***0.003** |
| | | AN:PI_TH | 0.040 | 0.037 | 1.83 | 0.029 |
| | | AN:PI_UL | 0.080 | 0.064 | 2.37 | 0.008 |
| | | AN:PL_TH | 0.057 | 0.049 | 2.17 | 0.019 |
| | | AN:PL_UL | 0.054 | 0.052 | 1.73 | 0.039 |
| | | AN:SB_TH | 0.117 | 0.049 | 5.00 | ***0.001** |
| | Hestvatn and Hítarvatn | Hest:Hit | 0.114 | 0.075 | 3.22 | ***0.001** |
| | | Hest:LB_TH | 0.059 | 0.043 | 2.61 | ***0.007** |
| | | Hest:LB_UL | 0.027 | 0.050 | -0.16 | 0.573 |
| | | Hest:PI_TH | 0.050 | 0.043 | 2.11 | 0.020 |
| | | Hest:PI_UL | 0.055 | 0.068 | 1.00 | 0.155 |
| | | Hest:PL_TH | 0.087 | 0.047 | 3.83 | ***0.001** |
| | | Hest:PL_UL | 0.064 | 0.052 | 2.42 | ***0.005** |
| | | Hest:SB_TH | 0.136 | 0.047 | 5.61 | ***0.001** |
| | | Hit:LB_TH | 0.107 | 0.072 | 3.18 | ***0.001** |
| | | Hit:LB_UL | 0.130 | 0.079 | 3.53 | ***0.001** |
| | | Hit:PI_TH | 0.092 | 0.074 | 2.35 | 0.008 |
| | | Hit:PI_UL | 0.149 | 0.090 | 3.48 | ***0.001** |
| | | Hit:PL_TH | 0.055 | 0.075 | 0.69 | 0.249 |
| | | Hit:PL_UL | 0.101 | 0.080 | 2.46 | ***0.006** |
| | | Hit:SB_TH | 0.121 | 0.074 | 3.46 | ***0.001** |
| | Þingvallavatn | LB_TH:PI_TH | 0.022 | 0.039 | -0.12 | 0.545 |
| | | LB_TH:PL_TH | 0.069 | 0.051 | 2.69 | ***0.004** |
| | | LB_TH:SB_TH | 0.087 | 0.052 | 3.49 | ***0.001** |
| | | PI_TH:PL_TH | 0.057 | 0.052 | 1.92 | 0.026 |
| | | PI_TH:SB_TH | 0.090 | 0.052 | 3.72 | ***0.001** |
| | | PL_TH:SB_TH | 0.080 | 0.034 | 4.73 | ***0.001** |
| | Úlfljótsvatn | LB_UL:PI_UL | 0.037 | 0.073 | -0.18 | 0.551 |
| | | LB_UL:PL_UL | 0.069 | 0.056 | 2.38 | ***0.007** |
| | | PI_UL:PL_UL | 0.063 | 0.075 | 1.09 | 0.148 |
| | Across Þingvallavatn and Úlfljótsvatn | LB_TH:LB_UL | 0.059 | 0.047 | 2.43 | ***0.006** |
| | | LB_TH:PI_UL | 0.053 | 0.065 | 1.03 | 0.147 |
| | | LB_TH:PL_UL | 0.031 | 0.054 | -0.10 | 0.545 |
| | | LB_UL:PI_TH | 0.058 | 0.047 | 2.30 | 0.008 |
| | | LB_UL:PL_TH | 0.103 | 0.051 | 4.13 | ***0.001** |
| | | LB_UL:SB_TH | 0.143 | 0.052 | 6.07 | ***0.001** |
| | | PI_TH:PI_UL | 0.064 | 0.064 | 1.65 | 0.051 |
| | | PI_TH:PL_UL | 0.036 | 0.055 | 0.31 | 0.394 |
| | | PI_UL:PL_TH | 0.113 | 0.074 | 3.03 | ***0.001** |
| | | PI_UL:SB_TH | 0.133 | 0.075 | 3.63 | ***0.001** |
| | | PL_TH:PL_UL | 0.057 | 0.044 | 2.54 | ***0.004** |
| | | PL_UL:SB_TH | 0.087 | 0.042 | 4.45 | ***0.001** |

1 For Dentary and Premaxilla the full model (shape = CSbone x population) had the best fit.

2 For the Articular-angular, Quadrate, Maxilla and Supramaxilla the reduced model (shape = CSbone + population) had the best fit

D = pairwise differences between vector lengths; UCL = upper confidence level; Z = Z-statistic; P = P-value, ≤ 0.008 in bold with asterisk.

***Supplementary material 21***: *Results from pairwise comparisons on population differences in mean bone shape for the dentary and premaxilla, based on the reduced model (shape = $CS_{bone}$ + population) assuming no difference in allometry by population. AN-charr treated as five separate populations. Used to make Figure 4D and Figure 4C.*

| Bone | System | Pairs | D | UCL | Z | P-value |
|---|---|---|---|---|---|---|
| Dentary | AN populations | AN_Brúará:AN_Gufudalsá | 0.049 | 0.057 | 1.06 | 0.139 |
| | | AN_Brúará:AN_Lambeyrarkvísl | 0.050 | 0.063 | 0.94 | 0.168 |
| | | AN_Brúará:AN_Víðidalsá | 0.064 | 0.059 | 1.95 | 0.025 |
| | | AN_Brúará:AN_Vífilsdalsá | 0.065 | 0.065 | 1.68 | 0.050 |
| | | AN_Gufudalsá:AN_Lambeyrarkvísl | 0.046 | 0.045 | 1.78 | 0.036 |
| | | AN_Gufudalsá:AN_Víðidalsá | 0.049 | 0.041 | 2.33 | 0.010 |
| | | AN_Gufudalsá:AN_Vífilsdalsá | 0.057 | 0.050 | 2.09 | 0.012 |
| | | AN_Lambeyrarkvísl:AN_Víðidalsá | 0.064 | 0.045 | 2.77 | ***0.004** |
| | | AN_Lambeyrarkvísl:AN_Vífilsdalsá | 0.031 | 0.052 | -0.26 | 0.611 |
| | | AN_Víðidalsá:AN_Vífilsdalsá | 0.079 | 0.049 | 3.23 | ***0.001** |
| | Brúará to lake morphs | AN_Brúará:Hest | 0.056 | 0.055 | 1.77 | 0.043 |
| | | AN_Brúará:Hit | 0.073 | 0.068 | 1.91 | 0.032 |
| | | AN_Brúará:LB_TH | 0.092 | 0.052 | 3.53 | ***0.001** |
| | | AN_Brúará:LB_UL | 0.043 | 0.057 | 0.77 | 0.220 |
| | | AN_Brúará:PI_TH | 0.036 | 0.052 | 0.26 | 0.403 |
| | | AN_Brúará:PI_UL | 0.041 | 0.063 | 0.15 | 0.452 |
| | | AN_Brúará:PL_TH | 0.040 | 0.060 | 0.41 | 0.340 |
| | | AN_Brúará:PL_UL | 0.049 | 0.060 | 1.02 | 0.153 |
| | | AN_Brúará:SB_TH | 0.077 | 0.060 | 2.47 | ***0.006** |
| | Gufudalsá to lake morphs | AN_Gufudalsá:Hest | 0.048 | 0.035 | 2.70 | ***0.004** |
| | | AN_Gufudalsá:Hit | 0.079 | 0.052 | 3.19 | ***0.001** |
| | | AN_Gufudalsá:LB_TH | 0.116 | 0.033 | 5.88 | ***0.001** |
| | | AN_Gufudalsá:LB_UL | 0.064 | 0.036 | 3.62 | ***0.001** |
| | | AN_Gufudalsá:PI_TH | 0.054 | 0.033 | 3.28 | ***0.001** |
| | | AN_Gufudalsá:PI_UL | 0.059 | 0.048 | 2.41 | 0.012 |
| | | AN_Gufudalsá:PL_TH | 0.054 | 0.037 | 3.00 | ***0.003** |
| | | AN_Gufudalsá:PL_UL | 0.065 | 0.039 | 3.34 | ***0.001** |
| | | AN_Gufudalsá:SB_TH | 0.105 | 0.038 | 4.44 | ***0.001** |
| | Lambeyrarkvísl to lake morphs | AN_Lambeyrarkvísl:Hest | 0.073 | 0.042 | 3.52 | ***0.001** |
| | | AN_Lambeyrarkvísl:Hit | 0.076 | 0.057 | 2.70 | ***0.004** |
| | | AN_Lambeyrarkvísl:LB_TH | 0.117 | 0.039 | 5.44 | ***0.001** |
| | | AN_Lambeyrarkvísl:LB_UL | 0.060 | 0.042 | 2.90 | ***0.002** |
| | | AN_Lambeyrarkvísl:PI_TH | 0.056 | 0.039 | 2.89 | ***0.002** |
| | | AN_Lambeyrarkvísl:PI_UL | 0.060 | 0.050 | 2.27 | 0.013 |
| | | AN_Lambeyrarkvísl:PL_TH | 0.055 | 0.044 | 2.40 | 0.008 |
| | | AN_Lambeyrarkvísl:PL_UL | 0.056 | 0.047 | 2.24 | ***0.005** |
| | | AN_Lambeyrarkvísl:SB_TH | 0.098 | 0.048 | 4.08 | ***0.001** |
| | Víðidalsá to lake morphs | AN_Víðidalsá:Hest | 0.040 | 0.038 | 1.83 | 0.034 |
| | | AN_Víðidalsá:Hit | 0.065 | 0.056 | 2.15 | 0.015 |
| | | AN_Víðidalsá:LB_TH | 0.140 | 0.035 | 5.97 | ***0.001** |
| | | AN_Víðidalsá:LB_UL | 0.079 | 0.041 | 3.85 | ***0.001** |
| | | AN_Víðidalsá:PI_TH | 0.062 | 0.035 | 3.67 | ***0.001** |
| | | AN_Víðidalsá:PI_UL | 0.063 | 0.047 | 2.69 | ***0.004** |
| | | AN_Víðidalsá:PL_TH | 0.052 | 0.046 | 2.10 | 0.019 |
| | | AN_Víðidalsá:PL_UL | 0.082 | 0.047 | 3.32 | ***0.001** |
| | | AN_Víðidalsá:SB_TH | 0.118 | 0.050 | 4.74 | ***0.001** |
| | Vífisdalsá to lake morphs | AN_Vífilsdalsá:Hest | 0.089 | 0.045 | 4.02 | ***0.001** |
| | | AN_Vífilsdalsá:Hit | 0.093 | 0.060 | 3.02 | ***0.001** |
| | | AN_Vífilsdalsá:LB_TH | 0.119 | 0.044 | 4.76 | ***0.001** |
| | | AN_Vífilsdalsá:LB_UL | 0.076 | 0.047 | 3.01 | ***0.001** |
| | | AN_Vífilsdalsá:PI_TH | 0.072 | 0.044 | 3.10 | ***0.001** |
| | | AN_Vífilsdalsá:PI_UL | 0.076 | 0.054 | 2.70 | ***0.003** |
| | | AN_Vífilsdalsá:PL_TH | 0.074 | 0.048 | 3.01 | ***0.002** |
| | | AN_Vífilsdalsá:PL_UL | 0.069 | 0.052 | 2.53 | ***0.006** |

| | | | | | | |
|---|---|---|---|---|---|---|
| | | AN_Vífilsdalsá:SB_TH | 0.106 | 0.051 | 3.99 | ***0.001** |
| | Hestvatn and Hítarvatn | Hest:Hit | 0.065 | 0.049 | 2.68 | ***0.003** |
| | | Hest:LB_TH | 0.122 | 0.028 | 5.52 | ***0.001** |
| | | Hest:LB_UL | 0.068 | 0.032 | 3.98 | ***0.001** |
| | | Hest:PI_TH | 0.045 | 0.028 | 3.52 | ***0.001** |
| | | Hest:PI_UL | 0.053 | 0.043 | 2.35 | 0.011 |
| | | Hest:PL_TH | 0.043 | 0.032 | 2.52 | ***0.007** |
| | | Hest:PL_UL | 0.075 | 0.036 | 3.90 | ***0.001** |
| | | Hest:SB_TH | 0.106 | 0.035 | 4.93 | ***0.001** |
| | | Hit:LB_TH | 0.127 | 0.049 | 5.26 | ***0.001** |
| | | Hit:LB_UL | 0.088 | 0.049 | 3.85 | ***0.001** |
| | | Hit:PI_TH | 0.070 | 0.048 | 2.95 | ***0.002** |
| | | Hit:PI_UL | 0.084 | 0.057 | 3.14 | ***0.002** |
| | | Hit:PL_TH | 0.054 | 0.049 | 2.01 | 0.020 |
| | | Hit:PL_UL | 0.093 | 0.051 | 3.91 | ***0.001** |
| | | Hit:SB_TH | 0.094 | 0.051 | 3.67 | ***0.001** |
| | Þingvallavatn | LB_TH:PI_TH | 0.093 | 0.025 | 5.30 | ***0.001** |
| | | LB_TH:PL_TH | 0.111 | 0.032 | 5.22 | ***0.001** |
| | | LB_TH:SB_TH | 0.047 | 0.036 | 2.63 | ***0.002** |
| | | PI_TH:PL_TH | 0.029 | 0.034 | 1.08 | 0.145 |
| | | PI_TH:SB_TH | 0.079 | 0.037 | 3.98 | ***0.001** |
| | | PL_TH:SB_TH | 0.089 | 0.021 | 6.42 | ***0.001** |
| | Úlfljótsvatn | LB_UL:PI_UL | 0.028 | 0.043 | -0.01 | 0.487 |
| | | LB_UL:PL_UL | 0.026 | 0.036 | 0.37 | 0.356 |
| | | PI_UL:PL_UL | 0.036 | 0.052 | 0.45 | 0.318 |
| | Across Þingvallavatn and Úlfljótsvatn | LB_TH:LB_UL | 0.080 | 0.030 | 4.78 | ***0.001** |
| | | LB_TH:PI_UL | 0.097 | 0.041 | 4.46 | ***0.001** |
| | | LB_TH:PL_UL | 0.085 | 0.036 | 4.42 | ***0.001** |
| | | LB_UL:PI_TH | 0.037 | 0.030 | 2.32 | 0.010 |
| | | LB_UL:PL_TH | 0.057 | 0.033 | 3.68 | ***0.001** |
| | | LB_UL:SB_TH | 0.071 | 0.036 | 4.02 | ***0.001** |
| | | PI_TH:PI_UL | 0.029 | 0.041 | 0.48 | 0.321 |
| | | PI_TH:PL_UL | 0.047 | 0.037 | 2.50 | ***0.006** |
| | | PI_UL:PL_TH | 0.047 | 0.046 | 1.64 | 0.044 |
| | | PI_UL:SB_TH | 0.086 | 0.051 | 3.37 | ***0.001** |
| | | PL_TH:PL_UL | 0.065 | 0.028 | 4.34 | ***0.001** |
| | | PL_UL:SB_TH | 0.075 | 0.027 | 4.83 | ***0.001** |
| Premaxilla | AN populations | AN_Brúará:AN_Gufudalsá | 0.047 | 0.082 | -0.68 | 0.737 |
| | | AN_Brúará:AN_Lambeyrarkvísl | 0.059 | 0.086 | 0.03 | 0.500 |
| | | AN_Brúará:AN_Víðidalsá | 0.092 | 0.083 | 2.09 | 0.024 |
| | | AN_Brúará:AN_Vífilsdalsá | 0.066 | 0.090 | 0.26 | 0.398 |
| | | AN_Gufudalsá:AN_Lambeyrarkvísl | 0.047 | 0.065 | 0.21 | 0.415 |
| | | AN_Gufudalsá:AN_Víðidalsá | 0.079 | 0.059 | 3.00 | ***0.003** |
| | | AN_Gufudalsá:AN_Vífilsdalsá | 0.051 | 0.070 | 0.36 | 0.367 |
| | | AN_Lambeyrarkvísl:AN_Víðidalsá | 0.067 | 0.067 | 1.71 | 0.044 |
| | | AN_Lambeyrarkvísl:AN_Vífilsdalsá | 0.052 | 0.076 | 0.06 | 0.463 |
| | | AN_Víðidalsá:AN_Vífilsdalsá | 0.067 | 0.071 | 1.36 | 0.087 |
| | Brúará to lake morphs | AN_Brúará:Hest | 0.101 | 0.078 | 2.64 | ***0.006** |
| | | AN_Brúará:Hit | 0.086 | 0.095 | 1.17 | 0.114 |
| | | AN_Brúará:LB_TH | 0.103 | 0.075 | 3.03 | ***0.002** |
| | | AN_Brúará:LB_UL | 0.056 | 0.078 | 0.16 | 0.446 |
| | | AN_Brúará:PI_TH | 0.071 | 0.076 | 1.34 | 0.079 |
| | | AN_Brúará:PI_UL | 0.050 | 0.088 | -0.84 | 0.796 |
| | | AN_Brúará:PL_TH | 0.067 | 0.083 | 0.75 | 0.232 |
| | | AN_Brúará:PL_UL | 0.053 | 0.083 | -0.28 | 0.591 |
| | | AN_Brúará:SB_TH | 0.077 | 0.082 | 1.36 | 0.087 |
| | Gufudalsá to lake morphs | AN_Gufudalsá:Hest | 0.094 | 0.049 | 4.34 | ***0.001** |
| | | AN_Gufudalsá:Hit | 0.064 | 0.074 | 1.05 | 0.149 |
| | | AN_Gufudalsá:LB_TH | 0.116 | 0.049 | 5.48 | ***0.001** |
| | | AN_Gufudalsá:LB_UL | 0.072 | 0.053 | 3.15 | ***0.001** |
| | | AN_Gufudalsá:PI_TH | 0.077 | 0.047 | 3.81 | ***0.001** |

| | | | | | | |
|---|---|---|---|---|---|---|
| | | AN_Gufudalsá:PI_UL | 0.056 | 0.067 | 0.90 | 0.186 |
| | | AN_Gufudalsá:PL_TH | 0.068 | 0.051 | 2.97 | ***0.002** |
| | | AN_Gufudalsá:PL_UL | 0.081 | 0.052 | 3.66 | ***0.003** |
| | | AN_Gufudalsá:SB_TH | 0.094 | 0.050 | 4.59 | ***0.001** |
| | Lambeyrarkvísl to lake morphs | AN_Lambeyrarkvísl:Hest | 0.082 | 0.059 | 3.09 | **0.002** |
| | | AN_Lambeyrarkvísl:Hit | 0.076 | 0.084 | 1.29 | 0.104 |
| | | AN_Lambeyrarkvísl:LB_TH | 0.127 | 0.057 | 5.16 | ***0.001** |
| | | AN_Lambeyrarkvísl:LB_UL | 0.095 | 0.061 | 3.97 | ***0.001** |
| | | AN_Lambeyrarkvísl:PI_TH | 0.065 | 0.058 | 2.23 | 0.017 |
| | | AN_Lambeyrarkvísl:PI_UL | 0.077 | 0.076 | 1.75 | 0.044 |
| | | AN_Lambeyrarkvísl:PL_TH | 0.068 | 0.064 | 1.83 | 0.034 |
| | | AN_Lambeyrarkvísl:PL_UL | 0.091 | 0.065 | 3.25 | ***0.001** |
| | | AN_Lambeyrarkvísl:SB_TH | 0.104 | 0.065 | 3.89 | ***0.001** |
| | Víðidalsá to lake morphs | AN_Víðidalsá:Hest | 0.083 | 0.054 | 3.71 | ***0.001** |
| | | AN_Víðidalsá:Hit | 0.087 | 0.077 | 2.07 | 0.022 |
| | | AN_Víðidalsá:LB_TH | 0.160 | 0.050 | 7.03 | ***0.001** |
| | | AN_Víðidalsá:LB_UL | 0.128 | 0.054 | 6.30 | ***0.001** |
| | | AN_Víðidalsá:PI_TH | 0.090 | 0.050 | 4.28 | ***0.001** |
| | | AN_Víðidalsá:PI_UL | 0.099 | 0.068 | 3.29 | ***0.002** |
| | | AN_Víðidalsá:PL_TH | 0.080 | 0.063 | 2.72 | ***0.003** |
| | | AN_Víðidalsá:PL_UL | 0.123 | 0.063 | 5.05 | ***0.001** |
| | | AN_Víðidalsá:SB_TH | 0.135 | 0.062 | 5.38 | ***0.001** |
| | Vífisdalsá to lake morphs | AN_Vífilsdalsá:Hest | 0.094 | 0.066 | 3.17 | ***0.001** |
| | | AN_Vífilsdalsá:Hit | 0.084 | 0.086 | 1.56 | 0.056 |
| | | AN_Vífilsdalsá:LB_TH | 0.130 | 0.061 | 4.64 | ***0.001** |
| | | AN_Vífilsdalsá:LB_UL | 0.091 | 0.066 | 3.13 | ***0.002** |
| | | AN_Vífilsdalsá:PI_TH | 0.085 | 0.061 | 3.01 | ***0.003** |
| | | AN_Vífilsdalsá:PI_UL | 0.075 | 0.077 | 1.50 | 0.066 |
| | | AN_Vífilsdalsá:PL_TH | 0.080 | 0.072 | 2.13 | 0.022 |
| | | AN_Vífilsdalsá:PL_UL | 0.091 | 0.071 | 2.70 | ***0.005** |
| | | AN_Vífilsdalsá:SB_TH | 0.102 | 0.071 | 3.09 | ***0.002** |
| | Hestvatn and Hítarvatn | Hest:Hit | 0.085 | 0.071 | 2.49 | ***0.007** |
| | | Hest:LB_TH | 0.159 | 0.040 | 7.18 | ***0.001** |
| | | Hest:LB_UL | 0.132 | 0.045 | 6.63 | ***0.001** |
| | | Hest:PI_TH | 0.075 | 0.040 | 4.63 | ***0.001** |
| | | Hest:PI_UL | 0.098 | 0.063 | 3.56 | ***0.001** |
| | | Hest:PL_TH | 0.075 | 0.045 | 4.18 | ***0.001** |
| | | Hest:PL_UL | 0.135 | 0.047 | 6.53 | ***0.001** |
| | | Hest:SB_TH | 0.145 | 0.043 | 7.22 | ***0.001** |
| | | Hit:LB_TH | 0.155 | 0.071 | 4.73 | ***0.001** |
| | | Hit:LB_UL | 0.117 | 0.074 | 3.44 | ***0.001** |
| | | Hit:PI_TH | 0.101 | 0.068 | 3.40 | ***0.001** |
| | | Hit:PI_UL | 0.094 | 0.085 | 2.10 | 0.018 |
| | | Hit:PL_TH | 0.089 | 0.072 | 2.50 | 0.008 |
| | | Hit:PL_UL | 0.116 | 0.072 | 3.57 | ***0.001** |
| | | Hit:SB_TH | 0.134 | 0.071 | 4.13 | ***0.001** |
| | Þingvallavatn | LB_TH:PI_TH | 0.094 | 0.036 | 5.45 | ***0.001** |
| | | LB_TH:PL_TH | 0.098 | 0.049 | 5.25 | ***0.001** |
| | | LB_TH:SB_TH | 0.045 | 0.048 | 1.38 | 0.085 |
| | | PI_TH:PL_TH | 0.037 | 0.048 | 0.51 | 0.294 |
| | | PI_TH:SB_TH | 0.087 | 0.047 | 4.75 | ***0.001** |
| | | PL_TH:SB_TH | 0.088 | 0.030 | 6.12 | ***0.001** |
| | Úlfljótsvatn | LB_UL:PI_UL | 0.045 | 0.062 | 0.23 | 0.404 |
| | | LB_UL:PL_UL | 0.044 | 0.051 | 0.94 | 0.174 |
| | | PI_UL:PL_UL | 0.065 | 0.067 | 1.44 | 0.068 |
| | Across Þingvallavatn and Úlfljótsvatn | LB_TH:LB_UL | 0.065 | 0.042 | 3.56 | ***0.001** |
| | | LB_TH:PI_UL | 0.091 | 0.059 | 3.45 | ***0.001** |
| | | LB_TH:PL_UL | 0.078 | 0.049 | 3.76 | ***0.001** |
| | | LB_UL:PI_TH | 0.085 | 0.043 | 4.74 | ***0.001** |
| | | LB_UL:PL_TH | 0.084 | 0.050 | 3.83 | ***0.001** |
| | | LB_UL:SB_TH | 0.048 | 0.049 | 1.52 | 0.069 |

| | | | | | | |
|---|---|---|---|---|---|---|
| | | PI_TH:PI_UL | 0.064 | 0.060 | 1.92 | 0.027 |
| | | PI_TH:PL_UL | 0.086 | 0.048 | 4.27 | ***0.001** |
| | | PI_UL:PL_TH | 0.066 | 0.066 | 1.50 | 0.059 |
| | | PI_UL:SB_TH | 0.073 | 0.067 | 2.01 | 0.020 |
| | | PL_TH:PL_UL | 0.089 | 0.039 | 5.66 | ***0.001** |
| | | PL_UL:SB_TH | 0.051 | 0.038 | 2.98 | ***0.003** |

D = pairwise differences between vector lengths; UCL = upper confidence level; Z = Z-statistic; P = P-value, ≤ 0.008 in bold with asterisk.

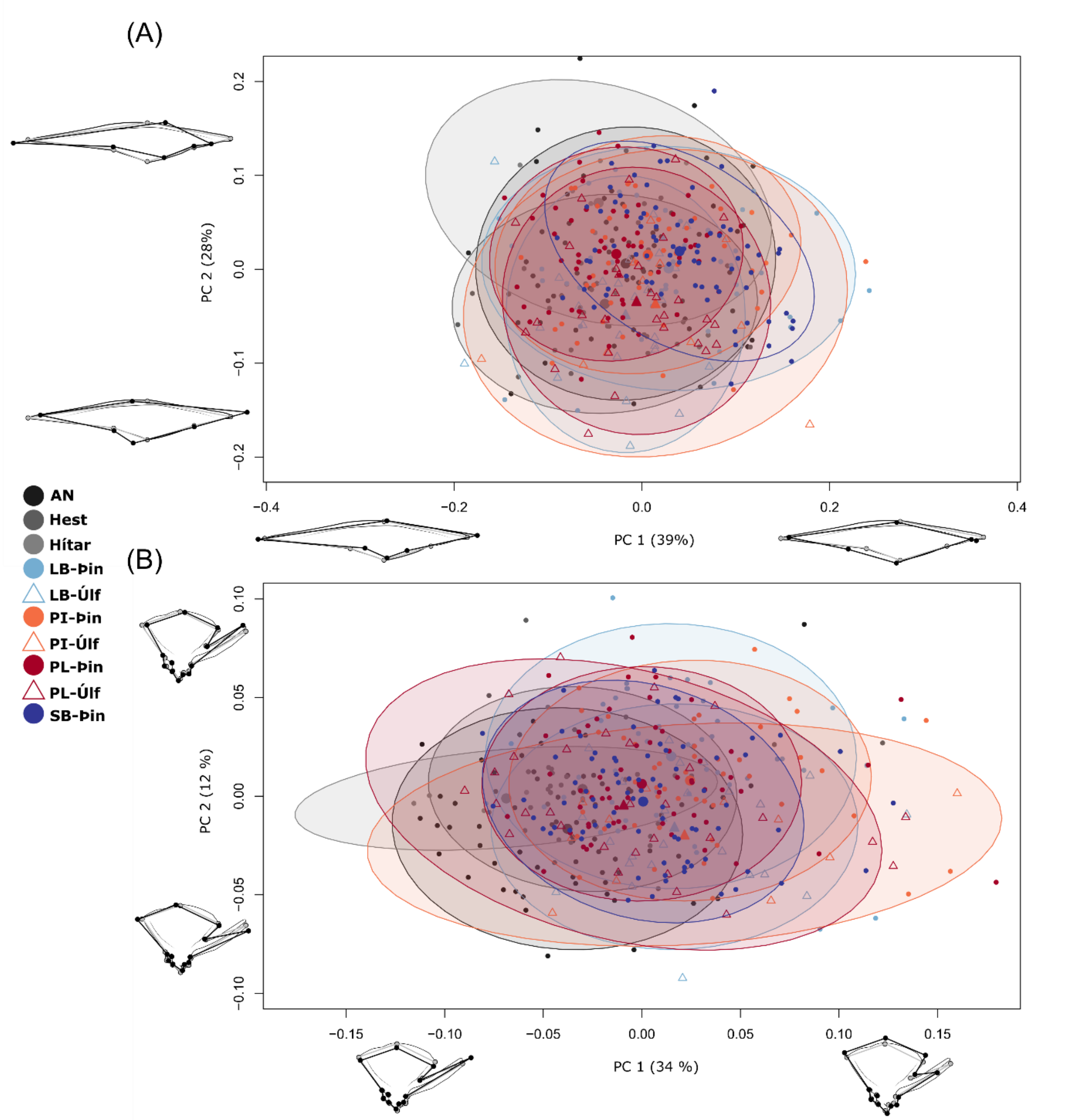


***Supplementary material 22**: PC-plot with deformation grids showing variation in shape for the A) supramaxilla and B) quadrate, size effects have been removed in both plots. None of the deformation grids are magnified, grey outline represents mean shape and black lines the two extremes for each PC. Each dot or triangle represents an individual and the ellipses 95% CI for the distribution by group (large dots or triangles represent the mean shape of each group in these dimensions). The percentage of total variation which each PC explains is written next to the PC.*

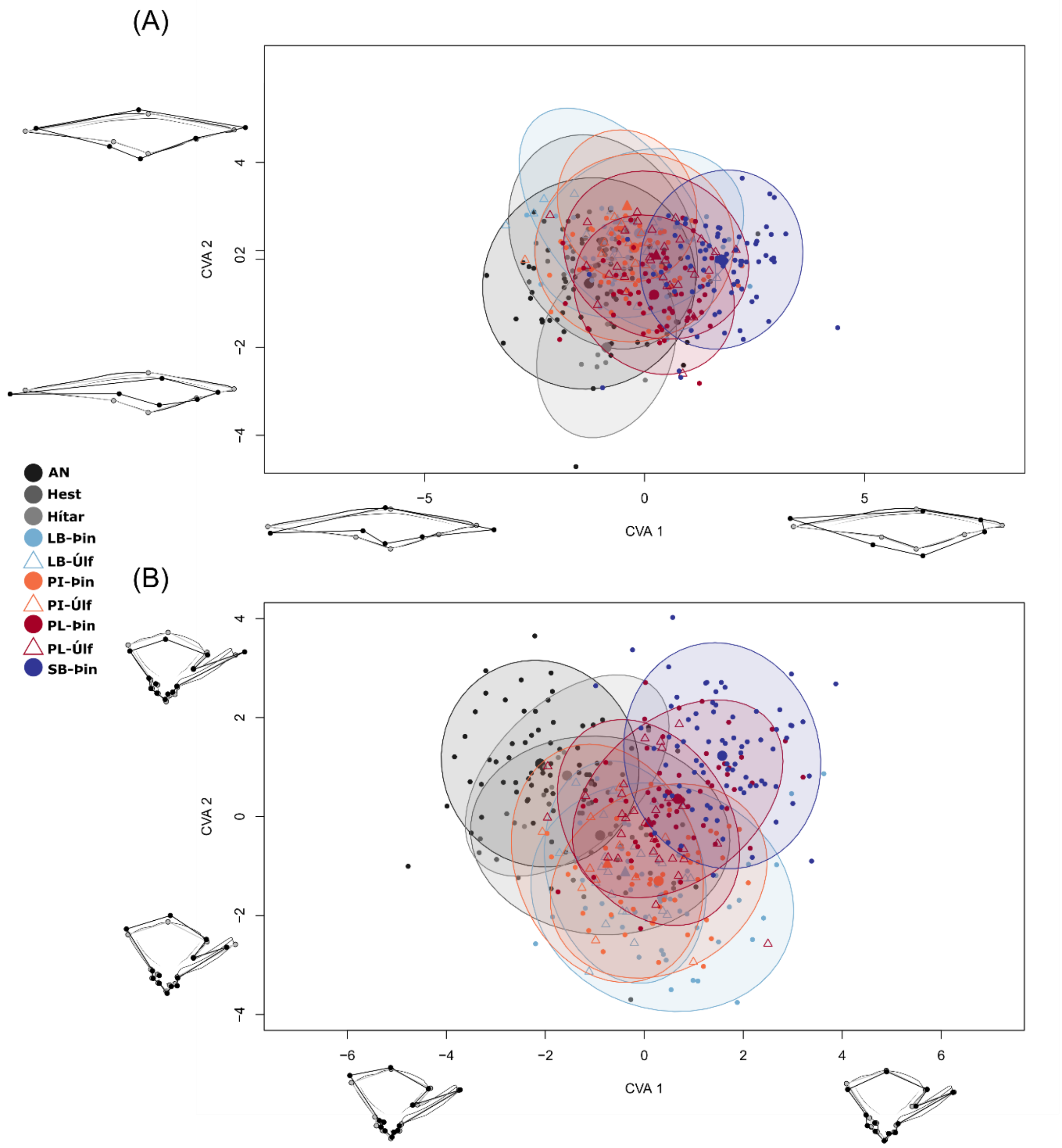


***Supplementary material 23****: CVA plot, with deformation grids showing variation in shape for the A) supramaxilla and B) quadrate. None of the deformation grids are magnified, grey outline represents mean shape and black lines the two extremes for each CVA. Each dot or triangle represents an individual and the ellipses 95% CI for the distribution by group (large dots or triangles represent the mean shape of each group in these dimensions).*

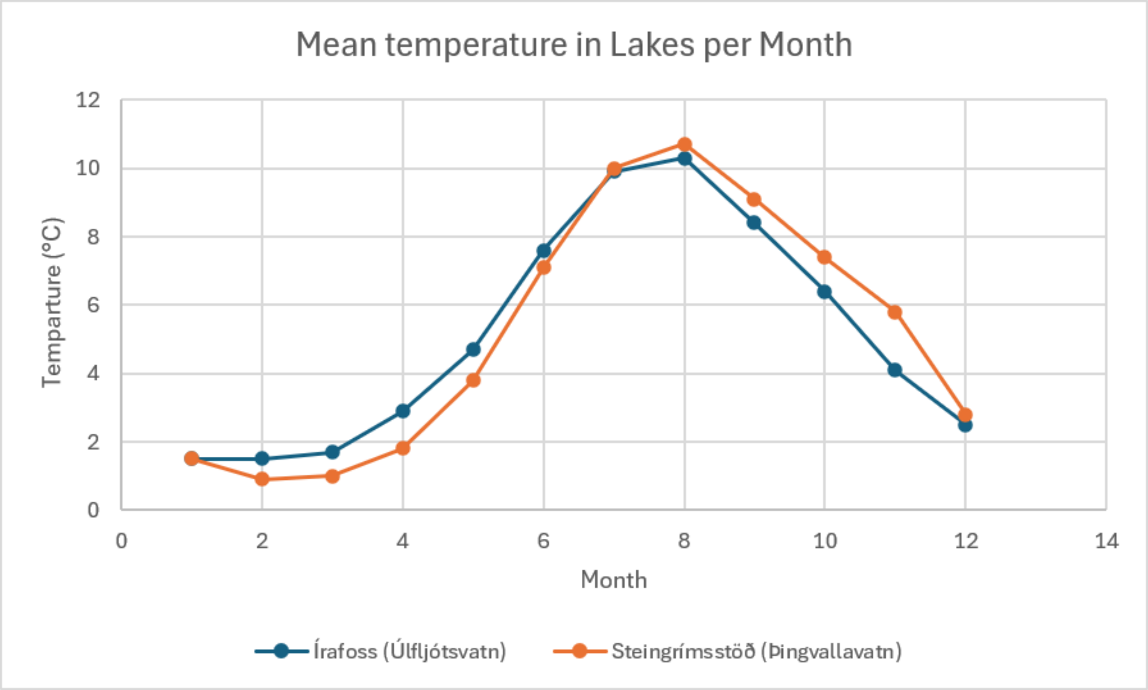


***Supplementary material 24****: Scatterplot showing average water temperature measurements per month in Þingvallavatn and Úlfljótsvatn. Benóný Jónsson at the Icelandic Marine & Freshwater Research Institute provided measurements from the years 2003 -2019 from Írafoss (station in the south of lake Úlfljótsvatn) and Steingrímsstöð (station in the south of lake Þingvallavatn).*
